# Phylogenomics of Cypriniformes, the most diverse order of freshwater fishes: consensus, challenges and limitations

**DOI:** 10.64898/2026.01.14.699467

**Authors:** Hiranya Sudasinghe, Ralf Britz, Michael Matschiner, Kevin Conway, Heok Hui Tan, Maurice Kottelat, Rajeev Raghavan, Neelesh Dahanukar, V.K. Anoop, Jörg Bohlen, Kerstin Howe, Richard Durbin, Walter Salzburger, Catherine L. Peichel, Lukas Rüber

## Abstract

Cypriniformes, the most species-rich order of freshwater fishes (∼5,000 species), represents a key lineage for understanding vertebrate diversification in freshwater ecosystems. This clade includes several highly miniaturized and understudied lineages whose phylogenetic placements have long remained contentious. Here, we present the first phylogenomic analysis of Cypriniformes with complete family-level representation and broad genus-level coverage, encompassing 316 species comprising approximately 30% of all described genera. Our dataset integrates 257 newly assembled genomes with publicly available resources and analyzes multiple sets of genome-wide markers using both concatenation-based and coalescent-aware approaches. The general concordance among analytical frameworks indicates that the backbone topology of Cypriniformes is now established, allowing clear identification between well-supported clades and regions of persistent conflict. Our results strengthen the evolutionary relationships of several miniaturized lineages, while identifying recalcitrant relationships shaped by both biological processes and model artefacts that can yield superficially similar patterns of phylogenetic conflict. Our study substantially expands genomic representation and establishes a phylogenomic foundation for future comparative, developmental, and evolutionary research in this freshwater radiation.

## Introduction

Understanding evolutionary relationships among organisms through phylogenetic inference is a central pursuit in biology. The testing of these phylogenetic hypotheses provides the foundation for recognizing patterns of organismal diversification and forms the basis for comparative biology (1–3). With the advent of high-throughput sequencing technologies, genome-scale datasets have revolutionized phylogenetic inference, giving rise to the field of phylogenomics, which is now considered the gold standard for testing molecular evolutionary hypotheses (4, 5).

With more than 35,000 described species, ray-finned fishes (Actinopterygii) are the most species-rich vertebrate clade, accounting for approximately half of all extant vertebrate species (6, 7). Remarkably, although freshwater habitats cover less than 1% of Earth’s surface, they are home to nearly 50% of all known fish species (8). The order Cypriniformes, comprising nearly 5,000 species, is the most species-rich clade of freshwater fishes (6, 7). Cypriniformes are distributed across a wide range of freshwater habitats in Asia, Europe, Africa, and North America and exhibit extraordinary morphological, physiological, and ecological adaptations (7, 9, 10). Cypriniformes contribute to our understanding of development, genetics, physiology and neuroscience through multiple model organisms (11–13) and benefit human-wellbeing through substantial economic, cultural and recreational fisheries, aquaculture and aquarium trade species (14, 15). Given this exceptional diversity and broad significance across multiple disciplines, a phylogenetic framework is essential for understanding the evolutionary history and systematics of Cypriniformes, in addition to informing comparative research.

Despite their scientific, economic, and cultural importance, the evolutionary history of Cypriniformes remains incompletely understood. Previous molecular phylogenetic studies have been limited by sparse taxon sampling, reliance on only a few genetic markers or targeted capture approaches, and incomplete representation of the order’s major lineages (e.g., 16–24). While some of these studies have recovered congruent relationships among certain lineages, they also showcase persistent phylogenetic conflict in others (25). The evolutionary placement of some of the most enigmatic lineages within Cypriniformes has remained uncertain typically owing to extreme morphological reduction, accelerated molecular evolution, or long-branch effects(26–28). A notable case is *Paedocypris*, a genus of progenetic miniature fishes comprising some of the smallest known vertebrates (29). Competing hypotheses place *Paedocypris* as the sister-group to either all other Cypriniformes or the suborder Cyprinoidei, or within the family Danionidae (e.g., 20, 21, 23, 26, 30). Thus, understanding the phylogenetic placement of highly miniaturized and morphologically derived taxa (*Paedocypris*, *Sundadanio*, *Fangfangia*, and *Danionella*) remains a challenge in vertebrate phylogenetics (24, 31). Phylogenetic inference in Cypriniformes is further complicated by the frequent occurrence of polyploid species (32–34). Improving phylogenetic resolution within Cypriniformes therefore requires not only broader genomic and taxonomic sampling, but also analytical strategies that can accommodate biological sources of incongruence such as incomplete lineage sorting (ILS), introgression, rate heterogeneity, and compositional bias as well as systematic sources, such as, model misspecification (2, 35–38).

In this study, we present the first phylogenomic study of Cypriniformes with a complete family-level representation and a broad taxonomic sampling across the order. Our dataset comprises more than 250 new genomes assembled from Illumina short reads as well as four new high-quality genome assemblies based on PacBio long reads. By integrating these new resources with publicly available cypriniform genomes, we construct and analyze multiple genome-scale datasets using different sets of genome-wide markers and phylogenetic methods. We then evaluate alternative phylogenetic hypotheses both among and within cypriniform lineages. By substantially expanding genomic resources for Cypriniformes, this study provides a phylogenomic framework that advances research in systematics, evolutionary biology, and comparative genomics, while placing important model organisms within an evolutionary framework.

## Materials and methods

### Taxon sampling

Previous phylogenetic investigations of Cypriniformes have been constrained by limited genetic marker sampling, relying primarily on mitochondrial DNA or a small number of nuclear loci, or genomic datasets based on anchored hybrid enrichment (AHE) with limited locus sampling (e.g., 17, 19–23). To resolve higher-level relationships within Cypriniformes, we therefore compiled a comprehensive phylogenomic dataset with a complete family-level coverage and extensive genus-level representation (Supplementary Fig. 1). The sampling design was established by downloading the classification for all Cypriniformes genera from the Catalog of Fishes(6) (accessed 01 December 2020), which was compared with the classification proposed by Tan & Armbruster (39). Based on this taxonomic framework and a review of the published literature, we designed a sampling scheme to represent all major lineages within Cypriniformes. We generated new genomic resources for 257 species, including Illumina short-read data for 253 species and long-read data for *Paedocypris* sp. “Kalimantan Tengah”(hereafter *Paedocypris* sp.), *Sundadanio atomus*, *Boraras brigittae*, and *Rasbora kalochroma* using the PacBio HiFi technology (Supplementary Table 1). This sampling extends the genomic coverage to lineages previously absent from all molecular databases, most notably the progenetic miniature species, *Fangfangia spinicleithralis*. Furthermore, whole-genome data are reported here for the first time for the cypriniform families Barbuccidae, Ellopostomatidae, Leptobarbidae, and Psilorhynchidae. In addition, our sampling includes several rare lineages that have been poorly represented or are entirely absent from previous phylogenetic studies, such as *Thryssocypris tonlesapensis, Pangio bhujia*, *Theriodes sandakanensis*, *Katibasia insidiosa*, *Plesiomyzon baotingensis*, *Protomyzon borneensis*, *Sundoreonectes sabanus*, *Eechathalakenda ophicephalus*, *Hypselobarbus periyarensis*, *Lepidopygopsis typus*, *Systomus binduchitra*, *Tariqilabeo periyarensis*, and *Barilius mesopotamicus,* thereby substantially broadening genomic representation across the order. Our sampling includes approximately 30% of the ∼527 described Cypriniformes genera, providing the broadest genus-level representation achieved to date for this order (Supplementary Fig. 1).

### DNA extraction, library preparation, and sequencing

Genomic DNA for Illumina short-read sequencing was extracted from museum-preserved specimens using the DNeasy Blood & Tissue Kit (Qiagen) according to the manufacturer’s protocol. The quantity, purity and length of the DNA was assessed using a Thermo Fisher Scientific Qubit 4.0 fluorometer. Sequencing libraries were made using Illumina TruSeq DNA PCR-free libraries starting with 1100ng. For gDNA less than 1000 ng in quantity, Illumina TruSeq nano DNA samples were generated using the Nano DNA High Throughput Library Prep Kit. The DNA libraries were evaluated using a Thermo Fisher Scientific Qubit 4.0 fluorometer with the Qubit dsDNA HS Assay Kit (Thermo Fisher Scientific, Q32854) and an Agilent Fragment Analyzer (Agilent) with a HS NGS Fragment Kit (Agilent, DNF-474), respectively. Library pools were sequenced paired-end 150 bp reads on an Illumina NovaSeq 6000 instrument. The quality of the sequencing run was assessed using Illumina Sequencing Analysis Viewer (Illumina version 2.4.7) and all base call files were demultiplexed and converted into FASTQ files using Illumina bcl2fastq conversion software v2.20. Because genome sizes were unknown for most species, sequencing depth was adjusted under the assumption of a genome size of 1 Gb and targeting a sequence coverage of approximately 30× for diploids and approximately 60× for polyploids.

For PacBio HiFi sequencing, high-molecular-weight DNA was isolated from multiple tissues of a single individual per species using Qiagen Genomic-tips 20/G columns (Qiagen, 10223) in conjunction with the Genomic DNA Buffer Set (Qiagen, 19060) and according to the QIAGEN Genomic DNA Handbook instructions. Prior to library preparation, genomic DNA was assessed for quantity, quality and purity. Between 1000-5000 ng of gDNA was used to prepare barcoded PacBio SMRTbell libraries for each species. *Paedocypris* sp. libraries (6–7 kb inserts) were sequenced on a PacBio Sequel IIe platform using a Single-Molecule Real-Time (SMRT) cell, whereas the remaining three species (*Sundadanio atomus*, *Boraras brigittae*, *Rasbora kalochroma*) were prepared with 10–12 kb inserts and sequenced on a PacBio Revio system.

The quality control assessments, generation of libraries and sequencing were performed at the Next Generation Sequencing (NGS) Platform, University of Bern, Switzerland, with the exception of 15 species sampled in India, which were sequenced at Agrigenome Labs Pvt. Ltd., India (Supplementary Table 1).

### Quality control, kmer profiling and genome assembly

Quality control (QC) of the Illumina short-read datasets was performed using FastQC(40) and summarized with MultiQC (41). Following QC, short reads were preprocessed in two sequential rounds using fastp (42). In a first round, adapters were trimmed (--detect_adapter_for_pe), base correction was applied to overlapping regions of paired-end reads (--correction), duplicate reads were removed (--dedup), polyG tails were trimmed (--trim_poly_g), and polyX tails were trimmed at the 3′ ends (--trim_poly_x). Quality filtering was disabled in this step (-- disable_quality_filtering). In a second round, reads were quality trimmed using a sliding window approach from the 5′ to 3′ direction (--cut_right), with a window size of 4 (-- cut_right_window_size), requiring a mean Phred quality score of 20 (--cut_right_mean_quality) and a minimum qualified Phred score of 20 (--qualified_quality_phred).

K-mer profiling and genome characterization were conducted using KMC 3 (43) and GenomeScope 2.0(44) with k = 21 and k = 31 to evaluate k-mer coverage and infer genome size, heterozygosity, and repeat content.

*De novo* assembly of the Illumina short reads was performed using two assemblers: ABySS 2.0 (45) and SPAdes (46). ABySS assemblies were first optimized across 20 representative species spanning divergent lineages, with parameters for k = 70, 80, 90, and 95. Based on this initial assembly statistics, the overall optimal parameters (k = 90, kc = 2) were subsequently applied to all taxa. SPAdes assemblies were generated using default settings.

Assembly quality was then assessed using QUAST (47) for general assembly statistics, and genome completeness was evaluated using Benchmarking Universal Single-Copy Orthologs (BUSCO) (48, 49), as implemented in compleasm (50), with the actinopterygii_odb10 (n = 3640) lineage dataset as the reference set of universal single-copy orthologs. We then selected the respectively best assembly from ABySS and SPAdes per each species based on the complete single-copy BUSCO score.

For the PacBio long-read datasets, HiFiAdapterFilt (51) was used to remove adapter sequences and other sequencing artifacts from the raw PacBio HiFi reads. To estimate genome size, heterozygosity, and repetitive content, k-mer analysis was performed using Meryl(52) and GenomeScope 2.0 with k = 21 and k = 31.

Genome assembly followed the Vertebrate Genome Project (VGP) standard workflow (53), with minor modifications at specific stages. hifiasm (54) was used in solo mode with light purge settings to generate a pseudohaplotype assembly, producing both primary and alternate contig sets. The primary assembly was processed with purge_dups (55) to identify and remove haplotypic duplications. The haplotigs extracted during this step were concatenated with the alternate contig set, after which purge_dups was executed a second time to further refine the alternate assembly. This iterative procedure produced a final, purged primary and alternate assembly pair.

Assembly quality was evaluated at multiple stages using: Merqury (52) to assess consensus QV and k-mer completeness; QUAST for contiguity and assembly statistics; and BUSCO to evaluate genome completeness.

### Taxonomic validation and contamination screening

To ensure accurate species identification and to detect potential sample contamination, we conducted taxonomic validations for all newly sequenced samples. Because our material originated from multiple sources, it was important to confirm species identity, ideally to the species level, but at minimum to the genus level before including the respective samples in the downstream phylogenomic analyses.

For each Illumina short-read dataset, we first assembled the mitochondrial genome using MITObim (56), with either *Cyprinus carpio* (NC_001606.1) or *Danio rerio* (NC_002333.2) as the initial reference. When these references failed to produce a mitogenome, we used the mitogenome of a closely related species as bait for iterative mapping.

The assembled mitogenomes from the Illumina short-read dataset were annotated using MITOS (57). From each annotated mitogenome, the *cytochrome c oxidase subunit I* (*cox1*) locus was extracted and trimmed to approximately 650 bp, corresponding to the standard DNA barcoding fragment, using Geneious (58). This barcode fragment is widely represented across taxa in the National Center for Biotechnology Information (NCBI) database (59, 60), facilitating taxonomic comparison. Each *cox1* sequence was queried against NCBI using BLASTN (61), and the top 10 hits per sample were retrieved. All sequences were aligned, and a neighbour-joining (NJ) tree was constructed in Geneious to visualize relationships and verify sample identity.

Samples for which *cox1* could not be extracted, or insufficient closely related sequences were available in GenBank, we additionally extracted and analyzed the mitochondrial *cytochrome b* (*cytb*) gene following the same workflow.

If the original identification of the sample showed high sequence similarity (≥95%) in *cox1* and/or *cytb* hits, these were considered correctly identified at the species or genus level. If samples lack close hits (<95% identity) or samples for which *cox1* or *cytb* sequences indicated a different taxonomic identity than recorded were cross-checked with sample providers who are taxonomic experts to confirm or revise identification. Samples in which *cox1* and *cytb* sequences yielded markedly discordant identifications were interpreted as potential contaminations and excluded from the downstream analyses.

### Incorporation of publicly available genomic resources

In addition to the newly generated datasets, we incorporated publicly available genomic data from the NCBI. All available genome assemblies for Cypriniformes and relevant outgroups were reviewed as of 1 July 2024.

Genomes were selected for inclusion according to the following criteria: (*i*) The genome assembly was associated with a published article. (*ii*) If not associated with a published article, the genome was explicitly declared publicly available for unrestricted use; in such cases, we contacted the corresponding data providers to confirm permission. (*iii*) The genome had been generated at the Wellcome Sanger Institute by some of us (KH, RD) and were made available for public or collaborative use. Genomes with unclear data-use status or active embargoes were not considered.

The application of these criteria resulted in the inclusion of 95 publicly available Cypriniformes genomes and 19 outgroup genomes (Supplementary Table 2) representing Characiformes (Characoidei and Citharinoidei both included), Gymnotiformes, Siluriformes, Gonorynchiformes, and Clupeiformes within the Otomorpha clade(62–64). The assembly quality for these 114 genomes was evaluated using BUSCO.

Additionally, 18 SRA datasets representing species of Danionidae, *Paedocypris* sp., and *Sundadanio* sp. were downloaded (Supplementary Table 2). These datasets were either associated with a published article or generated by the Wellcome Sanger Institute by some of us (KH, RD). Quality control, k-mer profiling, genome assembly and taxonomic validation for these SRA samples followed the same workflow described above for the newly generated Illumina short-read dataset.

### Genome dataset curation and filtering

The combination of newly generated genomic data and publicly available resources resulted in a total set of 389 genome assemblies representing Cypriniformes and relevant outgroups. To ensure data quality, an initial round of filtering was applied based on contamination detection, hybrid status, QC and assembly quality metrics.

Two samples (LR12807, LR02128) were identified as probable contaminants based on incongruent mitochondrial signals and inconsistent taxonomic placement and were excluded. One sample (LR15718) was identified as a probable hybrid individual based on morphology and excluded. Four samples (LR15296, LR14717, LR10789, 1736_2012) had insufficient assembly quality thresholds and completeness metrics and were excluded. These four samples had elevated read error rates and poor model fit in k-mer profiling with 10% discrepancy between genome size estimates for k = 21 and k = 31 models, unrealistically high inferred heterozygosity, and low BUSCO single-copy ortholog recovery across both ABySS and SPAdes assemblies.

Across the combined dataset, 47 species were represented by more than one genome assembly. In such cases, we retained the assembly with the highest BUSCO single-copy completeness score for downstream phylogenomic analyses.

After these filtering steps, the final curated dataset comprised 335 species (Supplementary Table 3; Supplementary Fig. 1c-d.

### Construction of phylogenomic datasets

We extracted multiple genome-wide marker datasets comprising both protein-coding exons and non-coding regions to construct phylogenomic matrices for our dataset. Four primary datasets were developed: three exon-based datasets (BUSCO, HUGHES, and MALMSTRØM) and one dataset based on ultraconserved elements (UCEs). The extraction, alignment, and trimming workflows are described below.

#### a. BUSCO dataset

The first exon dataset was derived from the BUSCO complete single-copy gene set (actinopterygii_odb10) (48, 65). BUSCO loci represent near-universal, single-copy orthologs and have been used in phylogenomic studies across diverse taxa (66–69), though their utility has not been extensively tested in large-scale phylogenomic analyses in teleosts.

BUSCO analyses were initially performed to assess genome completeness for all assemblies. From the BUSCO output files, we extracted the coordinates of complete single-copy loci, which encompass both coding sequences (CDS) and introns. Using custom scripts, we retrieved only the CDS regions from each assembly to generate nucleotide sequences for each BUSCO locus.

The BUSCO nucleotide sequences were concatenated locus-wise to generate multiple sequence alignments (MSAs). Each MSA was aligned and refined through several steps. First MACSE v2 (70) was used to align nucleotide sequences (NT) based on their amino acid (AA) translations (-prog alignSequences), producing both NT and AA alignments. Block Mapping and Gathering with Entropy (BMGE) (71) was used to trim the AA alignments (-m BLOSUM30), retaining only phylogenetically informative regions. Then TAPER (72) was applied to further clean alignments by distinguishing true sequence variation from putative alignment errors. The trimmed AA alignments were then used to mask the corresponding NT alignments in MACSE v2 (-prog reportMaskAA2NT, -mask_AA X, -min_percent_NT_at_ends 0.5, -dist_isolate_AA 3). The resulting masked NT alignments were then trimmed in MACSE v2 (-prog trimAlignment, - respect_first_RF_ON) to preserve the reading frame and remove low-coverage ends. BMGE was used again on the trimmed NT alignments (-e 0.8, -g 0.2) to eliminate highly variable or gappy sites. A final cleaning step using TAPER was carried out on the NT alignments.

#### b. HUGHES dataset

The second exon dataset was based on the teleost-specific exon probe set developed by Hughes et al. (73), designed for large-scale fish phylogenomic analyses and filtered for paralogs. Locus alignments (File3_ExonAlignments) were obtained from the authors’ Figshare repository (https://figshare.com/articles/dataset/Data_from_Exon_probe_sets_and_bioinformatics_pipelines_for_all_levels_of_fish_phylogenomics/11844783). For each locus alignment, a HMMER profile was constructed and used to search our genome assemblies following the pipeline outlined by Hughes et al. (73) (https://github.com/lilychughes/FishLifeExonHarvesting), with minor modifications. The detected exons were harvested and concatenated locus-wise to generate MSAs. Each nucleotide MSA was then aligned with MAFFT (74), trimmed using BMGE with default settings, and cleaned using TAPER.

#### c. MALMSTRØM dataset

The third exon dataset followed the approach of Malmstrøm et al. (75), who targeted orthologous nuclear genes to resolve deep teleost divergences. The orthologous exon set was downloaded from https://github.com/uio-cees/teleost_genomes_immune/tree/master/S11_S14_ortholog_detection_and_filtering/data/queries/nuclear. We used custom scripts implementing DIAMOND blastx (76) and SAMtools (77, 78) to retrieve orthologous sequences from each genome assembly. Loci were concatenated across taxa to form MSAs, and alignment and trimming followed the same steps as for the HUGHES dataset.

#### d. UCE dataset

In addition to the exon datasets, we harvested ultraconserved elements (UCE) loci, which comprise highly conserved core regions and variable flanking sequences that often include intronic and intergenic regions(79). The Ostariophysan UCE probe set (Ostariophysans-UCE- 2.7Kv1.fasta) was obtained from Faircloth et al. (80) (https://figshare.com/articles/dataset/Ostariophysans_6_737_baits_targeting_2_708_conserved_loci/7144199?file=13147316). UCE extraction followed the PHYLUCE(81) workflow for genome assemblies (https://phyluce.readthedocs.io/en/latest/tutorials/tutorial-3.html) with default parameters and 500 bp flanking regions. Internal alignments were generated with MAFFT using the --no-trim option, and the resulting MSAs were trimmed with Gblocks (82). Throughout the study, we use the term “locus” to refer collectively to both protein-coding exons and non-coding introns.

### Curation and quality filtering of phylogenomic datasets

Following the construction of the four phylogenomic datasets (BUSCO, HUGHES, MALMSTRØM, and UCE), we implemented a multi-step filtering strategy to remove low-quality alignments, poorly represented taxa, and potential paralogs. Filtering was performed at both the taxon and locus levels to ensure that only high-confidence orthologous loci were retained for subsequent phylogenomic inference.

#### a. Alignment-level and species-level filtering

For each exon dataset, we first removed taxa within an alignment that exhibited >50% missing characters (excluding gaps). Alignments were then checked for translation consistency. The presumed correct reading frame was identified, and loci containing more than two premature stop codons were discarded entirely. For loci containing one or two stop codons, only the affected species were excluded.

All exon alignments were manually inspected in AliView (83). Alignments exhibiting clear signs of misalignment, frame shifts, or paralogy (e.g., excessive sequence divergence or poor overlap among species) were removed. Alignments shorter than 150 bp were also discarded.

#### b. Missing data thresholds

To minimize missing data across the datasets, we removed any locus that was represented in fewer than 251 taxa (i.e., <75% of the total 335 species). This criterion was applied uniformly across the BUSCO, HUGHES, MALMSTRØM, and UCE datasets.

After these filtering steps, the retained loci were as follows: 769 BUSCO, 866 HUGHES, 1394 MALMSTRØM, and 896 UCE loci.

For each exon dataset (BUSCO, HUGHES, and MALMSTRØM), we generated three data partitions: full nucleotide sequences including all codon positions (NT.C123), nucleotide sequences excluding third codon positions (NT.C12), and translated amino acid sequences (AA). The UCE dataset, was composed mainly of noncoding loci and analyzed as a single nucleotide dataset.

In total, 10 datasets were generated: BUSCO.NT.C123, BUSCO.NT.C12, BUSCO.AA, HUGHES.NT.C123, HUGHES.NT.C12, HUGHES.AA, MALMSTRØM.NT.C123, MALMSTRØM.NT.C12, MALMSTRØM.AA, and UCE.

#### c. Phylogenetic reliability filtering with genesortR

For each locus in all datasets, we inferred maximum likelihood trees using IQ-TREE2(84) with ModelFinder (85) for model selection (-m MFP) and 1,000 ultrafast bootstrap replicates (86).

The resulting locus trees and alignments were subsequently filtered using genesortR (87), a multivariate method that optimizes phylogenetic signal while controlling for known sources of systematic bias. We ran genesortR with an outlier fraction of 0.3 and topological similarity disabled (topological_similarity = FALSE), thus excluding Robinson–Foulds distances from the scoring process.

After genesortR filtering, the final datasets comprised: 539 BUSCO, 607 HUGHES, 976 MALMSTRØM, and 628 UCE loci (Supplementary Fig. 2).

The locus properties estimated by genesortR were broadly consistent across datasets, with NT.C123, NT.C12, and AA showing similar patterns across BUSCO, HUGHES, and MALMSTRØM. The UCE dataset exhibited a pattern largely comparable to that of the NT.C12 datasets (Supplementary Fig. 3). Nonetheless, differences among NT.C123, NT.C12, and AA remained evident within datasets. For instance, NT.C123 datasets tended to display higher levels of saturation, faster evolutionary rates, and greater average bootstrap support and patristic distances relative to NT.C12, AA, and UCE (Supplementary Fig. 3). These curated and phylogenetically validated datasets formed the basis for subsequent concatenation- and coalescent-aware phylogenomic analyses.

### Assessing overlap among exon marker datasets

To evaluate potential redundancy among the three exon-based datasets (BUSCO, HUGHES, and MALMSTRØM), we examined whether any of the loci targeted the same orthologous regions in the zebrafish genome (*Danio rerio*). This analysis aimed to quantify overlap among marker sets and assess their relative independence within the phylogenomic framework.

For each multiple sequence alignment, the *D. rerio* nucleotide sequence was extracted when present. If *D. rerio* was absent, the longest sequence belonging to another species of *Danio* was used as a proxy. In cases where no *Danio* species were present, the longest sequence from any other danionid taxon was selected instead.

These representative sequences were then used as queries in BLASTN searches against the *D. rerio* reference coding sequence (CDS) database from Ensembl (GRCz11, release 114). Searches were performed and the resulting BLAST hits were sorted by e-value, retaining only the top hit per query.

For loci that did not yield significant hits against the CDS database, we subsequently performed BLASTx searches against the *D. rerio* protein database. Similarly, outputs were sorted by e-value, and the top hit per query was retained for downstream comparisons.

For the remaining loci that still did not produce any significant hits, we carried out manual BLAST searches using the Ensembl web interface (https://www.ensembl.org/Multi/Tools/Blast?db=core). The resulting hits were cross-referenced with the Ensembl GFF annotation files, and the corresponding Gene_ID, Transcript_ID, and Gene_Name were retrieved from the first annotated feature in the hit region.

In a few cases, the BLAST hit did not overlap with an annotated gene model. These loci were recorded as “unannotated hits”. When multiple loci from different datasets mapped to the same Ensembl Gene_ID, they were considered to represent the same orthologous locus.

In total, 45 loci lacked retrievable Ensembl Gene_IDs (due to incomplete or unannotated hits), and 39 alignments failed during BLAST searches. The remaining loci were mapped and used to visualize the extent of overlap among the three exon datasets. The overlap among the three exon datasets (BUSCO, HUGHES, and MALMSTRØM) was low, indicating that these marker sets are largely non-redundant and represent relatively independent samples of the genome within our phylogenomic framework (Supplementary Fig. 2).

### Phylogenomic inference

Phylogenomic inferences were performed for each of the ten primary datasets (BUSCO.NT.C123, BUSCO.NT.C12, BUSCO.AA, HUGHES.NT.C123, HUGHES.NT.C12, HUGHES.AA, MALMSTRØM.NT.C123, MALMSTRØM.NT.C12, MALMSTRØM.AA, and UCE), as well as for the combined BUSCO.NT.C12 + UCE nucleotide dataset. Multiple analytical frameworks were employed, encompassing both concatenation-based and coalescent-aware species tree approaches, as well as a novel reference-free genome-wide inference method.

Details of the datasets used in each phylogenomic inference, including the number of taxa, retained loci, and alignment lengths, are summarized in Supplementary Table 4.

#### a. Concatenation (supermatrix) analyses

For concatenated analyses, we used IQ-TREE2 with ModelFinder for model selection (-m MFP). We applied the --merge option to optimize partition merging and improve model fit.

#### b. Coalescent-aware species tree inference

To account for ILS under the multi-species coalescent (MSC) model (88), we used following complementary coalescent-aware methods implemented in the program ASTER (Accurate Species Tree EstimatoR) (89).

Species trees were inferred using ASTRAL-IV (89, 90) based on the individual locus trees obtained from IQ-TREE2 analyses. ASTRAL-IV estimates unrooted species trees from a set of unrooted single-copy locus trees and accommodates missing data effectively. Furthermore, ASTRAL-IV implements CASTLES-Pro (91), enabling the estimation of both terminal and internal branch lengths in substitution-per-site units.

We further applied Weighted ASTRAL (wASTRAL), which assigns weights to locus trees based on branch lengths and support values, thereby reducing the influence of poorly resolved or conflicting loci (92). Analyses were conducted using default settings. Unlike ASTRAL-IV, wASTRAL does not compute substitution-based branch lengths, its results were used primarily for topological comparison and branch support estimation.

We also implemented CASTER (93), a recently introduced coalescence-aware method that directly infers species trees from MSAs rather than pre-estimated locus trees. We ran two CASTER models: CASTER-site, which infers species trees from site pattern frequencies under the F84 model and allows arbitrary rate heterogeneity across sites; and CASTER-pair, which models substitution patterns across pairs of sites using one of three six-parameter submodels of GTR (93). The CASTER-site and CASTER-pair approaches, originally developed for whole-genome alignment data, yielded several biologically implausible topologies. These inconsistencies likely reflect limited sampling power and sensitivity of site-pattern-based models when applied to reduced genomic datasets such as ours (93); therefore, CASTER inferences are not presented in the results.

#### c. Reference-free, orthology-free species tree inference

In addition to the above analyses, we tested a fully automated, reference-free species tree estimation pipeline, ROADIES (94). ROADIES (Reference-free, Orthology-free, Annotation-free, Discordance-aware Estimation of Species trees) provides an end-to-end framework that infers species trees directly from raw genomic assemblies, eliminating the need for predefined ortholog sets.

ROADIES consists of six stages: starting from random subsequences sampled from each input genome, each treated as a putative “locus”. Then it carries out a pairwise alignment of each subsequence against all input assemblies using LASTZ (95) to identify homologous regions. Then low-quality and redundant alignments are filtered. Then the homologous regions are aligned using PASTA (96) to make MSA. Based on these MSA, locus trees are inferred using RAxML-NG under a maximum likelihood framework(97). Finally, a species tree is estimated based on these locus trees using ASTRAL-Pro-3 which can accommodate multi-copy genes (98).

ROADIES was executed thrice independently under Accurate Mode, with IDENTITY = 50, which defines the percentage of aligned base pairs (matches + mismatches) required in LASTZ.

#### d. Heterotachy model

To assess the impact of rate and compositional heterogeneity on phylogenetic inference, we applied the GHOST mixture model in IQ-TREE2, which explicitly accommodates heterotachy by allowing substitution rates and base composition to vary across sites and rate categories (99). Owing to the computational demands of this model, we analyzed a reduced dataset comprising 140 representative species that preserved complete family-level and major lineage coverage. The analysis was based on the BUSCO.NT.C12 + UCE nucleotide alignment. We evaluated ten alternative GHOST models, and model fit was assessed using Akaike Information Criterion (AIC) and Bayesian Information Criterion (BIC) scores to identify the optimal configuration (Supplementary Table 5). The resulting topology was then compared with both concatenation- and coalescent-aware species trees to evaluate the influence of heterotachy correction on the resolution of relationships within Cypriniformes.

Among nucleotide datasets, those including all codon positions (NT.C123) from BUSCO, HUGHES, and MALMSTRØM showed some incongruent relationships, likely driven by substitutional saturation at third codon positions, a common issue in deep-scale phylogenomic analyses (28, 36, 37). Accordingly, we focus subsequent interpretations in results based on the NT.C12, AA and UCE datasets, which exhibited higher overall resolution and reduced levels of homoplasy.

### Assessment of branch support, topological variation and relative rate variation

Statistical support for branches in the concatenated supermatrix phylogenies was evaluated using ultrafast bootstrap approximation with 1,000 replicates implemented in IQ-TREE2.

For coalescent-aware species trees inferred with ASTRAL-IV, wASTRAL, and ROADIES (which uses ASTRAL-Pro-3), branch support was assessed using local posterior probabilities (100). For species trees estimated with CASTER-site and CASTER-pair, branch support was quantified using 1,000 local block bootstrap replicates (93).

To characterize the degree of topological variation and biological heterogeneity among loci, we estimated gene concordance factors (gCF) and site concordance factors (sCF) using IQ-TREE2 (101–103). sCF values were first calculated using the ASTRAL-IV species tree as a reference topology with maximum likelihood approach (--scfl 100) for each dataset, and the resulting annotated tree was then used as input (--te) to compute gCF values across loci. For each dataset, we examined the relationship between branch-specific concordance and evolutionary rate by plotting gCF and sCF against log-transformed branch lengths. Pearson correlation coefficients were computed, and linear regression models were fitted to evaluate the strength and direction of these relationships (Supplementary Table 6).

Furthermore, we calculated quartet concordance factors (qCF) using the parameter -u 2 during the ASTRAL-IV species tree inference. Here we interpret all types of CFs as descriptors of topological variation rather than as measures of statistical support (101).

Together, these complementary metrics provide an integrated assessment of branch robustness and the extent of locus tree–species tree discordance within our phylogenomic datasets.

To explore relative substitution rate variation across Cypriniformes, we calculated root-to-tip distances based on the rooted concatenated phylogeny for each of the ten datasets, after pruning all outgroup taxa. Root-to-tip distances were computed in R Statistical Software 4.0.0 (104) using the package ape (105). The GC distribution of the nucleotide datasets was computed using summary statistics in AMAS (106). The root-to-tip distances and the GC distributions were visualized with ggplot2 (107).

### Phylogenetic Placement of Paedocyprididae and Sundadanionidae

Two lineages with progenetic miniaturized taxa within Cypriniformes, *Paedocypris* and *Sundadanio* have long been associated with substantial phylogenetic uncertainty (25), leading to their recognition as separate families (Paedocyprididae and Sundadanionidae, respectively). Competing hypotheses variously place *Paedocypris* as sister-group to (i) all other Cypriniformes, (ii) the suborder Cyprinoidei, or (iii) within Danionidae (e.g., 20, 21, 23, 24, 26, 30). Similarly, *Sundadanio* has been alternately recovered as part of Danionidae or as an independent lineage within Cyprinoidei (e.g., 21, 23, 24, 26, 30).

We examined concordance vectors (101) for branches associated with Paedocyprididae and Sundadanionidae. Concordance vectors summarize gene-tree heterogeneity at a focal branch into four proportions that sum to one: ψ₁, the concordance factor (proportion of gene/site/quartet supporting the focal species-tree branch), ψ₂ and ψ₃, the two dominant alternative topologies (discordance factors, ψ₂ always greater than ψ₃), and ψ₄, the residual proportion representing all other discordant topologies. A low ψ₁ value indicates that, despite high species-tree certainty, most loci do not support the inferred history for that branch, thereby pinpointing potential sources of conflict or ILS (101).

To explicitly test alternative hypotheses for the placement of Paedocyprididae and Sundadanionidae, we employed Gene Genealogy Interrogation: GGI (108) using the BUSCO.NT.C12, HUGHES.NT.C12, MALMSTRØM.NT.C12, and UCE datasets. GGI evaluates competing topological hypotheses by comparing likelihoods under constrained maximum-likelihood searches for each locus, thereby identifying which hypothesis each locus most strongly supports while mitigating stochastic gene-tree error.

For each locus, input topologies were pruned to match the taxon sampling of each alignment using nw_prune from the newick_utils package (109). Then constrained ML tree searches were conducted in IQ-TREE2 under each alternative topology, followed by the approximately unbiased (AU) topology test (110) with parameters -zb 20000 -au --sitelh --ntop 40.

The hypotheses tested for Paedocyprididae were as follows: H1: Paedocyprididae plus Danionidae form a monophyletic group; H2: Cyprinoidei form a monophyletic group excluding Paedocyprididae; H3: Cyprinoidei form a monophyletic group including Paedocyprididae but excluding Danionidae; H4: Cypriniformes form a monophyletic group excluding Paedocyprididae; and H5: Cypriniformes form a monophyletic group including Paedocyprididae but excluding Danionidae. Here, H1–H3 and H1, H4, H5 were treated as alternative hypothesis sets and H2/H4 and H3/H5 are nested and were not compared directly.

The hypotheses tested for Sundadanionidae were as follows: H1: Sundadanionidae plus Danionidae are monophyletic; H2: Sundadanionidae form a clade with Leptobarbidae, Xenocyprididae, Acheilognathidae, Gobionidae, Tincidae, Leuciscidae, Tanichthyidae excluding Danionidae; H3: Danionidae form a clade with the above group excluding Sundadanionidae.

We further quantified gene-wise phylogenetic signal (111) for the above relationships. For each locus, the log-likelihood difference (ΔGLS) between two competing topologies was computed to assess the degree of support for each topology. Distributions of ΔGLS across loci were visualized to identify whether a small subset of loci disproportionately influenced overall support, an indicator of contentious branches in large phylogenomic datasets (111).

Finally, we examined whether locus properties estimated by genesortR (alignment length, average bootstrap support, average patristic distances, evolutionary rate, missing data, occupancy, proportion of variable sites, root to tip variance, saturation, tree length, and treeness) varied among loci supporting alternative hypotheses. Patterns in these properties can reveal whether biases such as rate heterogeneity, missing data, or alignment quality correlate with the topological preferences inferred from GGI and ΔGLS analyses.

All visualization and hypothesis comparison analyses were conducted in R, and locus trees lacking any representatives of Paedocyprididae or Sundadanionidae were excluded in the above analyses.

## Results

### Expanding the phylogenomic framework for Cypriniformes

The phylogenomic framework of this study expands both the taxonomic and genomic coverage for Cypriniformes. Our analyses include 316 Cypriniformes species, representing a ∼84% increase in taxon sampling over Stout et al.’s (23) coverage. The largest combined dataset in our study, the BUSCO.NT.C12 + UCE matrix, comprises 541,983 bp, representing a ∼229% increase in overall alignment size (sites x species) over the dataset of Stout et al.(23).

Across all analyses, both concatenated supermatrix and coalescent-aware species tree inferences (ASTRAL-IV and wASTRAL) recovered broadly congruent phylogenetic relationships (Fig. 1; Supplementary Fig. 4-5). Species trees inferred using the reference-free ROADIES framework which follows a fundamentally different inference workflow were also largely consistent with these results (Supplementary Fig. 6). The taxonomically reduced dataset analyzed under the GHOST model, which accounts for heterotachy, revealed broadly similar relationships to the main analyses (Supplementary Fig. 7).

**Fig. 1.**
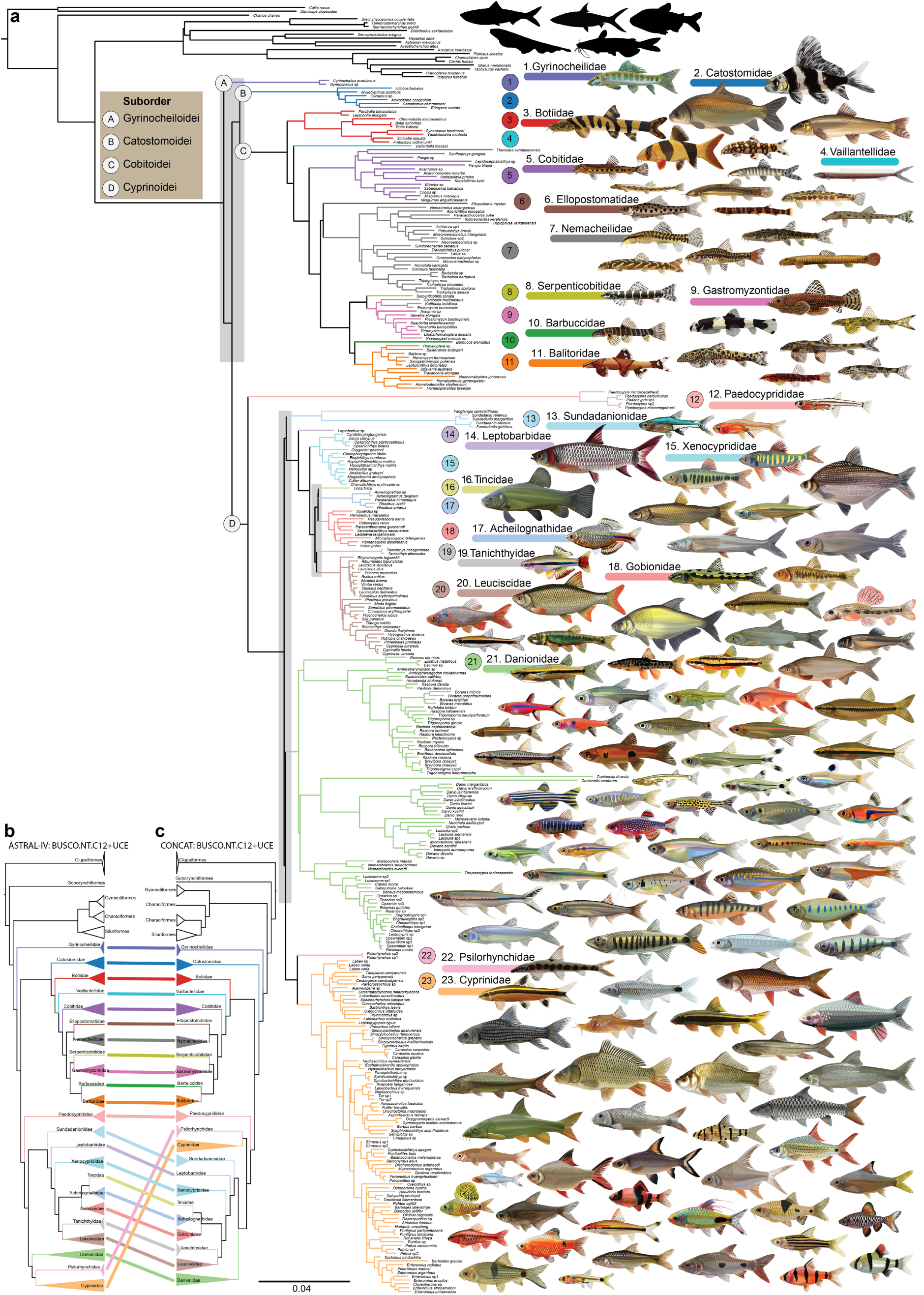
**(a)** ASTRAL-IV species tree showing the hypothesis of phylogenomic relationships of Cypriniformes inferred from 335 species and 1,167 loci (total alignment length: 541,983 bp), based on combined first and second codon positions of BUSCO.NT.C12 and UCE nucleotide dataset. The numbers indicate the different Cypriniformes families. The major conflicting zones of phylogenetic relationships are highlighted in grey (see Fig. 2; Supplementary Fig. 10-11). Simplified family-level topologies are shown for **(b)** the ASTRAL-IV species tree and **(c)** the IQ-TREE2 concatenated phylogeny for the same dataset.

Overall, our phylogenomic results corroborate the prevailing understanding of Cypriniformes evolutionary relationships established through decades of morphological and molecular systematic research (Fig. 1). Concatenation-based analyses generally recovered similar topologies across all datasets, whereas coalescent-aware analyses revealed modest variation among marker sets.

Statistical support for major relationships was uniformly high across datasets, as expected for genome-scale analyses with low sampling variance (101, 112). Nonetheless, considerable topological heterogeneity was detected between loci, as reflected by concordance factors (CFs). BUSCO.NT.C12 and UCE datasets exhibited the highest overall concordance, whereas HUGHES.NT.C12 and MALMSTRØM.NT.C12 showed greater discordance.

All family level taxa represented by multiple species were consistently recovered as monophyletic with strong statistical support across inference frameworks (Supplementary Fig. 8). However, the degree of underlying gene-tree discordance varied substantially between families.

Taken together, the overall stability of well-established relationships and the distribution of contentious branches appear unevenly spread across the tree, with both strongly supported and conflicting relationships occurring throughout the phylogenetic tree (Fig. 1).

### Stable phylogenetic relationships within Cypriniformes

Across all datasets and inference approaches, multiple regions of the Cypriniformes phylogeny exhibited strong and consistent support, indicating a broadly stable backbone topology for the group.

The phylogenetic relationships among loaches were consistently and strongly supported across all datasets and inference frameworks (branch C in Fig. 1; Supplementary Fig. 9). Within Cobitoidei, Botiidae was consistently recovered as the sister lineage to all other loaches, followed successively by Vaillantellidae and Cobitidae. A well-supported monophyletic group comprising Gastromyzontidae, Serpenticobitidae, Balitoridae, and Barbuccidae was recovered across all analyses, with Gastromyzontidae + Serpenticobitidae forming a sister-group relationship to Balitoridae + Barbuccidae in most cases (Fig. 1; Supplementary Fig. 9).

The enigmatic loach family Ellopostomatidae was recovered as sister-group to Nemacheilidae in most analyses; however, this relationship received comparatively weak support relative to other loach clades and was characterized by a short internal branch (Fig. 1). The clade formed by Gastromyzontidae + Serpenticobitidae + Balitoridae + Barbuccidae was inferred as the sister-group to the Nemacheilidae + Ellopostomatidae in most of the analyses (Fig. 1; Supplementary Fig. 9).

Within Cyprinoidei, several relationships also showed strong and consistent support across all analyses. The sister-group relationship between Cyprinidae and Psilorhynchidae (numbers 22 and 23 in Fig. 1; Supplementary Fig. 10) was recovered in every dataset, corroborating previous molecular studies (113, 114). Another consistently supported relationship within Cyprinoidei was the placement of Leptobarbidae and Xenocyprididae as successive sister-groups to the clade comprising Acheilognathidae, Gobionidae, Leuciscidae, Tanichthyidae, and Tincidae (numbers 14 and 15 in Fig. 1; Supplementary Fig. 11).

While these clades represent the most stable components of the Cypriniformes backbone, several other relationships exhibited notable topological variation among datasets and inference frameworks.

### Conflicting phylogenetic relationships within Cypriniformes

Several regions of the Cypriniformes phylogeny are characterized by ILS (Fig. 1), a hallmark of extremely short internal branches and/or rapid lineage diversification (88, 115). These regions correspond to three principal zones of phylogenetic conflict consistently recovered across our analyses (grey bars in Fig. 1).

The first major area of topological incongruence was observed among the early-branching relationships of the four Cypriniform suborders: Gyrinocheiloidei, Catostomoidei, Cobitoidei, and Cyprinoidei (Fig. 2; Supplementary Fig. 12). In all concatenated and most of coalescent-aware analyses, Gyrinocheiloidei was recovered as the sister lineage to all remaining Cypriniformes. In all concatenated supermatrix phylogenies, Catostomoidei was inferred as the sister-group to Cobitoidei + Cyprinoidei. In contrast, none of the coalescent-aware species tree inferences supported this topology. Instead, these analyses generally placed Catostomoidei as sister-group to Cobitoidei, with the combined Catostomoidei + Cobitoidei clade sister-group to Cyprinoidei (Fig. 2; Supplementary Fig. 12).

**Fig. 2.**
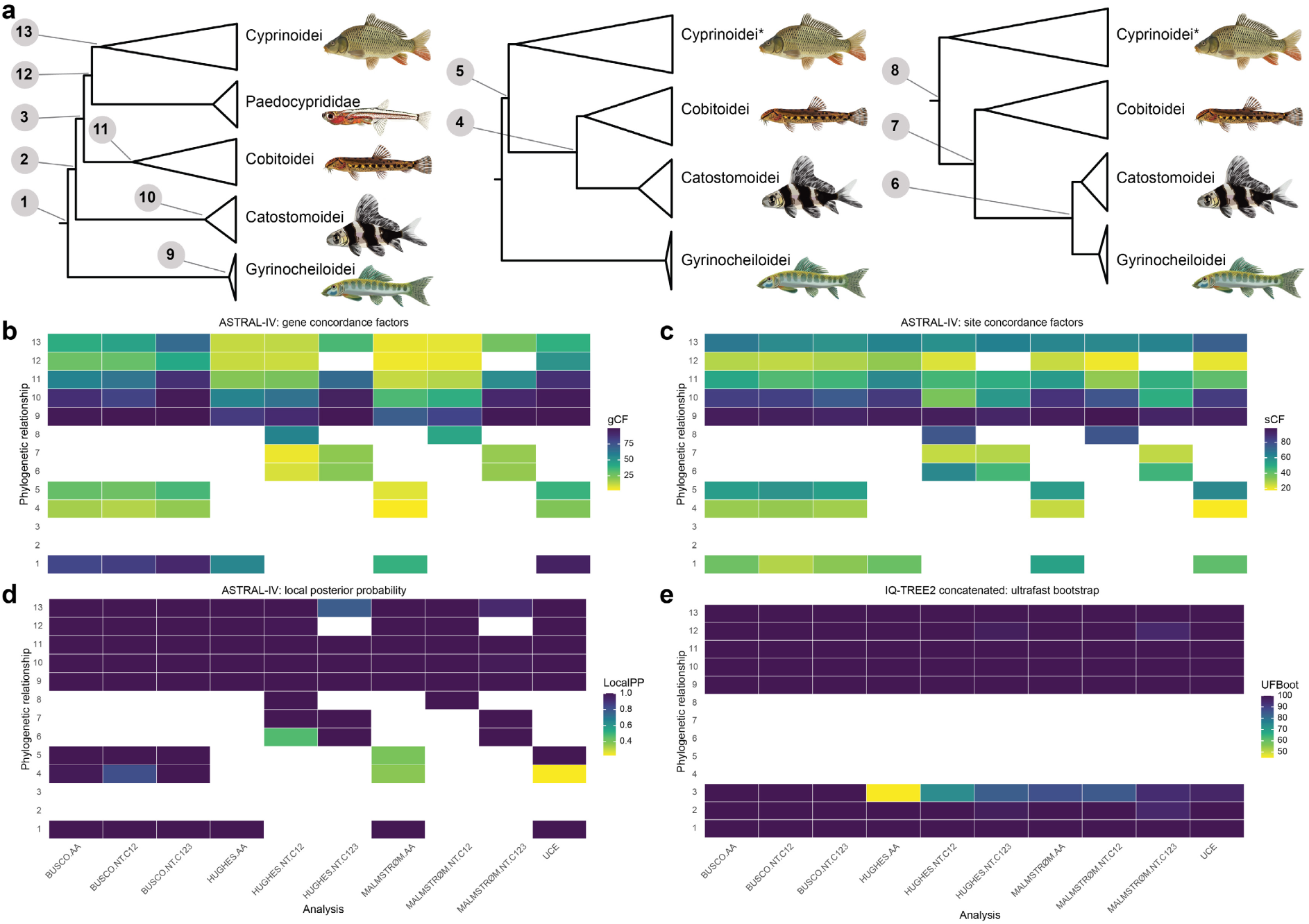
**(a)** Phylogenetic relationships among the suborders of Cypriniformes recovered from different analyses. Less frequently recovered relationships are not shown. Topological variation is illustrated based on **(b)** gene and **(c)** site concordance factors. Statistical support is indicated by **(d)** local posterior probabilities for the ASTRAL-IV species tree and **(e)** ultrafast bootstrap values for the IQ-TREE2 concatenated phylogeny. Increasing blue intensity indicate greater concordance or statistical support. An asterisk following Cyprinoidei indicates that Paedocyprididae is also included within this clade. The numbers in y-axis in **(b-e)** correspond to the phylogenetic relationships shown in panel **(a)**.

A second major zone of incongruence occurred at the base of Cyprinoidei, the most species-rich and taxonomically complex suborder of Cypriniformes. The conflicting placements primarily involved the position of early branching in Cyprinoidei (excluding Paedocyprididae), which was variably recovered as Danionidae as (i) the sister lineage to all other Cyprinoidei, (ii) sister-group to the Cyprinidae + Psilorhynchidae clade, or (iii) sister-group to all other Cyprinoidei excluding Cyprinidae + Psilorhynchidae (Supplementary Fig. 10).

A third region of topological conflict was found within the Cyprinoidei, involving the families Acheilognathidae, Gobionidae, Leuciscidae, Tanichthyidae, and Tincidae. Relationships among these lineages differed markedly among datasets and inference methods and exhibited some of the lowest gCF and sCF values across the phylogeny (Supplementary Fig. 11).

### Phylogenomic insights into zebrafish relatives

With approximately 364 species in 39 genera, the family Danionidae is one of the most species-rich lineages within Cypriniformes, and includes key vertebrate model organisms. Our sampling covers representatives from 34 genera, approximately 87% of all valid genera, providing one of the most comprehensive phylogenomic frameworks for the family to date and a foundation for comparative evolutionary and developmental studies.

Across all analyses, the phylogenetic position of *Danionella* was stable and consistently recovered as the sister lineage to all remaining members of the subfamily Danioninae, despite exhibiting pronounced rate heterogeneity (Fig. 1; Supplementary Fig. 10; see below). Within Danionidae, subfamilial relationships displayed varying levels of stability (Supplementary Fig. 10). Esominae showed topological shifts among analyses, being recovered as sister-group to Rasborinae in most concatenated trees and several coalescent-aware species trees, but as sister-group to the remaining Danionidae in others (Supplementary Fig. 10). Chedrinae was consistently inferred as sister-group to Danioninae, with the combined clade (Chedrinae + Danioninae) forming the sister-group to either Rasborinae + Esominae or to Rasborinae alone in most analyses (Supplementary Fig. 10).

### Elevated rate variation in progenetic miniatures

Progenetic miniatures exhibit profound developmental truncation, retaining larval-like morphology into adulthood (26, 116, 117). Within Cypriniformes, four genera, *Paedocypris*, *Danionella*, *Sundadanio*, and *Fangfangia* exhibit extreme progenetic miniaturization and remarkable morphological innovation (26, 29, 116–122).

Resolving the phylogenetic placement of these developmentally truncated lineages has long been problematic, as their extreme morphological reduction and possible molecular rate shifts can obscure deep evolutionary signal (26, 123, 124). Across all datasets, the four progenetic miniature genera, particularly *Paedocypris* and *Danionella* exhibited markedly higher substitution rates relative to other Cypriniformes, as indicated by the distribution of root-to-tip distances in the rooted concatenated phylogenies (Fig. 3a-b; Supplementary Fig. 13). Such rate heterogeneities can complicate phylogenetic reconstructions by introducing long-branch attractions (LBA) or by amplifying substitutional saturation in rapidly evolving lineages leading to the erosion of phylogenetic signal (5, 28, 125).

**Fig. 3.**
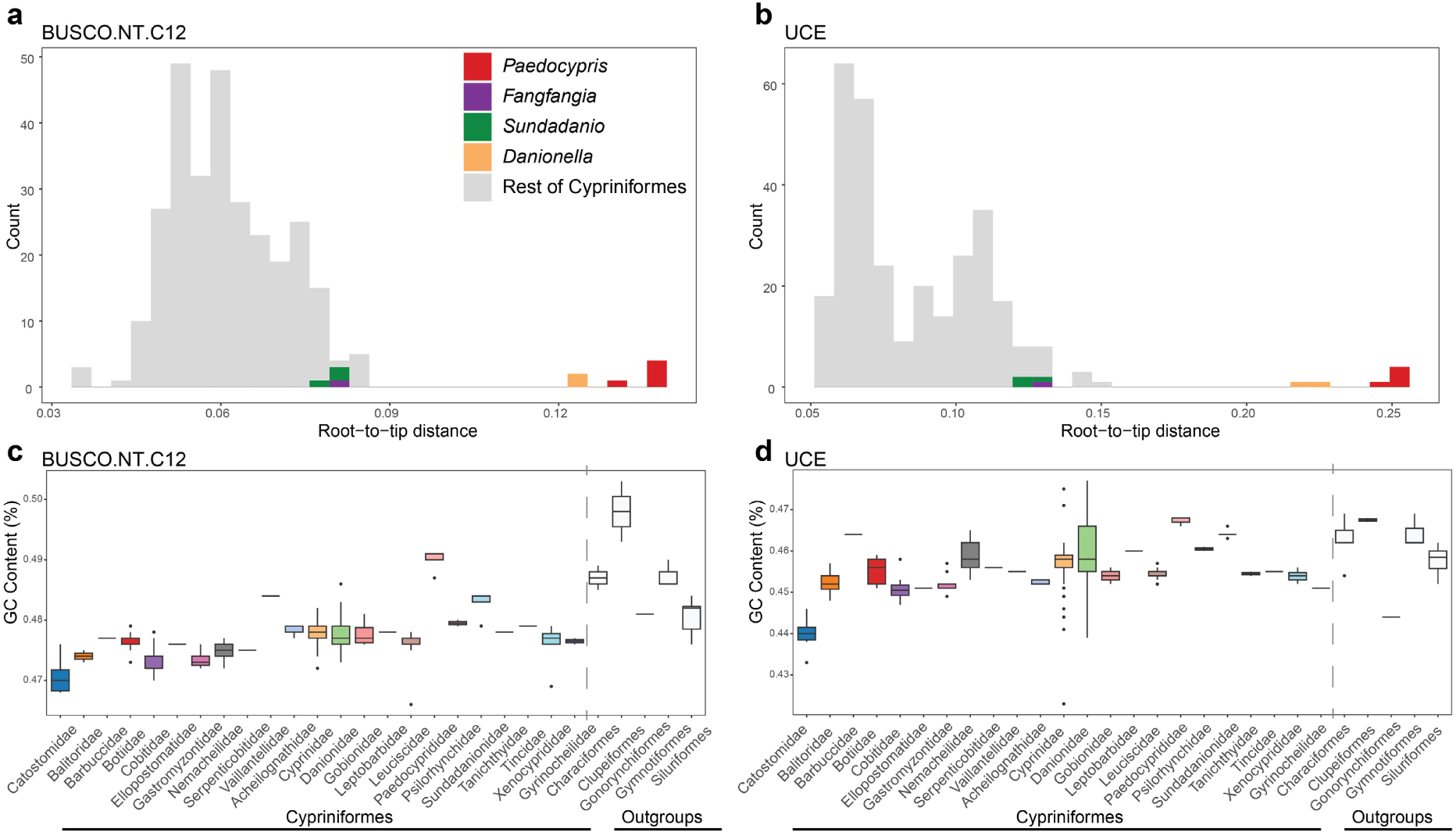
Patterns of rate and compositional variation across Cypriniformes. **(a–b)** Substitution rate variation estimated from the **(a)** BUSCO.NT.C12 and **(b)** UCE datasets, highlighting elevated evolutionary rates in the progenetic miniature genera *Paedocypris* and *Danionella*. **(c–d)** GC-content variation across taxa in the **(c)** BUSCO.NT.C12 and **(d)** UCE datasets, showing markedly higher GC% in *Paedocypris* relative to other Cypriniformes lineages.

### The phylogenetic position of *Fangfangia*

The progenetic miniature *Fangfangia spinicleithralis* was previously unrepresented in any genetic database and known only from its original morphological description(119). Based on shared derived unique morphological characters, Britz et al.(119) hypothesized a close relationship between *Fangfangia* and *Sundadanio*. Our phylogenomic analyses across all datasets support this relationship, consistently recovering *Fangfangia* and *Sundadanio* as sister taxa with high statistical support and high concordance in both gCF and sCF, strengthening this hypothesis (Fig. 1; Supplementary Fig. 10).

### Short internal branches and incomplete lineage sorting obscures the phylogenetic signal of Sundadanionidae

The discordant phylogenetic signal surrounding Sundadanionidae (*Fangfangia* + *Sundadanio*) likely reflects genuine biological incongruence, driven by ILS during an early, rapid radiation rather than analytical artefacts. Across all datasets, Sundadanionidae was consistently recovered as the sister lineage to the clade comprising Leptobarbidae, Xenocyprididae, Acheilognathidae, Gobionidae, Leuciscidae, Tanichthyidae, and Tincidae within Cyprinoidei, with strong statistical support, albeit with a very short internal branch (Fig. 4a-b; Supplementary Fig. 14-15).

**Fig. 4.**
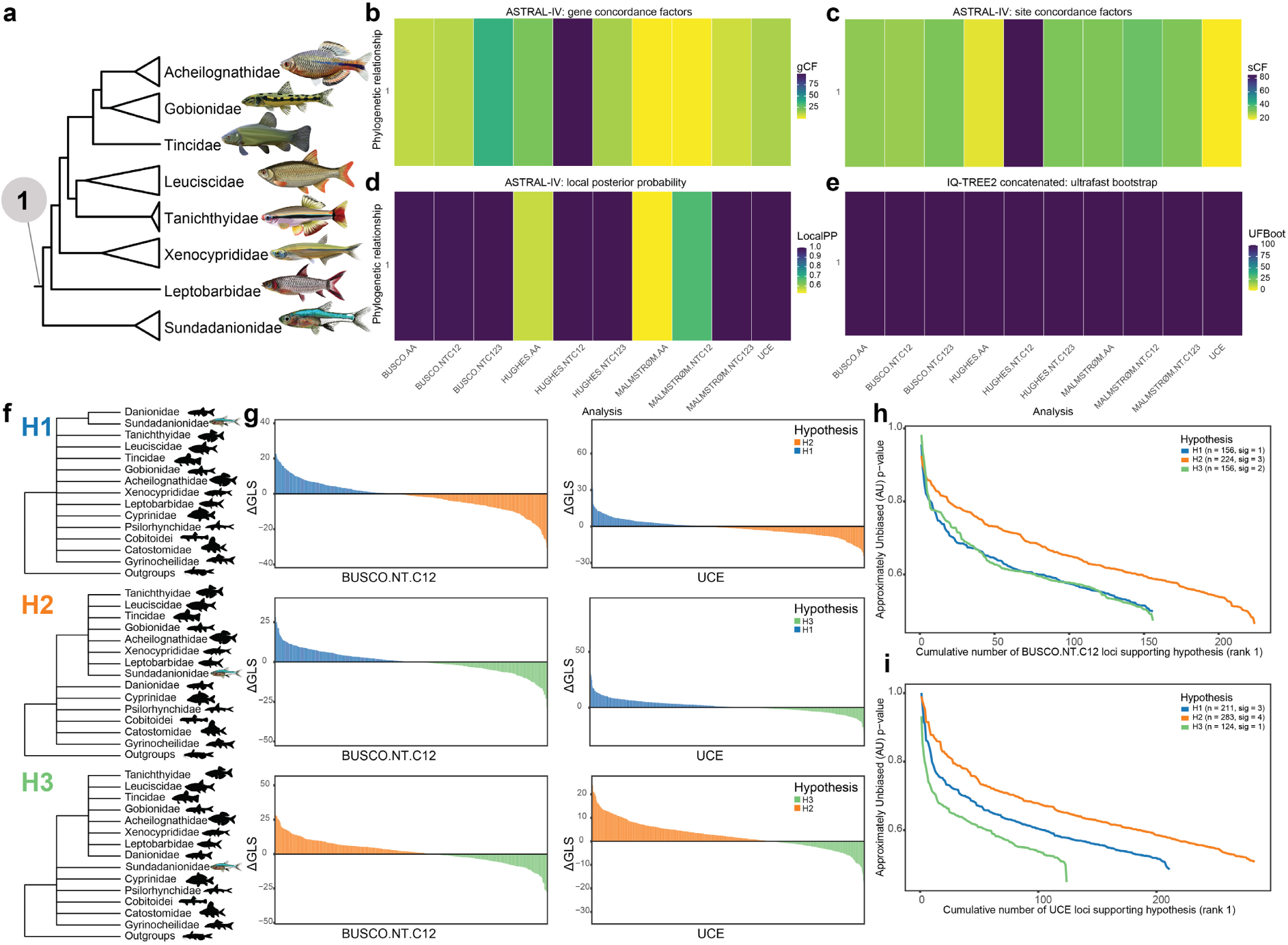
**(a)** Phylogenetic position of Sundadanionidae recovered from different analyses. Topological variation is illustrated based on **(b)** gene and **(c)** site concordance factors. Statistical support is indicated by **(d)** local posterior probabilities for the ASTRAL-IV species tree and **(e)** ultrafast bootstrap values for the IQ-TREE2 concatenated phylogeny. Increasing blue intensity indicate greater concordance or statistical support. The number in y-axis in **(b-e)** correspond to the phylogenetic relationship shown in panel **(a)**. **(f)** Topologies of alternative hypotheses tested for the phylogenetic position of Sundadanionidae. Gene-wise phylogenetic signal for alternative hypotheses in **(g)** BUSCO.NT.C12 and UCE datasets, with the y-axis representing differences in gene-wise log-likelihood scores (ΔGLS). Gene genealogy interrogation (GGI) applied to test alternative phylogenetic hypotheses for **(h)** BUSCO.NT.C12 and **(i)** UCE datasets. Plotted lines show the cumulative number of loci (x-axis) supporting each topology as the highest-ranked hypothesis (rank 1), with AU test P-values on the y-axis.

Patterns of concordance and discordance for Sundadanionidae were consistent across datasets (Supplementary Fig. 16). Gene-level concordance vectors revealed high residual discordance (ψ₄ > 78.5%) in all analyses. Site-level concordances were mixed: in UCE, BUSCO.NT.C12, BUSCO.AA, and MALMSTRØM.AA datasets, discordance (ψ₂ = 47–67%) exceeded concordance (ψ₁ = 19–33%), whereas in other datasets ψ₁ and ψ₂ were roughly balanced. Quartet-level concordance vectors showed generally even distributions (ψ₁ = 34–40%, ψ₂ = 30–34%) in NT.C12 and AA datasets, while UCE dataset exhibited higher concordance (ψ₁ = 42%) and lower discordance (ψ₂ = 29%: Supplementary Fig. 16).

To evaluate competing hypotheses, we performed 8,208 constrained ML searches and corresponding GGI and ΔGLS analyses. GGI ranked H₂ (in which Sundadanionidae forms a clade with Leptobarbidae, Xenocyprididae, Acheilognathidae, Gobionidae, Leuciscidae, Tanichthyidae, Tincidae) highest, followed by H₁ (Sundadanionidae plus Danionidae are monophyletic) and H₃ (Danionidae form a clade with the above group excluding Sundadanionidae), respectively. However, no single hypothesis received an overwhelming higher rank than others and many loci (>94%) were non-significant between alternative hypotheses (Fig. 4h-i; Supplementary Fig. 17). Such balanced GGI results indicate genuine genealogical conflict, consistent with ILS(108). Pairwise ΔGLS comparisons revealed no strong outliers, and the proportions of loci supporting alternative topologies were relatively balanced, suggesting that the observed conflict likely reflects biological signal rather than analytical noise (Fig. 4g; Supplementary Fig. 18). Gene-property comparisons across hypotheses showed no apparent differences among loci (Supplementary Fig. 19).

### Conflicting signals and rate heterogeneity obscure the placement of Paedocyprididae

In contrast to Sundadanionidae, the instability in the phylogenetic placement of Paedocyprididae appears to arise from a combination of weak phylogenetic signal, extensive rate heterogeneity, LBA, and potential model biases, rather than from ILS or stochastic error alone. Across most analyses, Paedocyprididae was recovered as the sister-group to all other Cyprinoidei, with strong statistical support (Fig. 5; Supplementary Fig. 20). However, in the HUGHES.NT.C123 and MALMSTRØM.NT.C123 datasets, coalescent-aware species trees inferred with ASTRAL-IV and wASTRAL placed Paedocyprididae as sister-group to all remaining Cypriniformes (Fig. 5; Supplementary Fig. 20). In contrast to the internal branch of Sundadanionidae, the internal branch of Paedocyprididae was longer across all the datasets (Supplementary Fig. 15).

**Fig. 5.**
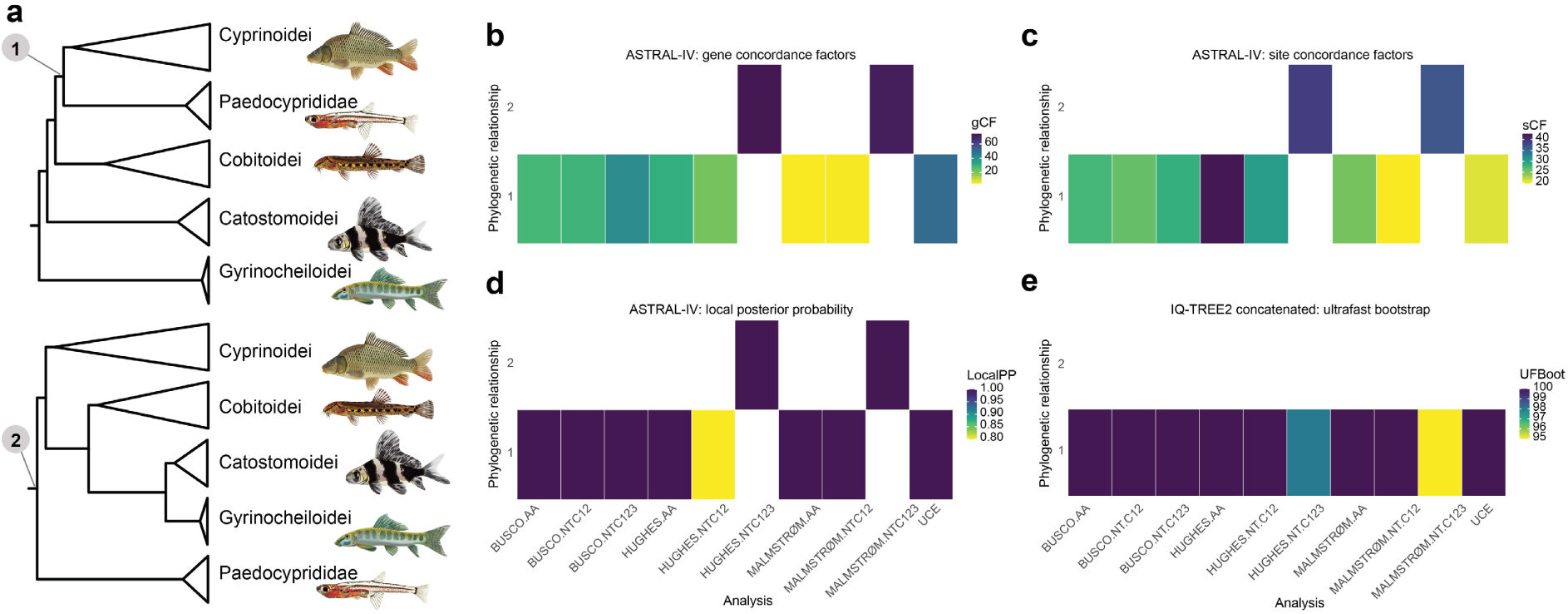
**(a)** Phylogenetic position of Paedocyprididae recovered from different analyses. Topological variation is illustrated based on **(b)** gene and **(c)** site concordance factors. Statistical support is indicated by **(d)** local posterior probabilities for the ASTRAL-IV species tree and **(e)** ultrafast bootstrap values for the IQ-TREE2 concatenated phylogeny. Increasing blue intensity indicate greater concordance or statistical support. The numbers in y-axis in **(b-e)** correspond to the phylogenetic relationships shown in panel **(a)**.

Exploration of topological variation using concordance vectors (ψ₁–ψ₄) revealed substantial gene-tree conflict. In the NT.C12, and AA datasets, residual discordance (ψ₄) was markedly greater than ψ₂ and ψ₃, with the HUGHES and MALMSTRØM datasets showing the strongest gene-tree discordance (ψ₄ > 79.8%) and very low gene concordance (ψ₁ = 4.5–10.8%; Supplementary Fig. 21). In contrast, the UCE dataset exhibited substantially higher overall concordance (ψ₁ = 50.4%) and lower residual discordance (ψ₄ = 38.8%; Supplementary Fig. 21). At the site level, most datasets displayed higher discordance (ψ₂ = 34.7–64.9%) relative to concordance (ψ₁ = 20– 33%) (Supplementary Fig. 21). Quartet-based concordance factors indicated variable but moderate-to-high concordance (ψ₁ = 38.9–66.9%), with the highest concordance observed in the UCE dataset (Supplementary Fig. 21).

To further evaluate alternative hypotheses, we conducted 12,265 constrained maximum-likelihood (ML) analyses using GGI and ΔGLS approaches on the NT.C12 and UCE datasets. In GGI analyses of hypotheses H₁–H₃, topology H₂ (Cyprinoidei monophyletic excluding Paedocyprididae) consistently ranked highest across all datasets (Fig. 6c; Supplementary Fig. 22). Similarly, for H₁ (Paedocyprididae plus Danionidae are monophyletic), H₄ (Cypriniformes monophyletic excluding Paedocyprididae), and H₅ (Cypriniformes form a monophyletic group including Paedocyprididae but excluding Danionidae), topology H₄ had the highest probability (Fig. 6d; Supplementary Fig. 22). Nonetheless, both analyses revealed a large proportion of non-significant loci (>91%) lacking sufficient signal to decisively favor any topology, indicating generally weak per-locus support. A greater proportion of loci favored H₂ and H₄ over alternative topologies based on the pairwise ΔGLS comparisons (Fig. 6b; Supplementary Fig. 23). Similarly, in comparisons between H₁ and H₃/H₅, a modest majority of loci favored H₁ (Fig. 6b; Supplementary Fig. 23).

**Fig. 6.**
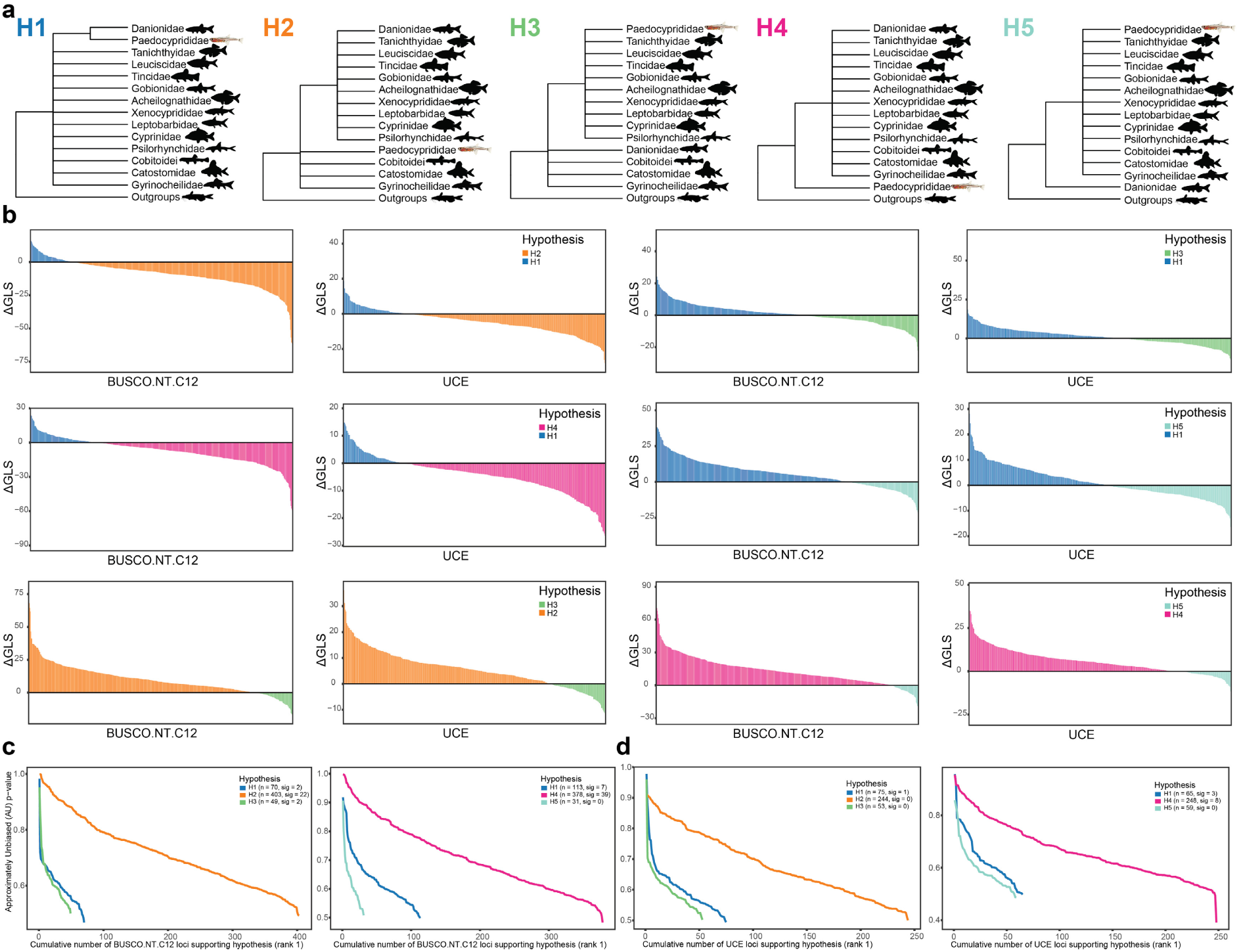
**(a)** Topologies of alternative hypotheses tested for the phylogenetic position of Paedocyprididae. Gene-wise phylogenetic signal for alternative hypotheses in **(b)** BUSCO.NT.C12 and UCE datasets, with the y-axis representing differences in gene-wise log-likelihood scores (ΔGLS). Gene genealogy interrogation (GGI) applied to test alternative phylogenetic hypotheses for **(c)** BUSCO.NT.C12 and **(d)** UCE datasets. Plotted lines show the cumulative number of loci (x-axis) supporting each topology as the highest-ranked hypothesis (rank 1), with AU test P-values on the y-axis.

Gene-property comparisons among loci supporting different topologies revealed no clear differences (Supplementary Fig. 24-25), suggesting no locus-specific biases. Furthermore, the comparison of GC content distributions across the nucleotide datasets indicates that Paedocyprididae exhibits a markedly elevated GC% relative to other Cypriniformes families (Fig. 3c-d; Supplementary Fig. 26). Notably, the GC% distribution of Paedocyprididae closely resembles that of the outgroup taxa, suggesting a potential case of compositional convergence. Such GC-biased similarity between distantly related lineages can artificially group them together as a result of LBA (28, 36). In the presence of LBA, GGI and ΔGLS on single-locus analyses may erroneously favor the majority species-tree topology (126).

## Discussion

Despite decades of morphological and molecular research, our understanding of evolutionary history of Cypriniformes has remained incomplete. Limited taxonomic and genomic sampling, coupled with pervasive phylogenetic conflict, has hindered efforts to establish an evolutionary backbone for the order. By integrating newly generated genome assemblies with available public resources, and using multiple genome-wide loci analyzed under both concatenation-based and coalescent-aware methods, our study provides the first complete family-level phylogenomic framework for Cypriniformes including a broad representation across genera to date.

The general congruence among concatenation-based and coalescent-aware frameworks suggests that the backbone topology of Cypriniformes is now established, enabling identification of well-supported clades versus regions of persistent conflict. The relationships within Cobitoidei are consistently well supported. Within Cyprinoidei, the sister-group relationship between Cyprinidae and Psilorhynchidae, and the placement of Leptobarbidae and Xenocyprididae, are strongly supported across analyses. These findings corroborate previous molecular studies (e.g., 17, 114, 127, 128).

Nevertheless, several regions of persistent phylogenetic conflict remain, concentrated along short internal branches that separate major lineages. The three main regions of discordance were among the early branching of the four suborders, the base of Cyprinoidei, and the diversification of the families Acheilognathidae, Gobionidae, Leuciscidae, Tanichthyidae, and Tincidae. The combination of extremely short internodes and heterogeneous gene histories is consistent with ILS arising from rapid successive speciation events. Under the multispecies coalescent model, when speciation events occur in rapid succession, ancestral polymorphisms can persist across successive lineages before fixation, producing stochastic variation in gene genealogies and consequently conflicting topologies among loci (88, 115). The fact that such incongruences are evident in both deep and recent divergences suggests that the Cypriniformes may have undergone repeated bursts of diversification during its evolutionary timescale. Similar patterns of both deep and shallow phylogenetic discordance have been observed in other rapidly diversifying groups, suggesting that such conflict may be a general feature of species radiations (e.g., 129–133). Collectively, these results indicate that much of the observed topological conflict within Cypriniformes likely reflects biological processes, principally rapid lineage divergence and ILS, rather than methodological artifacts at both deep and shallow phylogenetic levels.

One of the most striking results emerging from our analyses is the pronounced rate heterogeneity associated with progenetic miniature lineages, particularly in *Paedocypris* and *Danionella*. These taxa exhibit accelerated substitution rates and elevated GC content relative to other Cypriniformes, consistent with long-branch effects and compositional biases that can distort phylogenetic inference (5, 28, 30). Such elevated rates indicate heterogeneous molecular evolution in progenetic miniatures, potentially reflecting differences in life history, genome architecture, or metabolic constraints linked to extreme miniaturization and developmental truncation (134, 135). Nevertheless, in the case of *Danionella*, despite its long branch and elevated substitution rate, its phylogenetic position has been remarkably stable across all datasets and inference frameworks. *Danionella* was consistently recovered as the sister-group to the subfamily Danioninae within the family Danionidae with relatively strong sCFs and qCFs (Supplementary Fig. 27). This pattern corroborates previous morphological and molecular evidence for a close relationship between *Danionella* and other danionines (118, 136). This stable and well-supported close relationship between *Danionella* and *Danio rerio* establishes a valuable evolutionary framework for comparative developmental genomics and the study of miniaturization.

Our analyses provide the first molecular confirmation of the hypothesized sister-group relationship between *Fangfangia* and *Sundadanio*, a link previously inferred from morphological evidence. The consistent recovery of this relationship across multiple, independent datasets strengthens the phylogenetic hypothesis of this rare lineage.

Among phylogenetic relationships within Cypriniformes, the placement of progenetic miniatures has long remained contentious. Our analyses now strengthen the placement for *Danionella* and *Fangfangia*. However, the relationship of *Paedocypris* (Paedocyprididae) and *Sundadanio* + *Fangfangia* (Sundadanionidae) remain recalcitrant, exhibiting persistent incongruence across datasets and analytical frameworks. Interestingly, these two cases appear to reflect mechanistically distinct sources of conflict. For Sundadanionidae, balanced gene-tree support and very short internodes suggest that discordance is primarily biological in origin, most plausibly the result of ILS during a rapid early radiation at the base of Cyprinoidei. In contrast, the instability in the placement of Paedocyprididae appears to stem from extensive rate heterogeneity and GC-biased compositional similarity to outgroups, conditions that foster LBA rather than stochastic error or ILS alone.

These contrasting patterns underscore that both biological processes (e.g., ILS) and methodological artefacts (e.g., LBA) can yield superficially similar signatures of phylogenetic conflict within the same radiation, yet they require different methodological strategies for resolution. Even genome-scale datasets may fail to fully resolve such recalcitrant relationships (132), underscoring the inherent limitations of sequence-based phylogenomics (137). Future progress will likely depend on models incorporating recombination landscapes and novel genomic characters such as rare genomic changes or conserved gene synteny enabled by the growing availability of chromosome-level genome assemblies, together with new morphological datasets especially early developmental stages (137–140). However, sources of phylogenomic noise and systematic error remain largely underexplored in these novel approaches, which may introduce new analytical challenges even as they offer powerful complementary perspectives on genome evolution (137). These emerging tools nevertheless hold promise for disentangling the most persistent phylogenetic ambiguities within Cypriniformes and they have already been applied for the early branching of the cypriniform suborders providing novel evolutionary hypotheses (141).

Together, our findings establish a family-complete phylogenomic framework for Cypriniformes, strengthening the evolutionary placement of several rare and enigmatic lineages while identifying key recalcitrant groups that continue to challenge current analytical approaches. This framework not only strengthens the systematic foundation of the order but also provides a powerful reference for future research spanning evolutionary biology, and comparative genomics. By integrating these phylogenomic insights with emerging developmental, morphological, and genomic data, it will now be possible to explore the mechanisms driving genome evolution, diversification dynamics, and the repeated emergence of extreme miniaturization in freshwater fishes. As genomic resources and analytical models continue to advance, Cypriniformes will remain a central model for understanding how genome architecture, developmental constraints, and rapid speciation interact to shape one of the most diverse and evolutionarily dynamic vertebrate radiations on Earth.

## Acknowledgments

We would like to thank Pamela Nicholson and the staff of the Next Generation Sequencing (NGS) Platform, University of Bern, Switzerland, for sequencing support; Ulrich Schliewen (Bavarian State Collection of Zoology, Germany), Thomas Turner and Emily DeArmon (Museum of Southwestern Biology, USA), and Sven Kullander (Swedish Museum of Natural History, Sweden) for providing tissue samples; Conor Waldock for constructive comments on an earlier version of the manuscript; and Nadeela Hirimuthugoda (https://nadeela.weebly.com/illustrations.html) for providing the fish illustrations. HS further thanks Fernando Alda, Lily Hughes, Daniel Jeffries, Nicolás Mongiardino Koch, Zuyao Liu, and Fabrizia Ronco for their valuable guidance with specific analyses.

## Supplementary information

**Supplementary Fig. 1.**
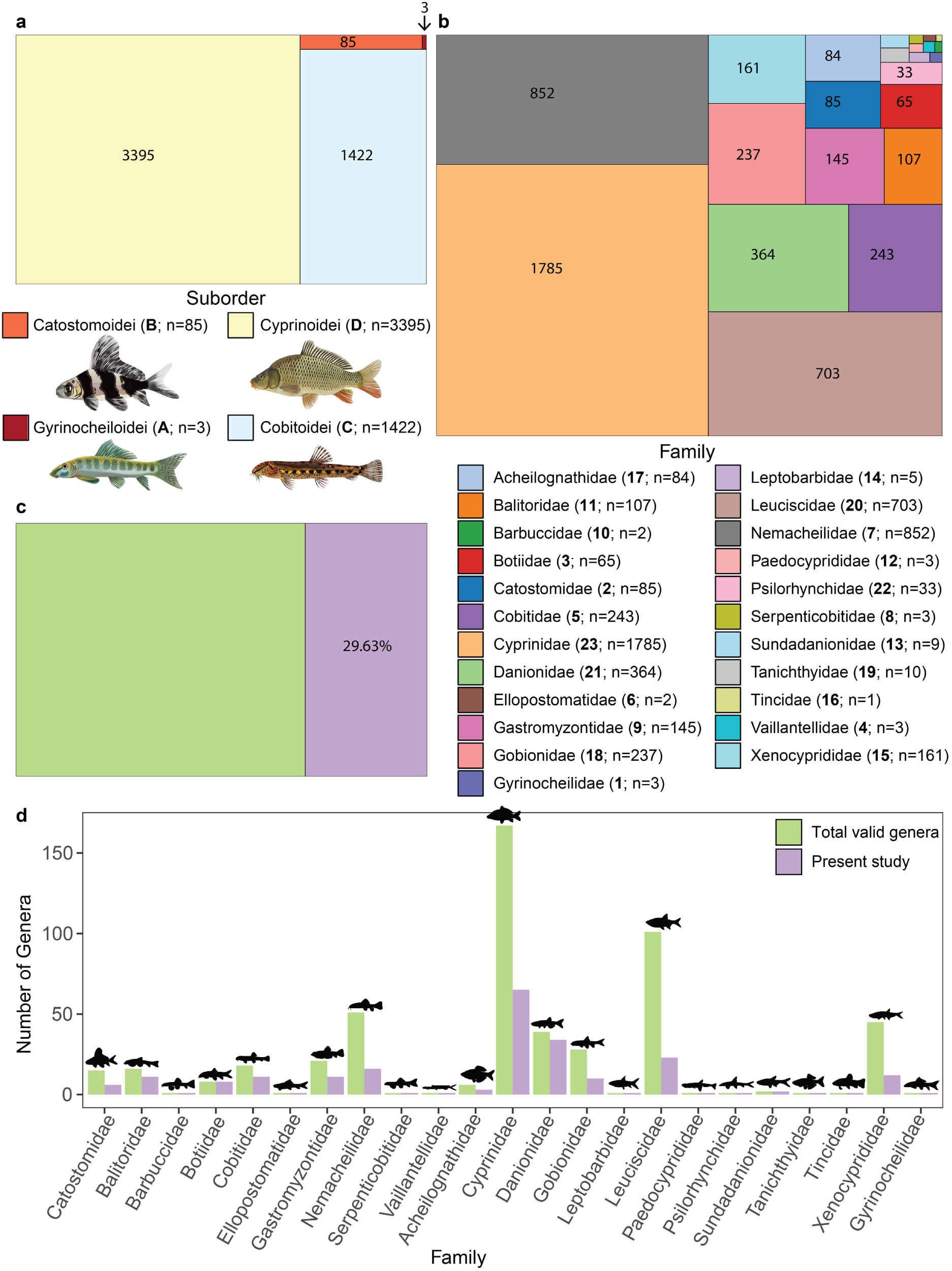
**(a)** Composition of Cypriniformes at the suborder level. **(b)** Composition at the family level. The number used for families in Fig.1 and the number of valid species based on the Catalog of Fishes is indicated within parenthesis. **(c)** Composition of genera included in this study as a proportion of the total number of valid Cypriniformes genera. **(d)** Breakdown of genera by family and their representation in this study. Rectangle sizes are proportional to the number of species in panels **(a)** and **(b)** and to the number of genera in panel **(c)**.

**Supplementary Fig. 2.**
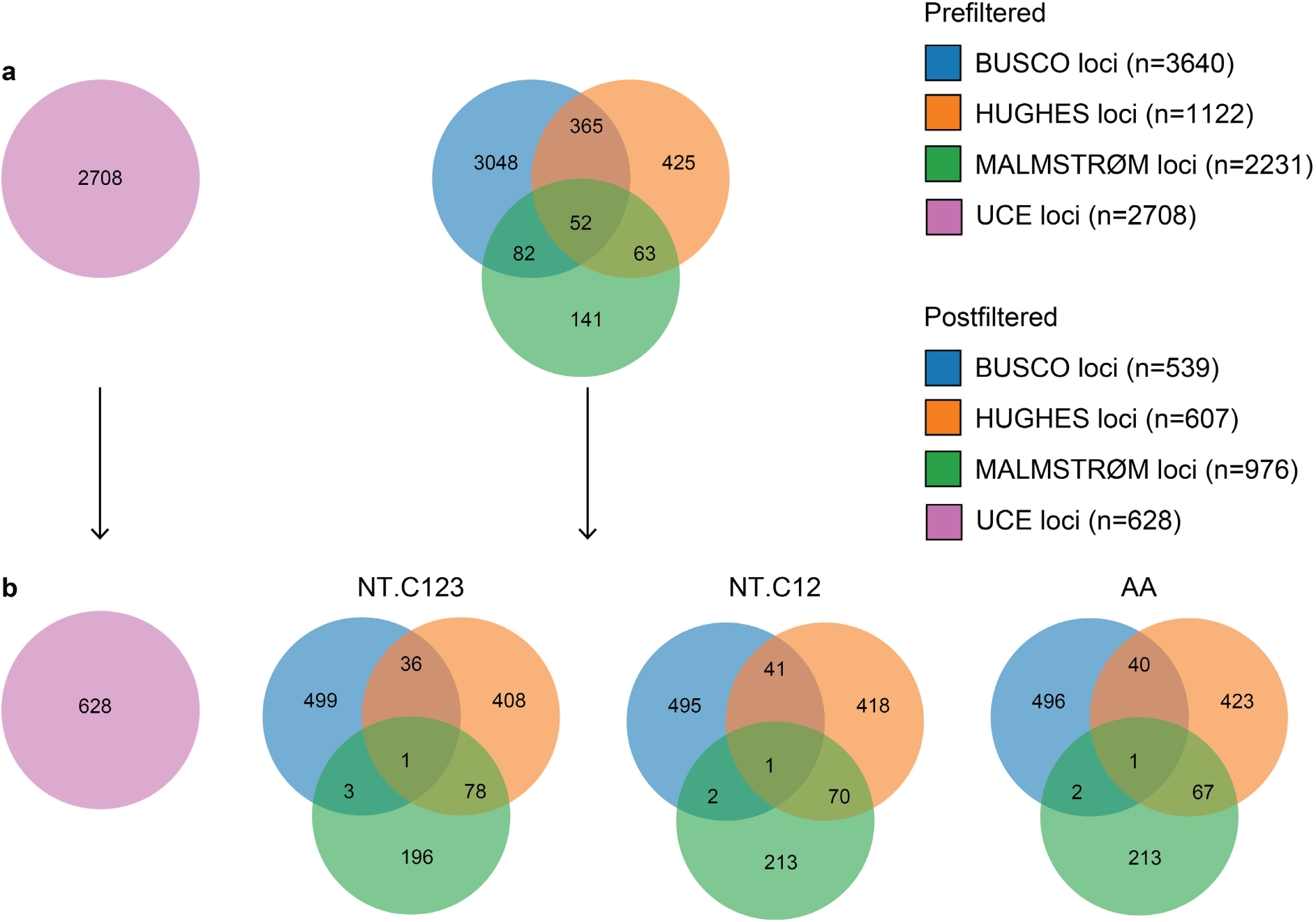
Number of UCE, BUSCO, HUGHES, and MALMSTRØM loci **(a)** before filtering and **(b)** after filtering for quality, orthology, and phylogenetic signal, which were then used in the subsequent phylogenomic analyses. Overlaps among BUSCO, HUGHES, and MALMSTRØM loci were identified using BLAST-based mapping of representative exon sequences to the zebrafish genome (*Danio rerio*, Ensembl GRCz11, release 114). When multiple exons matched the same Ensembl gene ID, they were counted as a single locus. Ensembl gene IDs could not be retrieved for 45 loci and alignments failed for 39 loci.

**Supplementary Fig. 3.**
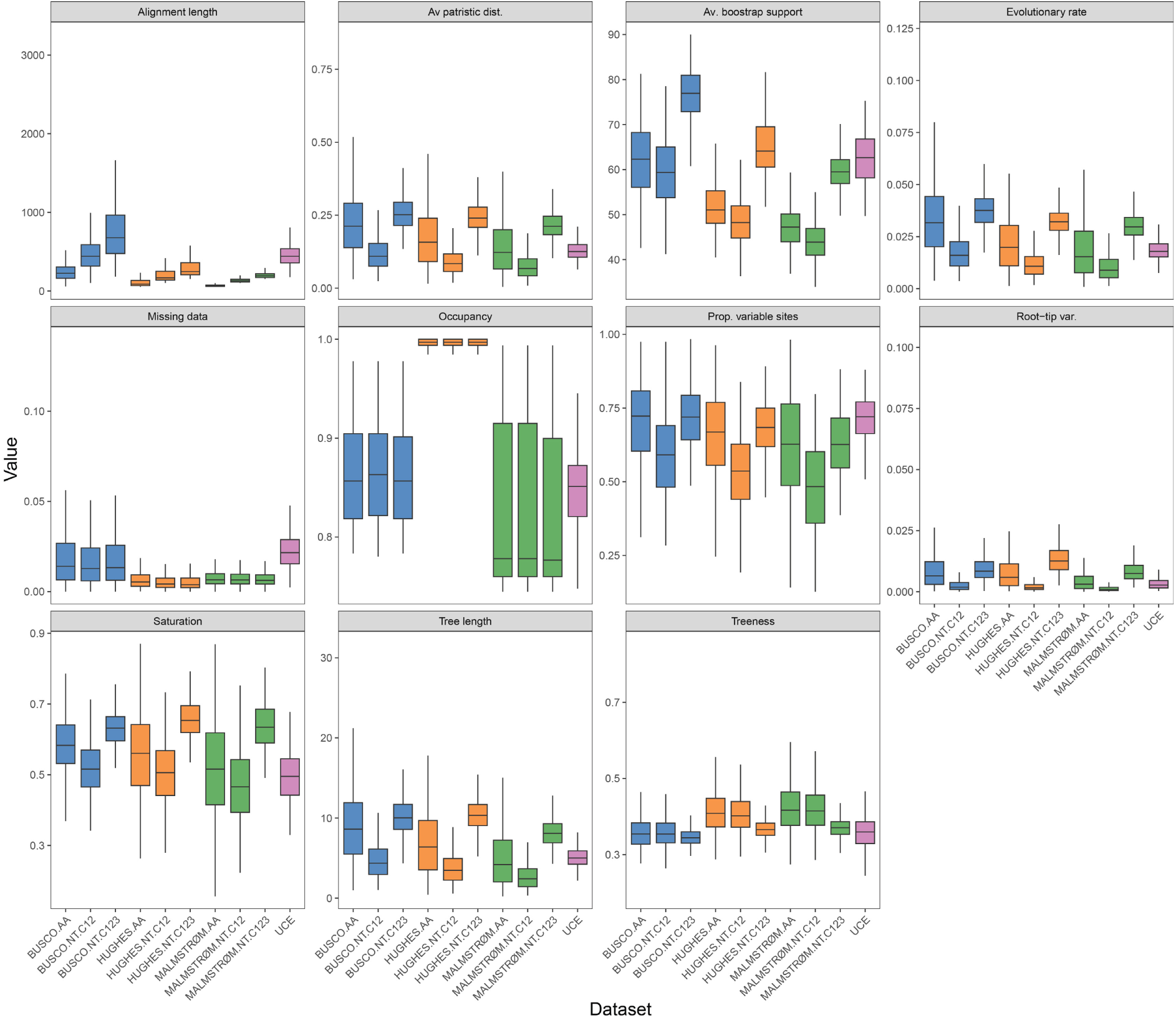
Comparison of gene properties estimated by genesortR for the different datasets used in the subsequent phylogenomic analyses.

**Supplementary Fig. 4.**
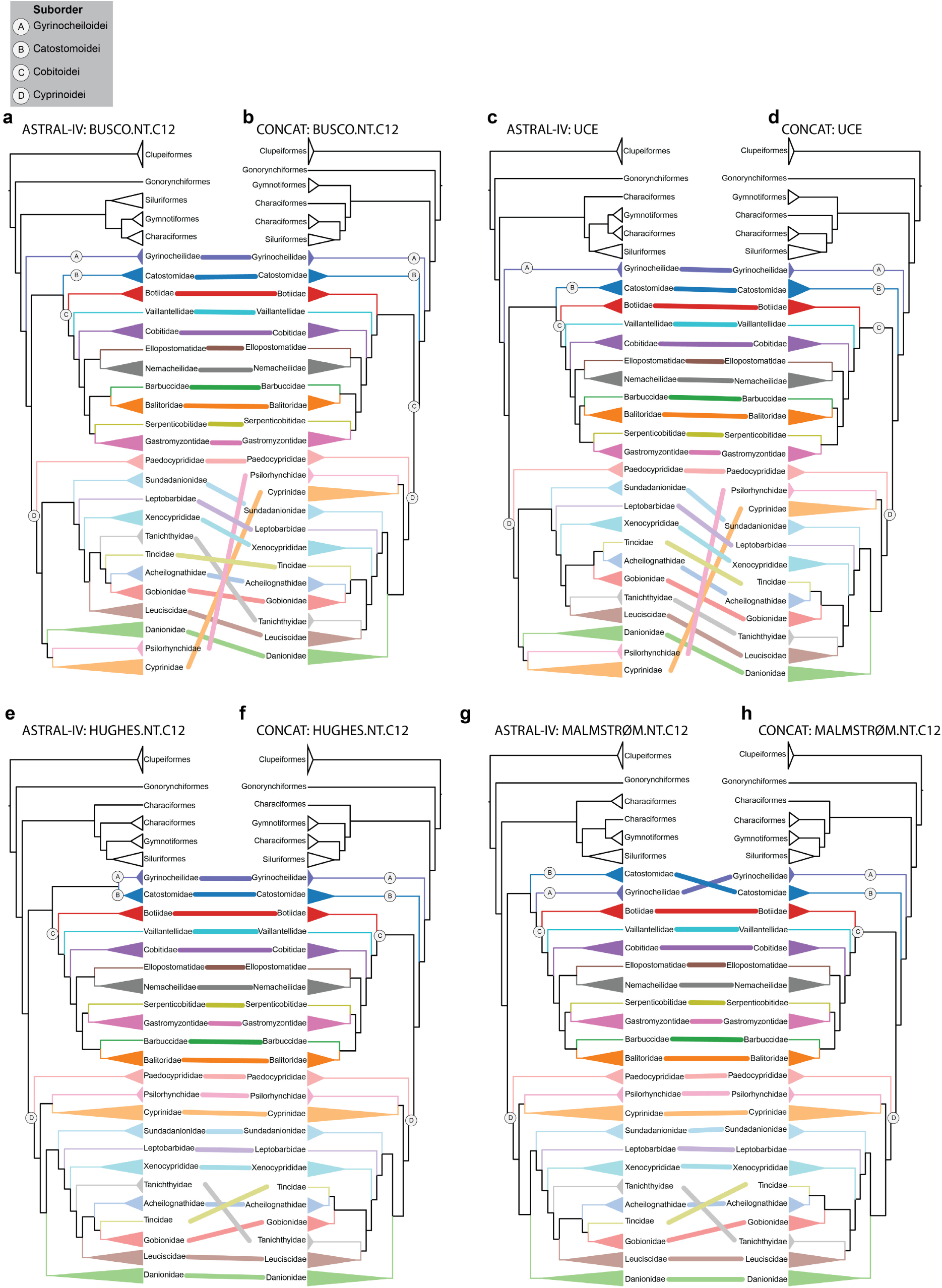
Comparison of family-level phylogenetic topologies obtained from **(a,c,e,g)** ASTRAL-IV species trees and **(b,d,f,h)** IQ-TREE2 concatenated phylogenies for different datasets. **(a,b)** BUSCO.NT.C12, **(c,d)** UCE, **(e,f)** HUGHES.NT.C12, and **(g,h)** MALMSTRØM.NT.C12.

**Supplementary Fig. 5.**
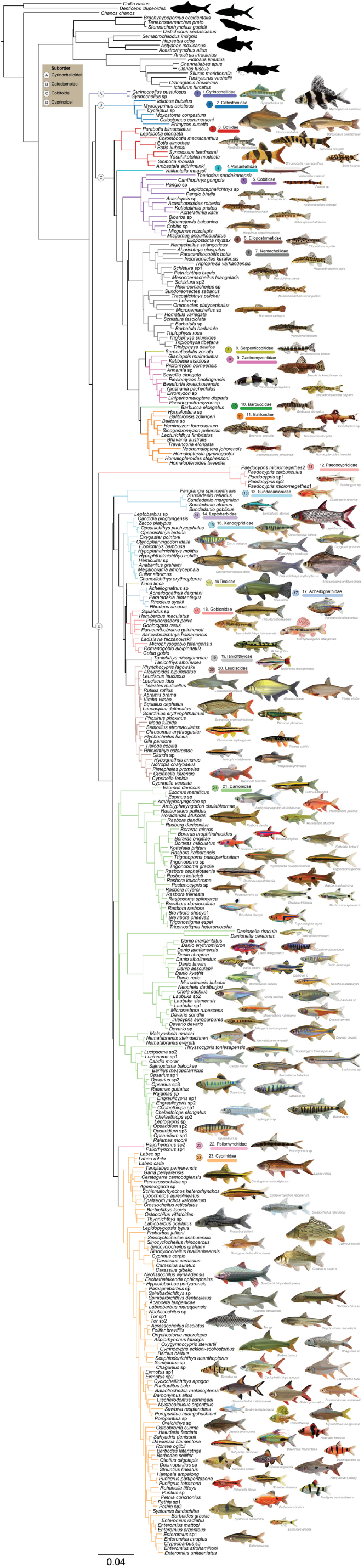
The expanded ASTRAL-IV species tree showing the hypothesis of phylogenomic relationships of Cypriniformes inferred from 335 species and 1,167 loci (total alignment length: 541,983 bp), based on combined first and second codon positions of BUSCO.NT.C12 and UCE nucleotide dataset. The numbers indicate the different Cypriniformes families.

**Supplementary Fig. 6.**
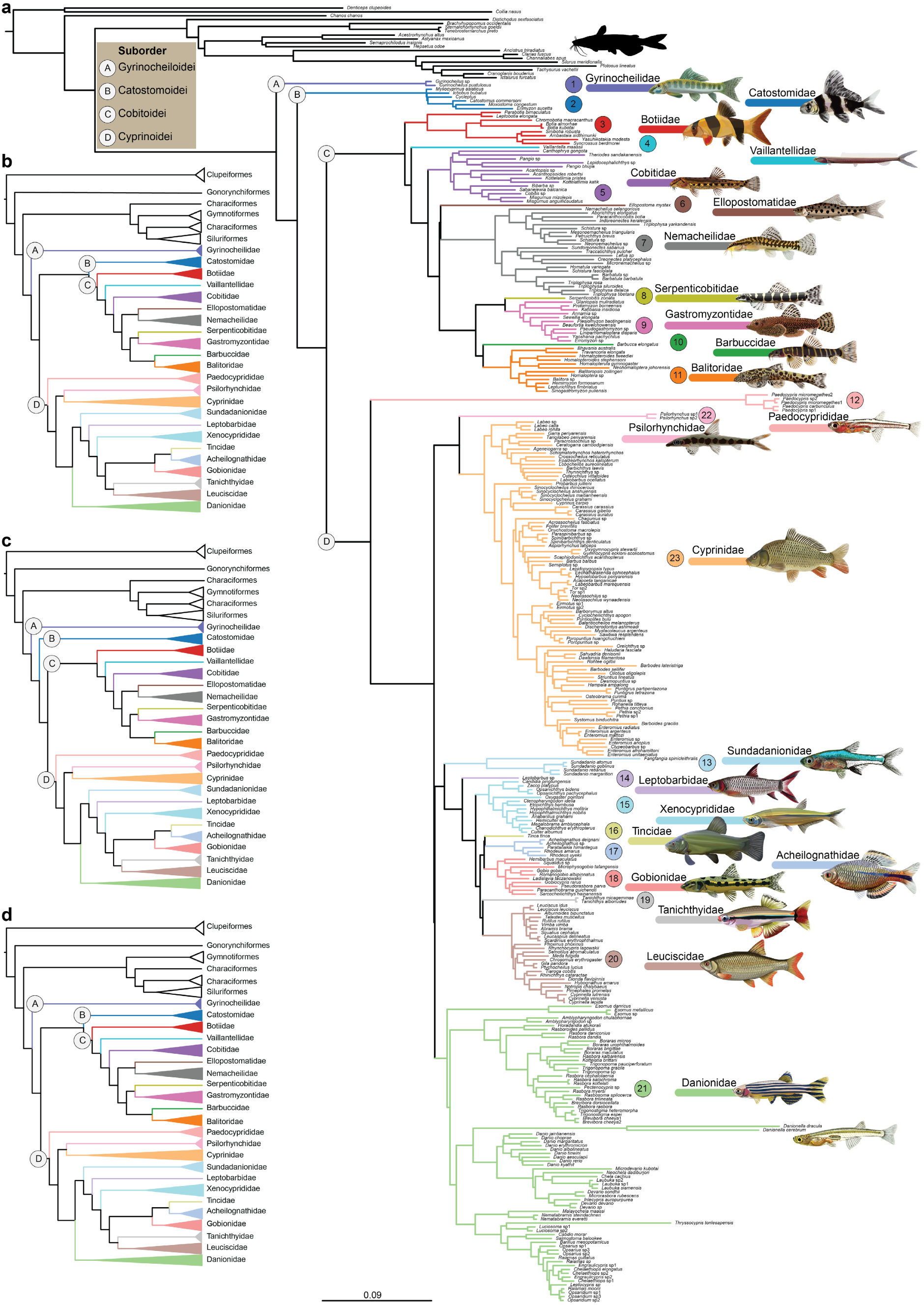
**(a)** Species tree of Cypriniformes inferred using the ROADIES pipeline under the ASTRAL-Pro3 framework from 335 species and 2,662 loci (total alignment length: 1769786 bp). Individual locus trees were estimated with maximum likelihood in RAxML-NG. The numbers indicate the different Cypriniformes families. **(b–d)** Simplified family-level topologies from three independent ROADIES runs based on 1,337, 2,662, and 1,352 loci, respectively. The first and third runs converged after four iterations, while the second run converged after five iterations.

**Supplementary Fig. 7.**
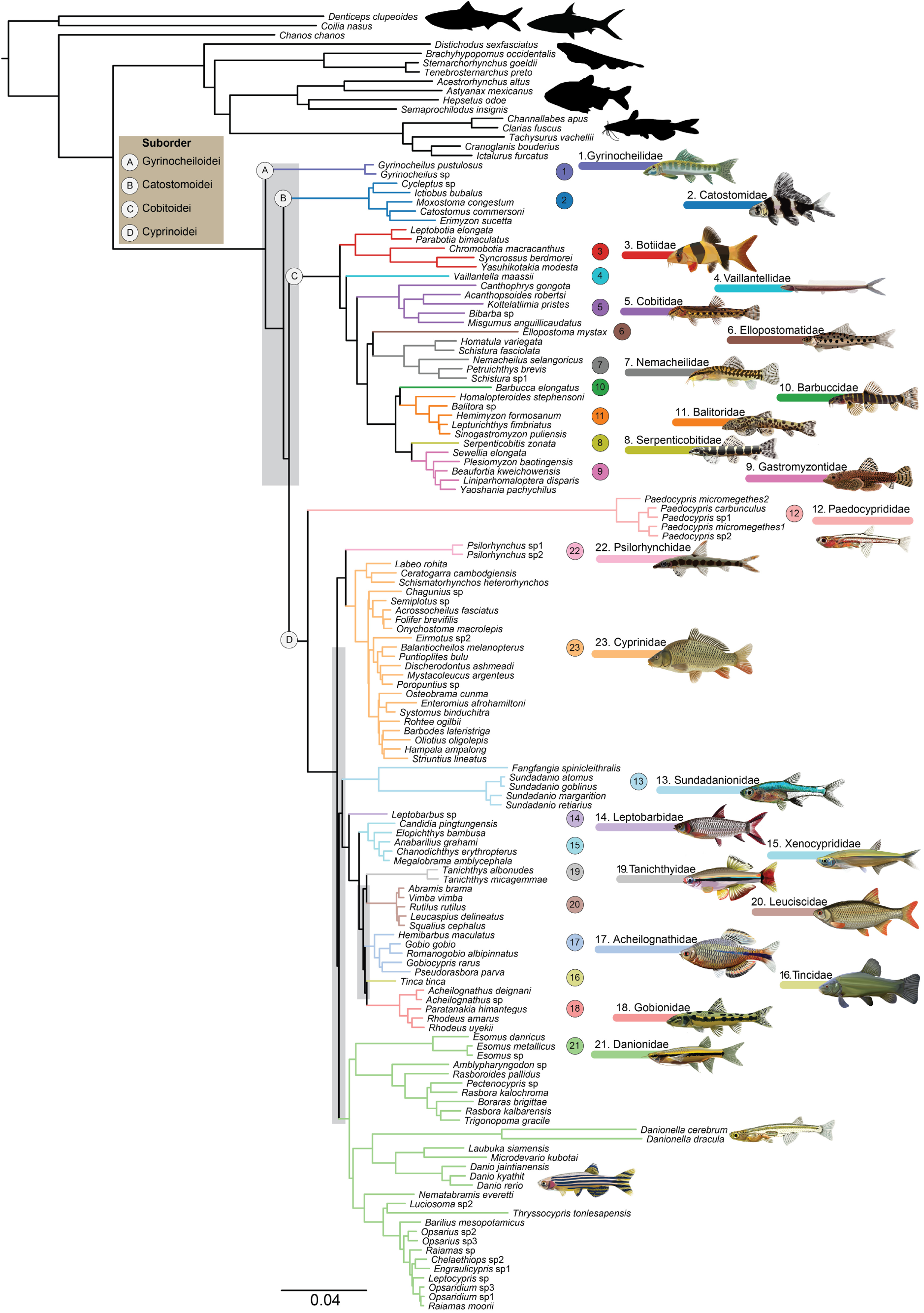
Phylogenomic relationships of Cypriniformes inferred from 140 species and 1,167 loci (total alignment length: 541,983 bp), based on combined first and second codon positions of BUSCO.NT.C12 and UCE nucleotide dataset using the GHOST model (GTR + H6) accounting for heterotachy in IQ-TREE2. The numbers indicate the different Cypriniformes families. The major conflicting zones of phylogenetic relationships is highlighted in grey.

**Supplementary Fig. 8.**
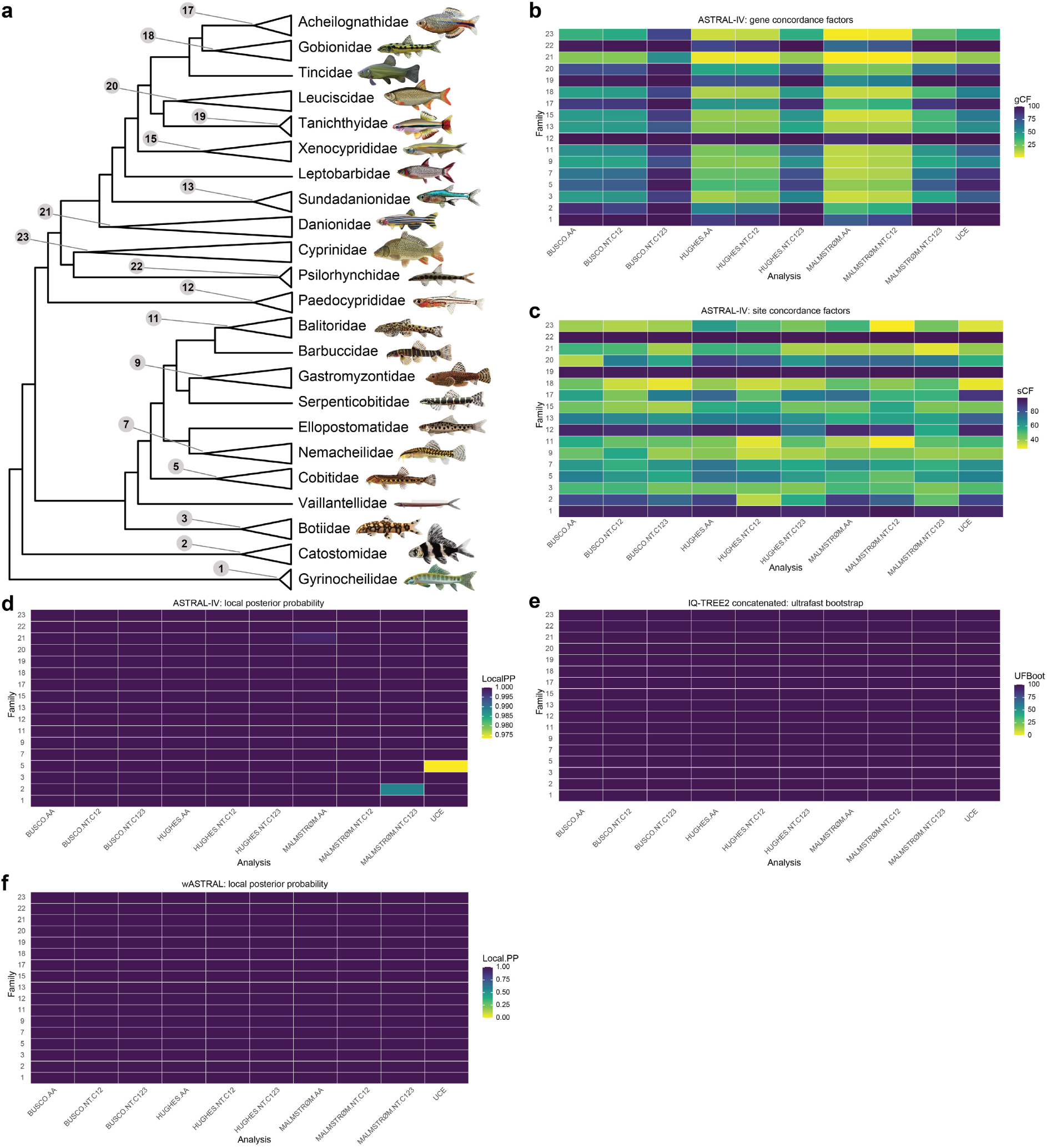
**(a)** Family-level monophyly of Cypriniformes recovered from different analyses. Topological variation is illustrated based on **(b)** gene and **(c)** site concordance factors. Statistical support is indicated by **(d)** local posterior probabilities for the ASTRAL-IV species tree, **(e)** ultrafast bootstrap values for the IQ-TREE2 concatenated phylogeny and **(f)** local posterior probabilities for the wASTRAL species tree. Increasing blue intensity indicate greater concordance or statistical support. The numbers in y-axis in **(b-f)** correspond to the monophyly of family shown in panel **(a)**.

**Supplementary Fig. 9.**
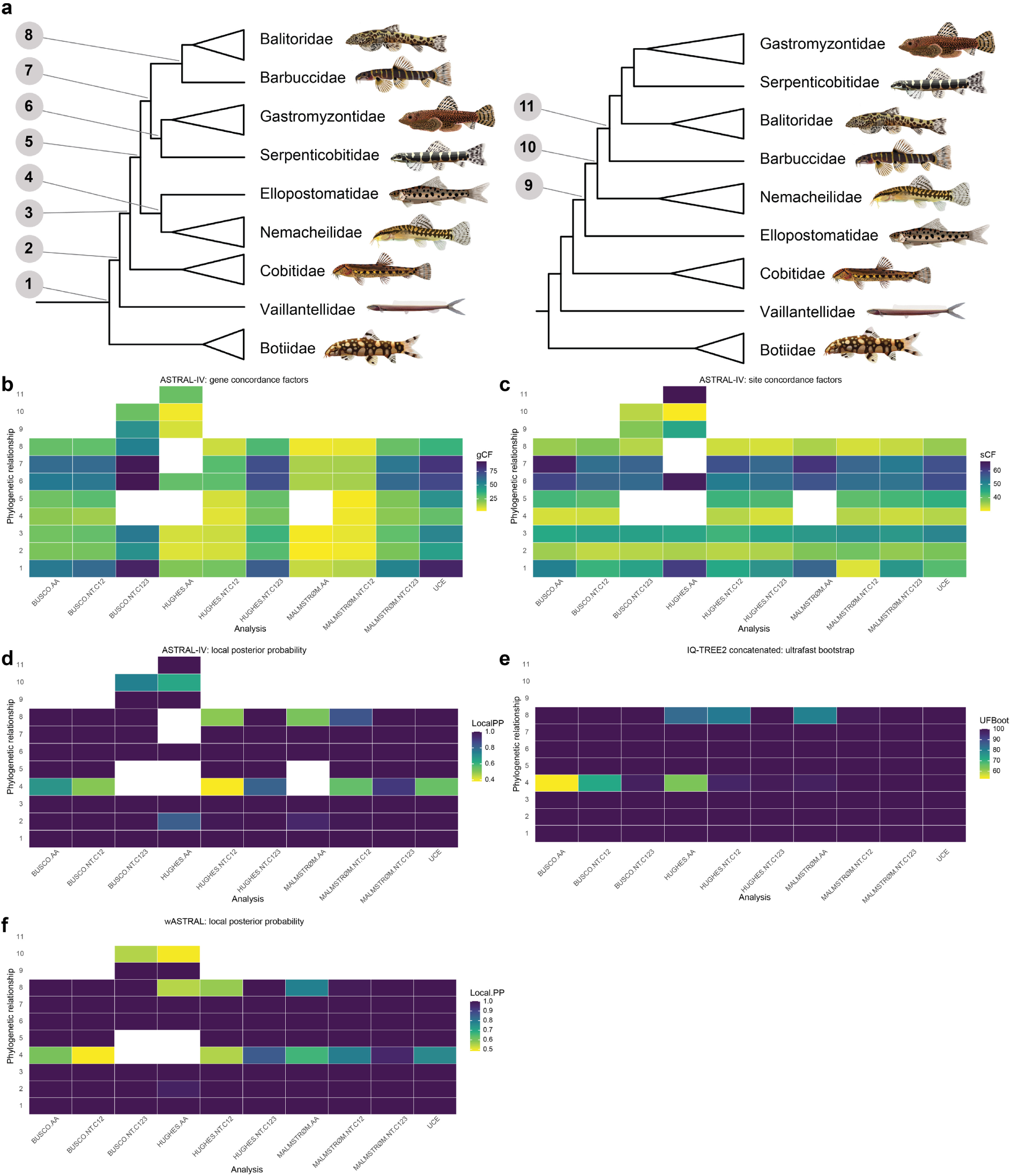
Phylogenetic relationships of Cobiotoidei recovered from different analyses. Less frequent and less plausible relationships are not shown. Topological variation is illustrated based on **(b)** gene and **(c)** site concordance factors. Statistical support is indicated by **(d)** local posterior probabilities for the ASTRAL-IV species tree, **(e)** ultrafast bootstrap values for the IQ-TREE2 concatenated phylogeny and **(f)** local posterior probabilities for the wASTRAL species tree. Increasing blue intensity indicate greater concordance or statistical support. The numbers in y-axis in **(b-f)** correspond to the phylogenetic relationships shown in panel **(a)**.

**Supplementary Fig. 10.**
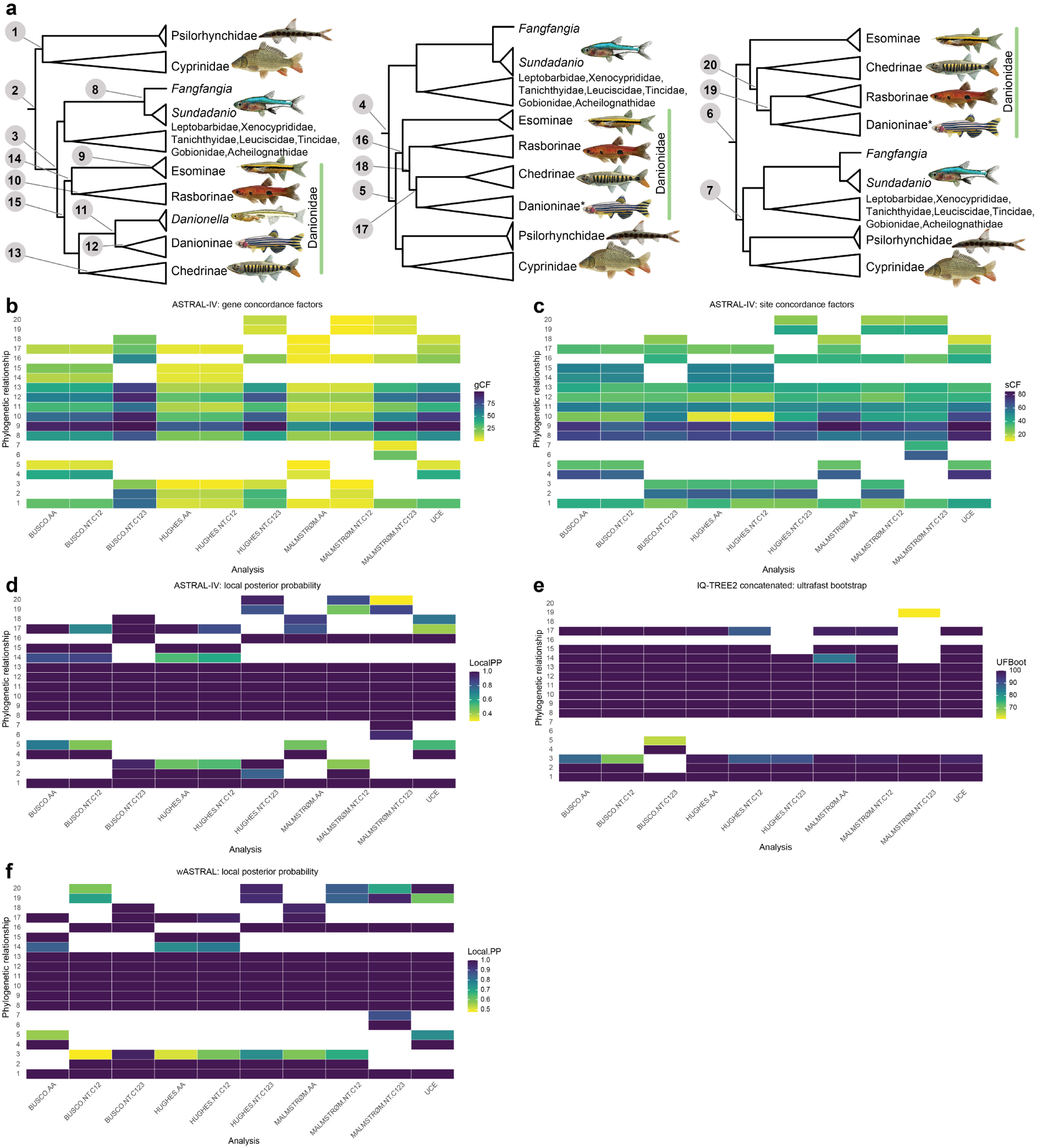
Phylogenetic relationships among Psilorhynchidae, Cyprinidae, Sundadanionidae, and Danionidae of Cyprinoidei recovered from different analyses. Less frequent and less plausible relationships are not shown. Topological variation is illustrated based on **(b)** gene and **(c)** site concordance factors. Statistical support is indicated by **(d)** local posterior probabilities for the ASTRAL-IV species tree, **(e)** ultrafast bootstrap values for the IQ-TREE2 concatenated phylogeny and **(f)** local posterior probabilities for the wASTRAL species tree. Increasing blue intensity indicate greater concordance or statistical support. The numbers in y-axis in **(b-f)** correspond to the phylogenetic relationships shown in panel **(a)**. An asterisk following Danioninae indicates that *Danionella* is also included within this clade.

**Supplementary Fig. 11.**
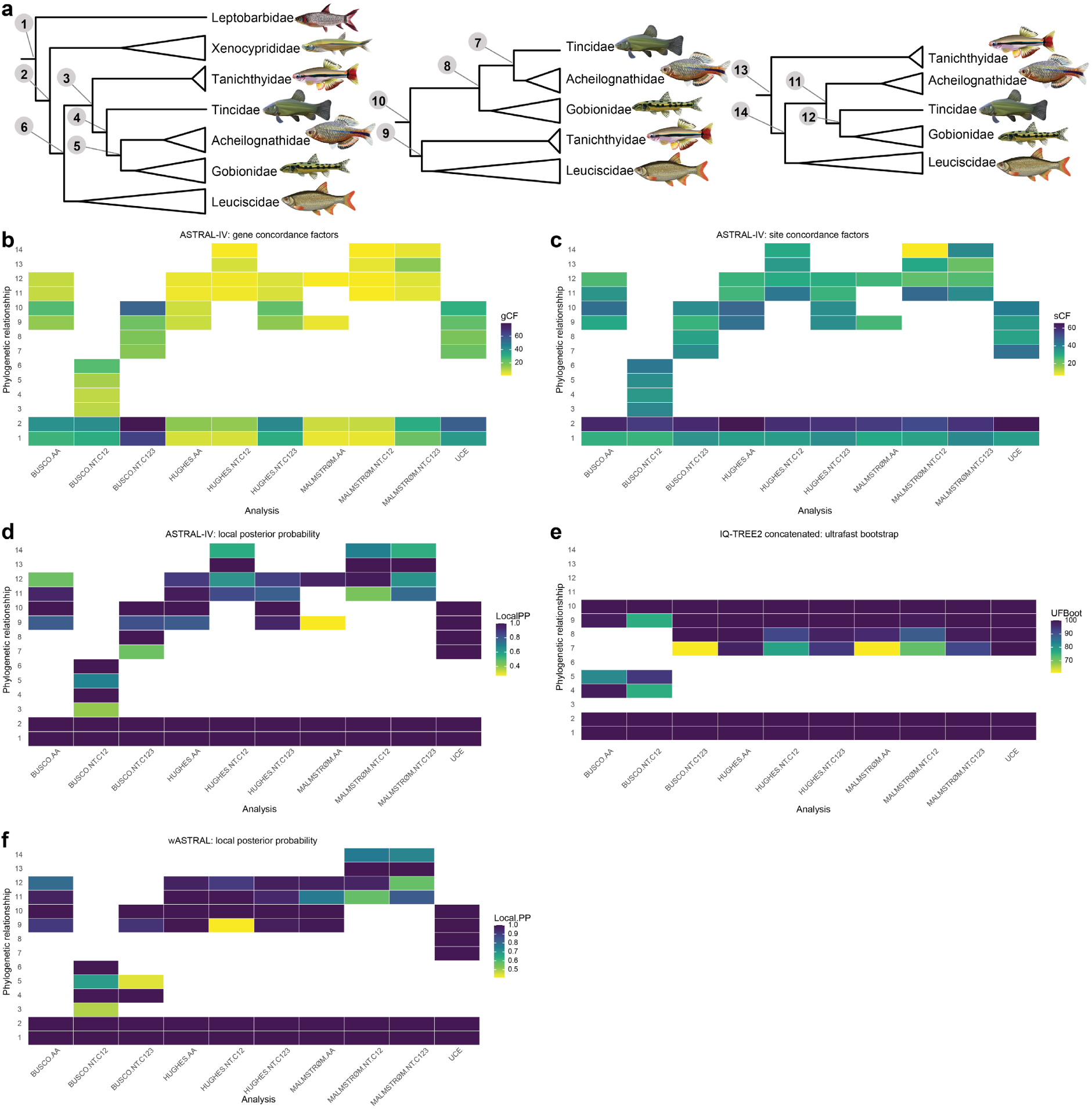
Phylogenetic relationships Leptobarbidae, Xenocyprididae, Tanichthyidae, Leuciscidae, Tincidae, Gobionidae, Acheilognathidae of Cyprinoidei recovered from different analyses. Less frequent and less plausible relationships are not shown. Topological variation is illustrated based on **(b)** gene and **(c)** site concordance factors. Statistical support is indicated by **(d)** local posterior probabilities for the ASTRAL-IV species tree, **(e)** ultrafast bootstrap values for the IQ-TREE2 concatenated phylogeny and **(f)** local posterior probabilities for the wASTRAL species tree. Increasing blue intensity indicate greater concordance or statistical support. The numbers in y-axis in **(b-f)** correspond to the phylogenetic relationships shown in panel **(a)**.

**Supplementary Fig. 12.**
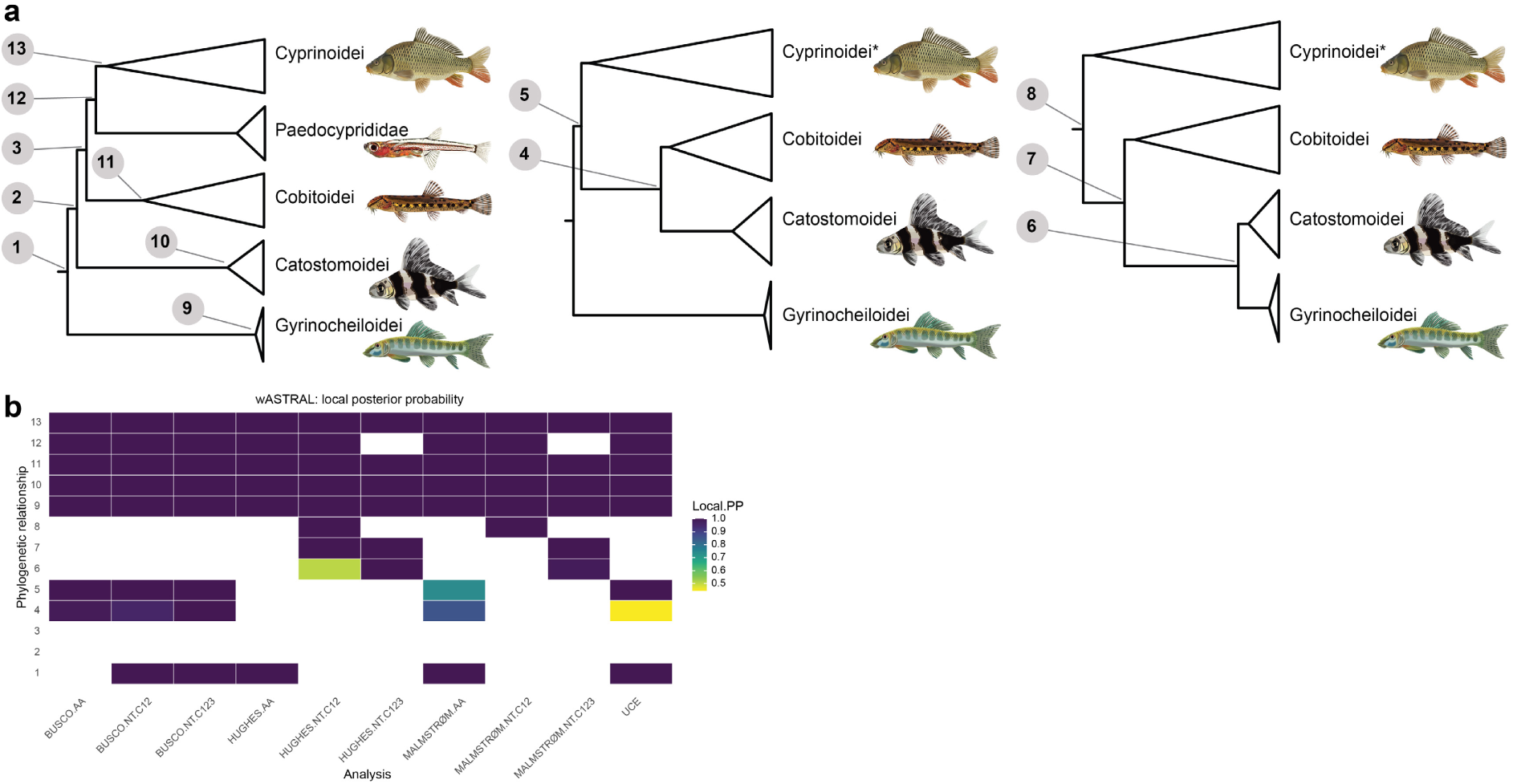
**(a)** Phylogenetic relationships among the suborders of Cypriniformes recovered from different analyses. Less frequent relationships are not shown. Statistical support is indicated by **(b)** local posterior probabilities for the wASTRAL species tree. Increasing blue intensity indicate greater concordance or statistical support. The numbers in y-axis in **(b)** correspond to the phylogenetic relationships shown in panel **(a)**. An asterisk following Cyprinoidei indicates that Paedocyprididae is also included within this clade.

**Supplementary Fig. 13.**
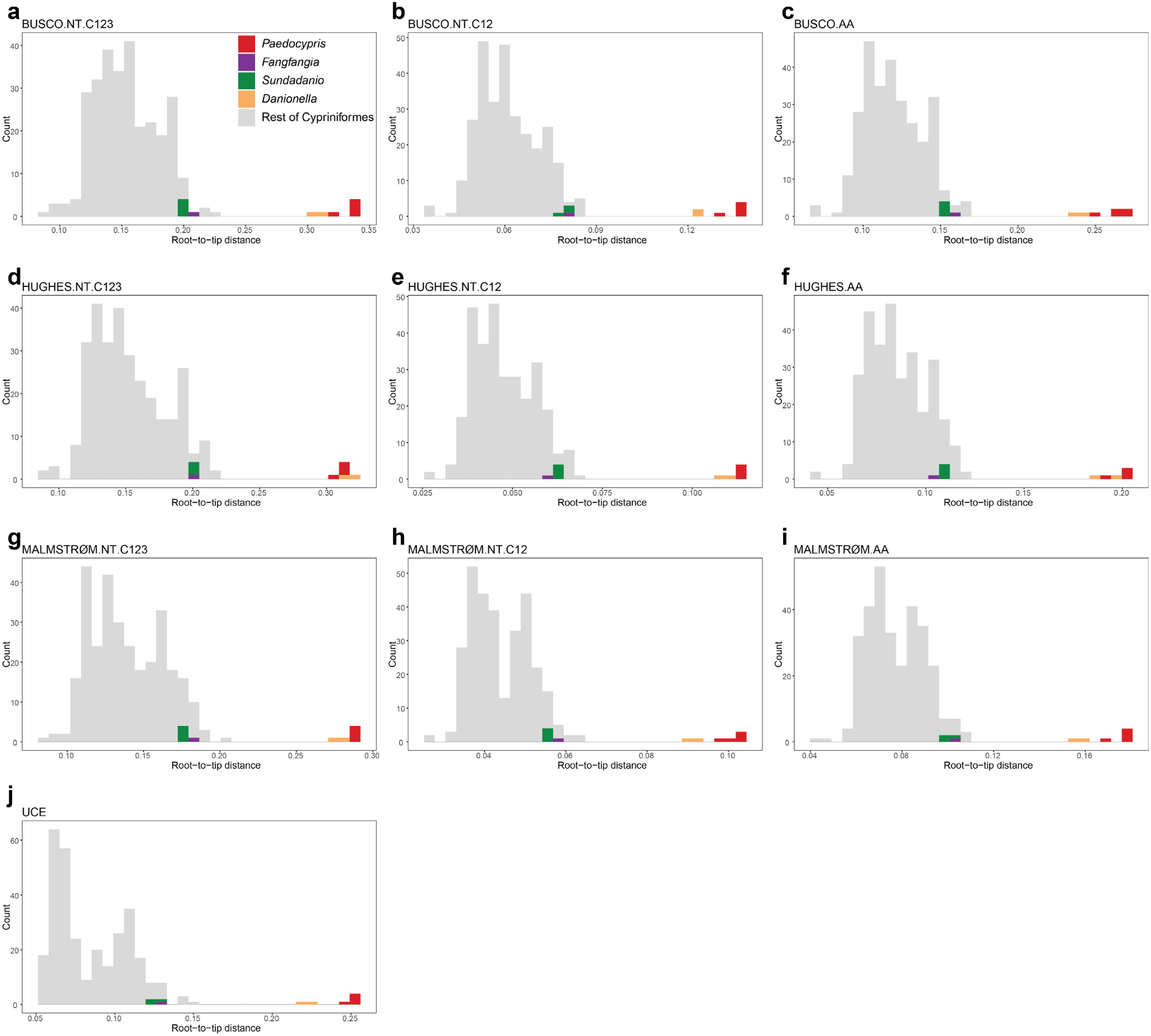
Comparison of root-to-tip distance in Cypriniformes for the IQ-TREE2 concatenated phylogenies inferred from different datasets: **(a)** BUSCO.NT.C123, **(b)** BUSCO.NT.C12, **(c)** BUSCO.AA, **(d)** HUGHES.NT.C123, **(e)** HUGHES.NT.C12, **(f)** HUGHES.AA, **(g)** MALMSTRØM.NT.C123, **(h)** MALMSTRØM.NT.C12, **(i)** MALMSTRØM.AA, and **(j)** UCE. Progenetic miniature genera are highlighted.

**Supplementary Fig. 14.**
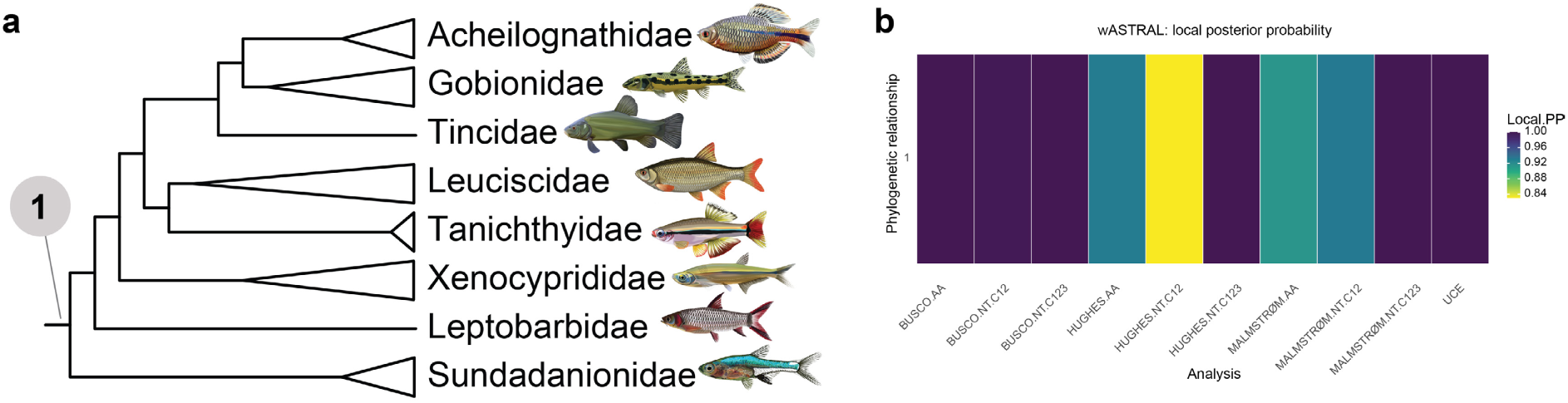
**(a)** Phylogenetic position of Sundadanionidae recovered from different analyses. Less frequent and less plausible relationships are not shown. Statistical support is indicated by **(b)** local posterior probabilities for the wASTRAL species tree. Increasing blue intensity indicate greater concordance or statistical support. The number in y-axis in **(b)** correspond to the phylogenetic relationship shown in panel **(a)**.

**Supplementary Fig. 15.**
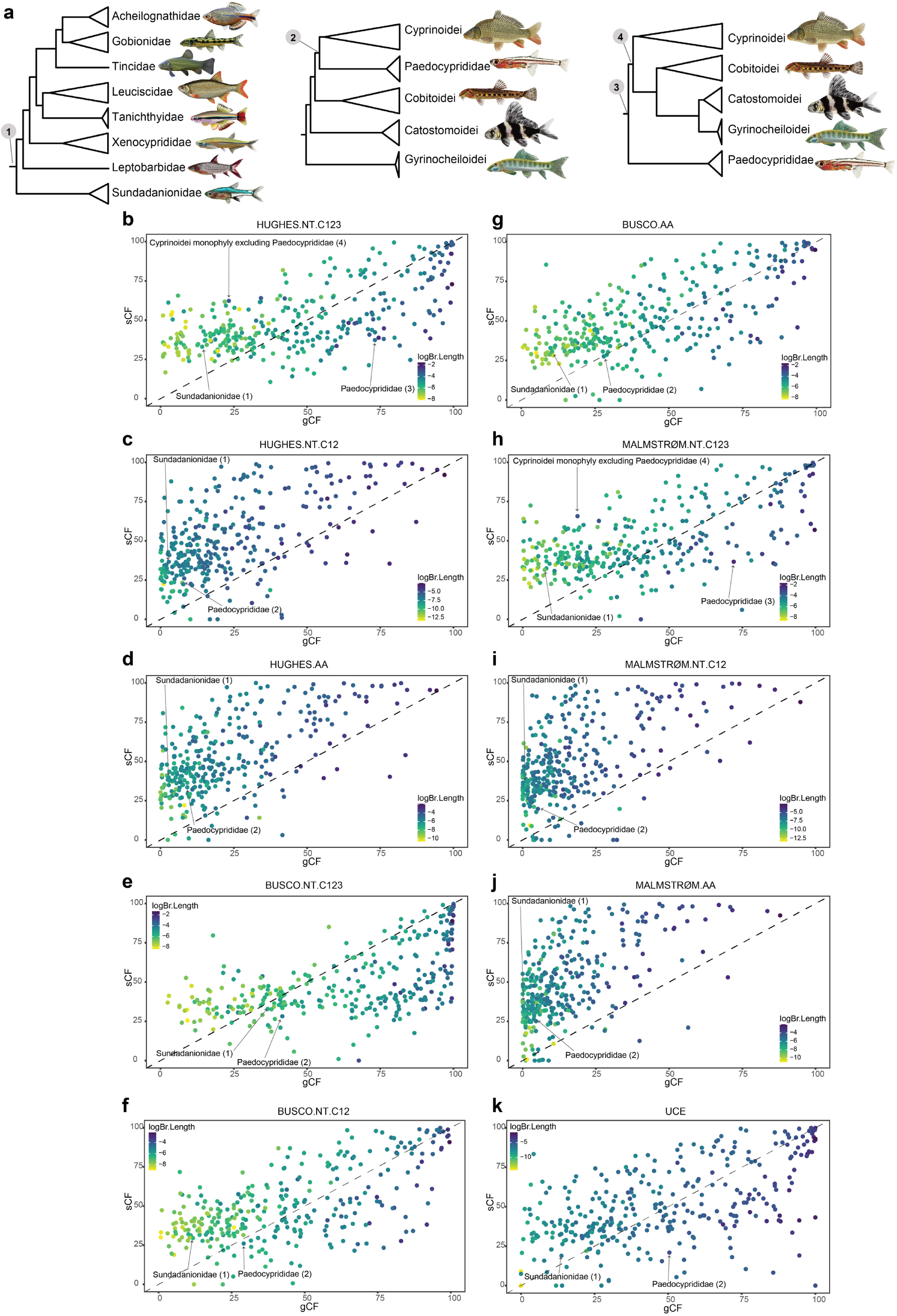
**(a)** Phylogenetic position of Sundadanionidae and Paedocyprididae recovered from different analyses. Panels **(b–k)** show the relationship between gene concordance factors, site concordance factors, and log-transformed branch lengths for ASTRAL-IV species trees inferred from different datasets: **(b)** HUGHES.NT.C123, **(c)** HUGHES.NT.C12, **(d)** HUGHES.AA, **(e)** BUSCO.NT.C123, **(f)** BUSCO.NT.C12, **(g)** BUSCO.AA, **(h)** MALMSTRØM.NT.C123, **(i)** MALMSTRØM.NT.C12, **(j)** MALMSTRØM.AA, and **(k)** UCE. Branches of interest are highlighted in each panel, and the numbers in parentheses correspond to the phylogenetic positions shown in panel **(a)**. Increasing blue intensity indicate longer log-transformed branch lengths.

**Supplementary Fig. 16.**
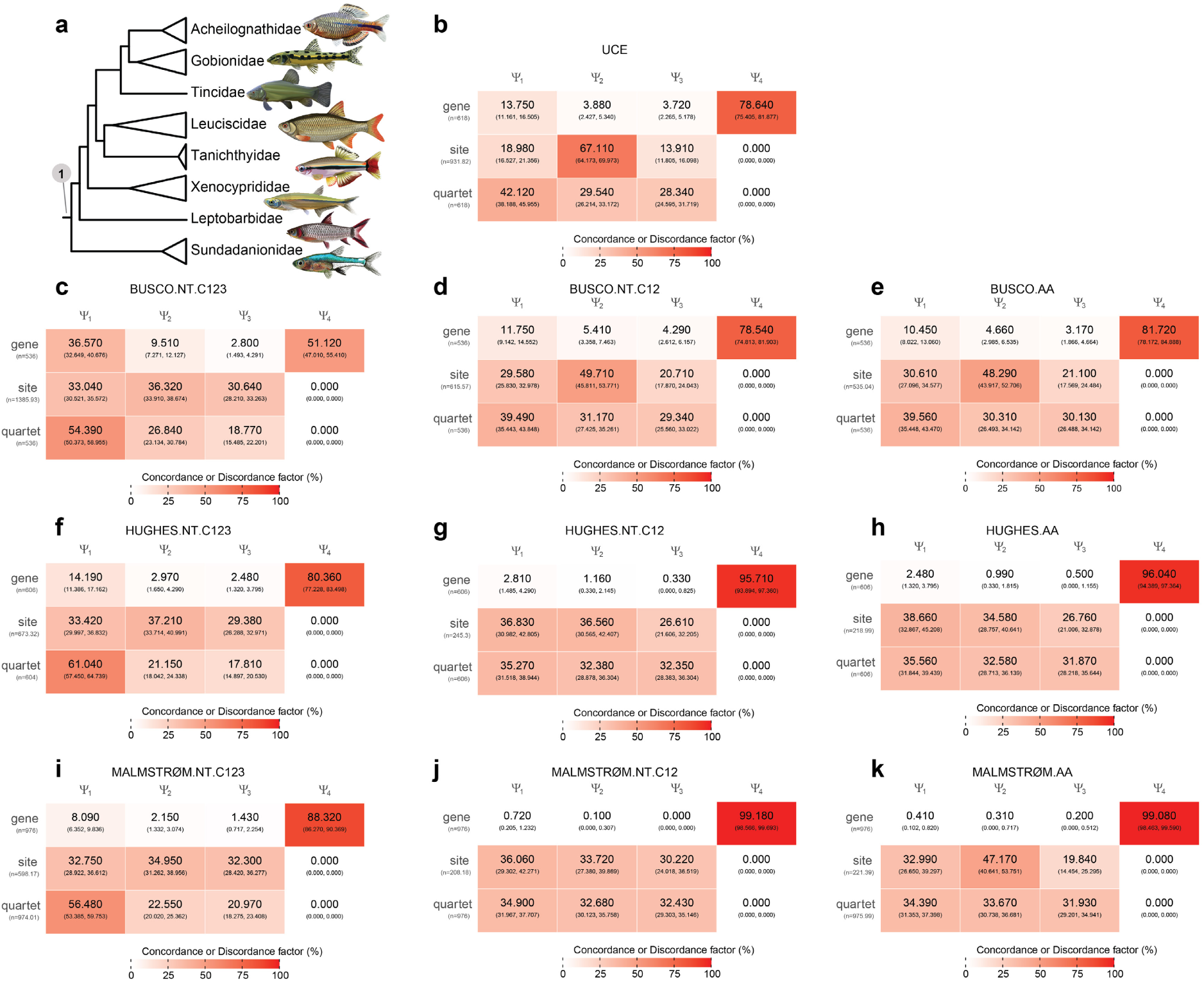
**(a)** Phylogenetic position of Sundadanionidae recovered from different analyses. Panels **(b–k)** show the gene, site, and quartet concordance vectors for ASTRAL-IV species trees inferred from different datasets: **(b)** UCE, **(c)** BUSCO.NT.C123, **(d)** BUSCO.NT.C12, **(e)** BUSCO.AA, **(f)** HUGHES.NT.C123, **(g)** HUGHES.NT.C12, **(h)** HUGHES.AA, **(i)** MALMSTRØM.NT.C123, **(j)** MALMSTRØM.NT.C12, and **(k)** MALMSTRØM.AA. Values in each cell are percentages, and the numbers in parentheses represent bootstrap-based 95% confidence intervals. Increasing red intensity indicate greater concordance or discordance. ψ₁ represent the concordance factor supporting the focal branch while ψ₂-ψ₄ represent the discordance factors.

**Supplementary Fig. 17.**
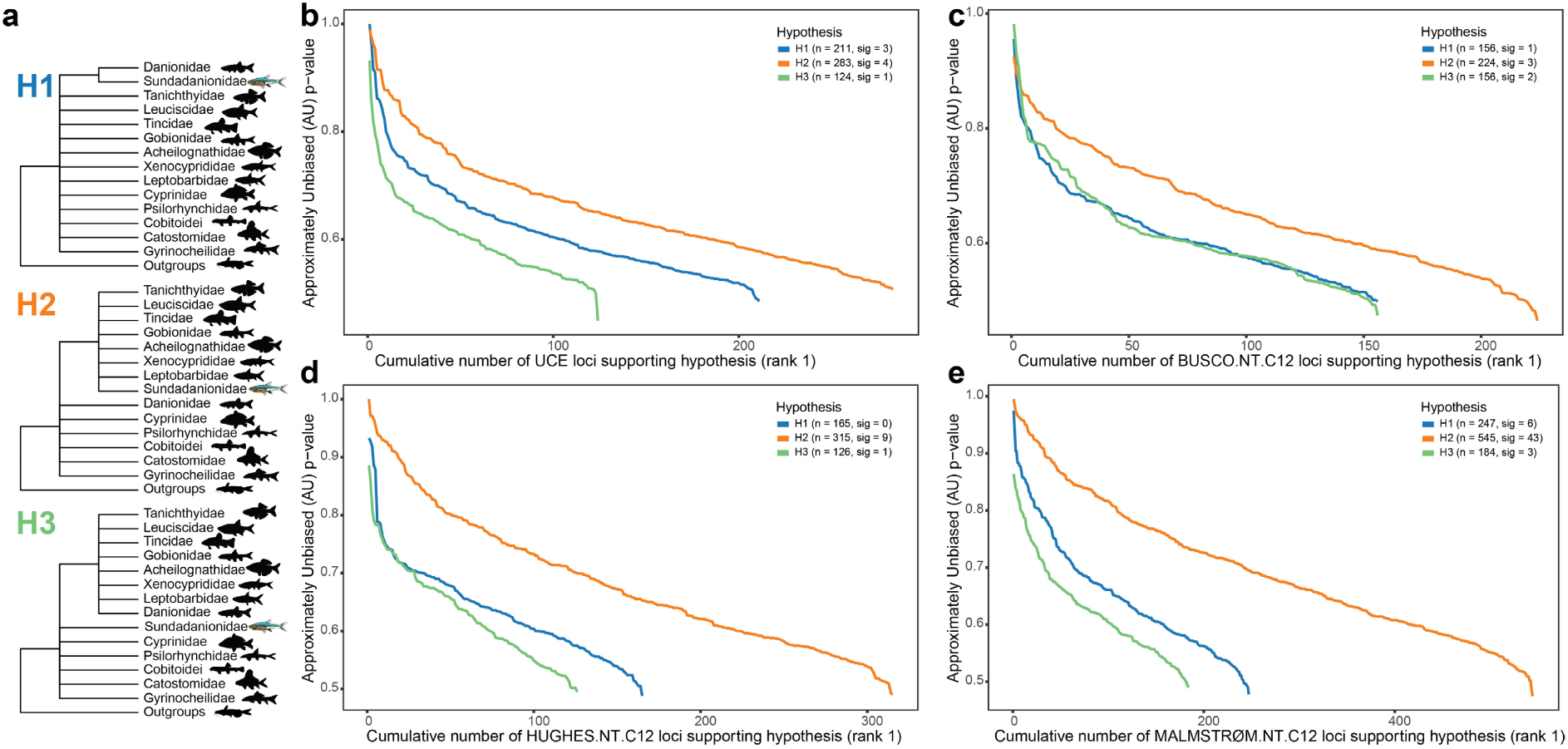
**(a)** Topologies of alternative hypotheses tested for the phylogenetic position of Sundadanionidae. Gene genealogy interrogation (GGI) applied to test alternative phylogenetic hypotheses for **(b)** UCE, **(c)** BUSCO.NT.C12, **(d)** HUGHES.NT.C12, **(e)** MALMSTRØM.NT.C12 datasets. Plotted lines show the cumulative number of loci (x-axis) supporting each topology as the highest-ranked hypothesis (rank 1), with AU test P-values on the y-axis.

**Supplementary Fig. 18.**
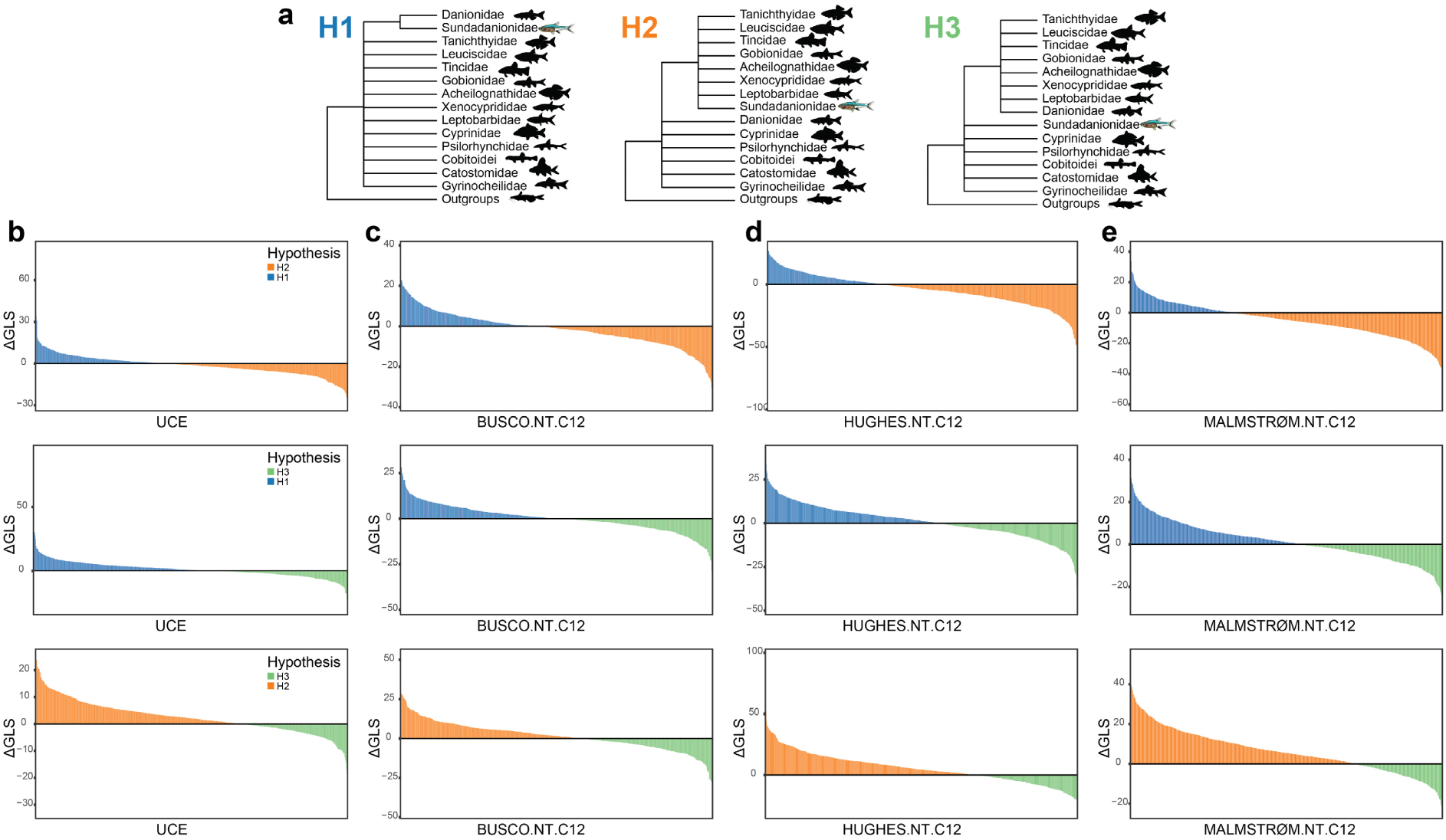
**(a)** Topologies of alternative hypotheses tested for the phylogenetic position of Sundadanionidae. Gene-wise phylogenetic signal for alternative hypotheses in **(b)** UCE, **(c)** BUSCO.NT.C12, **(d)** HUGHES.NT.C12, **(e)** MALMSTRØM.NT.C12 datasets, with the y-axis representing differences in gene-wise log-likelihood scores (ΔGLS).

**Supplementary Fig. 19.**
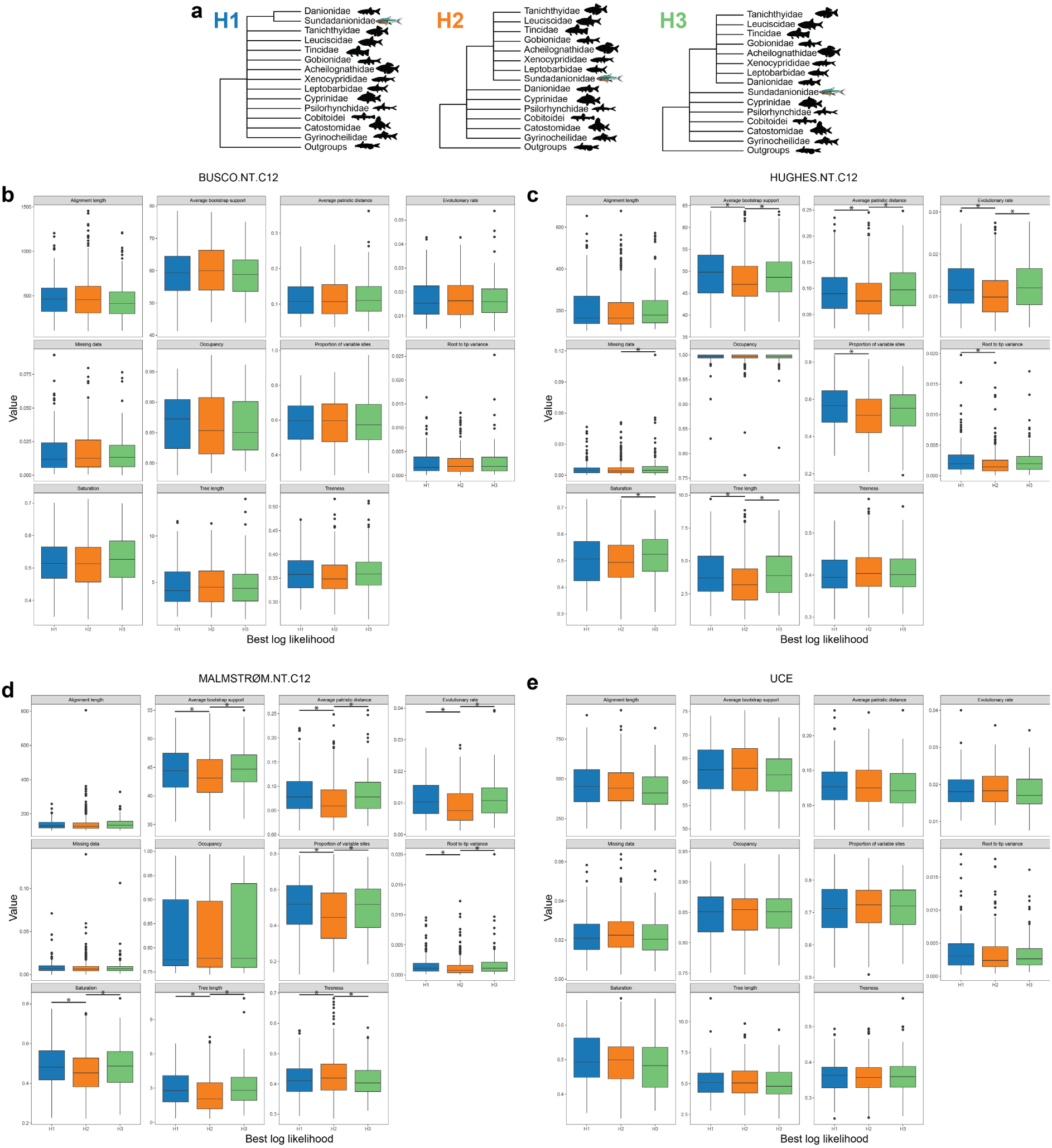
**(a)** Topologies of alternative hypotheses tested for the phylogenetic position of Sundadanionidae. Panels **(b– e)** compare loci properties estimated by genesortR for unconstrained **(b)** BUSCO.NT.C12, **(c)** HUGHES.NT.C12, **(d)** MALMSTRØM.NT.C12, and **(e)** UCE datasets. Each locus is assigned to H1, H2, or H3 based on the highest-ranked hypothesis according to log-likelihood scores for the constrained topologies. Asterisks indicate significant differences (p < 0.05) after pairwise t-tests, with false discovery rate (FDR) controlled using the Benjamini–Hochberg procedure.

**Supplementary Fig. 20.**
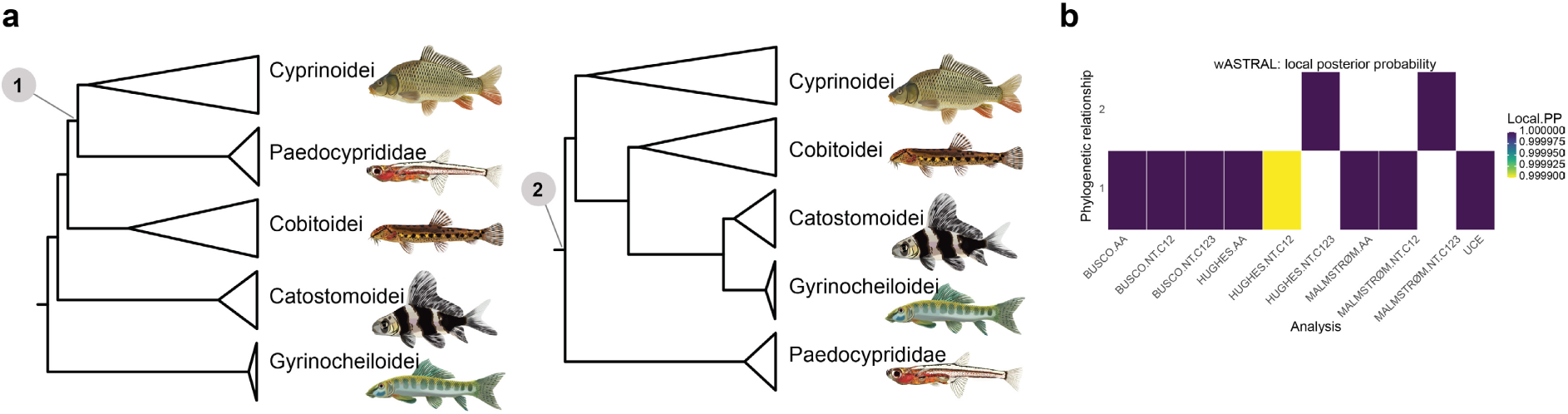
**(a)** Phylogenetic position of Paedocyprididae recovered from different analyses. Statistical support is indicated by **(b)** local posterior probabilities for the wASTRAL species tree. Increasing blue intensity indicate greater concordance or statistical support. The numbers in y-axis in **(b-f)** correspond to the phylogenetic relationship shown in panel **(a)**.

**Supplementary Fig. 21.**
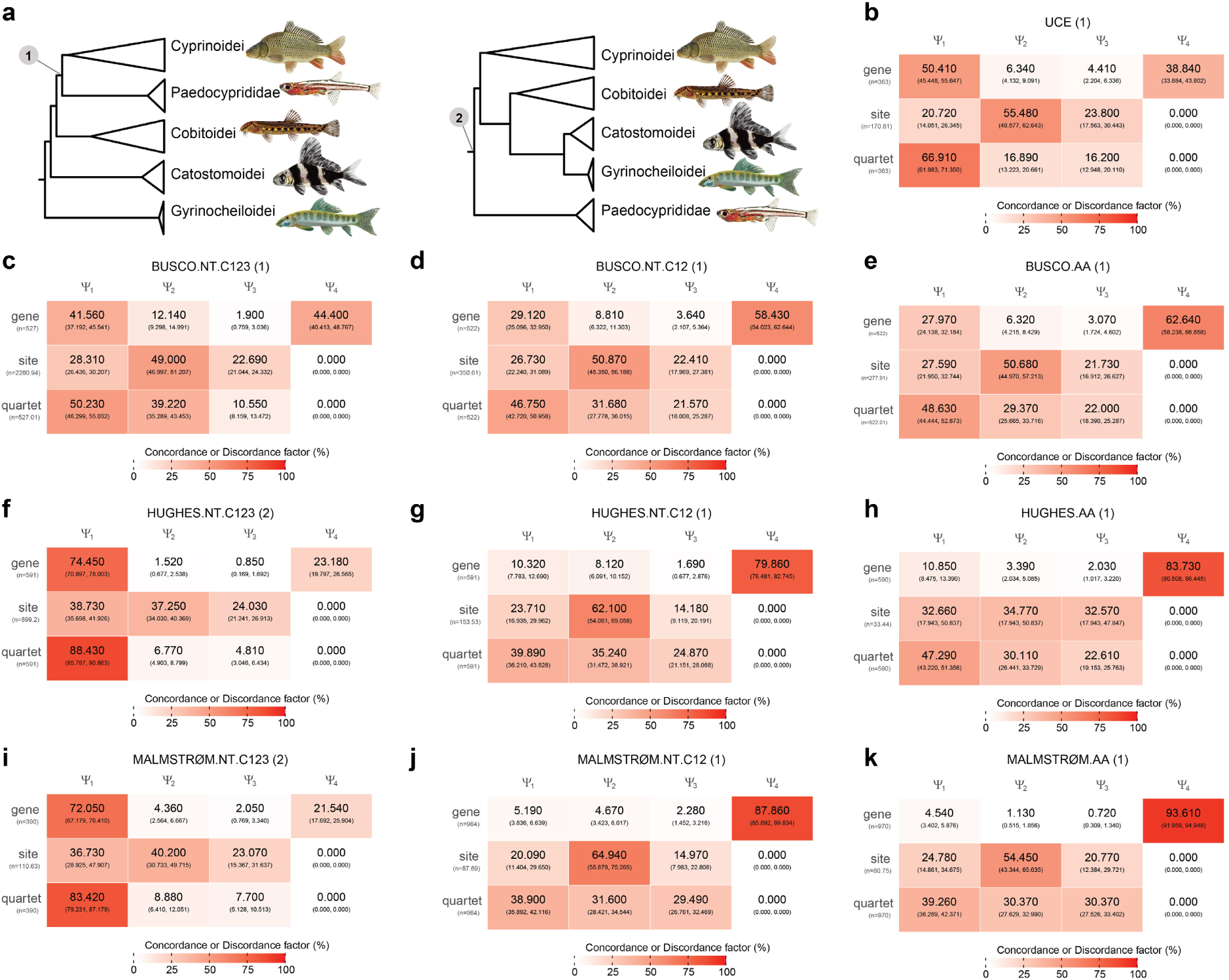
**(a)** Phylogenetic position of Paedocyprididae recovered from different analyses. Panels **(b–k)** show the gene, site, and quartet concordance vectors for ASTRAL-IV species trees inferred from different datasets: **(b)** UCE, **(c)** BUSCO.NT.C123, **(d)** BUSCO.NT.C12, **(e)** BUSCO.AA, **(f)** HUGHES.NT.C123, **(g)** HUGHES.NT.C12, **(h)** HUGHES.AA, **(i)** MALMSTRØM.NT.C123, **(j)** MALMSTRØM.NT.C12, and **(k)** MALMSTRØM.AA. The numbers in parentheses following each dataset name correspond to the phylogenetic relationship shown in panel **(a)**. Values in each cell are percentages, and the numbers in parentheses represent bootstrap-based 95% confidence intervals. Increasing red intensity indicate greater concordance or discordance. ψ₁ represent the concordance factor supporting the focal branch while ψ₂-ψ₄ represent the discordance factors.

**Supplementary Fig. 22.**
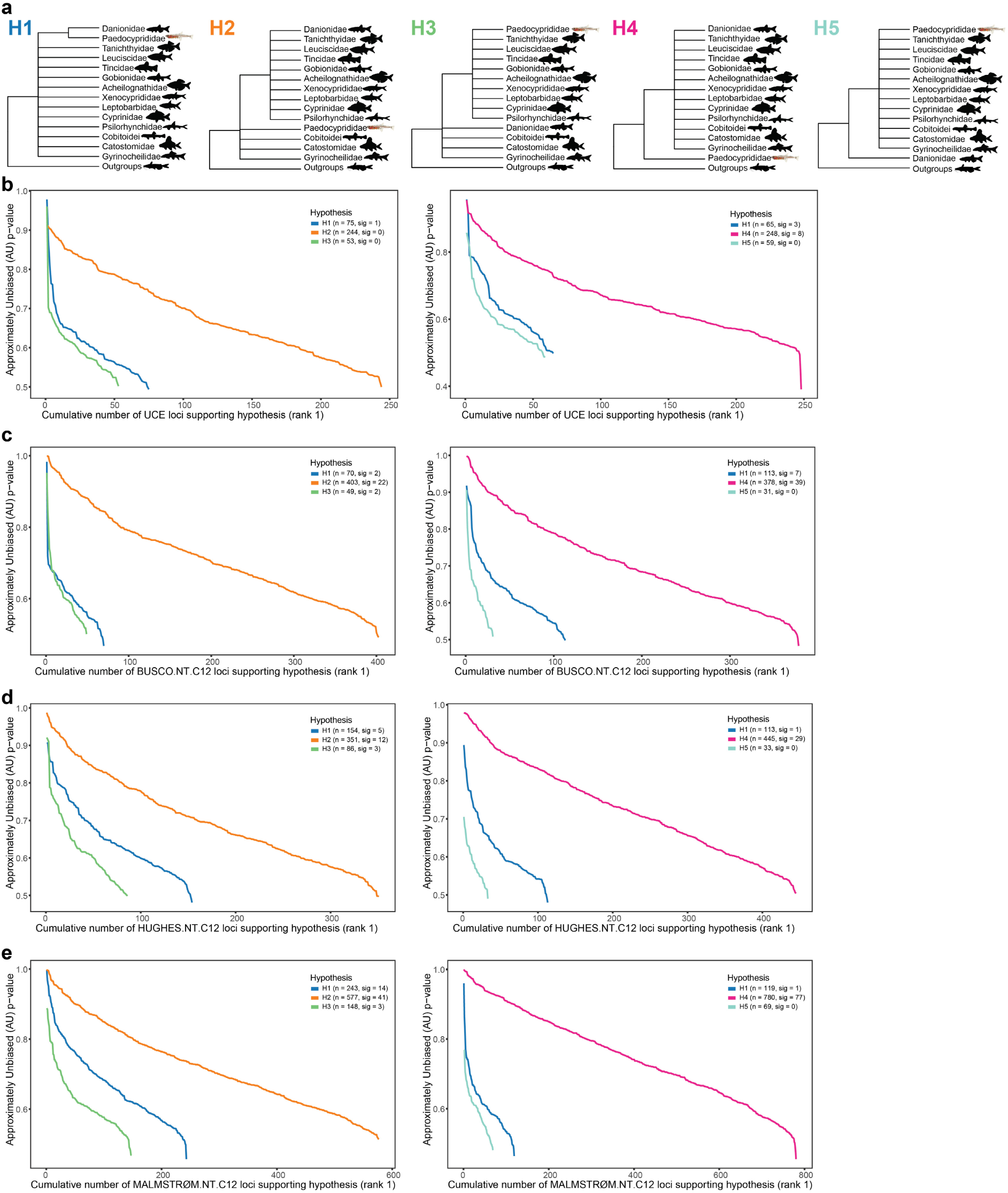
**(a)** Topologies of alternative hypotheses tested for the phylogenetic position of Paedocyprididae. Gene genealogy interrogation (GGI) applied to test alternative phylogenetic hypotheses for **(b)** UCE, **(c)** BUSCO.NT.C12, **(d)** HUGHES.NT.C12, **(e)** MALMSTRØM.NT.C12 datasets. Plotted lines show the cumulative number of loci (x-axis) supporting each topology as the highest-ranked hypothesis (rank 1), with AU test P-values on the y-axis.

**Supplementary Fig. 23.**
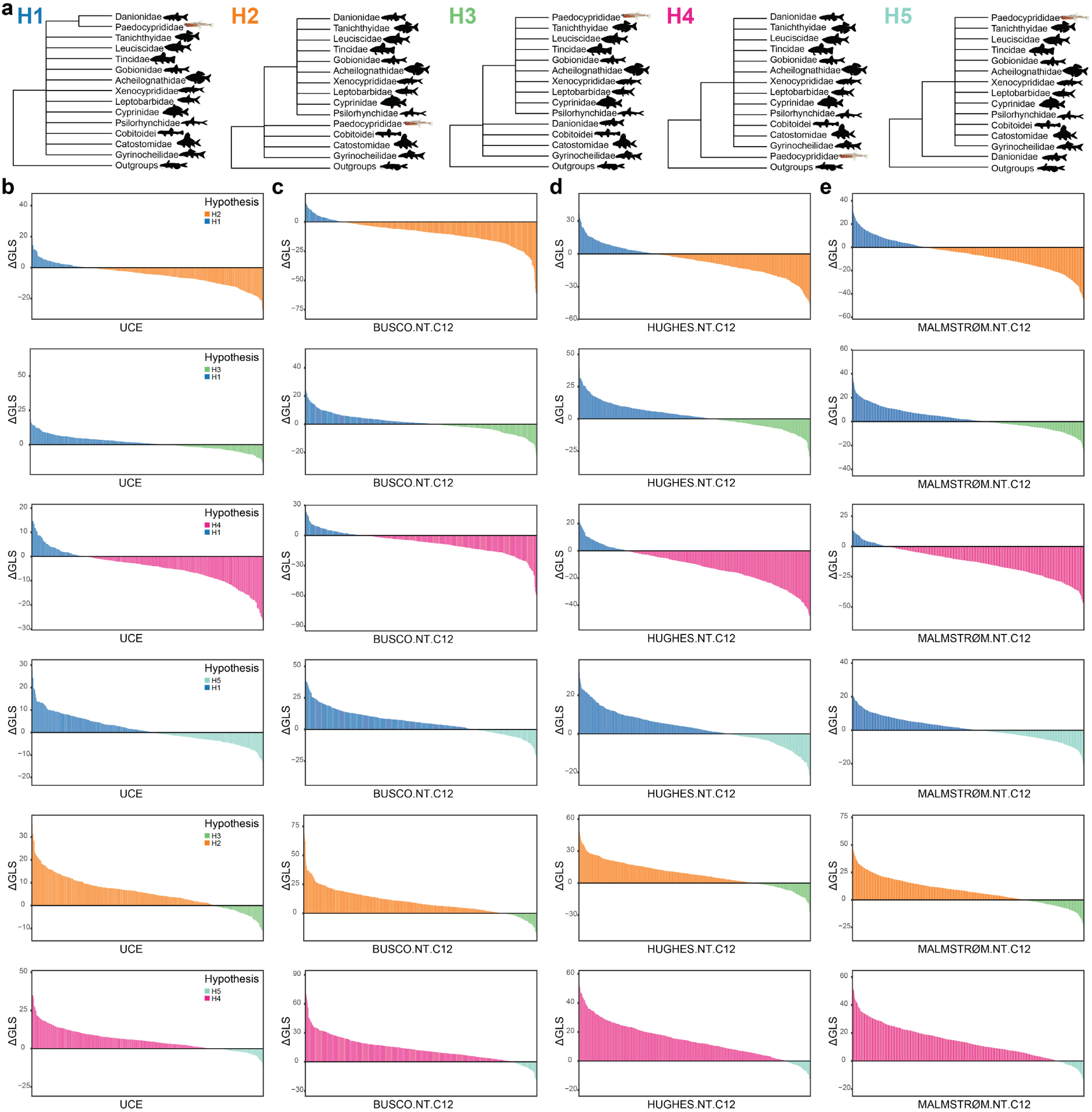
**(a)** Topologies of alternative hypotheses tested for the phylogenetic position of Paedocyprididae. Gene-wise phylogenetic signal for alternative hypotheses in **(b)** UCE, **(c)** BUSCO.NT.C12, **(d)** HUGHES.NT.C12, **(e)** MALMSTRØM.NT.C12 datasets, with the y-axis representing differences in gene-wise log-likelihood scores (ΔGLS).

**Supplementary Fig. 24.**
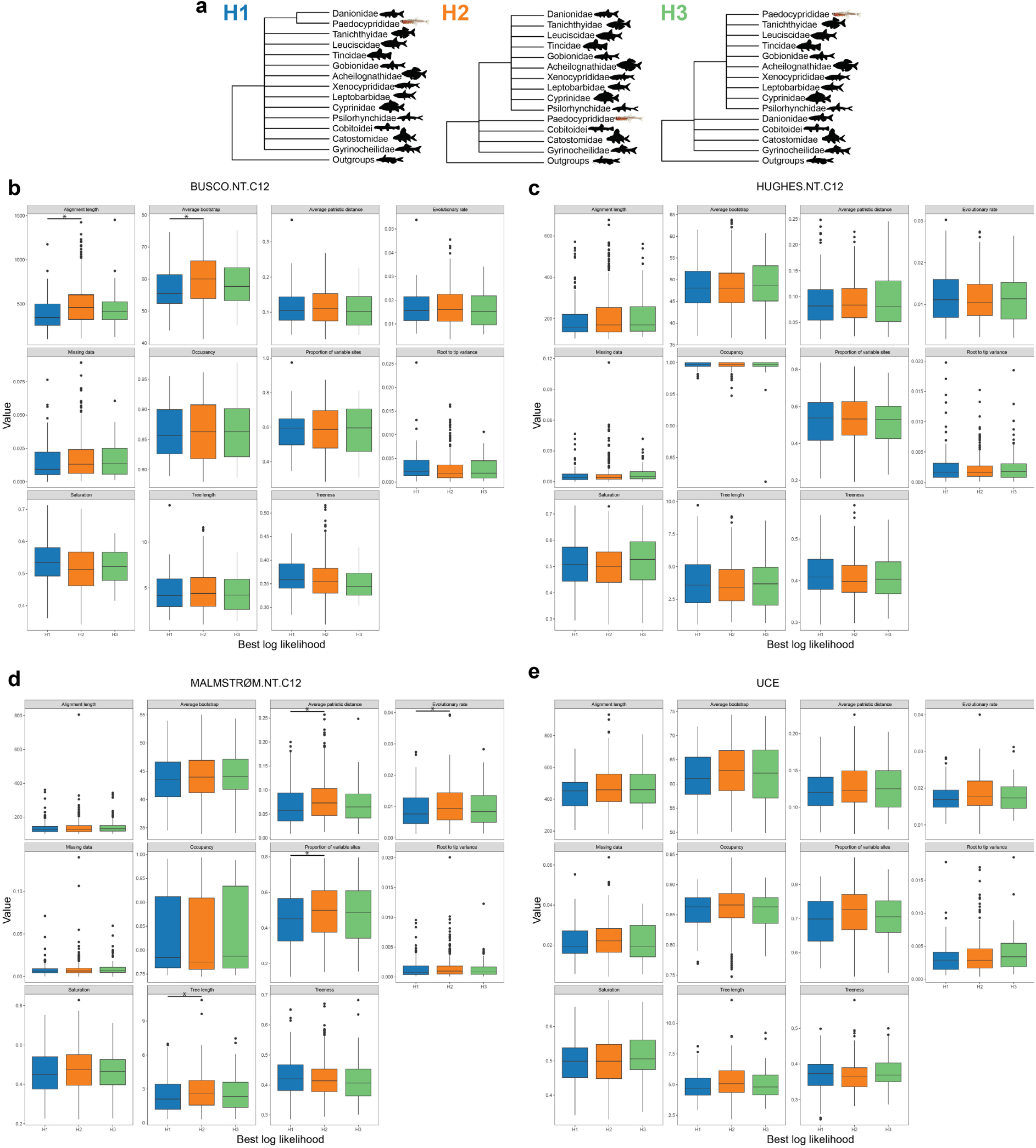
**(a)** Topologies of alternative hypotheses H1, H2, and H3 tested for the phylogenetic position of Paedocyprididae. Panels **(b–e)** compare loci properties estimated by genesortR for unconstrained **(b)** BUSCO.NT.C12, **(c)** HUGHES.NT.C12, **(d)** MALMSTRØM.NT.C12, and **(e)** UCE datasets. Each locus is assigned to H1, H2, or H3 based on the highest-ranked hypothesis according to log-likelihood scores for the constrained topologies. Asterisks indicate significant differences (p < 0.05) after pairwise t-tests, with false discovery rate (FDR) controlled using the Benjamini–Hochberg procedure.

**Supplementary Fig. 25.**
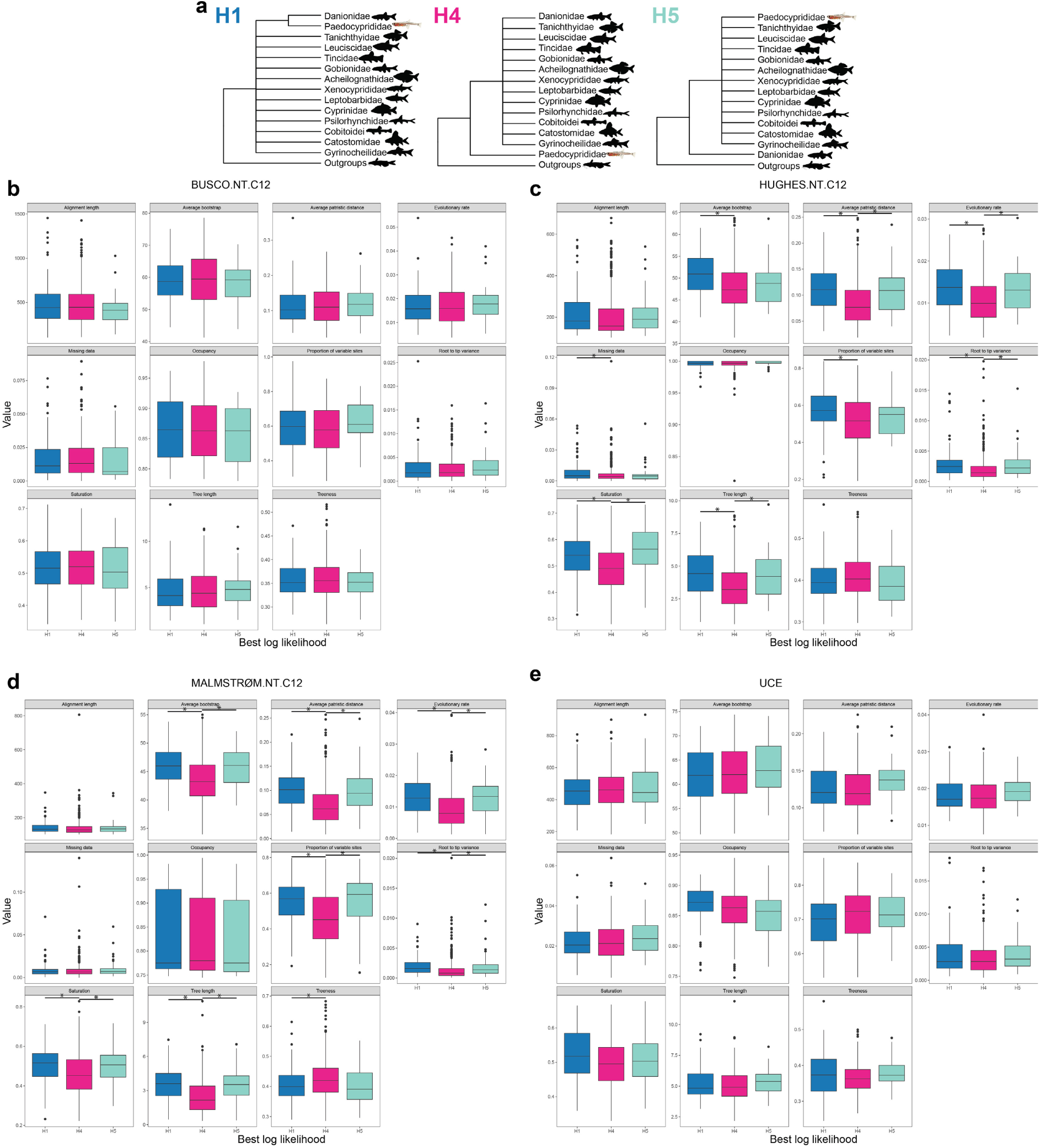
**(a)** Topologies of alternative hypotheses H1, H4, and H5 tested for the phylogenetic position of Paedocyprididae. Panels **(b–e)** compare loci properties estimated by genesortR for unconstrained **(b)** BUSCO.NT.C12, **(c)** HUGHES.NT.C12, **(d)** MALMSTRØM.NT.C12, and **(e)** UCE datasets. Each locus is assigned to H1, H4, or H5 based on the highest-ranked hypothesis according to log-likelihood scores for the constrained topologies. Asterisks indicate significant differences (p < 0.05) after pairwise t-tests, with false discovery rate (FDR) controlled using the Benjamini–Hochberg procedure.

**Supplementary Fig. 26.**
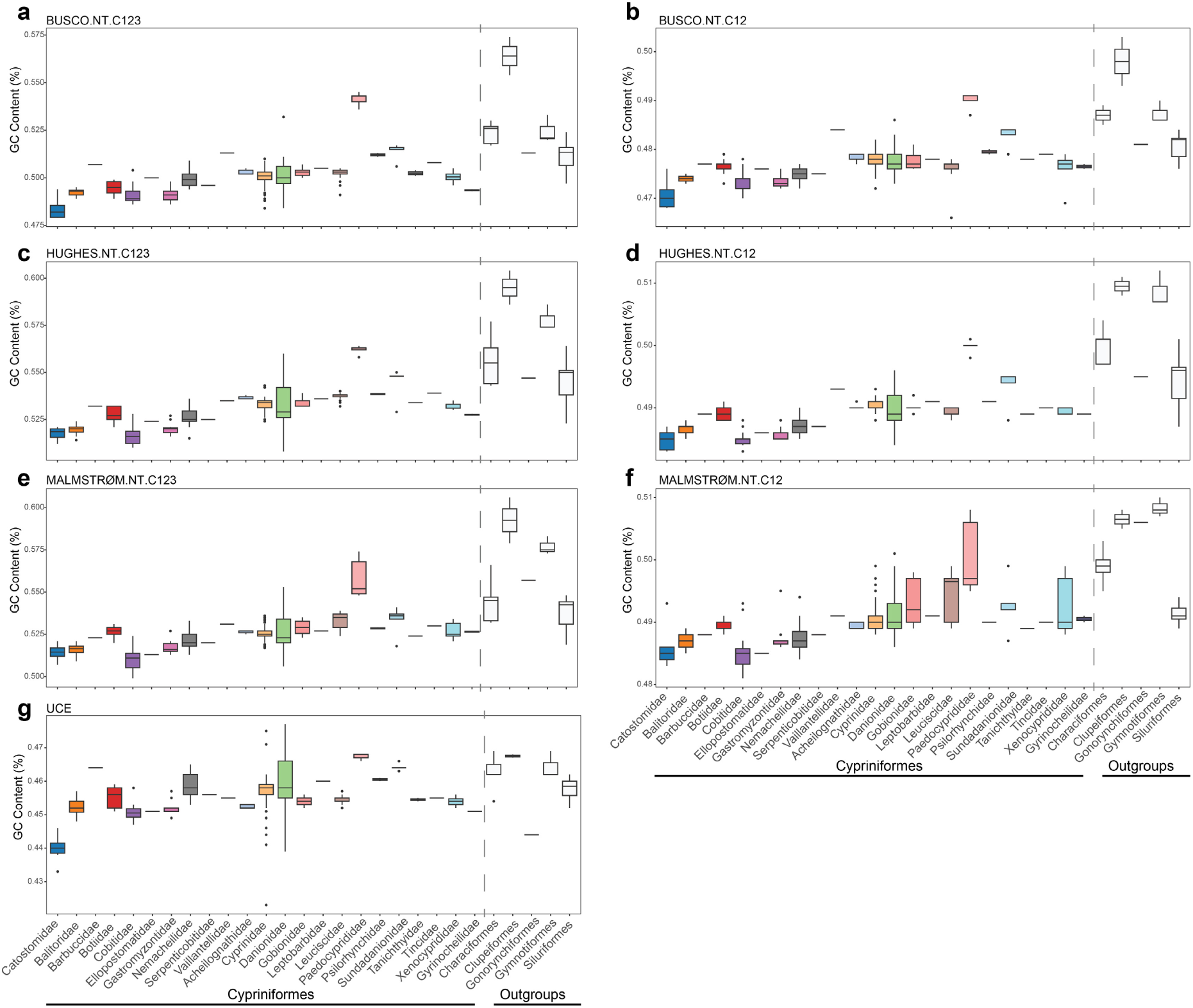
GC content distribution in the nucleotide phylogenomic datasets. **(a)** BUSCO.NT.C123, **(b)** BUSCO.NT.C12, **(c)** HUGHES.NT.C123, **(d)** HUGHES.NT.C12, **(e)** MALMSTRØM.NT.C123, **(f)** MALMSTRØM.NT.C12, and **(g)** UCE. Note the elevated GC% in Paedocyprididae relative to other Cypriniformes families.

**Supplementary Fig. 27.**
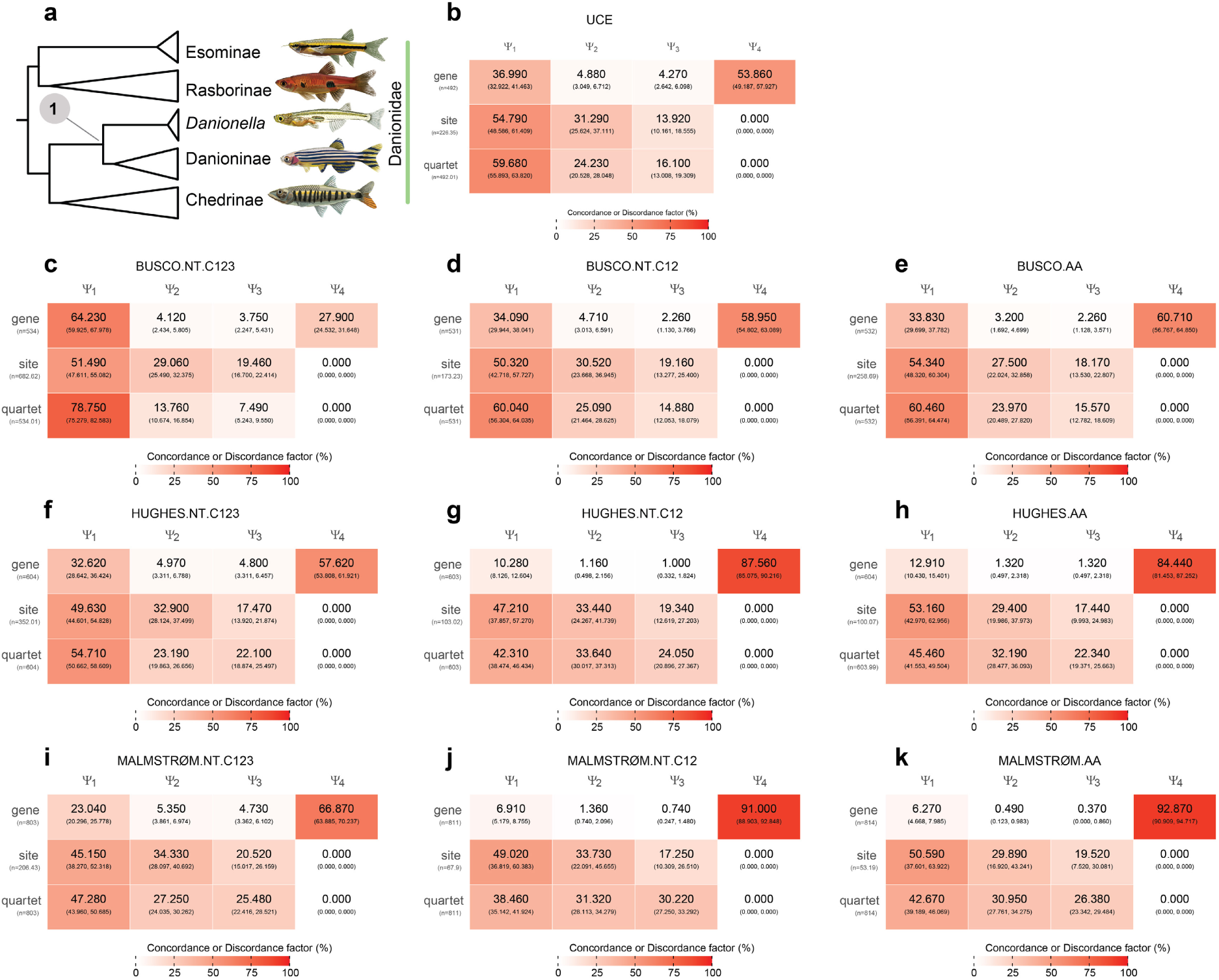
**(a)** Phylogenetic position of *Danionella* recovered from different analyses. Panels **(b–k)** show the gene, site, and quartet concordance vectors for ASTRAL-IV species trees inferred from different datasets: **(b)** UCE, **(c)** BUSCO.NT.C123, **(d)** BUSCO.NT.C12, **(e)** BUSCO.AA, **(f)** HUGHES.NT.C123, **(g)** HUGHES.NT.C12, **(h)** HUGHES.AA, **(i)** MALMSTRØM.NT.C123, **(j)** MALMSTRØM.NT.C12, and **(k)** MALMSTRØM.AA. Values in each cell are percentages, and the numbers in parentheses represent bootstrap-based 95% confidence intervals. Increasing red intensity indicate greater concordance or discordance. ψ₁ represent the concordance factor supporting the focal branch while ψ₂-ψ₄ represent the discordance factors.

**Supplementary Table 1.** Genomes newly generated in this study and their corresponding accession numbers.

**Supplementary Table 2.** NCBI genomes downloaded in this study.

**Supplementary Table 3**. Phylogenomic dataset of 335 species included in this study.

**Supplementary Table 4.** Summary of datasets used in the phylogenomic analyses, showing the number of taxa, retained loci, and alignment lengths.

**Supplementary Table 5.** Different GHOST models tested for the combined first and second codon positions of BUSCO.NT.C12 and UCE nucleotide dataset with 140 species.

**Supplementary Table 6.** Relationship between log-transformed branch lengths and gene concordance factors (gCF) and site concordance factors (sCF).

