## Supplementary Tables for "Phylogenomics of Cypriniformes, the most diverse order of freshwater fishes: consensus, challenges and limitations"

### **Supplementary Table 1.** Genomes newly generated in this study and their corresponding accession numbers.

| Voucher_reads | Voucher_original | Suborder | Family | Genus | Species | SeqFacility | Sequencing_platform | QC_filtering | Phylogenomics_dataset | Phylogenomics_dataset_Remarks |
|---|---|---|---|---|---|---|---|---|---|---|
| LR15244 | TCWC 15787.02 | Catostomoidei | Catostomidae | Catostomus | commersoni | Next Generation Sequencing (NGS) Platform, University of Bern, Switzerland | Illumina NovaSeq 6000 | PASS | Included | |
| LR15247 | TCWC 19725.02 | Catostomoidei | Catostomidae | Cycleptus | sp | Next Generation Sequencing (NGS) Platform, University of Bern, Switzerland | Illumina NovaSeq 6000 | PASS | Included | |
| LR15245 | TCWC 20294.01 | Catostomoidei | Catostomidae | Erimyzon | sucetta | Next Generation Sequencing (NGS) Platform, University of Bern, Switzerland | Illumina NovaSeq 6000 | PASS | Included | |
| LR15246 | TCWC 17181.02 | Catostomoidei | Catostomidae | Ictiobus | bubalus | Next Generation Sequencing (NGS) Platform, University of Bern, Switzerland | Illumina NovaSeq 6000 | PASS | Included | |
| LR15248 | TCWC 20297.11 | Catostomoidei | Catostomidae | Moxostoma | congestum | Next Generation Sequencing (NGS) Platform, University of Bern, Switzerland | Illumina NovaSeq 6000 | PASS | Included | |
| LR15059 | LR15059 | Catostomoidei | Catostomidae | Myxocyprinus | asiaticus | Next Generation Sequencing (NGS) Platform, University of Bern, Switzerland | Illumina NovaSeq 6000 | PASS | Not included | Replaced with GCF_019703515.2 Myxocyprinus asiaticus |
| LR15354 | LR15354 | Cobitoidei | Balitoridae | Balitora | sp | Next Generation Sequencing (NGS) Platform, University of Bern, Switzerland | Illumina NovaSeq 6000 | PASS | Included | |
| LR15094 | LR15094 | Cobitoidei | Balitoridae | Balitoropsis | zollingeri | Next Generation Sequencing (NGS) Platform, University of Bern, Switzerland | Illumina NovaSeq 6000 | PASS | Included | |
| KUFOS-AN-2019.1.1 | KUFOS-AN-2019.1.1 | Cobitoidei | Balitoridae | Bhavania | australis | Agrigenome Labs Pvt. Ltd., India | Illumina NovaSeq 6000 | PASS | Included | |
| LR15176 | LR15176 | Cobitoidei | Balitoridae | Hemimyzon | formosanum | Next Generation Sequencing (NGS) Platform, University of Bern, Switzerland | Illumina NovaSeq 6000 | PASS | Included | |
| LR15139 | LR15139 | Cobitoidei | Balitoridae | Homaloptera | sp | Next Generation Sequencing (NGS) Platform, University of Bern, Switzerland | Illumina NovaSeq 6000 | PASS | Included | |
| LR15140 | LR15140 | Cobitoidei | Balitoridae | Homalopteroides | stephensoni | Next Generation Sequencing (NGS) Platform, University of Bern, Switzerland | Illumina NovaSeq 6000 | PASS | Included | |
| LR15093 | LR15093 | Cobitoidei | Balitoridae | Homalopteroides | tweediei | Next Generation Sequencing (NGS) Platform, University of Bern, Switzerland | Illumina NovaSeq 6000 | PASS | Included | |
| LR15091 | LR15091 | Cobitoidei | Balitoridae | Homalopterula | gymnogaster | Next Generation Sequencing (NGS) Platform, University of Bern, Switzerland | Illumina NovaSeq 6000 | PASS | Included | |
| LR15172 | LR15172 | Cobitoidei | Balitoridae | Lepturichthys | fimbriatus | Next Generation Sequencing (NGS) Platform, University of Bern, Switzerland | Illumina NovaSeq 6000 | PASS | Included | |
| LR15487 | LR15487 | Cobitoidei | Balitoridae | Neohomaloptera | johorensis | Next Generation Sequencing (NGS) Platform, University of Bern, Switzerland | Illumina NovaSeq 6000 | PASS | Included | |
| LR15088 | LR15088 | Cobitoidei | Balitoridae | Sinogastromyzon | puliensis | Next Generation Sequencing (NGS) Platform, University of Bern, Switzerland | Illumina NovaSeq 6000 | PASS | Included | |
| LR15181 | LR15181 | Cobitoidei | Balitoridae | Travancoria | elongata | Next Generation Sequencing (NGS) Platform, University of Bern, Switzerland | Illumina NovaSeq 6000 | PASS | Included | |
| LR15360 | LR15360 | Cobitoidei | Barbuccidae | Barbucca | elongatus | Next Generation Sequencing (NGS) Platform, University of Bern, Switzerland | Illumina NovaSeq 6000 | PASS | Included | |
| LR15330 | LR15330 | Cobitoidei | Botiidae | Ambastaia | sidthimunki | Next Generation Sequencing (NGS) Platform, University of Bern, Switzerland | Illumina NovaSeq 6000 | PASS | Included | |
| LR15149 | LR15149 | Cobitoidei | Botiidae | Botia | kubotai | Next Generation Sequencing (NGS) Platform, University of Bern, Switzerland | Illumina NovaSeq 6000 | PASS | Included | |
| LR15148 | LR15148 | Cobitoidei | Botiidae | Chromobotia | macracanthus | Next Generation Sequencing (NGS) Platform, University of Bern, Switzerland | Illumina NovaSeq 6000 | PASS | Included | |
| LR15074 | LR15074 | Cobitoidei | Botiidae | Leptobotia | elongata | Next Generation Sequencing (NGS) Platform, University of Bern, Switzerland | Illumina NovaSeq 6000 | PASS | Included | |
| 112LR15712 | LR15712 | Cobitoidei | Botiidae | Parabotia | bimaculatus | Next Generation Sequencing (NGS) Platform, University of Bern, Switzerland | Illumina NovaSeq 6000 | PASS | Included | |
| LR15151 | LR15151 | Cobitoidei | Botiidae | Sinibotia | robusta | Next Generation Sequencing (NGS) Platform, University of Bern, Switzerland | Illumina NovaSeq 6000 | PASS | Included | |
| LR15446 | LR15446 | Cobitoidei | Botiidae | Syncrossus | berdmorei | Next Generation Sequencing (NGS) Platform, University of Bern, Switzerland | Illumina NovaSeq 6000 | PASS | Included | |
| 115LR15715 | LR15715 | Cobitoidei | Cobitidae | Acanthopsoides | robertsi | Next Generation Sequencing (NGS) Platform, University of Bern, Switzerland | Illumina NovaSeq 6000 | PASS | Included | |
| LR15340 | LR15340 | Cobitoidei | Cobitidae | Acantopsis | sp | Next Generation Sequencing (NGS) Platform, University of Bern, Switzerland | Illumina NovaSeq 6000 | PASS | Included | |
| LR15193 | LR15193 | Cobitoidei | Cobitidae | Bibarba | sp | Next Generation Sequencing (NGS) Platform, University of Bern, Switzerland | Illumina NovaSeq 6000 | PASS | Included | |
| LR15575 | LR15575 | Cobitoidei | Cobitidae | Canthophrys | gongota | Next Generation Sequencing (NGS) Platform, University of Bern, Switzerland | Illumina NovaSeq 6000 | PASS | Included | |
| LR15465 | LR15465 | Cobitoidei | Cobitidae | Cobitis | sp | Next Generation Sequencing (NGS) Platform, University of Bern, Switzerland | Illumina NovaSeq 6000 | PASS | Included | |
| LR15170 | LR15170 | Cobitoidei | Cobitidae | Kottelatlimia | katik | Next Generation Sequencing (NGS) Platform, University of Bern, Switzerland | Illumina NovaSeq 6000 | PASS | Included | |
| 114LR15714 | LR15714 | Cobitoidei | Cobitidae | Kottelatlimia | pristes | Next Generation Sequencing (NGS) Platform, University of Bern, Switzerland | Illumina NovaSeq 6000 | PASS | Included | |
| LR15467 | LR15467 | Cobitoidei | Cobitidae | Lepidocephalichthys | sp | Next Generation Sequencing (NGS) Platform, University of Bern, Switzerland | Illumina NovaSeq 6000 | PASS | Included | |
| LR15162 | LR15162 | Cobitoidei | Cobitidae | Misgurnus | sp | Next Generation Sequencing (NGS) Platform, University of Bern, Switzerland | Illumina NovaSeq 6000 | PASS | Not included | Replaced with GCF_027580225.1 Misgurnus anguillicaudatus (95.11) and GCA_025389145.1 Misgurnus mizolepis |
| BNHS-FWF-981 | BNHS-FWF-981 | Cobitoidei | Cobitidae | Pangio | bhujia | Agrigenome Labs Pvt. Ltd., India | Illumina NovaSeq 6000 | PASS | Included | |
| LR15141 | LR15141 | Cobitoidei | Cobitidae | Pangio | sp | Next Generation Sequencing (NGS) Platform, University of Bern, Switzerland | Illumina NovaSeq 6000 | PASS | Included | |
| LR15609 | LR15609 | Cobitoidei | Cobitidae | Sabanejewia | balcanica | Next Generation Sequencing (NGS) Platform, University of Bern, Switzerland | Illumina NovaSeq 6000 | PASS | Included | |
| LR15146 | LR15146 | Cobitoidei | Cobitidae | Theriodes | sandakanensis | Next Generation Sequencing (NGS) Platform, University of Bern, Switzerland | Illumina NovaSeq 6000 | PASS | Included | |
| 141LR15748 | LR15748 | Cobitoidei | Ellopostomatidae | Ellopostoma | mystax | Next Generation Sequencing (NGS) Platform, University of Bern, Switzerland | Illumina NovaSeq 6000 | PASS | Included | |
| LR15180 | LR15180 | Cobitoidei | Gastromyzontidae | Annamia | sp | Next Generation Sequencing (NGS) Platform, University of Bern, Switzerland | Illumina NovaSeq 6000 | PASS | Included | |
| LR15085 | LR15085 | Cobitoidei | Gastromyzontidae | Beaufortia | kweichowensis | Next Generation Sequencing (NGS) Platform, University of Bern, Switzerland | Illumina NovaSeq 6000 | PASS | Not included | Replaced with GCA_019155185.1 Beaufortia kweichowensis |
| LR15491 | LR15491 | Cobitoidei | Gastromyzontidae | Erromyzon | sp | Next Generation Sequencing (NGS) Platform, University of Bern, Switzerland | Illumina NovaSeq 6000 | PASS | Included | |
| LR15090 | LR15090 | Cobitoidei | Gastromyzontidae | Glaniopsis | muliradiatus | Next Generation Sequencing (NGS) Platform, University of Bern, Switzerland | Illumina NovaSeq 6000 | PASS | Included | |
| LR15133 | LR15133 | Cobitoidei | Gastromyzontidae | Katibasia | insidiosa | Next Generation Sequencing (NGS) Platform, University of Bern, Switzerland | Illumina NovaSeq 6000 | PASS | Included | |
| LR15175 | LR15175 | Cobitoidei | Gastromyzontidae | Liniparhomaloptera | disparis | Next Generation Sequencing (NGS) Platform, University of Bern, Switzerland | Illumina NovaSeq 6000 | PASS | Included | |
| LR15476 | LR15476 | Cobitoidei | Gastromyzontidae | Plesiomyzon | baotingensis | Next Generation Sequencing (NGS) Platform, University of Bern, Switzerland | Illumina NovaSeq 6000 | PASS | Included | |
| LR15425 | LR15425 | Cobitoidei | Gastromyzontidae | Protomyzon | borneensis | Next Generation Sequencing (NGS) Platform, University of Bern, Switzerland | Illumina NovaSeq 6000 | PASS | Included | |
| LR15067 | LR15067 | Cobitoidei | Gastromyzontidae | Pseudogastromyzon | sp | Next Generation Sequencing (NGS) Platform, University of Bern, Switzerland | Illumina NovaSeq 6000 | PASS | Included | |
| LR15561 | LR15561 | Cobitoidei | Gastromyzontidae | Sewellia | elongata | Next Generation Sequencing (NGS) Platform, University of Bern, Switzerland | Illumina NovaSeq 6000 | PASS | Included | |
| LR15188 | LR15188 | Cobitoidei | Gastromyzontidae | Vanmanenia | sp | Next Generation Sequencing (NGS) Platform, University of Bern, Switzerland | Illumina NovaSeq 6000 | PASS | Not included | Same lineage as LR15175 |
| 113LR15713 | LR15713 | Cobitoidei | Gastromyzontidae | Yaoshania | pachychilus | Next Generation Sequencing (NGS) Platform, University of Bern, Switzerland | Illumina NovaSeq 6000 | PASS | Included | |
| LR15343 | LR15343 | Cobitoidei | Nemacheilidae | Aborichthys | elongatus | Next Generation Sequencing (NGS) Platform, University of Bern, Switzerland | Illumina NovaSeq 6000 | PASS | Included | |
| LR15190 | LR15190 | Cobitoidei | Nemacheilidae | Barbatula | sp | Next Generation Sequencing (NGS) Platform, University of Bern, Switzerland | Illumina NovaSeq 6000 | PASS | Included | |
| LR15073 | LR15073 | Cobitoidei | Nemacheilidae | Homatula | variegata | Next Generation Sequencing (NGS) Platform, University of Bern, Switzerland | Illumina NovaSeq 6000 | PASS | Included | |
| KUFOS-AN-2019.2.1 | KUFOS-AN-2019.2.1 | Cobitoidei | Nemacheilidae | Indoreonectes | keralensis | Agrigenome Labs Pvt. Ltd., India | Illumina NovaSeq 6000 | PASS | Included | |
| LR15462 | LR15462 | Cobitoidei | Nemacheilidae | Lefua | sp | Next Generation Sequencing (NGS) Platform, University of Bern, Switzerland | Illumina NovaSeq 6000 | PASS | Included | |
| KUFOS-AN-2021.1.1 | KUFOS-AN-2021.1.1 | Cobitoidei | Nemacheilidae | Mesonoemacheilus | triangularis | Agrigenome Labs Pvt. Ltd., India | Illumina NovaSeq 6000 | PASS | Included | |
| LR15337 | LR15337 | Cobitoidei | Nemacheilidae | Micronemacheilus | sp | Next Generation Sequencing (NGS) Platform, University of Bern, Switzerland | Illumina NovaSeq 6000 | PASS | Included | |
| LR15459 | LR15459 | Cobitoidei | Nemacheilidae | Nemacheilus | selangoricus | Next Generation Sequencing (NGS) Platform, University of Bern, Switzerland | Illumina NovaSeq 6000 | PASS | Included | |
| LR15611 | LR15611 | Cobitoidei | Nemacheilidae | Neonoemacheilus | sp | Next Generation Sequencing (NGS) Platform, University of Bern, Switzerland | Illumina NovaSeq 6000 | PASS | Included | |
| LR15490 | LR15490 | Cobitoidei | Nemacheilidae | Oreonectes | platycephalus | Next Generation Sequencing (NGS) Platform, University of Bern, Switzerland | Illumina NovaSeq 6000 | PASS | Included | |
| LR15504 | LR15504 | Cobitoidei | Nemacheilidae | Paracanthocobitis | botia | Next Generation Sequencing (NGS) Platform, University of Bern, Switzerland | Illumina NovaSeq 6000 | PASS | Included | |
| LR15189 | LR15189 | Cobitoidei | Nemacheilidae | Petruichthys | brevis | Next Generation Sequencing (NGS) Platform, University of Bern, Switzerland | Illumina NovaSeq 6000 | PASS | Included | |
| LR15152 | LR15152 | Cobitoidei | Nemacheilidae | Schistura | fasciolata | Next Generation Sequencing (NGS) Platform, University of Bern, Switzerland | Illumina NovaSeq 6000 | PASS | Included | |
| LR15454 | LR15454 | Cobitoidei | Nemacheilidae | Schistura | sp1 | Next Generation Sequencing (NGS) Platform, University of Bern, Switzerland | Illumina NovaSeq 6000 | PASS | Included | |
| LR15496 | LR15496 | Cobitoidei | Nemacheilidae | Schistura | sp2 | Next Generation Sequencing (NGS) Platform, University of Bern, Switzerland | Illumina NovaSeq 6000 | PASS | Included | |
| LR15455 | LR15455 | Cobitoidei | Nemacheilidae | Sundoreonectes | sabanus | Next Generation Sequencing (NGS) Platform, University of Bern, Switzerland | Illumina NovaSeq 6000 | PASS | Included | |
| LR15072 | LR15072 | Cobitoidei | Nemacheilidae | Traccatichthys | pulcher | Next Generation Sequencing (NGS) Platform, University of Bern, Switzerland | Illumina NovaSeq 6000 | PASS | Included | |
| LR15357 | LR15357 | Cobitoidei | Serpenticobitidae | Serpenticobitis | zonata | Next Generation Sequencing (NGS) Platform, University of Bern, Switzerland | Illumina NovaSeq 6000 | PASS | Included | |
| LR15334 | LR15334 | Cobitoidei | Vaillantellidae | Vaillantella | maassii | Next Generation Sequencing (NGS) Platform, University of Bern, Switzerland | Illumina NovaSeq 6000 | PASS | Included | |
| LR15562 | LR15562 | Cyprinoidei | Acheilognathidae | Acheilognathus | deignani | Next Generation Sequencing (NGS) Platform, University of Bern, Switzerland | Illumina NovaSeq 6000 | PASS | Included | |
| LR15063 | LR15063 | Cyprinoidei | Acheilognathidae | Acheilognathus | sp | Next Generation Sequencing (NGS) Platform, University of Bern, Switzerland | Illumina NovaSeq 6000 | PASS | Included | |
| LR15636 | LR15636 | Cyprinoidei | Acheilognathidae | Paratanakia | himantegus | Next Generation Sequencing (NGS) Platform, University of Bern, Switzerland | Illumina NovaSeq 6000 | PASS | Included | |
| LR15661 | LR15661 | Cyprinoidei | Acheilognathidae | Rhodeus | amarus | Next Generation Sequencing (NGS) Platform, University of Bern, Switzerland | Illumina NovaSeq 6000 | PASS | Included | |
| LR15266 | LR15266 | Cyprinoidei | Cyprinidae | Acapoeta | tanganicae | Next Generation Sequencing (NGS) Platform, University of Bern, Switzerland | Illumina NovaSeq 6000 | PASS | Included | |
| 123LR15726 | LR15726 | Cyprinoidei | Cyprinidae | Acrossocheilus | sp | Next Generation Sequencing (NGS) Platform, University of Bern, Switzerland | Illumina NovaSeq 6000 | PASS | Not included | Replaced with GCA_039880705.1 Acrossocheilus fasciatus |
| LR15615 | LR15615 | Cyprinoidei | Cyprinidae | Ageneiogarra | sp | Next Generation Sequencing (NGS) Platform, University of Bern, Switzerland | Illumina NovaSeq 6000 | PASS | Included | |
| LR15639 | LR15639 | Cyprinoidei | Cyprinidae | Balantiocheilos | melanopterus | Next Generation Sequencing (NGS) Platform, University of Bern, Switzerland | Illumina NovaSeq 6000 | PASS | Included | |
| LR15079 | LR15079 | Cyprinoidei | Cyprinidae | Barbichthys | laevis | Next Generation Sequencing (NGS) Platform, University of Bern, Switzerland | Illumina NovaSeq 6000 | PASS | Included | |
| LR15102 | LR15102 | Cyprinoidei | Cyprinidae | Barbodes | lateristriga | Next Generation Sequencing (NGS) Platform, University of Bern, Switzerland | Illumina NovaSeq 6000 | PASS | Included | |
| 120LR15721 | LR15721 | Cyprinoidei | Cyprinidae | Barbodes | sellifer | Next Generation Sequencing (NGS) Platform, University of Bern, Switzerland | Illumina NovaSeq 6000 | PASS | Included | |
| LR15256 | LR15256 | Cyprinoidei | Cyprinidae | Barboides | gracilis | Next Generation Sequencing (NGS) Platform, University of Bern, Switzerland | Illumina NovaSeq 6000 | PASS | Included | |
| EA0124 | EA0124 | Cyprinoidei | Cyprinidae | Barbonymus | altus | Next Generation Sequencing (NGS) Platform, University of Bern, Switzerland | Illumina NovaSeq 6000 | PASS | Included | |
| LR15111 | LR15111 | Cyprinoidei | Cyprinidae | Ceratogarra | cambodgiensis | Next Generation Sequencing (NGS) Platform, University of Bern, Switzerland | Illumina NovaSeq 6000 | PASS | Included | |
| EA0357 | EA0357 | Cyprinoidei | Cyprinidae | Chagunius | sp | Next Generation Sequencing (NGS) Platform, University of Bern, Switzerland | Illumina NovaSeq 6000 | PASS | Not included | Same lineage as EA0401 |
| EA0401 | EA0401 | Cyprinoidei | Cyprinidae | Chagunius | sp | Next Generation Sequencing (NGS) Platform, University of Bern, Switzerland | Illumina NovaSeq 6000 | PASS | Included | |
| LR15270 | LR15270 | Cyprinoidei | Cyprinidae | Clypeobarbus | sp | Next Generation Sequencing (NGS) Platform, University of Bern, Switzerland | Illumina NovaSeq 6000 | PASS | Included | |
| LR10955 | LR10955 | Cyprinoidei | Cyprinidae | Crossocheilus | reticulatus | Next Generation Sequencing (NGS) Platform, University of Bern, Switzerland | Illumina NovaSeq 6000 | PASS | Included | |
| LR10341 | LR10341 | Cyprinoidei | Cyprinidae | Cyclocheilichthys | apogon | Next Generation Sequencing (NGS) Platform, University of Bern, Switzerland | Illumina NovaSeq 6000 | PASS | Included | |
| KUFOS-AN-2021.4.1 | KUFOS-AN-2021.4.1 | Cyprinoidei | Cyprinidae | Dawkinsia | filamentosa | Agrigenome Labs Pvt. Ltd., India | Illumina NovaSeq 6000 | PASS | Included | |
| 140LR15737 | LR15737 | Cyprinoidei | Cyprinidae | Desmopuntius | sp | Next Generation Sequencing (NGS) Platform, University of Bern, Switzerland | Illumina NovaSeq 6000 | PASS | Included | |
| 122LR15725 | LR15725 | Cyprinoidei | Cyprinidae | Discherodontus | ashmeadi | Next Generation Sequencing (NGS) Platform, University of Bern, Switzerland | Illumina NovaSeq 6000 | PASS | Included | |
| KUFOS-AN-2019.3.1 | KUFOS-AN-2019.3.1 | Cyprinoidei | Cyprinidae | Eechathalakenda | ophicephalus | Agrigenome Labs Pvt. Ltd., India | Illumina NovaSeq 6000 | PASS | Included | |
| 119LR15720 | LR15720 | Cyprinoidei | Cyprinidae | Eirmotus | sp1 | Next Generation Sequencing (NGS) Platform, University of Bern, Switzerland | Illumina NovaSeq 6000 | PASS | Included | |
| LR09582 | LR09582 | Cyprinoidei | Cyprinidae | Eirmotus | sp2 | Next Generation Sequencing (NGS) Platform, University of Bern, Switzerland | Illumina NovaSeq 6000 | PASS | Included | |
| LR15276 | LR15276 | Cyprinoidei | Cyprinidae | Enteromius | sp | Next Generation Sequencing (NGS) Platform, University of Bern, Switzerland | Illumina NovaSeq 6000 | PASS | Not included | Replaced with GCA_032359435.1 Enteromius unitaeniatus |
| LR15296 | LR15296 | Cyprinoidei | Cyprinidae | Enteromius | sp | Next Generation Sequencing (NGS) Platform, University of Bern, Switzerland | Illumina NovaSeq 6000 | FAIL: high read error and poor model fit in kmer profiling; genome size estimates has a % diff > 10% between k=21 and k=31 models; unrealistic high level of heterozygozity; very low BUSCO single-copy recovery for both abyss and spades assemblies | Not included | |
| LR15606 | LR15606 | Cyprinoidei | Cyprinidae | Enteromius | sp1 | Next Generation Sequencing (NGS) Platform, University of Bern, Switzerland | Illumina NovaSeq 6000 | PASS | Included | |
| LR01655 | LR01655 | Cyprinoidei | Cyprinidae | Epalzeorhynchos | kalopterum | Next Generation Sequencing (NGS) Platform, University of Bern, Switzerland | Illumina NovaSeq 6000 | PASS | Included | |
| LR15563 | LR15563 | Cyprinoidei | Cyprinidae | Folifer | brevifilis | Next Generation Sequencing (NGS) Platform, University of Bern, Switzerland | Illumina NovaSeq 6000 | PASS | Included | |
| KUFOS-RR-2022.P.4 | KUFOS-RR-2022.P.4 | Cyprinoidei | Cyprinidae | Garra | periyarensis | Agrigenome Labs Pvt. Ltd., India | Illumina NovaSeq 6000 | PASS | Included | |
| LR15635 | LR15635 | Cyprinoidei | Cyprinidae | Haludaria | fasciata | Next Generation Sequencing (NGS) Platform, University of Bern, Switzerland | Illumina NovaSeq 6000 | PASS | Included | |
| LR15078 | LR15078 | Cyprinoidei | Cyprinidae | Hampala | ampalong | Next Generation Sequencing (NGS) Platform, University of Bern, Switzerland | Illumina NovaSeq 6000 | PASS | Included | |
| KUFOS-RR-2022.P.3 | KUFOS-RR-2022.P.3 | Cyprinoidei | Cyprinidae | Hypselobarbus | periyarensis | Agrigenome Labs Pvt. Ltd., India | Illumina NovaSeq 6000 | PASS | Included | |
| LR15278 | LR15278 | Cyprinoidei | Cyprinidae | Labeo | cylindricus | Next Generation Sequencing (NGS) Platform, University of Bern, Switzerland | Illumina NovaSeq 6000 | PASS | Included | |
| LR15282 | LR15282 | Cyprinoidei | Cyprinidae | Labeobarbus | sp | Next Generation Sequencing (NGS) Platform, University of Bern, Switzerland | Illumina NovaSeq 6000 | PASS | Not included | Replaced with GCA_034780575.1 Labeobarbus marequensis |
| LR12884 | LR12884 | Cyprinoidei | Cyprinidae | Labiobarbus | ocellatus | Next Generation Sequencing (NGS) Platform, University of Bern, Switzerland | Illumina NovaSeq 6000 | PASS | Included | |
| KUFOS-RR-2022.P.2 | KUFOS-RR-2022.P.2 | Cyprinoidei | Cyprinidae | Lepidopygopsis | typus | Agrigenome Labs Pvt. Ltd., India | Illumina NovaSeq 6000 | PASS | Included | |
| LR15104 | LR15104 | Cyprinoidei | Cyprinidae | Lobocheilos | aureolineatus | Next Generation Sequencing (NGS) Platform, University of Bern, Switzerland | Illumina NovaSeq 6000 | PASS | Included | |
| LR15565 | LR15565 | Cyprinoidei | Cyprinidae | Mystacoleucus | argenteus | Next Generation Sequencing (NGS) Platform, University of Bern, Switzerland | Illumina NovaSeq 6000 | PASS | Included | |
| LR15612 | LR15612 | Cyprinoidei | Cyprinidae | Neolissochilus | sp | Next Generation Sequencing (NGS) Platform, University of Bern, Switzerland | Illumina NovaSeq 6000 | PASS | Included | |
| KUFOS-AN-2021.7.1 | KUFOS-AN-2021.7.1 | Cyprinoidei | Cyprinidae | Neolissochilus | wynaadensis | Agrigenome Labs Pvt. Ltd., India | Illumina NovaSeq 6000 | PASS | Included | |
| 143LR15682 | LR15682 | Cyprinoidei | Cyprinidae | Oliotius | oligolepis | Next Generation Sequencing (NGS) Platform, University of Bern, Switzerland | Illumina NovaSeq 6000 | PASS | Included | |
| EA1291 | EA1291 | Cyprinoidei | Cyprinidae | Oreichthys | sp | Next Generation Sequencing (NGS) Platform, University of Bern, Switzerland | Illumina NovaSeq 6000 | PASS | Included | |
| LR09367 | LR09367 | Cyprinoidei | Cyprinidae | Osteobrama | cunma | Next Generation Sequencing (NGS) Platform, University of Bern, Switzerland | Illumina NovaSeq 6000 | PASS | Included | |
| LR15096 | LR15096 | Cyprinoidei | Cyprinidae | Osteochilus | vittatoides | Next Generation Sequencing (NGS) Platform, University of Bern, Switzerland | Illumina NovaSeq 6000 | PASS | Included | |
| LR01656 | LR01656 | Cyprinoidei | Cyprinidae | Paracrossochilus | sp | Next Generation Sequencing (NGS) Platform, University of Bern, Switzerland | Illumina NovaSeq 6000 | PASS | Included | |
| LR15618 | LR15618 | Cyprinoidei | Cyprinidae | Paraspinibarbus | sp | Next Generation Sequencing (NGS) Platform, University of Bern, Switzerland | Illumina NovaSeq 6000 | PASS | Included | |
| KUFOS-AN-2021.5.1 | KUFOS-AN-2021.5.1 | Cyprinoidei | Cyprinidae | Pethia | conchonius | Agrigenome Labs Pvt. Ltd., India | Illumina NovaSeq 6000 | PASS | Included | |
| LR06584 | LR06584 | Cyprinoidei | Cyprinidae | Pethia | sp1 | Next Generation Sequencing (NGS) Platform, University of Bern, Switzerland | Illumina NovaSeq 6000 | PASS | Included | |
| LR15646 | LR15646 | Cyprinoidei | Cyprinidae | Pethia | sp2 | Next Generation Sequencing (NGS) Platform, University of Bern, Switzerland | Illumina NovaSeq 6000 | PASS | Included | |
| LR15083 | LR15083 | Cyprinoidei | Cyprinidae | Poropuntius | sp | Next Generation Sequencing (NGS) Platform, University of Bern, Switzerland | Illumina NovaSeq 6000 | PASS | Included | |
| LR15065 | LR15065 | Cyprinoidei | Cyprinidae | Probarbus | jullieni | Next Generation Sequencing (NGS) Platform, University of Bern, Switzerland | Illumina NovaSeq 6000 | PASS | Included | |
| 125LR15728 | LR15728 | Cyprinoidei | Cyprinidae | Puntigrus | partipentazona | Next Generation Sequencing (NGS) Platform, University of Bern, Switzerland | Illumina NovaSeq 6000 | PASS | Included | |
| 146LR15695 | LR15695 | Cyprinoidei | Cyprinidae | Puntigrus | tetrazona | Next Generation Sequencing (NGS) Platform, University of Bern, Switzerland | Illumina NovaSeq 6000 | PASS | Not included | Replaced with GCF_018831695.1 Puntigrus tetrazona |
| LR15108 | LR15108 | Cyprinoidei | Cyprinidae | Puntioplites | bulu | Next Generation Sequencing (NGS) Platform, University of Bern, Switzerland | Illumina NovaSeq 6000 | PASS | Included | |
| EA1300 | EA1300 | Cyprinoidei | Cyprinidae | Puntius | sp | Next Generation Sequencing (NGS) Platform, University of Bern, Switzerland | Illumina NovaSeq 6000 | PASS | Included | |
| LR15257 | LR15257 | Cyprinoidei | Cyprinidae | Rohanella | titteya | Next Generation Sequencing (NGS) Platform, University of Bern, Switzerland | Illumina NovaSeq 6000 | PASS | Included | |
| 124LR15727 | LR15727 | Cyprinoidei | Cyprinidae | Rohtee | ogilbii | Next Generation Sequencing (NGS) Platform, University of Bern, Switzerland | Illumina NovaSeq 6000 | PASS | Included | |

| | | | | | | | | | | |
|---|---|---|---|---|---|---|---|---|---|---|
| 145LR15691 | LR15691 | Cyprinoidei | Cyprinidae | Sahyadria | denisonii | Next Generation Sequencing (NGS) Platform, University of Bern, Switzerland | Illumina NovaSeq 6000 | PASS | Included | |
| LR15061 | LR15061 | Cyprinoidei | Cyprinidae | Sawbwa | resplendens | Next Generation Sequencing (NGS) Platform, University of Bern, Switzerland | Illumina NovaSeq 6000 | PASS | Included | |
| LR15564 | LR15564 | Cyprinoidei | Cyprinidae | Scaphiodonichthys | acanthopterus | Next Generation Sequencing (NGS) Platform, University of Bern, Switzerland | Illumina NovaSeq 6000 | PASS | Included | |
| LR15101 | LR15101 | Cyprinoidei | Cyprinidae | Schismatorhynchos | heterorhynchos | Next Generation Sequencing (NGS) Platform, University of Bern, Switzerland | Illumina NovaSeq 6000 | PASS | Included | |
| LR09392 | LR09392 | Cyprinoidei | Cyprinidae | Semiplotus | sp | Next Generation Sequencing (NGS) Platform, University of Bern, Switzerland | Illumina NovaSeq 6000 | PASS | Included | |
| 111LR15711 | LR15711 | Cyprinoidei | Cyprinidae | Spinibarbichthys | denticulatus | Next Generation Sequencing (NGS) Platform, University of Bern, Switzerland | Illumina NovaSeq 6000 | PASS | Included | |
| LR15617 | LR15617 | Cyprinoidei | Cyprinidae | Spinibarbichthys | sp | Next Generation Sequencing (NGS) Platform, University of Bern, Switzerland | Illumina NovaSeq 6000 | PASS | Included | |
| 126LR15731 | LR15731 | Cyprinoidei | Cyprinidae | Striuntius | lineatus | Next Generation Sequencing (NGS) Platform, University of Bern, Switzerland | Illumina NovaSeq 6000 | PASS | Included | |
| LR06590 | LR06590 | Cyprinoidei | Cyprinidae | Systomus | binduchitra | Next Generation Sequencing (NGS) Platform, University of Bern, Switzerland | Illumina NovaSeq 6000 | PASS | Included | |
| KUFOS-RR-2022.P.1 | KUFOS-RR-2022.P.1 | Cyprinoidei | Cyprinidae | Tariqilabeo | periyarensis | Agrigenome Labs Pvt. Ltd., India | Illumina NovaSeq 6000 | PASS | Included | |
| LR12880 | LR12880 | Cyprinoidei | Cyprinidae | Thynnichthys | sp | Next Generation Sequencing (NGS) Platform, University of Bern, Switzerland | Illumina NovaSeq 6000 | PASS | Included | |
| LR09401 | LR09401 | Cyprinoidei | Cyprinidae | Tor | sp1 | Next Generation Sequencing (NGS) Platform, University of Bern, Switzerland | Illumina NovaSeq 6000 | PASS | Included | |
| LR15100 | LR15100 | Cyprinoidei | Cyprinidae | Tor | sp2 | Next Generation Sequencing (NGS) Platform, University of Bern, Switzerland | Illumina NovaSeq 6000 | PASS | Included | |
| LR01638 | LR01638 | Cyprinoidei | Danionidae | Amblypharyngodon | chulabhornae | Next Generation Sequencing (NGS) Platform, University of Bern, Switzerland | Illumina NovaSeq 6000 | PASS | Included | |
| LR15576 | LR15576 | Cyprinoidei | Danionidae | Barilius | mesopotamicus | Next Generation Sequencing (NGS) Platform, University of Bern, Switzerland | Illumina NovaSeq 6000 | PASS | Included | |
| LR15871 | LR15871 | Cyprinoidei | Danionidae | Boraras | brigittae | Next Generation Sequencing (NGS) Platform, University of Bern, Switzerland | PacBio HiFi Revio | PASS | Included | |
| LR12206 | LR12206 | Cyprinoidei | Danionidae | Boraras | micros | Next Generation Sequencing (NGS) Platform, University of Bern, Switzerland | Illumina NovaSeq 6000 | PASS | Included | |
| LR12190 | LR12190 | Cyprinoidei | Danionidae | Boraras | urophthalmoides | Next Generation Sequencing (NGS) Platform, University of Bern, Switzerland | Illumina NovaSeq 6000 | PASS | Included | |
| LR09154 | LR09154 | Cyprinoidei | Danionidae | Brevibora | cheeya1 | Next Generation Sequencing (NGS) Platform, University of Bern, Switzerland | Illumina NovaSeq 6000 | PASS | Included | |
| LR10130 | LR10130 | Cyprinoidei | Danionidae | Brevibora | cheeya2 | Next Generation Sequencing (NGS) Platform, University of Bern, Switzerland | Illumina NovaSeq 6000 | PASS | Included | |
| LR11616 | LR11616 | Cyprinoidei | Danionidae | Brevibora | dorsiocellata | Next Generation Sequencing (NGS) Platform, University of Bern, Switzerland | Illumina NovaSeq 6000 | PASS | Included | |
| LR15567 | LR15567 | Cyprinoidei | Danionidae | Cabdio | morar | Next Generation Sequencing (NGS) Platform, University of Bern, Switzerland | Illumina NovaSeq 6000 | PASS | Included | |
| 131LR08589 | LR08589 | Cyprinoidei | Danionidae | Chelaethiops | elongatus | Next Generation Sequencing (NGS) Platform, University of Bern, Switzerland | Illumina NovaSeq 6000 | PASS | Included | |
| LR15596 | LR15596 | Cyprinoidei | Danionidae | Chelaethiops | sp1 | Next Generation Sequencing (NGS) Platform, University of Bern, Switzerland | Illumina NovaSeq 6000 | PASS | Included | |
| LR15597 | LR15597 | Cyprinoidei | Danionidae | Chelaethiops | sp2 | Next Generation Sequencing (NGS) Platform, University of Bern, Switzerland | Illumina NovaSeq 6000 | PASS | Included | |
| LR01642 | LR01642 | Cyprinoidei | Danionidae | Danio | erythromicron | Next Generation Sequencing (NGS) Platform, University of Bern, Switzerland | Illumina NovaSeq 6000 | PASS | Included | |
| LR02056 | LR02056 | Cyprinoidei | Danionidae | Danio | margaritatus | Next Generation Sequencing (NGS) Platform, University of Bern, Switzerland | Illumina NovaSeq 6000 | PASS | Included | |
| 144LR15698 | LR15698 | Cyprinoidei | Danionidae | Devario | sondhii | Next Generation Sequencing (NGS) Platform, University of Bern, Switzerland | Illumina NovaSeq 6000 | PASS | Included | |
| LR14717 | LR14717 | Cyprinoidei | Danionidae | Devario | sp | Next Generation Sequencing (NGS) Platform, University of Bern, Switzerland | Illumina NovaSeq 6000 | FAIL: high read error and poor model fit in kmer profiling; genome size estimates has a % diff > 10% between k=21 and k=31 models; unrealistic high level of heterozygozity; very low BUSCO single-copy recovery for both abyss and spades assemblies | Not included | |
| LR15598 | LR15598 | Cyprinoidei | Danionidae | Engraulicypris | sp1 | Next Generation Sequencing (NGS) Platform, University of Bern, Switzerland | Illumina NovaSeq 6000 | PASS | Included | |
| LR15656 | LR15656 | Cyprinoidei | Danionidae | Engraulicypris | sp2 | Next Generation Sequencing (NGS) Platform, University of Bern, Switzerland | Illumina NovaSeq 6000 | PASS | Included | |
| 116LR15716 | LR15716 | Cyprinoidei | Danionidae | Esomus | danricus | Next Generation Sequencing (NGS) Platform, University of Bern, Switzerland | Illumina NovaSeq 6000 | PASS | Included | |
| LR01653 | LR01653 | Cyprinoidei | Danionidae | Esomus | metallicus | Next Generation Sequencing (NGS) Platform, University of Bern, Switzerland | Illumina NovaSeq 6000 | PASS | Included | |
| LR12503 | LR12503 | Cyprinoidei | Danionidae | Esomus | sp | Next Generation Sequencing (NGS) Platform, University of Bern, Switzerland | Illumina NovaSeq 6000 | PASS | Included | |
| LR01646 | LR01646 | Cyprinoidei | Danionidae | Inlecypris | auropurpurea | Next Generation Sequencing (NGS) Platform, University of Bern, Switzerland | Illumina NovaSeq 6000 | PASS | Included | |
| LR01674 | LR01674 | Cyprinoidei | Danionidae | Kottelatia | brittani | Next Generation Sequencing (NGS) Platform, University of Bern, Switzerland | Illumina NovaSeq 6000 | PASS | Included | |
| 132LR09413 | LR09413 | Cyprinoidei | Danionidae | Laubuka | sp1 | Next Generation Sequencing (NGS) Platform, University of Bern, Switzerland | Illumina NovaSeq 6000 | PASS | Included | |
| KUFOS-AN-2021.3.1 | KUFOS-AN-2021.3.1 | Cyprinoidei | Danionidae | Laubuka | sp2 | Agrigenome Labs Pvt. Ltd., India | Illumina NovaSeq 6000 | PASS | Included | |
| LR02238 | LR02238 | Cyprinoidei | Danionidae | Laubuka | sp1 | Next Generation Sequencing (NGS) Platform, University of Bern, Switzerland | Illumina NovaSeq 6000 | PASS | Not included | Same lineage as 132LR09413 |
| LR15601 | LR15601 | Cyprinoidei | Danionidae | Leptocypris | sp | Next Generation Sequencing (NGS) Platform, University of Bern, Switzerland | Illumina NovaSeq 6000 | PASS | Included | |
| LR01659 | LR01659 | Cyprinoidei | Danionidae | Luciosoma | sp 2 | Next Generation Sequencing (NGS) Platform, University of Bern, Switzerland | Illumina NovaSeq 6000 | PASS | Included | |
| LR15098 | LR15098 | Cyprinoidei | Danionidae | Malayochela | maassi | Next Generation Sequencing (NGS) Platform, University of Bern, Switzerland | Illumina NovaSeq 6000 | PASS | Included | |
| LR15378 | LR15378 | Cyprinoidei | Danionidae | Malayochela | maassi | Next Generation Sequencing (NGS) Platform, University of Bern, Switzerland | Illumina NovaSeq 6000 | PASS | Not included | Same lineage as LR15098 |
| LR15109 | LR15109 | Cyprinoidei | Danionidae | Microdevario | kubotai | Next Generation Sequencing (NGS) Platform, University of Bern, Switzerland | Illumina NovaSeq 6000 | PASS | Not included | Same lineage as LR15568 |
| LR15568 | LR15568 | Cyprinoidei | Danionidae | Microdevario | kubotai | Next Generation Sequencing (NGS) Platform, University of Bern, Switzerland | Illumina NovaSeq 6000 | PASS | Included | |
| LR15650 | LR15650 | Cyprinoidei | Danionidae | Microrasbora | rubescens | Next Generation Sequencing (NGS) Platform, University of Bern, Switzerland | Illumina NovaSeq 6000 | PASS | Included | |
| LR15651 | LR15651 | Cyprinoidei | Danionidae | Microrasbora | rubescens | Next Generation Sequencing (NGS) Platform, University of Bern, Switzerland | Illumina NovaSeq 6000 | PASS | Not included | Same lineage as LR15650 |
| 121LR15723 | LR15723 | Cyprinoidei | Danionidae | Nematabramis | everetti | Next Generation Sequencing (NGS) Platform, University of Bern, Switzerland | Illumina NovaSeq 6000 | PASS | Included | |
| LR15099 | LR15099 | Cyprinoidei | Danionidae | Nematabramis | steindachneri | Next Generation Sequencing (NGS) Platform, University of Bern, Switzerland | Illumina NovaSeq 6000 | PASS | Included | |
| LR14062 | LR14062 | Cyprinoidei | Danionidae | Neochela | dadiburjori | Next Generation Sequencing (NGS) Platform, University of Bern, Switzerland | Illumina NovaSeq 6000 | PASS | Included | |
| LR12807 | LR12807 | Cyprinoidei | Danionidae | Pectenocypris | sp | Next Generation Sequencing (NGS) Platform, University of Bern, Switzerland | Illumina NovaSeq 6000 | FAIL: generic ID doubtful; probable sample contamination with Rasbora and Pectenocypris | Not included | |
| LR15289 | LR15289 | Cyprinoidei | Danionidae | Opsaridium | sp1 | Next Generation Sequencing (NGS) Platform, University of Bern, Switzerland | Illumina NovaSeq 6000 | PASS | Included | |
| LR15290 | LR15290 | Cyprinoidei | Danionidae | Opsaridium | sp2 | Next Generation Sequencing (NGS) Platform, University of Bern, Switzerland | Illumina NovaSeq 6000 | PASS | Included | |
| LR15602 | LR15602 | Cyprinoidei | Danionidae | Opsaridium | sp3 | Next Generation Sequencing (NGS) Platform, University of Bern, Switzerland | Illumina NovaSeq 6000 | PASS | Included | |
| 142LR15666 | LR15666 | Cyprinoidei | Danionidae | Opsarius | sp1 | Next Generation Sequencing (NGS) Platform, University of Bern, Switzerland | Illumina NovaSeq 6000 | PASS | Included | |
| LR12240 | LR12240 | Cyprinoidei | Danionidae | Opsarius | sp3 | Next Generation Sequencing (NGS) Platform, University of Bern, Switzerland | Illumina NovaSeq 6000 | PASS | Included | |
| LR02136 | LR02136 | Cyprinoidei | Danionidae | Opsarius | sp 2 | Next Generation Sequencing (NGS) Platform, University of Bern, Switzerland | Illumina NovaSeq 6000 | PASS | Included | |
| LR15081 | LR15081 | Cyprinoidei | Danionidae | Pectenocypris | sp | Next Generation Sequencing (NGS) Platform, University of Bern, Switzerland | Illumina NovaSeq 6000 | PASS | Included | |
| LR15349 | LR15349 | Cyprinoidei | Danionidae | Pectenocypris | sp | Next Generation Sequencing (NGS) Platform, University of Bern, Switzerland | Illumina NovaSeq 6000 | PASS | Not included | Same lineage as LR15081 |
| LR15292 | LR15292 | Cyprinoidei | Danionidae | Raiamas | moorii | Next Generation Sequencing (NGS) Platform, University of Bern, Switzerland | Illumina NovaSeq 6000 | PASS | Included | |
| 105LR15605 | LR15605 | Cyprinoidei | Danionidae | Raiamas | sp | Next Generation Sequencing (NGS) Platform, University of Bern, Switzerland | Illumina NovaSeq 6000 | PASS | Included | |
| LR10786 | LR10786 | Cyprinoidei | Danionidae | Rasbora | cephalotaenia | Next Generation Sequencing (NGS) Platform, University of Bern, Switzerland | Illumina NovaSeq 6000 | PASS | Included | |
| KUFOS-AN-2021.2.1 | KUFOS-AN-2021.2.1 | Cyprinoidei | Danionidae | Rasbora | dandia | Agrigenome Labs Pvt. Ltd., India | Illumina NovaSeq 6000 | PASS | Included | |
| LR10789 | LR10789 | Cyprinoidei | Danionidae | Rasbora | einthovenii | Next Generation Sequencing (NGS) Platform, University of Bern, Switzerland | Illumina NovaSeq 6000 | FAIL: high read error and poor model fit in kmer profiling; genome size estimates has a % diff > 10% between k=21 and k=31 models; unrealistic high level of heterozygozity; very low BUSCO single-copy recovery for both abyss and spades assemblies | Not included | |
| LR15080 | LR15080 | Cyprinoidei | Danionidae | Rasbora | kalbarensis | Next Generation Sequencing (NGS) Platform, University of Bern, Switzerland | Illumina NovaSeq 6000 | PASS | Not included | Same lineage as LR15347 |
| LR15347 | LR15347 | Cyprinoidei | Danionidae | Rasbora | kalbarensis | Next Generation Sequencing (NGS) Platform, University of Bern, Switzerland | Illumina NovaSeq 6000 | PASS | Included | |
| LR15877 | LR15877 | Cyprinoidei | Danionidae | Rasbora | kalochroma | Next Generation Sequencing (NGS) Platform, University of Bern, Switzerland | PacBio HiFi Revio | PASS | Included | |
| LR10804 | LR10804 | Cyprinoidei | Danionidae | Rasbora | kottelati | Next Generation Sequencing (NGS) Platform, University of Bern, Switzerland | Illumina NovaSeq 6000 | PASS | Included | |
| LR09739 | LR09739 | Cyprinoidei | Danionidae | Rasbora | myersi | Next Generation Sequencing (NGS) Platform, University of Bern, Switzerland | Illumina NovaSeq 6000 | PASS | Included | |
| LR09847 | LR09847 | Cyprinoidei | Danionidae | Rasbora | trilineata | Next Generation Sequencing (NGS) Platform, University of Bern, Switzerland | Illumina NovaSeq 6000 | PASS | Included | |
| LR05463 | LR05463 | Cyprinoidei | Danionidae | Rasboroides | pallidus | Next Generation Sequencing (NGS) Platform, University of Bern, Switzerland | Illumina NovaSeq 6000 | PASS | Included | |
| LR12212 | LR12212 | Cyprinoidei | Danionidae | Rasbosoma | spilocerca | Next Generation Sequencing (NGS) Platform, University of Bern, Switzerland | Illumina NovaSeq 6000 | PASS | Included | |
| LR12213 | LR12213 | Cyprinoidei | Danionidae | Rasbosoma | spilocerca | Next Generation Sequencing (NGS) Platform, University of Bern, Switzerland | Illumina NovaSeq 6000 | PASS | Not included | Same lineage as LR12212 |
| KUFOS-AN-2021.6.1 | KUFOS-AN-2021.6.1 | Cyprinoidei | Danionidae | Salmostoma | balookee | Agrigenome Labs Pvt. Ltd., India | Illumina NovaSeq 6000 | PASS | Included | |
| 129LR02128 | LR02128 | Cyprinoidei | Danionidae | Salmostoma | sp | Next Generation Sequencing (NGS) Platform, University of Bern, Switzerland | Illumina NovaSeq 6000 | FAIL: generic ID doubtful: probable library contamination with Salmostoma and Mastacembelus | Not included | |
| EA0634 | EA0634 | Cyprinoidei | Danionidae | Thryssocypris | tonlesapensis | Next Generation Sequencing (NGS) Platform, University of Bern, Switzerland | Illumina NovaSeq 6000 | PASS | Included | |
| EA0635 | EA0635 | Cyprinoidei | Danionidae | Thryssocypris | tonlesapensis | Next Generation Sequencing (NGS) Platform, University of Bern, Switzerland | Illumina NovaSeq 6000 | PASS | Not included | Same lineage as EA0634 |
| LR08172 | LR08172 | Cyprinoidei | Danionidae | Trigonopoma | gracile | Next Generation Sequencing (NGS) Platform, University of Bern, Switzerland | Illumina NovaSeq 6000 | PASS | Included | |
| LR10181 | LR10181 | Cyprinoidei | Danionidae | Trigonopoma | pauciperforatum | Next Generation Sequencing (NGS) Platform, University of Bern, Switzerland | Illumina NovaSeq 6000 | PASS | Included | |
| LR11656 | LR11656 | Cyprinoidei | Danionidae | Trigonopoma | sp | Next Generation Sequencing (NGS) Platform, University of Bern, Switzerland | Illumina NovaSeq 6000 | PASS | Included | |
| LR08812 | LR08812 | Cyprinoidei | Danionidae | Trigonostigma | espei | Next Generation Sequencing (NGS) Platform, University of Bern, Switzerland | Illumina NovaSeq 6000 | PASS | Included | |
| LR11086 | LR11086 | Cyprinoidei | Danionidae | Trigonostigma | heteromorpha | Next Generation Sequencing (NGS) Platform, University of Bern, Switzerland | Illumina NovaSeq 6000 | PASS | Included | |
| 1736_2012 | NMBE 1064463 | Cyprinoidei | Gobionidae | Gobio | gobio | Next Generation Sequencing (NGS) Platform, University of Bern, Switzerland | Illumina NovaSeq 6000 | FAIL: high read error and poor model fit in kmer profiling; genome size estimates has a % diff > 10% between k=21 and k=31 models; unrealistic high level of heterozygozity; very low BUSCO single-copy recovery for both abyss and spades assemblies | Not included | |
| LR15569 | LR15569 | Cyprinoidei | Gobionidae | Hemibarbus | maculatus | Next Generation Sequencing (NGS) Platform, University of Bern, Switzerland | Illumina NovaSeq 6000 | PASS | Included | |
| LR15570 | LR15570 | Cyprinoidei | Gobionidae | Ladislavia | taczanowskii | Next Generation Sequencing (NGS) Platform, University of Bern, Switzerland | Illumina NovaSeq 6000 | PASS | Included | |
| LR15062 | LR15062 | Cyprinoidei | Gobionidae | Microphysogobio | tafangensis | Next Generation Sequencing (NGS) Platform, University of Bern, Switzerland | Illumina NovaSeq 6000 | PASS | Included | |
| LR15571 | LR15571 | Cyprinoidei | Gobionidae | Sarcocheilichthys | hainanensis | Next Generation Sequencing (NGS) Platform, University of Bern, Switzerland | Illumina NovaSeq 6000 | PASS | Included | |
| LR15579 | LR15579 | Cyprinoidei | Gobionidae | Squalidus | sp | Next Generation Sequencing (NGS) Platform, University of Bern, Switzerland | Illumina NovaSeq 6000 | PASS | Included | |
| LR15106 | LR15106 | Cyprinoidei | Leptobarbidae | Leptobarbus | sp | Next Generation Sequencing (NGS) Platform, University of Bern, Switzerland | Illumina NovaSeq 6000 | PASS | Included | |
| 1360_2014 | NMBE 1070731 | Cyprinoidei | Leuciscidae | Abramis | brama | Next Generation Sequencing (NGS) Platform, University of Bern, Switzerland | Illumina NovaSeq 6000 | PASS | Not included | Replaced with GCA_963993115.1 Abramis brama |
| 2180_2012 | NMBE 1065116 | Cyprinoidei | Leuciscidae | Alburnoides | bipunctatus | Next Generation Sequencing (NGS) Platform, University of Bern, Switzerland | Illumina NovaSeq 6000 | PASS | Included | |
| 108LR15742 | TCWC 20551.01 | Cyprinoidei | Leuciscidae | Chrosomus | erythrogaster | Next Generation Sequencing (NGS) Platform, University of Bern, Switzerland | Illumina NovaSeq 6000 | PASS | Included | |
| LR15250 | TCWC 20295.02 | Cyprinoidei | Leuciscidae | Cyprinella | lutrensis | Next Generation Sequencing (NGS) Platform, University of Bern, Switzerland | Illumina NovaSeq 6000 | PASS | Not included | Replaced with GCA_027497735.1 Cyprinella lepida, GCA_027361055.1 Cyprinella venusta, GCA_027497675.1 Cyprinella lutrensis |
| LR15252 | TCWC 20317.01 | Cyprinoidei | Leuciscidae | Dionda | flavipinnis | Next Generation Sequencing (NGS) Platform, University of Bern, Switzerland | Illumina NovaSeq 6000 | PASS | Included | |
| 110LR15746 | MSB 114786.001 | Cyprinoidei | Leuciscidae | Gila | pandora | Next Generation Sequencing (NGS) Platform, University of Bern, Switzerland | Illumina NovaSeq 6000 | PASS | Included | |
| LR15253 | TCWC 20318.01 | Cyprinoidei | Leuciscidae | Hybognathus | nuchalis | Next Generation Sequencing (NGS) Platform, University of Bern, Switzerland | Illumina NovaSeq 6000 | PASS | Not included | Replaced with GCA_030378365.1 Hybognathus amarus |
| 2223_2013 | NMBE 1068595 | Cyprinoidei | Leuciscidae | Leuciscus | leuciscus | Next Generation Sequencing (NGS) Platform, University of Bern, Switzerland | Illumina NovaSeq 6000 | PASS | Included | |
| LR15254 | TCWC 20317.03 | Cyprinoidei | Leuciscidae | Notropis | chalybaeus | Next Generation Sequencing (NGS) Platform, University of Bern, Switzerland | Illumina NovaSeq 6000 | PASS | Included | |
| 2564_2011 | NMBE 1061900 | Cyprinoidei | Leuciscidae | Phoxinus | csikii | Next Generation Sequencing (NGS) Platform, University of Bern, Switzerland | Illumina NovaSeq 6000 | PASS | Not included | Replaced with GCA_949152265.1 Phoxinus phoxinus |
| 109LR15744 | MSB 113977-28 | Cyprinoidei | Leuciscidae | Ptychocheilus | lucius | Next Generation Sequencing (NGS) Platform, University of Bern, Switzerland | Illumina NovaSeq 6000 | PASS | Included | |
| LR15251 | TCWC 19726.02 | Cyprinoidei | Leuciscidae | Rhinichthys | cataractae | Next Generation Sequencing (NGS) Platform, University of Bern, Switzerland | Illumina NovaSeq 6000 | PASS | Included | |
| LR15572 | LR15572 | Cyprinoidei | Leuciscidae | Rhynchocypris | lagowskii | Next Generation Sequencing (NGS) Platform, University of Bern, Switzerland | Illumina NovaSeq 6000 | PASS | Included | |
| 2333_2013 | NMBE 1068705 | Cyprinoidei | Leuciscidae | Rutilus | rutilus | Next Generation Sequencing (NGS) Platform, University of Bern, Switzerland | Illumina NovaSeq 6000 | PASS | Not included | Replaced with GCA_951802725.1 Rutilus rutilus |
| 1010_2016 | NMBE 1074568 | Cyprinoidei | Leuciscidae | Scardinius | sp | Next Generation Sequencing (NGS) Platform, University of Bern, Switzerland | Illumina NovaSeq 6000 | PASS | Not included | Replaced with GCA_024453875.1 Scardinius erythrophthalmus |
| LR15249 | TCWC 15788.06 | Cyprinoidei | Leuciscidae | Semotilus | atromaculatus | Next Generation Sequencing (NGS) Platform, University of Bern, Switzerland | Illumina NovaSeq 6000 | PASS | Not included | Replaced with GCA_031834385.1 Semotilus atromaculatus |
| 1104_2014 | NMBE 1069134 | Cyprinoidei | Leuciscidae | Squalius | cephalus | Next Generation Sequencing (NGS) Platform, University of Bern, Switzerland | Illumina NovaSeq 6000 | PASS | Not included | Replaced with GCA_949319135.1 Squalius cephalus |
| 1450_2013 | NMBE 1067637 | Cyprinoidei | Leuciscidae | Telestes | muticellus | Next Generation Sequencing (NGS) Platform, University of Bern, Switzerland | Illumina NovaSeq 6000 | PASS | Included | |
| LR15325 | LR15325 | Cyprinoidei | Paedocyprididae | Paedocypris | sp2 | Next Generation Sequencing (NGS) Platform, University of Bern, Switzerland | Illumina NovaSeq 6000 | PASS | Included | |
| LR15863A | LR15863A | Cyprinoidei | Paedocyprididae | Paedocypris | sp1 | Next Generation Sequencing (NGS) Platform, University of Bern, Switzerland | PacBio HiFi Sequel Iie | PASS | Included | |
| LR09467 | LR09467 | Cyprinoidei | Psilorhynchidae | Psilorhynchus | sp1 | Next Generation Sequencing (NGS) Platform, University of Bern, Switzerland | Illumina NovaSeq 6000 | PASS | Included | |
| LR15089 | LR15089 | Cyprinoidei | Psilorhynchidae | Psilorhynchus | sp2 | Next Generation Sequencing (NGS) Platform, University of Bern, Switzerland | Illumina NovaSeq 6000 | PASS | Included | |
| LR15870 | LR15870 | Cyprinoidei | Sundadanionidae | Sundadanio | atomus | Next Generation Sequencing (NGS) Platform, University of Bern, Switzerland | PacBio HiFi Revio | PASS | Included | |
| 100LR08982 | LR08982 | Cyprinoidei | Sundadanionidae | Fangfangia | spinicleithralis | Next Generation Sequencing (NGS) Platform, University of Bern, Switzerland | Illumina NovaSeq 6000 | PASS | Included | |
| LR10372 | LR10372 | Cyprinoidei | Sundadanionidae | Sundadanio | goblinus | Next Generation Sequencing (NGS) Platform, University of Bern, Switzerland | Illumina NovaSeq 6000 | PASS | Included | |
| LR15082 | LR15082 | Cyprinoidei | Sundadanionidae | Sundadanio | margarition | Next Generation Sequencing (NGS) Platform, University of Bern, Switzerland | Illumina NovaSeq 6000 | PASS | Included | |
| 101LR12751 | LR12751 | Cyprinoidei | Sundadanionidae | Sundadanio | retiarius | Next Generation Sequencing (NGS) Platform, University of Bern, Switzerland | Illumina NovaSeq 6000 | PASS | Included | |
| LR15110 | LR15110 | Cyprinoidei | Tanichthyidae | Tanichthys | albonudes | Next Generation Sequencing (NGS) Platform, University of Bern, Switzerland | Illumina NovaSeq 6000 | PASS | Included | |
| LR02462 | LR02462 | Cyprinoidei | Tanichthyidae | Tanichthys | micagemmae | Next Generation Sequencing (NGS) Platform, University of Bern, Switzerland | Illumina NovaSeq 6000 | PASS | Included | |
| 1545_2010 | NMBE 1058507 | Cyprinoidei | Tincidae | Tinca | tinca | Next Generation Sequencing (NGS) Platform, University of Bern, Switzerland | Illumina NovaSeq 6000 | PASS | Included | |
| 117LR15718 | LR15718 | Cyprinoidei | Xenocyprididae | Aphyocypris | chinensis | Next Generation Sequencing (NGS) Platform, University of Bern, Switzerland | Illumina NovaSeq 6000 | FAIL: probable hybrid sample of Nipponocypris X Aphyocypris | Not included | |
| 118LR15719 | LR15719 | Cyprinoidei | Xenocyprididae | Candidia | pingtungensis | Next Generation Sequencing (NGS) Platform, University of Bern, Switzerland | Illumina NovaSeq 6000 | PASS | Included | |
| 103LR15622 | LR15622 | Cyprinoidei | Xenocyprididae | Hemiculter | sp | Next Generation Sequencing (NGS) Platform, University of Bern, Switzerland | Illumina NovaSeq 6000 | PASS | Included | |
| 102LR15621 | LR15621 | Cyprinoidei | Xenocyprididae | Opsariichthys | bidens | Next Generation Sequencing (NGS) Platform, University of Bern, Switzerland | Illumina NovaSeq 6000 | PASS | Included | |
| 128LR15724 | LR15724 | Cyprinoidei | Xenocyprididae | Opsariichthys | pachycephalus | Next Generation Sequencing (NGS) Platform, University of Bern, Switzerland | Illumina NovaSeq 6000 | PASS | Included | |
| LR15574 | LR15574 | Cyprinoidei | Xenocyprididae | Oxygaster | pointoni | Next Generation Sequencing (NGS) Platform, University of Bern, Switzerland | Illumina NovaSeq 6000 | PASS | Included | |
| LR15097 | LR15097 | Gyrinocheiloidei | Gyrinocheilidae | Gyrinocheilus | pustulosus | Next Generation Sequencing (NGS) Platform, University of Bern, Switzerland | Illumina NovaSeq 6000 | PASS | Included | |
| 104EA0138 | EA0138 | Gyrinocheiloidei | Gyrinocheilidae | Gyrinocheilus | sp | Next Generation Sequencing (NGS) Platform, University of Bern, Switzerland | Illumina NovaSeq 6000 | PASS | Included | |

**Supplementary Table 2.** NCBI genomes downloaded in this study.

| Voucher reads | Group | Order | Suborder | Family | Genus | Species | Source | Assembly.Accession | Assembly.Stats.Total.N | Assembly.Sequencing.Tech | Assembly.BioProject.Accession | Assembly.BioSample.Accession | Assembly.Name | Annotation.Name | Assembly.Stats.Total.Sequence.Length | Assembly.Level | Assembly.Release.Date | WGS.project.accession | Assembly.Submitter | BioProject | BioSample | SRA | SRA RUNs | Publication | Phylogenomics dataset | Phylogenomics dataset Remarks |
|---|---|---|---|---|---|---|---|---|---|---|---|---|---|---|---|---|---|---|---|---|---|---|---|---|---|---|
| GCF 019703515. | Ingroup | Cypriniformes | Catostomoidei | Catostomidae | Myxocyprinus | asiaticus | NCBI | GCF 019703515.2 | 50 | Oxford Nanopore GridION; Illumina | PRJNA739167 | SAMN19774990 | UBuf Myxa 2 | GCF 019703515.2-RS 2023 02 | 2340848162 | Chromosome | 26/10/2022 | JAIGOD02 | University at Buffalo | NA | NA | NA | NA | https://doi.org/10.1093/gbe/evab190 | Included | NA |
| GCA 036877625. | Ingroup | Cypriniformes | Cobitoidei | Botiidae | Ambastaia | sidthimunki | NCBI | GCA 036877625.1 | | Illumina NovaSeq | PRJNA1067307 | SAMN39625708 | ASM3687762v1 | NA | 923471233 | Scaffold | 26/02/2024 | JAZGTK01 | University of Basel | NA | NA | NA | NA | https://doi.org/10.1016/j.cub.2024.02.022 | Not included | Replaced with LR15330 Ambastaia sidthimunki |
| GCA 036877575. | Ingroup | Cypriniformes | Cobitoidei | Botiidae | Botia | almorhae | NCBI | GCA 036877575.1 | | Illumina NovaSeq | PRJNA1067307 | SAMN39625707 | ASM3687757v1 | NA | 894053861 | Scaffold | 26/02/2024 | JAZGTJ01 | University of Basel | NA | NA | NA | NA | https://doi.org/10.1016/j.cub.2024.02.022 | Included | NA |
| GCA 036877585. | Ingroup | Cypriniformes | Cobitoidei | Botiidae | Botia | kubotai | NCBI | GCA 036877585.1 | | Illumina NovaSeq | PRJNA1067307 | SAMN39625706 | ASM3687758v1 | NA | 857539002 | Scaffold | 26/02/2024 | JAZGTI01 | University of Basel | NA | NA | NA | NA | https://doi.org/10.1016/j.cub.2024.02.022 | Not included | Replaced with LR15149 Botia kubotai |
| GCA 036877525. | Ingroup | Cypriniformes | Cobitoidei | Botiidae | Chromobotia | macracanthus | NCBI | GCA 036877525.1 | | Illumina NovaSeq; Oxford Nanopore PromethION | PRJNA1067307 | SAMN39546219 | ASM3687752v1 | NA | 783742438 | Scaffold | 26/02/2024 | JAZGTH01 | University of Basel | NA | NA | NA | NA | https://doi.org/10.1016/j.cub.2024.02.022 | Not included | Replaced with LR15148 Chromobotia macracanthus |
| GCA 036877545. | Ingroup | Cypriniformes | Cobitoidei | Botiidae | Yasuhikotakia | modesta | NCBI | GCA 036877545.1 | | Illumina NovaSeq | PRJNA1067307 | SAMN39625709 | ASM3687754v1 | NA | 636809230 | Scaffold | 26/02/2024 | JBAJNP01 | University of Basel | NA | NA | NA | NA | https://doi.org/10.1016/j.cub.2024.02.022 | Included | NA |
| GCF 027580225. | Ingroup | Cypriniformes | Cobitoidei | Cobitidae | Misgurnus | anguillicaudatus | NCBI | GCF 027580225.1 | 25 | Illumina HiSeq; PacBio | PRJNA812369 | SAMN26377316 | HAU Mang 1.0 | GCF 027580225.1-RS 2023 04 | 1104569478 | Chromosome | 09/01/2023 | JALDXJ01 | Huazhong Agriculture University | NA | NA | NA | NA | https://doi.org/10.1186/s12915-023-01517-1 | Included | NA |
| GCA 025389145. | Ingroup | Cypriniformes | Cobitoidei | Cobitidae | Misgurnus | mizolepis | NCBI | GCA 025389145.1 | | PacBio Sequel; Illumina NovaSeq | PRJNA751427 | SAMN20518172 | NIFS Mimiz 1.0 | NA | 1112094387 | Scaffold | 25/09/2022 | JAIFKI01 | National Institute of Fisheries Science | NA | NA | NA | NA | https://doi.org/10.3389/fmars.2021.799148 | Included | NA |
| GCA 019155185. | Ingroup | Cypriniformes | Cobitoidei | Gastromyzontidae | Beaufortia | kweichowensis | NCBI | GCA 019155185.1 | 25 | Oxford Nanopore GridION | PRJNA650219 | SAMN15699278 | ASM1915518v1 | NA | 448565924 | Chromosome | 07/07/2021 | JAGGEC01 | Institute of Hydrobiology | NA | NA | NA | NA | https://doi.org/10.1111/1755-0998.13400 | Included | NA |
| GCA 947034865. | Ingroup | Cypriniformes | Cobitoidei | Nemacheilidae | Barbatula | barbatula | NCBI | GCA 947034865.1 | 25 | PacBio,Arima2 | PRJEB56420 | SAMEA11296540 | fBarBar1.1 | NA | 617646710 | Chromosome | 07/10/2022 | CAMOOH01 | WELLCOME SANGER INSTITUTE | NA | NA | NA | NA | https://doi.org/10.12688/wellcomeopenres.20293.1 | Included | NA |
| GCF 015846415. | Ingroup | Cypriniformes | Cobitoidei | Nemacheilidae | Triplophysa | dalaica | NCBI | GCF 015846415.1 | 25 | PacBio Sequel II | PRJNA624716 | SAMN14582390 | ASM1584641v1 | GCF 015846415.1-RS 2023 06 | 607923607 | Chromosome | 11/12/2020 | JABZFK01 | HeNan Normal University and Wuhan diggers | NA | NA | NA | NA | https://doi.org/10.1093/gbe/evab153 | Included | NA |
| GCF 024868665. | Ingroup | Cypriniformes | Cobitoidei | Nemacheilidae | Triplophysa | rosa | NCBI | GCF 024868665.1 | 25 | Illumina; PacBio RSII | PRJNA673076 | SAMN16587819 | Trosa 1v2 | GCF 024868665.1-RS 2023 06 | 684805028 | Chromosome | 14/09/2022 | JAFHDT01 | the National Natural Science Foundation of China | NA | NA | NA | NA | https://doi.org/10.1111/mec.16700 | Included | NA |
| GCA 006030095. | Ingroup | Cypriniformes | Cobitoidei | Nemacheilidae | Triplophysa | siluroides | NCBI | GCA 006030095.1 | | PacBio Sequel; Illumina HiSeq | PRJNA543715 | SAMN11775600 | ASM603009v1 | NA | 583428323 | Scaffold | 05/06/2019 | VCII01 | Institute of Hydrobiology | NA | NA | NA | NA | https://doi.org/10.3389/fgene.2019.00991 | Included | NA |
| GCA 008369825. | Ingroup | Cypriniformes | Cobitoidei | Nemacheilidae | Triplophysa | tibetana | NCBI | GCA 008369825.1 | 25 | PacBio Sequel; Illumina HiSeq | PRJNA526292 | SAMN11094431 | ASM836982v1 | Annotation submitted by Googlgene | 652930317 | Chromosome | 12/09/2019 | SOYY01 | Googlgene | NA | NA | NA | NA | https://doi.org/10.1111/1755-0998.13021 | Included | NA |
| GCA 033220385. | Ingroup | Cypriniformes | Cobitoidei | Nemacheilidae | Triplophysa | yarkandensis | NCBI | GCA 033220385.1 | 25 | PacBio Sequel | PRJNA995909 | SAMN36510128 | ASM3322038v1 | NA | 520782176 | Chromosome | 03/11/2023 | JAWHON01 | Huazhong Agricultural University | NA | NA | NA | NA | https://doi.org/10.1038/s41597-023-02900-x | Included | NA |
| GCA 037043235. | Ingroup | Cypriniformes | Cyprinoidei | Acheilognathidae | Rhodeus | uyekii | NCBI | GCA 037043235.1 | | PacBio | PRJNA1033701 | SAMN38042216 | Rhuy ver1 | NA | 894549330 | Scaffold | 06/03/2024 | JAWRVZ01 | Nakdonggang National Institute of Biological Resources | NA | NA | NA | NA | https://doi.org/ 10.3390/ijms25126689 | Included | NA |
| GCA 036784905. | Ingroup | Cypriniformes | Cyprinoidei | Cyprinidae | Acrossocheilus | fasciatus | NCBI | GCA 036784905.1 | 25 | PacBio; Illumina | PRJNA1045882 | SAMN38725585 | ASM3678490v1 | NA | 899129631 | Chromosome | 23/02/2024 | JAXUIB01 | Shanghai Ocean University | NA | NA | NA | NA | https://doi.org/10.1038/s41597-024-02999-6 | Not included | Replaced with GCA 039880705.1 Acrossocheilus fasciatus |
| GCA 039880705. | Ingroup | Cypriniformes | Cyprinoidei | Cyprinidae | Acrossocheilus | fasciatus | NCBI | GCA 039880705.1 | 25 | Illumina; PacBio | PRJNA1049304 | SAMN39587862 | ASM3988070v1 | NA | 885609419 | Chromosome | 28/05/2024 | JAZDCR01 | Shanghai Ocean University | NA | NA | NA | NA | https://doi.org/10.1038/s41597-024-02999-6 | Included | NA |
| GCA 023376895. | Ingroup | Cypriniformes | Cyprinoidei | Cyprinidae | Aspiorhynchus | laticeps | NCBI | GCA 023376895.1 | 25 | PacBio Sequel | PRJNA832538 | SAMN27865501 | ASM2337689v1 | NA | 1583238279 | Chromosome | 17/05/2022 | JALXFT01 | Institute of Oceanology, Chinese Academy of Sciences | NA | NA | NA | NA | https://doi.org/10.1038/s41597-022-01671-1 | Included | NA |
| GCA 936440315. | Ingroup | Cypriniformes | Cyprinoidei | Cyprinidae | Barbus | barbus | NCBI | GCA 936440315.1 | 50 | NA | PRJEB52077 | SAMEA13335783 | fBarBab1.1 | NA | 1584913304 | Chromosome | 03/05/2022 | CAKZFL01 | WELLCOME SANGER INSTITUTE | NA | NA | NA | NA | https://doi.org/10.12688/wellcomeopenres.19332.1 | Included | NA |
| GCF 003368295. | Ingroup | Cypriniformes | Cyprinoidei | Cyprinidae | Carassius | auratus | NCBI | GCF 003368295.1 | 59 | PacBio RSII | PRJNA481500 | SAMN09670328 | ASM336829v1 | NCBI Carassius auratus Annotation Release 100 | 1820618472 | Chromosome | 09/08/2018 | OPKE01 | National Institutes of Health | NA | NA | NA | NA | https://doi.org/10.1126/sciadv.aav0547 | Included | NA |
| GCF 963082965. | Ingroup | Cypriniformes | Cyprinoidei | Cyprinidae | Carassius | carassius | NCBI | GCF 963082965.1 | 50 | PacBio,Arima2 | PRJEB64837 | SAMEA11296534 | fCarCar2.1 | GCF 963082965.1-RS 2023 09 | 1684296438 | Chromosome | 09/08/2023 | CAUJCD01 | WELLCOME SANGER INSTITUTE | NA | NA | NA | NA | NA | Included | NA |
| GCA 023724075. | Ingroup | Cypriniformes | Cyprinoidei | Cyprinidae | Carassius | gibelio | NCBI | GCA 023724075.1 | 50 | PacBio HiFi; OmniC Illumina reads | PRJNA779620 | SAMN23046545 | carGib1.2-hapl.a | NA | 1949148655 | Chromosome | 16/06/2022 | JAJOLA01 | IGB Leibniz-Institute of Freshwater Ecology and Inland Fisheries | NA | NA | NA | NA | https://doi.org/10.1038/s41467-022-31515-w | Included | NA |
| GCF 018340385. | Ingroup | Cypriniformes | Cyprinoidei | Cyprinidae | Cyprinus | carpio | NCBI | GCF 018340385.1 | 50 | PacBio; Oxford Nanopore; Illumina HiSeq | PRJNA682709 | SAMN17005855 | ASM1834038v1 | NCBI Cyprinus carpio Annotation Release 101 | 1680118328 | Chromosome | 12/05/2021 | JAEOAB01 | Chinese Academy of Fishery Sciences | NA | NA | NA | NA | https://doi.org/10.1038/s41588-021-00933-9 | Included | NA |
| GCA 034779775. | Ingroup | Cypriniformes | Cyprinoidei | Cyprinidae | Enteromius | afrohamiltoni | NCBI | GCA 034779775.1 | | Illumina | PRJNA915624 | SAMN32526004 | ASM3477977v1 | NA | 663517611 | Scaffold | 28/12/2023 | JAUYWG01 | Iridian Genomes | NA | NA | NA | NA | https://doi.org/10.56179/001c.116676 | Included | NA |
| GCA 032359395. | Ingroup | Cypriniformes | Cyprinoidei | Cyprinidae | Enteromius | anoplus | NCBI | GCA 032359395.1 | | Illumina | PRJNA917146 | SAMN32526002 | ASM3235939v1 | NA | 737678727 | Scaffold | 04/10/2023 | JAUMHG01 | Iridian Genomes | NA | NA | NA | NA | https://doi.org/10.56179/001c.116676 | Included | NA |
| GCA 032357425. | Ingroup | Cypriniformes | Cyprinoidei | Cyprinidae | Enteromius | argenteus | NCBI | GCA 032357425.1 | | Illumina | PRJNA917131 | SAMN32525989 | ASM3235742v1 | NA | 681601004 | Scaffold | 04/10/2023 | JAUCMR01 | Iridian Genomes | NA | NA | NA | NA | https://doi.org/10.56179/001c.116676 | Included | NA |
| GCA 032361145. | Ingroup | Cypriniformes | Cyprinoidei | Cyprinidae | Enteromius | fasciolatus | NCBI | GCA 032361145.1 | | Illumina | PRJNA917138 | SAMN32525995 | ASM3236114v1 | NA | 719794475 | Scaffold | 04/10/2023 | JAUMHB01 | Iridian Genomes | NA | NA | NA | NA | https://doi.org/10.56179/001c.116676 | Not included | Replaced with GCA 032359395.1 Enteromius anoplus |
| GCA 032359235. | Ingroup | Cypriniformes | Cyprinoidei | Cyprinidae | Enteromius | mattozi | NCBI | GCA 032359235.1 | | Illumina | PRJNA915750 | SAMN32525988 | ASM3235923v1 | NA | 727350904 | Scaffold | 04/10/2023 | JAUMHE01 | Iridian Genomes | NA | NA | NA | NA | https://doi.org/10.56179/001c.116676 | Included | NA |
| GCA 032361165. | Ingroup | Cypriniformes | Cyprinoidei | Cyprinidae | Enteromius | paludinosus | NCBI | GCA 032361165.1 | | Illumina | PRJNA917134 | SAMN32525992 | ASM3236116v1 | NA | 745488820 | Scaffold | 04/10/2023 | JAUMHO01 | Iridian Genomes | NA | NA | NA | NA | https://doi.org/10.56179/001c.116676 | Not included | Replaced with GCA 032359435.1 Enteromius unitaeniatus |
| GCA 032357445. | Ingroup | Cypriniformes | Cyprinoidei | Cyprinidae | Enteromius | radiatus | NCBI | GCA 032357445.1 | | Illumina | PRJNA917141 | SAMN32525998 | ASM3235744v1 | NA | 616678458 | Scaffold | 04/10/2023 | JAUCMS01 | Iridian Genomes | NA | NA | NA | NA | https://doi.org/10.56179/001c.116676 | Included | NA |
| GCA 032357475. | Ingroup | Cypriniformes | Cyprinoidei | Cyprinidae | Enteromius | thamalakanensis | NCBI | GCA 032357475.1 | | Illumina | PRJNA917139 | SAMN32525996 | ASM3235747v1 | NA | 673856647 | Scaffold | 04/10/2023 | JAUCMT01 | Iridian Genomes | NA | NA | NA | NA | https://doi.org/10.56179/001c.116676 | Not included | Replaced with GCA 032357425.1 Enteromius argenteus |
| GCA 032275025. | Ingroup | Cypriniformes | Cyprinoidei | Cyprinidae | Enteromius | treurensis | NCBI | GCA 032275025.1 | | Illumina | PRJNA917145 | SAMN32526001 | ASM3227502v1 | NA | 665121873 | Scaffold | 02/10/2023 | JAUCMU01 | Iridian Genomes | NA | NA | NA | NA | https://doi.org/10.56179/001c.116676 | Not included | Replaced with GCA 032357425.1 Enteromius argenteus |
| GCA 032361065. | Ingroup | Cypriniformes | Cyprinoidei | Cyprinidae | Enteromius | trimaculatus | NCBI | GCA 032361065.1 | | Illumina | PRJNA917133 | SAMN32525991 | ASM3236106v1 | NA | 714608295 | Scaffold | 04/10/2023 | JAUMHP01 | Iridian Genomes | NA | NA | NA | NA | https://doi.org/10.56179/001c.116676 | Not included | Replaced with GCA 032359435.1 Enteromius unitaeniatus |
| GCA 032359435. | Ingroup | Cypriniformes | Cyprinoidei | Cyprinidae | Enteromius | unitaeniatus | NCBI | GCA 032359435.1 | | Illumina | PRJNA917137 | SAMN32525994 | ASM3235943v1 | NA | 643365342 | Scaffold | 04/10/2023 | JAUMHI01 | Iridian Genomes | NA | NA | NA | NA | https://doi.org/10.56179/001c.116676 | Included | NA |
| GCA 032353835. | Ingroup | Cypriniformes | Cyprinoidei | Cyprinidae | Enteromius | viviparus | NCBI | GCA 032353835.1 | | Illumina | PRJNA917151 | SAMN35339798 | ASM3235383v1 | NA | 689049118 | Scaffold | 04/10/2023 | JAUCNO01 | Iridian Genomes | NA | NA | NA | NA | https://doi.org/10.56179/001c.116676 | Not included | Replaced with GCA 032357445.1 ASM3235744v1 Enteromius radiatus |
| GCA 024082105. | Ingroup | Cypriniformes | Cyprinoidei | Cyprinidae | Gymnocypris | eckloni | NCBI | GCA 024082105.1 | | PacBio | PRJNA835611 | SAMN28105652 | ASM2408210v1 | NA | 918681488 | Scaffold | 30/06/2022 | JAMHKY01 | Qinghai University | NA | NA | NA | NA | https://doi.org/10.1038/s41597-022-01595-w | Not included | Replaced with GCA 027564155.1 Gymnocypris eckloni scoliostomus |
| GCA 027564155. | Ingroup | Cypriniformes | Cyprinoidei | Cyprinidae | Gymnocypris | eckloni-scoliostomus | NCBI | GCA 027564155.1 | 25 | Oxford Nanopore | PRJNA771798 | SAMN22344571 | ASM2756415v1 | NA | 948071110 | Chromosome | 09/01/2023 | JAJHNS01 | Institute of Hydrobiology, Chinese Academy of Sciences | NA | NA | NA | NA | https://doi.org/10.1093/nsr/nwac291 | Included | NA |
| GCA 012976165. | Ingroup | Cypriniformes | Cyprinoidei | Cyprinidae | Labeo | catla | NCBI | GCA 012976165.1 | | Illumina NextSeq; Oxford Nanopore MinION | PRJNA557138 | SAMN12390082 | CIFA Catla 1.0 | NA | 1019782285 | Scaffold | 05/05/2020 | VONZ01 | ICAR-Central Institute of Freshwater Aquaculture | NA | NA | NA | NA | https://doi.org/10.1186/s13104-020-05240-w | Included | NA |
| GCF 022985175. | Ingroup | Cypriniformes | Cyprinoidei | Cyprinidae | Labeo | rohita | NCBI | GCF 022985175.1 | 25 | Oxford Nanopore GridION; Illumina HiSeq | PRJNA650519 | SAMN15716776 | IGBB LRoh.1.0 | GCF 022985175.1-RS 2023 02 | 1126541939 | Chromosome | 19/04/2022 | JACTAM01 | Mississippi State University | NA | NA | NA | NA | https://doi.org/10.1093/g3journal/jkad009 | Included | NA |
| GCA 034780575. | Ingroup | Cypriniformes | Cyprinoidei | Cyprinidae | Labeobarbus | marequensis | NCBI | GCA 034780575.1 | | Illumina | PRJNA996131 | SAMN32526010 | ASM3478057v1 | NA | 1571828143 | Scaffold | 28/12/2023 | JAUZSZ01 | Iridian Genomes | NA | NA | NA | NA | NA | Included | NA |
| GCF 012432095. | Ingroup | Cypriniformes | Cyprinoidei | Cyprinidae | Onychostoma | macrolepis | NCBI | GCF 012432095.1 | 25 | Oxford Nanopore PromethION; Illumina HiSeq; Hi-C; Bionano optical mapping | PRJNA608076 | SAMN14163390 | ASM1243209v1 | GCF 012432095.1-RS 2023 08 | 886569841 | Chromosome | 17/04/2020 | JAAMOB01 | SOUTHWEST UNIVERSITY | NA | NA | NA | NA | https://doi.org/10.1111/1755-0998.13190 | Included | NA |
| GCA 003573665. | Ingroup | Cypriniformes | Cyprinoidei | Cyprinidae | Oxygymnocypris | stewartii | NCBI | GCA 003573665.1 | | PacBio Sequel | PRJNA485858 | SAMN09762613 | Novo Ost 1.0 | NA | 1849224471 | Scaffold | 18/09/2018 | OVTF01 | Novogene Bioinformatics Technology Corporation | NA | NA | NA | NA | https://doi.org/10.1038/sdata.2019.9 | Included | NA |
| GCA 004124795. | Ingroup | Cypriniformes | Cyprinoidei | Cyprinidae | Poropuntius | huangchuchieni | NCBI | GCA 004124795.1 | | Illumina HiSeq | PRJNA511031 | SAMN10662616 | ASM412479v1 | NA | 760177161 | Scaffold | 31/01/2019 | SCHK01 | Xiamen University | NA | NA | NA | NA | https://doi.org/10.1111/1755-0998.13365 | Included | NA |
| GCF 018831695. | Ingroup | Cypriniformes | Cyprinoidei | Cyprinidae | Puntigrus | tetrazona | NCBI | GCF 018831695.1 | 25 | Illumina HiSeq; Nanopore | PRJNA683758 | SAMN17036344 | ASM1883169v1 | NCBI Puntigrus tetrazona Annotation Release 100 | 730801986 | Chromosome | 10/06/2021 | JAEOBD01 | China Academy of Fishery Sciencey | NA | NA | NA | NA | https://doi.org/10.1038/s41588-021-00933-9 | Included | NA |
| GCF 001515605. | Ingroup | Cypriniformes | Cyprinoidei | Cyprinidae | Sinocyclocheilus | anshuiensis | NCBI | GCF 001515605.1 | | Illumina Hiseq2000 | PRJNA274017 | SAMN03320099 | SAMN03320099.WGS v1. | NCBI Sinocyclocheilus anshuiensis Annotation Release 100 | 1632701648 | Scaffold | 14/12/2015 | LAVE01 | BGI, Shenzhen | NA | NA | NA | NA | https://doi.org/10.1186/s12915-015-0223-4 | Included | NA |
| GCF 001515645. | Ingroup | Cypriniformes | Cyprinoidei | Cyprinidae | Sinocyclocheilus | grahami | NCBI | GCF 001515645.1 | | Illumina Hiseq2000 | PRJNA274017 | SAMN03320097 | SAMN03320097.WGS v1. | NCBI Sinocyclocheilus grahami Annotation Release 100 | 1750271176 | Scaffold | 16/12/2015 | LCYO01 | BGI, Shenzhen | NA | NA | NA | NA | https://doi.org/10.1186/s12915-015-0223-4 | Included | NA |
| GCA 018148995. | Ingroup | Cypriniformes | Cyprinoidei | Cyprinidae | Sinocyclocheilus | maitianheensis | NCBI | GCA 018148995.1 | | Illumina HiSeq | PRJNA669120 | SAMN16435219 | ASM1814899v1 | NA | 1680372801 | Scaffold | 28/04/2021 | JADLZM01 | BGI Academy of Marine Sciences | NA | NA | NA | NA | https://doi.org/10.3389/fgene.2021.736500 | Included | NA |
| GCF 001515625. | Ingroup | Cypriniformes | Cyprinoidei | Cyprinidae | Sinocyclocheilus | rhinocerous | NCBI | GCF 001515625.1 | | Illumina Hiseq2000 | PRJNA274017 | SAMN03320098 | SAMN03320098 v1.1 | NCBI Sinocyclocheilus rhinocerous Annotation Release 100 | 1655769822 | Scaffold | 14/12/2015 | LAVF01 | BGI, Shenzhen | NA | NA | NA | NA | https://doi.org/10.1186/s12915-015-0223-4 | Included | NA |
| fAmbSpp1 | Ingroup | Cypriniformes | Cyprinoidei | Danionidae | Amblypharyngodon | sp | NCBI-SRA | | | NA | NA | NA | NA | NA | NA | NA | NA | NA | NA | PRJEB19273 | SAMEA10413023 | ERS358363 | ERR3332378;ERR333238 | NA | Included | NA |
| fBorBri1 | Ingroup | Cypriniformes | Cyprinoidei | Danionidae | Boraras | brigittae | NCBI-SRA | | | NA | NA | NA | NA | NA | NA | NA | NA | NA | NA | PRJEB19273 | SAMEA10402641 | ERS358366 | ERR3332380;ERR333238 | NA | Not included | Replaced with LR15871 Boraras brigittae |
| fBorMac1 | Ingroup | Cypriniformes | Cyprinoidei | Danionidae | Boraras | maculatus | NCBI-SRA | | | NA | NA | NA | NA | NA | NA | NA | NA | NA | NA | PRJEB19273 | SAMEA10402640 | ERS358368 | ERR3332319;ERR333232 | NA | Included | NA |
| fSalSpp1 | Ingroup | Cypriniformes | Cyprinoidei | Danionidae | Cabdio | sp | NCBI-SRA | | | NA | NA | NA | NA | NA | NA | NA | NA | NA | NA | PRJEB19273 | SAMEA10413023 | ERS358367 | ERR3332344;ERR333234 | NA | Not included | Replaced with LR15567 Cabdio morar |
| fCheCac1 | Ingroup | Cypriniformes | Cyprinoidei | Danionidae | Chela | cachius | NCBI-SRA | | | NA | NA | NA | NA | NA | NA | NA | NA | NA | NA | PRJEB19273 | SAMEA10413024 | ERS358366 | ERR3332366;ERR333238 | NA | Included | NA |
| GCF 903798145. | Ingroup | Cypriniformes | Cyprinoidei | Danionidae | Danio | aesculapii | NCBI | GCF 903798145.1 | 25 | NA | PRJEB38583 | SAMEA104240212 | fDanAes4.1 | GCF 903798145.1-RS 2023 05 | 1381485597 | Chromosome | 27/06/2020 | CAIGOR01 | SC | NA | NA | NA | NA | https://doi.org/10.12688/wellcomeopenres.25028.1 | Included | NA |
| GCA 903798035. | Ingroup | Cypriniformes | Cyprinoidei | Danionidae | Danio | albolineatus | NCBI | GCA 903798035.1 | | NA | PRJEB38565 | SAMEA104026392 | fDanAlb1.1 | NA | 1463018096 | Scaffold | 16/06/2020 | CAIGOO01 | SC | NA | NA | NA | NA | https://doi.org/10.12688/wellcomeopenres.25116.1 | Included | NA |
| GCA 903798125. | Ingroup | Cypriniformes | Cyprinoidei | Danionidae | Danio | choprai | NCBI | GCA 903798125.1 | | NA | PRJEB38567 | SAMEA104026429 | fDanCho1.1 | NA | 1102162037 | Scaffold | 16/06/2020 | CAIGOW01 | SC | NA | NA | NA | NA | https://doi.org/10.12688/wellcomeopenres.25116.1 | Included | NA |
| GCA 903798115. | Ingroup | Cypriniformes | Cyprinoidei | Danionidae | Danio | jaintianensis | NCBI | GCA 903798115.1 | | NA | PRJEB38569 | SAMEA104026422 | fDanJai1.1 | NA | 1122503336 | Scaffold | 16/06/2020 | CAIGOV01 | SC | NA | NA | NA | NA | https://doi.org/10.12688/wellcomeopenres.25116.1 | Included | NA |
| GCA 903798195. | Ingroup | Cypriniformes | Cyprinoidei | Danionidae | Danio | kyathit | NCBI | GCA 903798195.1 | 25 | NA | PRJEB38580 | SAMEA104130240 | fDanKya3.1 | NA | 1708446541 | Chromosome | 27/06/2020 | CAIGOS01 | SC | NA | NA | NA | NA | https://doi.org/10.12688/wellcomeopenres.25031.1 | Included | NA |
| GCA 944039275. | Ingroup | Cypriniformes | Cyprinoidei | Danionidae | Danio | rerio | NCBI | GCA 944039275.1 | 25 | NA | PRJEB53407 | SAMEA14088276 | fDanRer4.1 | NA | 1413659274 | Chromosome | 24/06/2022 | CALUEM01 | WELLCOME SANGER INSTITUTE | NA | NA | NA | NA | https://doi.org/10.12688/wellcomeopenres.24569.1 | Included | NA |
| GCF 000002035. | Ingroup | Cypriniformes | Cyprinoidei | Danionidae | Danio | rerio | NCBI | GCF 000002035.6 | 25 | NA | PRJNA11776 | SAMN06930106 | GRCz11 | NCBI Danio rerio Annotation Release 106 | 1373454788 | Chromosome | 09/05/2017 | NA | Genome Reference Consortium | NA | NA | NA | NA | NA | Not included | Replaced with GCA 944039275.1 Danio rerio |
| GCA 903798205. | Ingroup | Cypriniformes | Cyprinoidei | Danionidae | Danio | tinwini | NCBI | GCA 903798205.1 | | NA | PRJEB38571 | SAMEA104026413 | fDanTin1.1 | NA | 1497358328 | Scaffold | 16/06/2020 | CAIGOZ01 | SC | NA | NA | NA | NA | https://doi.org/10.12688/wellcomeopenres.25116.1 | Included | NA |
| GCA 007224835. | Ingroup | Cypriniformes | Cyprinoidei | Danionidae | Danionella | cerebrum | NCBI | GCA 007224835.1 | | Illumina HiSeq; Oxford Nanopore MinION | PRJNA445947 | SAMN11288624 | ASM722483v1 | Annotation submitted by Charite - Universitatsmedizin Berlin | 735303417 | Scaffold | 22/07/2019 | SRMA01 | Charite - Universitatsmedizin Berlin | NA | NA | NA | NA | https://doi.org/10.1038/s41597-019-0161-z | Not included | Replaced with GCA 903798025.1 Danionella cerebrum |
| GCA 903798025. | Ingroup | Cypriniformes | Cyprinoidei | Danionidae | Danionella | cerebrum | NCBI | GCA 903798025.1 | | NA | PRJEB38563 | SAMEA104129915 | fDanTra1.1 | NA | 657501202 | Scaffold | 16/06/2020 | CAIGOI01 | SC | NA | NA | NA | NA | NA | Included | NA |
| GCA 900490495. | Ingroup | Cypriniformes | Cyprinoidei | Danionidae | Danionella | dracula | NCBI | GCA 900490495.1 | | NA | PRJEB27320 | SAMEA104026433 | fDanDra1.1 | NA | 665208374 | Scaffold | 01/07/2018 | UELW01 | SC | NA | NA | NA | NA | https://doi.org/10.12688/wellcomeopenres.21117.1 | Included | NA |
| fDevDev2 | Ingroup | Cypriniformes | Cyprinoidei | Danionidae | Devario | devario | NCBI-SRA | | | NA | NA | NA | NA | NA | NA | NA | NA | NA | NA | PRJEB19273 | SAMEA10413245 | ERS358363 | ERR3332326;ERR333233 | NA | Included | NA |
| fDevDev3 | Ingroup | Cypriniformes | Cyprinoidei | Danionidae | Devario | devario | NCBI-SRA | | | NA | NA | NA | NA | NA | NA | NA | NA | NA | NA | PRJEB19273 | SAMEA10413245 | ERS358363 | ERR3332371;ERR333237 | NA | Not included | Replaced with fDevDev2 Devario devario |
| fDanAeq2 | Ingroup | Cypriniformes | Cyprinoidei | Danionidae | Devario | sp | NCBI-SRA | | | NA | NA | NA | NA | NA | NA | NA | NA | NA | NA | PRJEB19273 | SAMEA10413246 | ERS358367 | ERR3332370;ERR333237 | NA | Included | NA |
| fHorAtu1 | Ingroup | Cypriniformes | Cyprinoidei | Danionidae | Horadandia | atukorali | NCBI-SRA | | | NA | NA | NA | NA | NA | NA | NA | NA | NA | NA | PRJEB19273 | SAMEA10413024 | ERS358366 | ERR3332350;ERR333235 | NA | Included | NA |
| fLauMye1 | Ingroup | Cypriniformes | Cyprinoidei | Danionidae | Laubuka | siamensis | NCBI-SRA | | | NA | NA | NA | NA | NA | NA | NA | NA | NA | NA | PRJEB19273 | SAMEA10413024 | ERS358366 | ERR3332320;ERR333232 | NA | Included | NA |
| fLucSpp1 | Ingroup | Cypriniformes | Cyprinoidei | Danionidae | Luciosoma | sp1 | NCBI-SRA | | | NA | NA | NA | NA | NA | NA | NA | NA | NA | NA | PRJEB19273 | SAMEA10413023 | ERS358363 | ERR3332351;ERR333235 | NA | Included | NA |
| fRaiGut1 | Ingroup | Cypriniformes | Cyprinoidei | Danionidae | Raiamas | guttatus | NCBI-SRA | | | NA | NA | NA | NA | NA | NA | NA | NA | NA | NA | PRJEB19273 | SAMEA10413024 | ERS358366 | ERR3332360;ERR333236 | NA | Included | NA |
| fRasDan1 | Ingroup | Cypriniformes | Cyprinoidei | Danionidae | Rasbora | daniconius | NCBI-SRA | | | NA | NA | NA | NA | NA | NA | NA | NA | NA | NA | PRJEB19273 | SAMEA10413023 | ERS358368 | ERR3332361;ERR333236 | NA | Included | NA |
| fRasKal1 | Ingroup | Cypriniformes | Cyprinoidei | Danionidae | Rasbora | kalochroma | NCBI-SRA | | | NA | NA | NA | NA | NA | NA | NA | NA | NA | NA | PRJEB19273 | SAMEA10413024 | ERS358366 | ERR3332342;ERR333234 | NA | Not included | Replaced with LR15877 Rasbora kalochroma |
| fRasRas1 | Ingroup | Cypriniformes | Cyprinoidei | Danionidae | Rasbora | rasbora | NCBI-SRA | | | NA | NA | NA | NA | NA | NA | NA | NA | NA | NA | PRJEB19273 | SAMEA10413023 | ERS358362 | ERR3332343;ERR333234 | NA | Included | NA |
| GCA 949357685. | Ingroup | Cypriniformes | Cyprinoidei | Gobionidae | Gobio | gobio | NCBI | GCA 949357685.1 | 25 | PacBio,Arima2 | PRJEB60324 | SAMEA11296538 | fGobGob1.1 | NA | 1460662624 | Chromosome | 08/04/2023 | CASHTD01 | WELLCOME SANGER INSTITUTE | NA | NA | NA | NA | NA | Included | NA |
| GCA 023029165. | Ingroup | Cypriniformes | Cyprinoidei | Gobionidae | Gobiocypris | rarus | NCBI | GCA 023029165.1 | 27 | Oxford Nanopore PromethION | PRJNA669452 | SAMN16453471 | ASM2302916v1 | NA | 1134099513 | Chromosome | 18/04/2022 | JAFJHK01 | Center for Ecological and Environmental Sciences | NA | NA | NA | NA | https://doi.org/10.1126/sciadv.abl7253 | Included | NA |
| GCA 018749465. | Ingroup | Cypriniformes | Cyprinoidei | Gobionidae | Paracanthobrama | guichenoti | NCBI | GCA 018749465.1 | 25 | Oxford Nanopore; Illumina HiSeq | PRJNA686690 | SAMN17120673 | ASM1874946v1 | NA | 1087742959 | Chromosome | 08/06/2021 | JAGHMJ01 | Chinese Academy of Fishery Sciences | NA | NA | NA | NA | https://doi.org/10.1038/s41588-021-00933-9 | Included | NA |
| GCA 024679245. | Ingroup | Cypriniformes | Cyprinoidei | Gobionidae | Pseudorasbora | parva | NCBI | GCA 024679245.1 | 25 | PacBio Sequel; Illumina NovaSeq | PRJNA843929 | SAMN28772710 | ASM2467924v1 | NA | 1228852188 | Chromosome | 16/08/2022 | JANLNJ01 | The Max Planck Institute of Molecular Cell Biology and Genetics | NA | NA | NA | NA | NA | Included | NA |
| GCA 023566225. | Ingroup | Cypriniformes | Cyprinoidei | Gobionidae | Romanogobio | albipinnatus | NCBI | GCA 023566225.1 | 25 | PacBio Sequel; Illumina NovaSeq | PRJNA799923 | SAMN25220526 | ASM2356622v1 | NA | 1428567521 | Chromosome | 31/05/2022 | JAKRKY01 | The Max Planck Institute of Molecular Cell Biology and Genetics | NA | NA | NA | NA | NA | Included | NA |
| GCA 963993115. | Ingroup | Cypriniformes | Cyprinoidei | Leuciscidae | Abramis | brama | NCBI | GCA 963993115.1 | 25 | PacBio,Arima2 | PRJEB73975 | SAMEA11296536 | fAbrBra2.1 | NA | 1108201685 | Chromosome | 16/03/2024 | CAXDWV01 | WELLCOME SANGER INSTITUTE | NA | NA | NA | NA | NA | Included | NA |
| GCA 027497735. | Ingroup | Cypriniformes | Cyprinoidei | Leuciscidae | Cyprinella | lepida | NCBI | GCA 027497735.1 | | Illumina | PRJNA806520 | SAMN25889353 | ASM2749773v1 | NA | 768165911 | Scaffold | 30/12/2022 | JAOTRB01 | Iridian Genomes | NA | NA | NA | NA | NA | Included | NA |
| GCA 027497675. | Ingroup | Cypriniformes | Cyprinoidei | Leuciscidae | Cyprinella | lutrensis | NCBI | GCA 027497675.1 | | Illumina | PRJNA806518 | SAMN25889351 | ASM2749767v1 | NA | 748193997 | Scaffold | 30/12/2022 | JAOTRC01 | Iridian Genomes | NA | NA | NA | NA | NA | Included | NA |
| GCA 027361055. | Ingroup | Cypriniformes | Cyprinoidei | Leuciscidae | Cyprinella | venusta | NCBI | GCA 027361055.1 | | Illumina | PRJNA806519 | SAMN25889352 | ASM2736105v1 | NA | 805420949 | Scaffold | 27/12/2022 | JAOTRG01 | Iridian Genomes | NA | NA | NA | NA | NA | Included | NA |
| GCA 030378365. | Ingroup | Cypriniformes | Cyprinoidei | Leuciscidae | Hybognathus | amarus | NCBI | GCA 030378365.1 | | Oxford Nanopore GridION | PRJNA887477 | SAMN32668620 | UNM UB HamF1 1.0 | NA | 977412142 | Scaffold | 30/06/2023 | JAOIUO01 | University of New Mexico | NA | NA | NA | NA | https://doi.org/10.1093/jhered/esad039 | Included | NA |
| GCA 023718395. | Ingroup | Cypriniformes | Cyprinoidei | Leuciscidae | Leucaspius | delineatus | NCBI | GCA 023718395.1 | 25 | PacBio Sequel; Illumina NovaSeq | PRJNA818486 | SAMN26865216 | ASM2371839v1 | NA | 1153514525 | Chromosome | 22/06/2022 | JALDPO01 | The Max Planck Institute of Molecular Cell Biology and Genetics | NA | NA | NA | NA | NA | Included | NA |
| GCA 021554675. | Ingroup | Cypriniformes | Cyprinoidei | Leuciscidae | Leuciscus | idus | NCBI | GCA 021554675.1 | 25 | PacBio Sequel; Illumina NovaSeq | PRJNA768343 | SAMN22028852 | fLeuIdu1.pri.20210917 | NA | 1104030153 | Chromosome | 23/01/2022 | JAJAVR01 | The Max Planck Institute of Molecular Cell Biology and Genetics | NA | NA | NA | NA | NA | Included | NA |
| GCA 030578275. | Ingroup | Cypriniformes | Cyprinoidei | Leuciscidae | Meda | fulgida | NCBI | GCA 030578275.1 | 25 | PacBio | PRJNA947235 | SAMN33845343 | ASM3057827v1 | NA | 882111708 | Chromosome | 08/08/2023 | JAROOJ01 | UC Berkeley | NA | NA | NA | NA | https://doi.org/10.1093/g3journal/jkad157 | Included | NA |
| GCA 949152265. | Ingroup | Cypriniformes | Cyprinoidei | Leuciscidae | Phoxinus | phoxinus | NCBI | GCA 949152265.1 | 25 | PacBio,Arima2 | PRJEB59820 | SAMEA11296539 | fPhoPho1.1 | NA | 950479694 | Chromosome | 08/04/2023 | CASCJP01 | WELLCOME SANGER INSTITUTE | NA | NA | NA | NA | NA | Included | NA |
| GCF 016745375. | Ingroup | Cypriniformes | Cyprinoidei | Leuciscidae | Pimephales | promelas | NCBI | GCF 016745375.1 | | PacBio RSII; Illumina HiSeq | PRJNA565199 | SAMN12875914 | EPA FHM 2.0 | NCBI Pimephales promelas Annotation Release 100 | 1066412313 | Scaffold | 24/01/2021 | WIOS01 | US EPA | NA | NA | NA | NA | https://doi.org/10.1002/etc.5266 | Included | NA |
| GCA 951802725. | Ingroup | Cypriniformes | Cyprinoidei | Leuciscidae | Rutilus | rutilus | NCBI | GCA 951802725.1 | 25 | PacBio,Arima2 | PRJEB62672 | SAMEA11296543 | fRutRut2.1 | NA | 1100179043 | Chromosome | 30/05/2023 | CATOOP01 | WELLCOME SANGER INSTITUTE | NA | NA | NA | NA | NA | Included | NA |
| GCA 024453875. | Ingroup | Cypriniformes | Cyprinoidei | Leuciscidae | Scardinius | erythrophthalmus | NCBI | GCA 024453875.1 | 25 | PacBio Sequel; Illumina NovaSeq | PRJNA841808 | SAMN28618522 | ASM2445387v1 | NA | 1135838337 | Chromosome | 27/07/2022 | JAMKMT01 | The Max Planck Institute of Molecular Cell Biology and Genetics | NA | NA | NA | NA | NA | Included | NA |
| GCA 031834385. | Ingroup | Cypriniformes | Cyprinoidei | Leuciscidae | Semotilus | atromaculatus | NCBI | GCA 031834385.1 | | PacBio Sequel | PRJNA994924 | SAMN36452606 | ASM3183438v1 | NA | 1099257308 | Contig | 21/09/2023 | JAULUF01 | University of Guelph | NA | NA | NA | NA | https://doi.org/10.1093/g3journal/jkad283 | Included | NA |
| GCA 949319135. | Ingroup | Cypriniformes | Cyprinoidei | Leuciscidae | Squalius | cephalus | NCBI | GCA 949319135.1 | 25 | PacBio,Arima2 | PRJEB60239 | SAMEA11296547 | fSquCep2.1 | NA | 1101913912 | Chromosome | 08/04/2023 | CASGGZ01 | WELLCOME SANGER INSTITUTE | NA | NA | NA | NA | NA | Included | NA |
| GCA 030578255. | Ingroup | Cypriniformes | Cyprinoidei | Leuciscidae | Tiaroga | cobitis | NCBI | GCA 030578255.1 | 25 | PacBio Sequel | PRJNA947235 | SAMN33845349 | ASM3057825v1 | NA | 1320521697 | Chromosome | 08/08/2023 | JAROOI01 | UC Berkeley | NA | NA | NA | NA | https://doi.org/10.1093/g3journal/jkad157 | Included | NA |
| GCA 022828995. | Ingroup | Cypriniformes | Cyprinoidei | Leuciscidae | Vimba | vimba | NCBI | GCA 022828995.1 | 25 | PacBio Sequel; Illumina NovaSeq | PRJNA798300 | SAMN25064631 | ASM2282899v1 | NA | 1054310786 | Chromosome | 08/04/2022 | JAKKIH01 | The Max Planck Institute of Molecular Cell Biology and Genetics | NA | NA | NA | NA | NA | Included | NA |
| GCA 028128575. | Ingroup | Cypriniformes | Cyprinoidei | Paedocyprididae | Paedocypris | carbunculus | NCBI | GCA 028128575.1 | | Illumina | PRJNA474779 | SAMN11175355 | ASM2812857v1 | NA | 407240804 | Scaffold | 27/01/2023 | SOMO01 | Universiti Sains Malaysia | NA | NA | NA | NA | https://doi.org/10.3389/fevo.2021.662501 | Not included | Replaced with LR15863 Paedocypris sp. |
| LR12004 | Ingroup | Cypriniformes | Cyprinoidei | Paedocyprididae | Paedocypris | carbunculus | NCBI-SRA | | | Illumina | NA | NA | NA | NA | NA | NA | NA | NA | NA | PRJEB15524 | SAMEA4467423 | ERS136660 | ERR1662935;ERR166293 | https://doi.org/10.1093/gbe/evv058 | Included | NA |
| GCA 028031935. | Ingroup | Cypriniformes | Cyprinoidei | Paedocyprididae | Paedocypris | micromegethes1 | NCBI | GCA 028031935.1 | 15 | Illumina | PRJNA474779 | SAMN09373892 | ASM2803193v1 | NA | 421346737 | Chromosome | 31/01/2023 | OKNB01 | Universiti Sains Malaysia | NA | NA | NA | NA | https://doi.org/10.3389/fevo.2021.662501 | Included | NA |
| LR7898 | Ingroup | Cypriniformes | Cyprinoidei | Paedocyprididae | Paedocypris | micromegethes2 | NCBI-SRA | | | Illumina | NA | NA | NA | NA | NA | NA | NA | NA | NA | PRJEB15524 | SAMEA4467424 | ERS136660 | ERR1662937;ERR166293 | https://doi.org/10.1093/gbe/evv058 | Included | NA |
| GCA 017309765. | Ingroup | Cypriniformes | Cyprinoidei | Paedocyprididae | Paedocypris | sp.Pulau-Singkep | NCBI | GCA 017309765.1 | | Illumina HiSeq | PRJNA636913 | SAMN15089894 | IMCB Psing M 1.0 | NA | 402724664 | Scaffold | 02/03/2021 | JABUMV01 | Institute of Molecular and Cell Biology, A*STAR, Singapore | NA | NA | NA | NA | https://doi.org/10.1126/sciadv.abd8180 | Not included | Replaced with GCA 028031935.1 Paedocypris sp. |
| GCA 017309775. | Ingroup | Cypriniformes | Cyprinoidei | Paedocyprididae | Paedocypris | sp.Pulau-Singkep | NCBI | GCA 017309775.1 | | Illumina HiSeq | PRJNA636914 | SAMN15089893 | IMCB Psing F 1.0 | NA | 403662444 | Scaffold | 02/03/2021 | JABUMW01 | Institute of Molecular and Cell Biology, A*STAR, Singapore | NA | NA | NA | NA | https://doi.org/10.1126/sciadv.abd8180 | Not included | Replaced with GCA 028031935.1 Paedocypris sp. |
| fSunRet1 | Ingroup | Cypriniformes | Cyprinoidei | Sundadanionidae | Sundadanio | retiarius | NCBI-SRA | | | NA | NA | NA | NA | NA | NA | NA | NA | NA | NA | PRJEB19273 | SAMEA10402637 | ERS358367 | ERR3332345;ERR333234 | NA | Not included | Replaced with LR12751 Sundadanio retiarius |
| GCA 003731715. | Ingroup | Cypriniformes | Cyprinoidei | Xenocyprididae | Anabarilius | grahami | NCBI | GCA 003731715.1 | | Illumina HiSeq | PRJNA477399 | SAMN09469597 | BGI Agra 1.0 | Annotation submitted by BGI | 991887266 | Scaffold | 15/11/2018 | RJVU01 | BGI | NA | NA | NA | NA | https://doi.org/10.3389/fgene.2018.00614 | Included | NA |
| GCA 024489055. | Ingroup | Cypriniformes | Cyprinoidei | Xenocyprididae | Chanodichthys | erythropterus | NCBI | GCA 024489055.1 | 24 | Oxford Nanopore PromethION | PRJNA827856 | SAMN27624467 | ASM2448905v1 | NA | 1085510300 | Chromosome | 29/07/2022 | JALPSW01 | Oufu Normal University | NA | NA | NA | NA | https://doi.org/10.1038/s41597-022-01648-0 | Included | NA |
| GCF 019924925. | Ingroup | Cypriniformes | Cyprinoidei | Xenocyprididae | Ctenopharyngodon | idella | NCBI | GCF 019924925.1 | 24 | PacBio Sequel | PRJNA745929 | SAMN20170273 | HZGC01 | GCF 019924925.1-RS 2023 02 | 893198495 | Chromosome | 14/09/2021 | JAIKLG01 | HuaZhong Agricultural University | NA | NA | NA | NA | https://doi.org/10.1186/s12864-022-08503-x | Included | NA |
| GCA 028476615. | Ingroup | Cypriniformes | Cyprinoidei | Xenocyprididae | Culter | alburnus | NCBI | GCA 028476615.1 | | Illumina HiSeq; PacBio | PRJNA700093 | SAMN31113473 | ASM2847661v1 | NA | 1055142460 | Scaffold | 06/02/2023 | JAOTNS01 | Zhejiang Institute of Freshwater Fisheries | NA | NA | NA | NA | https://doi.org/10.3389/fgene.2023.1072506 | Included | NA |
| GCA 037101425. | Ingroup | Cypriniformes | Cyprinoidei | Xenocyprididae | Elopichthys | bambusa | NCBI | GCA 037101425.1 | 24 | Illumina NovaSeq | PRJNA1035000 | SAMN38080342 | HZAU Ebam v1 | NA | 827696973 | Chromosome | 19/03/2024 | JAZKXJ01 | Huazhong agricultural university | NA | NA | NA | NA | https://doi.org/10.1038/s41597-024-03262-8 | Included | NA |
| GCA 004764525. | Ingroup | Cypriniformes | Cyprinoidei | Xenocyprididae | Hypophthalmichthys | molitrix | NCBI | GCA 004764525.1 | | Illumina HiSeq | PRJNA305140 | SAMN04317213 | HypMol1.0 | NA | 1104676189 | Scaffold | 09/04/2019 | LOHV01 | University of Nebraska at Omaha | NA | NA | NA | NA | https://doi.org/10.1111/eva.12863 | Included | NA |
| GCA 004193235. | Ingroup | Cypriniformes | Cyprinoidei | Xenocyprididae | Hypophthalmichthys | nobilis | NCBI | GCA 004193235.1 | | Illumina HiSeq | PRJNA305141 | SAMN04317176 | HypNob1.0 | NA | 1012063666 | Scaffold | 12/02/2019 | LOHW01 | University of Nebraska at Omaha | NA | NA | NA | NA | https://doi.org/10.1111/eva.12863 | Not included | Replaced with GCA 019925145.1 Hypophthalmichthys nobilis |
| GCA 019925145. | Ingroup | Cypriniformes | Cyprinoidei | Xenocyprididae | Hypophthalmichthys | nobilis | NCBI | GCA 019925145.1 | 24 | Illumina HiSeq; Oxford Nanopore PromethION | PRJNA657698 | SAMN15835045 | ASM1992514v1 | NA | 857045951 | Chromosome | 14/09/2021 | JACTVH01 | Chinese Academy of Sciences | NA | NA | NA | NA | https://doi.org/10.3389/fgene.2021.728177 | Included | NA |
| GCF 018812025. | Ingroup | Cypriniformes | Cyprinoidei | Xenocyprididae | Megalobrama | amblycephala | NCBI | GCF 018812025.1 | 24 | PacBio RSII | PRJNA343584 | SAMN18522313 | ASM1881202v1 | NCBI Megalobrama amblycephala Annotation Release 100 | 1109871993 | Chromosome | 09/06/2021 | JAGTWI01 | Huazhong Agricultural University | NA | NA | NA | NA | https://doi.org/10.1093/molbev/msab152 | Included | NA |
| GCA 034642465. | Ingroup | Cypriniformes | Cyprinoidei | Xenocyprididae | Zacco | platypus | NCBI | GCA 034642465.1 | 24 | PacBio CLR | PRJNA1028840 | SAMN37853698 | ASM3464246v1 | NA | 815003113 | Chromosome | 27/12/2023 | JAYDZZ01 | Zhejiang Academy of Agricultural Sciences | NA | NA | NA | NA | https://doi.org/10.1038/s41597-024-03163-w | Included | NA |
| GCA 034509495. | Outgroup | Characiformes | | Acestrorhynchidae | Acestrorhynchus | altus | NCBI | GCA 034509495.1 | | | PRJNA1004608 | SAMN36947357 | ASM3450949v1 | NA | 950195876 | Chromosome | 26/12/2023 | JAVGWO01 | BGI Research | | | | | https://doi.org/10.1093/dnares/dsad022 | Included | NA |
| GCF 023375975. | Outgroup | Characiformes | | Characidae | Astyanax | mexicanus | NCBI | GCF 023375975.1 | 25 | PacBio Sequel | PRJNA807270 | SAMN25965548 | AstMex3 surface | NCBI Astyanax mexicanus Annotation Release 103 | 1373169186 | Chromosome | 19/05/2022 | JALAHU01 | University of Missouri | | | | | https://doi.org/10.1038/s41467-021-21733-z | Included | NA |
| GCA 032718565. | Outgroup | Characiformes | | Distichodontidae | Distichodus | sexfasciatus | NCBI | GCA 032718565.1 | | Oxford Nanopore GridION | PRJNA1004608 | SAMN36947359 | ASM3271856v1 | NA | 812893345 | Scaffold | 16/10/2023 | JAVGVS01 | BGI Research | | | | | https://doi.org/10.1093/dnares/dsad022 | Included | NA |
| GCA 017165825. | Outgroup | Characiformes | | Hepsetidae | Hepsetus | odoe | NCBI | GCA 017165825.1 | 29 | BGI SEQ 500 | PRJNA625402 | SAMN14603884 | ASM1716582v1 | NA | 862056194 | Chromosome | 09/03/2021 | JABMDN01 | BGI Research | | | | | https://doi.org/10.1093/dnares/dsad022 | Included | NA |
| GCA 034552555. | Outgroup | Characiformes | | Prochilodontidae | Semaprochilodus | insignis | NCBI | GCA 034552555.1 | 27 | Complete Genomics | PRJNA1004608 | SAMN36947358 | ASM3455255v1 | NA | 1178126082 | Chromosome | 26/12/2023 | JAVGWN01 | BGI Research | | | | | https://doi.org/10.1093/dnares/dsad022 | Included | NA |
| GCF 900700375. | Outgroup | Clupeiformes | | Denticipitidae | Denticeps | clupeoides | NCBI | GCF 900700375.1 | 20 | PacBio,Illumina,BioNano,Arima | PRJEB31545 | SAMEA104129917 | fDenClu1.1 | NCBI Denticeps clupeoides Annotation Release 100 | 567401054 | Chromosome | 16/03/2019 | CAADHS01 | Wellcome Sanger Institute | | | | | NA | Included | NA |
| GCA 027475355. | Outgroup | Clupeiformes | | Engraulidae | Coilia | nasus | NCBI | GCA 027475355.1 | 24 | PacBio | PRJNA895906 | SAMN31526683 | ASM2747535v1 | NA | 851665587 | Chromosome | 28/12/2022 | JAPTFL01 | Nanjing Agricultural University | | | | | https://doi.org/10.1038/s41597-023-02278-w | Included | NA |
| GCF 902362185. | Outgroup | Gonorynchiformes | | Chanidae | Chanos | chanos | NCBI | GCF 902362185.1 | 16 | PacBio,Illumina,BioNano,Dovetail | PRJEB33882 | SAMEA5159473 | fChaCha1.1 | NCBI Chanos chanos Annotation Release 100 | 656912816 | Chromosome | 14/08/2019 | CABIVN01 | Wellcome Sanger Institute | | | | | NA | Included | NA |
| GCA 028565775. | Outgroup | Gymnotiformes | | Apteronotidae | Sternarchorhynchus | goeldii | NCBI | GCA 028565775.1 | | Illumina | PRJNA547755 | SAMN11975474 | ASM2856577v1 | NA | 527534136 | Scaffold | 09/02/2023 | JAOXYF01 | IRIDIAN GENOMES | | | | | NA | Included | NA |
| GCA 025802075. | Outgroup | Gymnotiformes | | Apteronotidae | Tenebrosternarchus | preto | NCBI | GCA 025802075.1 | | Illumina | PRJNA547788 | SAMN11976056 | ASM2580207v1 | NA | 536308962 | Scaffold | 23/10/2022 | JAOYMR01 | IRIDIAN GENOMES | | | | | NA | Included | NA |
| GCA 020368025. | Outgroup | Gymnotiformes | | Hypopomidae | Brachyhypopomus | occidentalis | NCBI | GCA 020368025.1 | | Oxford Nanopore MinION; Illumina HiSeq | PRJNA727273 | SAMN19011640 | ASM2036802v1 | NA | 540332535 | Scaffold | 07/10/2021 | JAGXOD01 | Smithsonian Tropical Research Institute | | | | | https://doi.org/10.1093/gbe/evab223 | Included | NA |
| GCF 030014155. | Outgroup | Siluriformes | | Bagridae | Tachysurus | vachellii | NCBI | GCF 030014155.1 | | PacBio Sequel | PRJNA819563 | SAMN33769291 | HZAU Pvac v1 | GCF 030014155.1-RS 2023 11 | 692104899 | Chromosome | 16/05/2023 | JASFAO01 | Huazhong Agricultural University | | | | | https://doi.org/10.1038/s41597-023-02509-0 | Included | NA |
| GCA 030522415. | Outgroup | Siluriformes | | Clariidae | Channallabes | apus | NCBI | GCA 030522415.1 | 28 | PacBio Sequel; Illumina NovaSeq | PRJNA834615 | SAMN28052917 | ASM3052241v1 | NA | 1164854870 | Chromosome | 26/07/2023 | JAMASW01 | The Max Planck Institute of Molecular Cell Biology and Genetics | | | | | https://doi.org/10.1186/s12915-023-01661-8 | Included | NA |
| GCA 030347435. | Outgroup | Siluriformes | | Clariidae | Clarias | fuscus | NCBI | GCA 030347435.1 | 28 | PacBio | PRJNA822706 | SAMN27277226 | ASM3034743v1 | NA | 944286092 | Chromosome | 27/06/2023 | JARWBY01 | Guangdong Ocean University | | | | | https://doi.org/10.1186/s12864-023-09394-2 | Included | NA |
| GCA 023630585. | Outgroup | Siluriformes | | Cranoglanididae | Cranoglanis | bouderius | NCBI | GCA 023630585.1 | | Oxford Nanopore PromethION; PacBio RSII | PRJNA720277 | SAMN28232390 | ASM2363058v1 | NA | 881346651 | Chromosome | 02/06/2022 | JAMETE01 | Southwest University | | | | | https://doi.org/10.3389/fgene.2022.962406 | Included | NA |
| GCF 023375685. | Outgroup | Siluriformes | | Ictaluridae | Ictalurus | furcatus | NCBI | GCF 023375685.1 | 29 | PacBio CLR; Illumina | PRJNA834118 | SAMN28041099 | Billie 1.0 | GCF 023375685.1-RS 2023 03 | 825459183 | Chromosome | 02/06/2022 | JAMCCP01 | USDA, Agricultural Research Service | | | | | https://doi.org/10.1186/s12915-023-01556-8 | Included | NA |
| GCA 025084515. | Outgroup | Siluriformes | | Loricariidae | Ancistrus | triradiatus | NCBI | GCA 025084515.1 | | Illumina HiSeq; PacBio | PRJNA769304 | SAMN22109509 | ASM2508451v1 | NA | 994062695 | Scaffold | 08/09/2022 | JAJAGP01 | University of Geneva | | | | | https://doi.org/10.1007/s00438-022-01947-6 | Included | NA |
| GCA 024760905. | Outgroup | Siluriformes | | Plotosidae | Plotosus | lineatus | NCBI | GCA 024760905.1 | 24 | Oxford Nanopore GridION; PacBio Sequel | PRJNA720277 | SAMN27514949 | SWU Pli 1.0 | NA | 1333861300 | Chromosome | 29/08/2022 | JALRMN01 | Southwest University | | | | | https://doi.org/10.24272/j.issn.2095-8137.2022.387 | Included | NA |
| GCF 014805685. | Outgroup | Siluriformes | | Siluridae | Silurus | meridionalis | NCBI | GCF 014805685.1 | 29 | Oxford Nanopore PromethION | PRJNA623722 | SAMN14558653 | ASM1480568v1 | NCBI Silurus meridionalis Annotation Release 100 | 742579920 | Chromosome | 02/10/2020 | JABFDY01 | Southwest University | | | | | https://doi.org/10.1111/1755-0998.13338 | Included | NA |

### **Supplementary Table 3.** Phylogenomic dataset of 335 species included in this study.

| Voucher reads | Group | Order | Suborder | Family | Genus | Species | A Single complete pct | Single complete genes | B Duplicate complete pct | Duplicate complete genes | Fragmented subclass1 pct | Fragmented subclass1 genes | I Fragmented subclass2 pct | I Fragmented subclass2 genes | D Missing pct | Missing genes | total complete genes | total fragmented genes | C total fragmented pct | genome completeness | Source | Assembly | BUSCO.loci.dataset.320taxa | Hughes.loci.dataset.330taxa | Malmstrøm.loci.dataset.335taxa | UCE.loci.dataset.335.taxa | BUSCO.NT.C12+UCE.loci.dat | ROADIES.loci.dataset.335taxa |
|---|---|---|---|---|---|---|---|---|---|---|---|---|---|---|---|---|---|---|---|---|---|---|---|---|---|---|---|---|
| LR15244 | Ingroup | Cypriniformes | Catostomoidei | Catostomidae | Catostomus | commersoni | 31.32 | 1140 | 6.73 | 245 | 28.02 | 1020 | 0.36 | 13 | 33.57 | 1222 | 1385 | 1033 | 28.38 | C:1385[S:1140,D:245],F:1033,M:1222,n:3640 | This study | abyss | + | + | + | + | + | + |
| LR15247 | Ingroup | Cypriniformes | Catostomoidei | Catostomidae | Cycleptus | sp | 40.03 | 1457 | 10.99 | 400 | 25.66 | 934 | 0.41 | 15 | 22.91 | 834 | 1857 | 949 | 26.07 | C:1857[S:1457,D:400],F:949,M:834,n:3640 | This study | abyss | + | + | + | + | + | + |
| LR15245 | Ingroup | Cypriniformes | Catostomoidei | Catostomidae | Erimyzon | sucetta | 54.29 | 1976 | 14.81 | 539 | 16.84 | 613 | 0.16 | 6 | 13.9 | 506 | 2515 | 619 | 17 | C:2515[S:1976,D:539],F:619,M:506,n:3640 | This study | abyss | + | + | + | + | + | + |
| LR15246 | Ingroup | Cypriniformes | Catostomoidei | Catostomidae | Ictiobus | bubalus | 19.4 | 706 | 3.24 | 118 | 29.09 | 1059 | 0.44 | 16 | 47.83 | 1741 | 824 | 1075 | 29.53 | C:824[S:706,D:118],F:1075,M:1741,n:3640 | This study | spades | + | + | + | + | + | + |
| LR15248 | Ingroup | Cypriniformes | Catostomoidei | Catostomidae | Moxostoma | congestum | 50.63 | 1843 | 15.6 | 568 | 20.05 | 730 | 0.3 | 11 | 13.41 | 488 | 2411 | 741 | 20.35 | C:2411[S:1843,D:568],F:741,M:488,n:3640 | This study | spades | + | + | + | + | + | + |
| GCF_019703515.2 | Ingroup | Cypriniformes | Catostomoidei | Catostomidae | Myxocyprinus | asiaticus | 42.58 | 1550 | 57.01 | 2075 | 0.27 | 10 | 0 | 0 | 0.14 | 5 | 3625 | 10 | 0.27 | C:3625[S:1550,D:2075],F:10,M:5,n:3640 | NCBI | multiple assembly methods | + | + | + | + | + | + |
| LR15354 | Ingroup | Cypriniformes | Cobitoidei | Balitoridae | Balitora | sp | 92.45 | 3365 | 1.29 | 47 | 3.76 | 137 | 0.03 | 1 | 2.47 | 90 | 3412 | 138 | 3.79 | C:3412[S:3365,D:47],F:138,M:90,n:3640 | This study | abyss | + | + | + | + | + | + |
| LR15094 | Ingroup | Cypriniformes | Cobitoidei | Balitoridae | Balitoropsis | zollingeri | 86.7 | 3156 | 3.87 | 141 | 5.3 | 193 | 0.14 | 5 | 3.98 | 145 | 3297 | 198 | 5.44 | C:3297[S:3156,D:141],F:198,M:145,n:3640 | This study | abyss | + | + | + | + | + | + |
| KUFOS.AN.2019.1.1 | Ingroup | Cypriniformes | Cobitoidei | Balitoridae | Bhavania | australis | 85.66 | 3118 | 0.74 | 27 | 8.32 | 303 | 0.11 | 4 | 5.16 | 188 | 3145 | 307 | 8.43 | C:3145[S:3118,D:27],F:307,M:188,n:3640 | This study | abyss | - | + | + | + | + | + |
| LR15176 | Ingroup | Cypriniformes | Cobitoidei | Balitoridae | Hemimyzon | formosanum | 89.48 | 3257 | 1.37 | 50 | 5.69 | 207 | 0 | 0 | 3.46 | 126 | 3307 | 207 | 5.69 | C:3307[S:3257,D:50],F:207,M:126,n:3640 | This study | abyss | + | + | + | + | + | + |
| LR15139 | Ingroup | Cypriniformes | Cobitoidei | Balitoridae | Homaloptera | sp | 87.17 | 3173 | 1.24 | 45 | 6.51 | 237 | 0.03 | 1 | 5.05 | 184 | 3218 | 238 | 6.54 | C:3218[S:3173,D:45],F:238,M:184,n:3640 | This study | abyss | + | + | + | + | + | + |
| LR15140 | Ingroup | Cypriniformes | Cobitoidei | Balitoridae | Homalopteroides | stephensoni | 91.29 | 3323 | 1.04 | 38 | 4.53 | 165 | 0 | 0 | 3.13 | 114 | 3361 | 165 | 4.53 | C:3361[S:3323,D:38],F:165,M:114,n:3640 | This study | abyss | + | + | + | + | + | + |
| LR15093 | Ingroup | Cypriniformes | Cobitoidei | Balitoridae | Homalopteroides | tweediei | 87.53 | 3186 | 0.44 | 16 | 7.12 | 259 | 0.11 | 4 | 4.81 | 175 | 3202 | 263 | 7.23 | C:3202[S:3186,D:16],F:263,M:175,n:3640 | This study | spades | + | + | + | + | + | + |
| LR15091 | Ingroup | Cypriniformes | Cobitoidei | Balitoridae | Homalopterula | gymnogaster | 78.82 | 2869 | 0.8 | 29 | 11.29 | 411 | 0.19 | 7 | 8.9 | 324 | 2898 | 418 | 11.48 | C:2898[S:2869,D:29],F:418,M:324,n:3640 | This study | abyss | + | + | + | + | + | + |
| LR15172 | Ingroup | Cypriniformes | Cobitoidei | Balitoridae | Lepturichthys | fimbriatus | 81.76 | 2976 | 1.84 | 67 | 9.64 | 351 | 0.22 | 8 | 6.54 | 238 | 3043 | 359 | 9.86 | C:3043[S:2976,D:67],F:359,M:238,n:3640 | This study | abyss | + | + | + | + | + | + |
| LR15487 | Ingroup | Cypriniformes | Cobitoidei | Balitoridae | Neohomaloptera | johorensis | 91.57 | 3333 | 1.15 | 42 | 4.23 | 154 | 0.08 | 3 | 2.97 | 108 | 3375 | 157 | 4.31 | C:3375[S:3333,D:42],F:157,M:108,n:3640 | This study | abyss | + | + | + | + | + | + |
| LR15088 | Ingroup | Cypriniformes | Cobitoidei | Balitoridae | Sinogastromyzon | puliensis | 88.9 | 3236 | 0.71 | 26 | 6.54 | 238 | 0.08 | 3 | 3.76 | 137 | 3262 | 241 | 6.62 | C:3262[S:3236,D:26],F:241,M:137,n:3640 | This study | abyss | + | + | + | + | + | + |
| LR15181 | Ingroup | Cypriniformes | Cobitoidei | Balitoridae | Travancoria | elongata | 90.41 | 3291 | 0.52 | 19 | 5.55 | 202 | 0.11 | 4 | 3.41 | 124 | 3310 | 206 | 5.66 | C:3310[S:3291,D:19],F:206,M:124,n:3640 | This study | spades | + | + | + | + | + | + |
| LR15360 | Ingroup | Cypriniformes | Cobitoidei | Barbuccidae | Barbucca | elongatus | 94.01 | 3422 | 0.69 | 25 | 2.77 | 101 | 0 | 0 | 2.53 | 92 | 3447 | 101 | 2.77 | C:3447[S:3422,D:25],F:101,M:92,n:3640 | This study | abyss | + | + | + | + | + | + |
| LR15330 | Ingroup | Cypriniformes | Cobitoidei | Botiidae | Ambastaia | sidthimunki | 62.14 | 2262 | 17.17 | 625 | 10.96 | 399 | 0.16 | 6 | 9.56 | 348 | 2887 | 405 | 11.12 | C:2887[S:2262,D:625],F:405,M:348,n:3640 | This study | abyss | + | + | + | + | + | + |
| GCA_036877575.1 | Ingroup | Cypriniformes | Cobitoidei | Botiidae | Botia | almorhae | 62.53 | 2276 | 20.22 | 736 | 9.64 | 351 | 0.16 | 6 | 7.45 | 271 | 3012 | 357 | 9.8 | C:3012[S:2276,D:736],F:357,M:271,n:3640 | NCBI | multiple assembly methods | + | + | + | + | + | + |
| LR15149 | Ingroup | Cypriniformes | Cobitoidei | Botiidae | Botia | kubotai | 59.42 | 2163 | 22.09 | 804 | 9.53 | 347 | 0.05 | 2 | 8.9 | 324 | 2967 | 349 | 9.58 | C:2967[S:2163,D:804],F:349,M:324,n:3640 | This study | abyss | + | + | + | + | + | + |
| LR15148 | Ingroup | Cypriniformes | Cobitoidei | Botiidae | Chromobotia | macracanthus | 67.8 | 2468 | 21.07 | 767 | 6.84 | 249 | 0.03 | 1 | 4.26 | 155 | 3235 | 250 | 6.87 | C:3235[S:2468,D:767],F:250,M:155,n:3640 | This study | spades | + | + | + | + | + | + |
| LR15074 | Ingroup | Cypriniformes | Cobitoidei | Botiidae | Leptobotia | elongata | 86.13 | 3135 | 0.93 | 34 | 7.69 | 280 | 0.03 | 1 | 5.22 | 190 | 3169 | 281 | 7.72 | C:3169[S:3135,D:34],F:281,M:190,n:3640 | This study | abyss | + | + | + | + | + | + |
| 112LR15712 | Ingroup | Cypriniformes | Cobitoidei | Botiidae | Parabotia | bimaculatus | 86.07 | 3133 | 0.27 | 10 | 7.94 | 289 | 0 | 0 | 5.71 | 208 | 3143 | 289 | 7.94 | C:3143[S:3133,D:10],F:289,M:208,n:3640 | This study | spades | + | + | + | + | + | + |
| LR15151 | Ingroup | Cypriniformes | Cobitoidei | Botiidae | Sinibotia | robusta | 52.45 | 1909 | 21.98 | 800 | 13.21 | 481 | 0.16 | 6 | 12.2 | 444 | 2709 | 487 | 13.37 | C:2709[S:1909,D:800],F:487,M:444,n:3640 | This study | abyss | + | + | + | + | + | + |
| LR15446 | Ingroup | Cypriniformes | Cobitoidei | Botiidae | Syncrossus | berdmorei | 63.41 | 2308 | 18.9 | 688 | 9.31 | 339 | 0.11 | 4 | 8.27 | 301 | 2996 | 343 | 9.42 | C:2996[S:2308,D:688],F:343,M:301,n:3640 | This study | abyss | + | + | + | + | + | + |
| GCA_036877545.1 | Ingroup | Cypriniformes | Cobitoidei | Botiidae | Yasuhikotakia | modesta | 70.58 | 2569 | 15.96 | 581 | 7.64 | 278 | 0.11 | 4 | 5.71 | 208 | 3150 | 282 | 7.75 | C:3150[S:2569,D:581],F:282,M:208,n:3640 | NCBI | multiple assembly methods | + | + | + | + | + | + |
| 115LR15715 | Ingroup | Cypriniformes | Cobitoidei | Cobitidae | Acanthopsoides | robertsi | 82.72 | 3011 | 1.59 | 58 | 9.75 | 355 | 0.14 | 5 | 5.8 | 211 | 3069 | 360 | 9.89 | C:3069[S:3011,D:58],F:360,M:211,n:3640 | This study | abyss | + | + | + | + | + | + |
| LR15340 | Ingroup | Cypriniformes | Cobitoidei | Cobitidae | Acantopsis | sp | 36.15 | 1316 | 0.25 | 9 | 24.37 | 887 | 0.52 | 19 | 38.71 | 1409 | 1325 | 906 | 24.89 | C:1325[S:1316,D:9],F:906,M:1409,n:3640 | This study | spades | + | + | + | + | + | + |
| LR15193 | Ingroup | Cypriniformes | Cobitoidei | Cobitidae | Bibarba | sp | 88.05 | 3205 | 0.38 | 14 | 6.65 | 242 | 0.14 | 5 | 4.78 | 174 | 3219 | 247 | 6.79 | C:3219[S:3205,D:14],F:247,M:174,n:3640 | This study | spades | + | + | + | + | + | + |
| LR15575 | Ingroup | Cypriniformes | Cobitoidei | Cobitidae | Canthophrys | gongota | 87.69 | 3192 | 1.1 | 40 | 6.43 | 234 | 0.05 | 2 | 4.73 | 172 | 3232 | 236 | 6.48 | C:3232[S:3192,D:40],F:236,M:172,n:3640 | This study | abyss | + | + | + | + | + | + |
| LR15465 | Ingroup | Cypriniformes | Cobitoidei | Cobitidae | Cobitis | sp | 27.36 | 996 | 0.14 | 5 | 26.21 | 954 | 0.49 | 18 | 45.8 | 1667 | 1001 | 972 | 26.7 | C:1001[S:996,D:5],F:972,M:1667,n:3640 | This study | abyss | + | + | + | + | + | + |
| LR15170 | Ingroup | Cypriniformes | Cobitoidei | Cobitidae | Kottelatlimia | katik | 62.28 | 2267 | 0.66 | 24 | 19.92 | 725 | 0.22 | 8 | 16.92 | 616 | 2291 | 733 | 20.14 | C:2291[S:2267,D:24],F:733,M:616,n:3640 | This study | abyss | + | + | + | + | + | + |
| 114LR15714 | Ingroup | Cypriniformes | Cobitoidei | Cobitidae | Kottelatlimia | pristes | 56.1 | 2042 | 1.24 | 45 | 21.46 | 781 | 0.38 | 14 | 20.82 | 758 | 2087 | 795 | 21.84 | C:2087[S:2042,D:45],F:795,M:758,n:3640 | This study | abyss | + | + | + | + | + | + |
| LR15467 | Ingroup | Cypriniformes | Cobitoidei | Cobitidae | Lepidocephalichthys | sp | 80.19 | 2919 | 1.46 | 53 | 10.49 | 382 | 0.16 | 6 | 7.69 | 280 | 2972 | 388 | 10.65 | C:2972[S:2919,D:53],F:388,M:280,n:3640 | This study | abyss | + | + | + | + | + | + |
| GCF_027580225.1 | Ingroup | Cypriniformes | Cobitoidei | Cobitidae | Misgurnus | anguillicaudatus | 95.11 | 3462 | 2.12 | 77 | 0.49 | 18 | 0 | 0 | 2.28 | 83 | 3539 | 18 | 0.49 | C:3539[S:3462,D:77],F:18,M:83,n:3640 | NCBI | multiple assembly methods | + | + | + | + | + | + |
| GCA_025389145.1 | Ingroup | Cypriniformes | Cobitoidei | Cobitidae | Misgurnus | mizolepis | 85.82 | 3124 | 12.31 | 448 | 0.47 | 17 | 0 | 0 | 1.4 | 51 | 3572 | 17 | 0.47 | C:3572[S:3124,D:448],F:17,M:51,n:3640 | NCBI | multiple assembly methods | + | + | + | + | + | + |
| BNHS.FWF.981 | Ingroup | Cypriniformes | Cobitoidei | Cobitidae | Pangio | bhujia | 70.44 | 2564 | 0.96 | 35 | 16.18 | 589 | 0.27 | 10 | 12.14 | 442 | 2599 | 599 | 16.45 | C:2599[S:2564,D:35],F:599,M:442,n:3640 | This study | spades | - | + | + | + | + | + |
| LR15141 | Ingroup | Cypriniformes | Cobitoidei | Cobitidae | Pangio | sp | 43.85 | 1596 | 2.94 | 107 | 24.29 | 884 | 0.47 | 17 | 28.46 | 1036 | 1703 | 901 | 24.76 | C:1703[S:1596,D:107],F:901,M:1036,n:3640 | This study | abyss | + | + | + | + | + | + |
| LR15609 | Ingroup | Cypriniformes | Cobitoidei | Cobitidae | Sabanejewia | balcanica | 54.75 | 1993 | 1.1 | 40 | 20.6 | 750 | 0.27 | 10 | 23.27 | 847 | 2033 | 760 | 20.87 | C:2033[S:1993,D:40],F:760,M:847,n:3640 | This study | abyss | + | + | + | + | + | + |
| LR15146 | Ingroup | Cypriniformes | Cobitoidei | Cobitidae | Theriodes | sandakanensis | 90.99 | 3312 | 1.62 | 59 | 4.12 | 150 | 0.03 | 1 | 3.24 | 118 | 3371 | 151 | 4.15 | C:3371[S:3312,D:59],F:151,M:118,n:3640 | This study | abyss | + | + | + | + | + | + |
| 141LR15748 | Ingroup | Cypriniformes | Cobitoidei | Ellopostomatidae | Ellopostoma | mystax | 92.72 | 3375 | 0.69 | 25 | 3.79 | 138 | 0 | 0 | 2.8 | 102 | 3400 | 138 | 3.79 | C:3400[S:3375,D:25],F:138,M:102,n:3640 | This study | abyss | + | + | + | + | + | + |
| LR15180 | Ingroup | Cypriniformes | Cobitoidei | Gastromyzontidae | Annamia | sp | 92.01 | 3349 | 1.32 | 48 | 3.87 | 141 | 0.05 | 2 | 2.75 | 100 | 3397 | 143 | 3.92 | C:3397[S:3349,D:48],F:143,M:100,n:3640 | This study | abyss | + | + | + | + | + | + |
| GCA_019155185.1 | Ingroup | Cypriniformes | Cobitoidei | Gastromyzontidae | Beaufortia | kweichowensis | 98.08 | 3570 | 0.77 | 28 | 0.41 | 15 | 0 | 0 | 0.74 | 27 | 3598 | 15 | 0.41 | C:3598[S:3570,D:28],F:15,M:27,n:3640 | NCBI | multiple assembly methods | + | + | + | + | + | + |
| LR15491 | Ingroup | Cypriniformes | Cobitoidei | Gastromyzontidae | Erromyzon | sp | 79.75 | 2903 | 0.85 | 31 | 10.05 | 366 | 0.19 | 7 | 9.15 | 333 | 2934 | 373 | 10.24 | C:2934[S:2903,D:31],F:373,M:333,n:3640 | This study | abyss | + | + | + | + | + | + |
| LR15090 | Ingroup | Cypriniformes | Cobitoidei | Gastromyzontidae | Glaniopsis | muliradiatus | 85.66 | 3118 | 2.01 | 73 | 7.12 | 259 | 0.11 | 4 | 5.11 | 186 | 3191 | 263 | 7.23 | C:3191[S:3118,D:73],F:263,M:186,n:3640 | This study | abyss | + | + | + | + | + | + |
| LR15133 | Ingroup | Cypriniformes | Cobitoidei | Gastromyzontidae | Katibasia | insidiosa | 90.25 | 3285 | 0.74 | 27 | 5.36 | 195 | 0.05 | 2 | 3.6 | 131 | 3312 | 197 | 5.41 | C:3312[S:3285,D:27],F:197,M:131,n:3640 | This study | abyss | + | + | + | + | + | + |
| LR15175 | Ingroup | Cypriniformes | Cobitoidei | Gastromyzontidae | Liniparhomaloptera | disparis | 89.26 | 3249 | 0.36 | 13 | 5.85 | 213 | 0.05 | 2 | 4.48 | 163 | 3262 | 215 | 5.9 | C:3262[S:3249,D:13],F:215,M:163,n:3640 | This study | abyss | + | + | + | + | + | + |
| LR15476 | Ingroup | Cypriniformes | Cobitoidei | Gastromyzontidae | Plesiomyzon | baotingensis | 90.88 | 3308 | 0.71 | 26 | 4.15 | 151 | 0.08 | 3 | 4.18 | 152 | 3334 | 154 | 4.23 | C:3334[S:3308,D:26],F:154,M:152,n:3640 | This study | abyss | + | + | + | + | + | + |
| LR15425 | Ingroup | Cypriniformes | Cobitoidei | Gastromyzontidae | Protomyzon | borneensis | 84.42 | 3073 | 3.46 | 126 | 7.34 | 267 | 0.05 | 2 | 4.73 | 172 | 3199 | 269 | 7.39 | C:3199[S:3073,D:126],F:269,M:172,n:3640 | This study | abyss | + | + | + | + | + | + |
| LR15067 | Ingroup | Cypriniformes | Cobitoidei | Gastromyzontidae | Pseudogastromyzon | sp | 92.58 | 3370 | 0.52 | 19 | 4.18 | 152 | 0 | 0 | 2.72 | 99 | 3389 | 152 | 4.18 | C:3389[S:3370,D:19],F:152,M:99,n:3640 | This study | abyss | + | + | + | + | + | + |
| LR15561 | Ingroup | Cypriniformes | Cobitoidei | Gastromyzontidae | Sewellia | elongata | 92.31 | 3360 | 1.1 | 40 | 3.87 | 141 | 0 | 0 | 2.72 | 99 | 3400 | 141 | 3.87 | C:3400[S:3360,D:40],F:141,M:99,n:3640 | This study | abyss | + | + | + | + | + | + |
| 113LR15713 | Ingroup | Cypriniformes | Cobitoidei | Gastromyzontidae | Yaoshania | pachychilus | 90.96 | 3311 | 0.47 | 17 | 5.33 | 194 | 0 | 0 | 3.24 | 118 | 3328 | 194 | 5.33 | C:3328[S:3311,D:17],F:194,M:118,n:3640 | This study | abyss | + | + | + | + | + | + |
| LR15343 | Ingroup | Cypriniformes | Cobitoidei | Nemacheilidae | Aborichthys | elongatus | 87.61 | 3189 | 0.52 | 19 | 7.34 | 267 | 0.05 | 2 | 4.48 | 163 | 3208 | 269 | 7.39 | C:3208[S:3189,D:19],F:269,M:163,n:3640 | This study | abyss | + | + | + | + | + | + |
| GCA_947034865.1 | Ingroup | Cypriniformes | Cobitoidei | Nemacheilidae | Barbatula | barbatula | 97.17 | 3537 | 0.6 | 22 | 0.52 | 19 | 0 | 0 | 1.7 | 62 | 3559 | 19 | 0.52 | C:3559[S:3537,D:22],F:19,M:62,n:3640 | NCBI | multiple assembly methods | + | + | + | + | + | + |
| LR15190 | Ingroup | Cypriniformes | Cobitoidei | Nemacheilidae | Barbatula | sp | 85.14 | 3099 | 1.68 | 61 | 7.97 | 290 | 0.11 | 4 | 5.11 | 186 | 3160 | 294 | 8.08 | C:3160[S:3099,D:61],F:294,M:186,n:3640 | This study | abyss | + | + | + | + | + | + |
| LR15073 | Ingroup | Cypriniformes | Cobitoidei | Nemacheilidae | Homatula | variegata | 84.09 | 3061 | 0.66 | 24 | 9.01 | 328 | 0.05 | 2 | 6.18 | 225 | 3085 | 330 | 9.06 | C:3085[S:3061,D:24],F:330,M:225,n:3640 | This study | abyss | + | + | + | + | + | + |
| KUFOS.AN.2019.2.1 | Ingroup | Cypriniformes | Cobitoidei | Nemacheilidae | Indoreonectes | keralensis | 49.48 | 1801 | 0.55 | 20 | 23.19 | 844 | 0.44 | 16 | 26.35 | 959 | 1821 | 860 | 23.63 | C:1821[S:1801,D:20],F:860,M:959,n:3640 | This study | spades | - | + | + | + | + | + |
| LR15462 | Ingroup | Cypriniformes | Cobitoidei | Nemacheilidae | Lefua | sp | 85.52 | 3113 | 1.59 | 58 | 7.42 | 270 | 0.11 | 4 | 5.36 | 195 | 3171 | 274 | 7.53 | C:3171[S:3113,D:58],F:274,M:195,n:3640 | This study | abyss | + | + | + | + | + | + |
| KUFOS.AN.2021.1.1 | Ingroup | Cypriniformes | Cobitoidei | Nemacheilidae | Mesonoemacheilus | triangularis | 64.64 | 2353 | 3.65 | 133 | 16.4 | 597 | 0.19 | 7 | 15.11 | 550 | 2486 | 604 | 16.59 | C:2486[S:2353,D:133],F:604,M:550,n:3640 | This study | abyss | - | + | + | + | + | + |
| LR15337 | Ingroup | Cypriniformes | Cobitoidei | Nemacheilidae | Micronemacheilus | sp | 89.56 | 3260 | 0.55 | 20 | 5.52 | 201 | 0.03 | 1 | 4.34 | 158 | 3280 | 202 | 5.55 | C:3280[S:3260,D:20],F:202,M:158,n:3640 | This study | abyss | + | + | + | + | + | + |
| LR15459 | Ingroup | Cypriniformes | Cobitoidei | Nemacheilidae | Nemacheilus | selangoricus | 88.68 | 3228 | 1.13 | 41 | 6.02 | 219 | 0.08 | 3 | 4.09 | 149 | 3269 | 222 | 6.1 | C:3269[S:3228,D:41],F:222,M:149,n:3640 | This study | abyss | + | + | + | + | + | + |
| LR15611 | Ingroup | Cypriniformes | Cobitoidei | Nemacheilidae | Neonoemacheilus | sp | 76.07 | 2769 | 2.47 | 90 | 12.06 | 439 | 0.19 | 7 | 9.2 | 335 | 2859 | 446 | 12.25 | C:2859[S:2769,D:90],F:446,M:335,n:3640 | This study | abyss | + | + | + | + | + | + |
| LR15490 | Ingroup | Cypriniformes | Cobitoidei | Nemacheilidae | Oreonectes | platycephalus | 82.42 | 3000 | 2.47 | 90 | 8.21 | 299 | 0.14 | 5 | 6.76 | 246 | 3090 | 304 | 8.35 | C:3090[S:3000,D:90],F:304,M:246,n:3640 | This study | abyss | + | + | + | + | + | + |
| LR15504 | Ingroup | Cypriniformes | Cobitoidei | Nemacheilidae | Paracanthocobitis | botia | 75.36 | 2743 | 1.24 | 45 | 12.94 | 471 | 0.14 | 5 | 10.33 | 376 | 2788 | 476 | 13.08 | C:2788[S:2743,D:45],F:476,M:376,n:3640 | This study | abyss | + | + | + | + | + | + |
| LR15189 | Ingroup | Cypriniformes | Cobitoidei | Nemacheilidae | Petruichthys | brevis | 83.24 | 3030 | 0.77 | 28 | 9.62 | 350 | 0.08 | 3 | 6.29 | 229 | 3058 | 353 | 9.7 | C:3058[S:3030,D:28],F:353,M:229,n:3640 | This study | abyss | + | + | + | + | + | + |
| LR15152 | Ingroup | Cypriniformes | Cobitoidei | Nemacheilidae | Schistura | fasciolata | 78.21 | 2847 | 0.91 | 33 | 12.61 | 459 | 0.14 | 5 | 8.13 | 296 | 2880 | 464 | 12.75 | C:2880[S:2847,D:33],F:464,M:296,n:3640 | This study | abyss | + | + | + | + | + | + |
| LR15454 | Ingroup | Cypriniformes | Cobitoidei | Nemacheilidae | Schistura | sp1 | 92.12 | 3353 | 1.26 | 46 | 4.2 | 153 | 0 | 0 | 2.42 | 88 | 3399 | 153 | 4.2 | C:3399[S:3353,D:46],F:153,M:88,n:3640 | This study | abyss | + | + | + | + | + | + |
| LR15496 | Ingroup | Cypriniformes | Cobitoidei | Nemacheilidae | Schistura | sp2 | 43.46 | 1582 | 7.72 | 281 | 21.02 | 765 | 0.63 | 23 | 27.17 | 989 | 1863 | 788 | 21.65 | C:1863[S:1582,D:281],F:788,M:989,n:3640 | This study | abyss | + | + | + | + | + | + |
| LR15455 | Ingroup | Cypriniformes | Cobitoidei | Nemacheilidae | Sundoreonectes | sabanus | 82.45 | 3001 | 1.4 | 51 | 9.37 | 341 | 0.08 | 3 | 6.7 | 244 | 3052 | 344 | 9.45 | C:3052[S:3001,D:51],F:344,M:244,n:3640 | This study | abyss | + | + | + | + | + | + |
| LR15072 | Ingroup | Cypriniformes | Cobitoidei | Nemacheilidae | Traccatichthys | pulcher | 82.09 | 2988 | 1.46 | 53 | 9.78 | 356 | 0.08 | 3 | 6.59 | 240 | 3041 | 359 | 9.86 | C:3041[S:2988,D:53],F:359,M:240,n:3640 | This study | abyss | + | + | + | + | + | + |
| GCF_015846415.1 | Ingroup | Cypriniformes | Cobitoidei | Nemacheilidae | Triplophysa | dalaica | 96.37 | 3508 | 0.82 | 30 | 0.49 | 18 | 0 | 0 | 2.31 | 84 | 3538 | 18 | 0.49 | C:3538[S:3508,D:30],F:18,M:84,n:3640 | NCBI | multiple assembly methods | + | + | + | + | + | + |
| GCF_024868665.1 | Ingroup | Cypriniformes | Cobitoidei | Nemacheilidae | Triplophysa | rosa | 97.64 | 3554 | 0.49 | 18 | 0.66 | 24 | 0 | 0 | 1.21 | 44 | 3572 | 24 | 0.66 | C:3572[S:3554,D:18],F:24,M:44,n:3640 | NCBI | multiple assembly methods | + | + | + | + | + | + |
| GCA_006030095.1 | Ingroup | Cypriniformes | Cobitoidei | Nemacheilidae | Triplophysa | siluroides | 97.45 | 3547 | 0.66 | 24 | 0.55 | 20 | 0 | 0 | 1.35 | 49 | 3571 | 20 | 0.55 | C:3571[S:3547,D:24],F:20,M:49,n:3640 | NCBI | multiple assembly methods | + | + | + | + | + | + |
| GCA_008369825.1 | Ingroup | Cypriniformes | Cobitoidei | Nemacheilidae | Triplophysa | tibetana | 96.24 | 3503 | 2.09 | 76 | 0.41 | 15 | 0 | 0 | 1.26 | 46 | 3579 | 15 | 0.41 | C:3579[S:3503,D:76],F:15,M:46,n:3640 | NCBI | multiple assembly methods | + | + | + | + | + | + |
| GCA_033220385.1 | Ingroup | Cypriniformes | Cobitoidei | Nemacheilidae | Triplophysa | yarkandensis | 94.48 | 3439 | 2.55 | 93 | 0.88 | 32 | 0 | 0 | 2.09 | 76 | 3532 | 32 | 0.88 | C:3532[S:3439,D:93],F:32,M:76,n:3640 | NCBI | multiple assembly methods | + | + | + | + | + | + |
| LR15357 | Ingroup | Cypriniformes | Cobitoidei | Serpenticobitidae | Serpenticobitis | zonata | 89.53 | 3259 | 0.47 | 17 | 6.1 | 222 | 0.08 | 3 | 3.82 | 139 | 3276 | 225 | 6.18 | C:3276[S:3259,D:17],F:225,M:139,n:3640 | This study | spades | + | + | + | + | + | + |
| LR15334 | Ingroup | Cypriniformes | Cobitoidei | Vaillantellidae | Vaillantella | maassii | 93.74 | 3412 | 0.38 | 14 | 3.52 | 128 | 0 | 0 | 2.36 | 86 | 3426 | 128 | 3.52 | C:3426[S:3412,D:14],F:128,M:86,n:3640 | This study | abyss | + | + | + | + | + | + |
| LR15562 | Ingroup | Cypriniformes | Cyprinoidei | Acheilognathidae | Acheilognathus | deignani | 57.66 | 2099 | 0.16 | 6 | 21.13 | 769 | 0.22 | 8 | 20.82 | 758 | 2105 | 777 | 21.35 | C:2105[S:2099,D:6],F:777,M:758,n:3640 | This study | spades | + | + | + | + | + | + |
| LR15063 | Ingroup | Cypriniformes | Cyprinoidei | Acheilognathidae | Acheilognathus | sp | 44.95 | 1636 | 0.25 | 9 | 24.37 | 887 | 0.41 | 15 | 30.03 | 1093 | 1645 | 902 | 24.78 | C:1645[S:1636,D:9],F:902,M:1093,n:3640 | This study | spades | + | + | + | + | + | + |
| LR15636 | Ingroup | Cypriniformes | Cyprinoidei | Acheilognathidae | Paratanakia | himantegus | 57.5 | 2093 | 0.77 | 28 | 20.27 | 738 | 0.19 | 7 | 21.26 | 774 | 2121 | 745 | 20.46 | C:2121[S:2093,D:28],F:745,M:774,n:3640 | This study | abyss | + | + | + | + | + | + |
| LR15661 | Ingroup | Cypriniformes | Cyprinoidei | Acheilognathidae | Rhodeus | amarus | 52.2 | 1900 | 0.27 | 10 | 22.75 | 828 | 0.16 | 6 | 24.62 | 896 | 1910 | 834 | 22.91 | C:1910[S:1900,D:10],F:834,M:896,n:3640 | This study | abyss | + | + | + | + | + | + |
| GCA_037043235.1 | Ingroup | Cypriniformes | Cyprinoidei | Acheilognathidae | Rhodeus | uyekii | 97.28 | 3541 | 1.02 | 37 | 0.93 | 34 | 0 | 0 | 0.77 | 28 | 3578 | 34 | 0.93 | C:3578[S:3541,D:37],F:34,M:28,n:3640 | NCBI | multiple assembly methods | + | + | + | + | + | + |
| LR15266 | Ingroup | Cypriniformes | Cyprinoidei | Cyprinidae | Acapoeta | tanganicae | 35 | 1274 | 13.74 | 500 | 25.82 | 940 | 0.6 | 22 | 24.84 | 904 | 1774 | 962 | 26.42 | C:1774[S:1274,D:500],F:962,M:904,n:3640 | This study | abyss | + | + | + | + | + | + |
| GCA_039880705.1 | Ingroup | Cypriniformes | Cyprinoidei | Cyprinidae | Acrossocheilus | fasciatus | 99.15 | 3609 | 0.41 | 15 | 0.33 | 12 | 0 | 0 | 0.11 | 4 | 3624 | 12 | 0.33 | C:3624[S:3609,D:15],F:12,M:4,n:3640 | NCBI | multiple assembly methods | + | - | + | + | + | + |
| LR15615 | Ingroup | Cypriniformes | Cyprinoidei | Cyprinidae | Ageneiogarra | sp | 31.07 | 1131 | 0.33 | 12 | 30.36 | 1105 | 0.52 | 19 | 37.72 | 1373 | 1143 | 1124 | 30.88 | C:1143[S:1131,D:12],F:1124,M:1373,n:3640 | This study | spades | + | + | + | + | + | + |
| GCA_023376895.1 | Ingroup | Cypriniformes | Cyprinoidei | Cyprinidae | Aspiorhynchus | laticeps | 69.45 | 2528 | 27.8 | 1012 | 0.58 | 21 | 0 | 0 | 2.17 | 79 | 3540 | 21 | 0.58 | C:3540[S:2528,D:1012],F:21,M:79,n:3640 | NCBI | multiple assembly methods | + | + | + | + | + | + |
| LR15639 | Ingroup | Cypriniformes | Cyprinoidei | Cyprinidae | Balantiocheilos | melanopterus | 67.12 | 2443 | 0.36 | 13 | 17.88 | 651 | 0.27 | 10 | 14.37 | 523 | 2456 | 661 | 18.15 | C:2456[S:2443,D:13],F:661,M:523,n:3640 | This study | spades | + | + | + | + | + | + |
| LR15079 | Ingroup | Cypriniformes | Cyprinoidei | Cyprinidae | Barbichthys | laevis | 66.92 | 2436 | 0.38 | 14 | 18.1 | 659 | 0.25 | 9 | 14.34 | 522 | 2450 | 668 | 18.35 | C:2450[S:2436,D:14],F:668,M:522,n:3640 | This study | spades | + | + | + | + | + | + |
| LR15102 | Ingroup | Cypriniformes | Cyprinoidei | Cyprinidae | Barbodes | lateristriga | 84.2 | 3065 | 0.41 | 15 | 10.41 | 379 | 0.11 | 4 | 4.86 | 177 | 3080 | 383 | 10.52 | C:3080[S:3065,D:15],F:383,M:177,n:3640 | This study | abyss | + | + | + | + | + | + |
| 120LR15721 | Ingroup | Cypriniformes | Cyprinoidei | Cyprinidae | Barbodes | sellifer | 42.99 | 1565 | 0.82 | 30 | 24.86 | 905 | 0.44 | 16 | 30.88 | 1124 | 1595 | 921 | 25.3 | C:1595[S:1565,D:30],F:921,M:1124,n:3640 | This study | abyss | + | + | + | + | + | + |
| LR15256 | Ingroup | Cypriniformes | Cyprinoidei | Cyprinidae | Barboides | gracilis | 79.95 | 2910 | 0.47 | 17 | 12.55 | 457 | 0.25 | 9 | 6.79 | 247 | 2927 | 466 | 12.8 | C:2927[S:2910,D:17],F:466,M:247,n:3640 | This study | spades | + | + | + | + | + | + |
| EA0124 | Ingroup | Cypriniformes | Cyprinoidei | Cyprinidae | Barbonymus | altus | 51.21 | 1864 | 0.3 | 11 | 21.48 | 782 | 0.41 | 15 | 26.59 | 968 | 1875 | 797 | 21.89 | C:1875[S:1864,D:11],F:797,M:968,n:3640 | This study | spades | + | + | + | + | + | + |
| GCA_936440315.1 | Ingroup | Cypriniformes | Cyprinoidei | Cyprinidae | Barbus | barbus | 20.47 | 745 | 79.04 | 2877 | 0.41 | 15 | 0 | 0 | 0.08 | 3 | 3622 | 15 | 0.41 | C:3622[S:745,D:2877],F:15,M:3,n:3640 | NCBI | multiple assembly methods | + | + | + | + | + | + |
| GCF_003368295.1 | Ingroup | Cypriniformes | Cyprinoidei | Cyprinidae | Carassius | auratus | 38.08 | 1386 | 61.29 | 2231 | 0.47 | 17 | 0 | 0 | 0.16 | 6 | 3617 | 17 | 0.47 | C:3617[S:1386,D:2231],F:17,M:6,n:3640 | NCBI | multiple assembly methods | + | + | + | + | + | + |
| GCF_963082965.1 | Ingroup | Cypriniformes | Cyprinoidei | Cyprinidae | Carassius | carassius | 47.5 | 1729 | 51.7 | 1882 | 0.47 | 17 | 0 | 0 | 0.33 | 12 | 3611 | 17 | 0.47 | C:3611[S:1729,D:1882],F:17,M:12,n:3640 | NCBI | multiple assembly methods | + | + | + | + | + | + |
| GCA_023724075.1 | Ingroup | Cypriniformes | Cyprinoidei | Cyprinidae | Carassius | gibelio | 44.23 | 1610 | 55.16 | 2008 | 0.41 | 15 | 0 | 0 | 0.19 | 7 | 3618 | 15 | 0.41 | C:3618[S:1610,D:2008],F:15,M:7,n:3640 | NCBI | multiple assembly methods | + | + | + | + | + | + |
| LR15111 | Ingroup | Cypriniformes | Cyprinoidei | Cyprinidae | Ceratogarra | cambodgiensis | 56.54 | 2058 | 0.44 | 16 | 21.37 | 778 | 0.49 | 18 | 21.15 | 770 | 2074 | 796 | 21.86 | C:2074[S:2058,D:16],F:796,M:770,n:3640 | This study | spades | + | + | + | + | + | + |
| EA0401 | Ingroup | Cypriniformes | Cyprinoidei | Cyprinidae | Chagunius | sp | 75.08 | 2733 | 0.49 | 18 | 14.73 | 536 | 0.25 | 9 | 9.45 | 344 | 2751 | 545 | 14.98 | C:2751[S:2733,D:18],F:545,M:344,n:3640 | This study | spades | + | + | + | + | + | + |
| LR15270 | Ingroup | Cypriniformes | Cyprinoidei | Cyprinidae | Clypeobarbus | sp | 66.59 | 2424 | 0.66 | 24 | 17.66 | 643 | 0.27 | 10 | 14.81 | 539 | 2448 | 653 | 17.93 | C:2448[S:2424,D:24],F:653,M:539,n:3640 | This study | abyss | + | + | + | + | + | + |
| LR10955 | Ingroup | Cypriniformes | Cyprinoidei | Cyprinidae | Crossocheilus | reticulatus | 36.32 | 1322 | 0.3 | 11 | 28.98 | 1055 | 0.27 | 10 | 34.12 | 1242 | 1333 | 1065 | 29.25 | C:1333[S:1322,D:11],F:1065,M:1242,n:3640 | This study | spades | + | + | + | + | + | + |
| LR10341 | Ingroup | Cypriniformes | Cyprinoidei | Cyprinidae | Cyclocheilichthys | apogon | 64.86 | 2361 | 0.6 | 22 | 18.32 | 667 | 0.3 | 11 | 15.91 | 579 | 2383 | 678 | 18.62 | C:2383[S:2361,D:22],F:678,M:579,n:3640 | This study | spades | + | + | + | + | + | + |
| GCF_018340385.1 | Ingroup | Cypriniformes | Cyprinoidei | Cyprinidae | Cyprinus | carpio | 41.57 | 1513 | 57.53 | 2094 | 0.41 | 15 | 0 | 0 | 0.49 | 18 | 3607 | 15 | 0.41 | C:3607[S:1513,D:2094],F:15,M:18,n:3640 | NCBI | multiple assembly methods | + | + | + | + | + | + |
| KUFOS.AN.2021.4.1 | Ingroup | Cypriniformes | Cyprinoidei | Cyprinidae | Dawkinsia | filamentosa | 41.87 | 1524 | 0.52 | 19 | 25.47 | 927 | 0.36 | 13 | 31.79 | 1157 | 1543 | 940 | 25.83 | C:1543[S:1524,D:19],F:940,M:1157,n:3640 | This study | spades | - | + | + | + | + | + |
| 140LR15737 | Ingroup | Cypriniformes | Cyprinoidei | Cyprinidae | Desmopuntius | sp | 81.24 | 2957 | 1.51 | 55 | 10.6 | 386 | 0.19 | 7 | 6.46 | 235 | 3012 | 393 | 10.79 | C:3012[S:2957,D:55],F:393,M:235,n:3640 | This study | abyss | + | + | + | + | + | + |
| 122LR15725 | Ingroup | Cypriniformes | Cyprinoidei | Cyprinidae | Discherodontus | ashmeadi | 74.92 | 2727 | 0.47 | 17 | 14.4 | 524 | 0.16 | 6 | 10.05 | 366 | 2744 | 530 | 14.56 | C:2744[S:2727,D:17],F:530,M:366,n:3640 | This study | spades | + | + | + | + | + | + |
| KUFOS.AN.2019.3.1 | Ingroup | Cypriniformes | Cyprinoidei | Cyprinidae | Eechathalakenda | ophicephalus | 29.15 | 1061 | 11.68 | 425 | 26.43 | 962 | 0.55 | 20 | 32.2 | 1172 | 1486 | 982 | 26.98 | C:1486[S:1061,D:425],F:982,M:1172,n:3640 | This study | spades | - | + | + | + | + | + |
| 119LR15720 | Ingroup | Cypriniformes | Cyprinoidei | Cyprinidae | Eirmotus | sp1 | 70.08 | 2551 | 0.52 | 19 | 16.73 | 609 | 0.27 | 10 | 12.39 | 451 | 2570 | 619 | 17 | C:2570[S:2551,D:19],F:619,M:451,n:3640 | This study | spades | + | + | + | + | + | + |
| LR09582 | Ingroup | Cypriniformes | Cyprinoidei | Cyprinidae | Eirmotus | sp2 | 72.12 | 2625 | 0.38 | 14 | 15.85 | 577 | 0.27 | 10 | 11.37 | 414 | 2639 | 587 | 16.12 | C:2639[S:2625,D:14],F:587,M:414,n:3640 | This study | spades | + | + | + | + | + | + |
| GCA_034779775.1 | Ingroup | Cypriniformes | Cyprinoidei | Cyprinidae | Enteromius | afrohamiltoni | 92.77 | 3377 | 0.36 | 13 | 4.2 | 153 | 0 | 0 | 2.66 | 97 | 3390 | 153 | 4.2 | C:3390[S:3377,D:13],F:153,M:97,n:3640 | NCBI | multiple assembly methods | + | + | + | + | + | + |
| GCA_032359395.1 | Ingroup | Cypriniformes | Cyprinoidei | Cyprinidae | Enteromius | anoplus | 88.02 | 3204 | 0.41 | 15 | 7.09 | 258 | 0.03 | 1 | 4.45 | 162 | 3219 | 259 | 7.12 | C:3219[S:3204,D:15],F:259,M:162,n:3640 | NCBI | multiple assembly methods | + | + | + | + | + | + |
| GCA_032357425.1 | Ingroup | Cypriniformes | Cyprinoidei | Cyprinidae | Enteromius | argenteus | 95.58 | 3479 | 0.3 | 11 | 2.55 | 93 | 0.03 | 1 | 1.54 | 56 | 3490 | 94 | 2.58 | C:3490[S:3479,D:11],F:94,M:56,n:3640 | NCBI | multiple assembly methods | + | + | + | + | + | + |
| GCA_032359235.1 | Ingroup | Cypriniformes | Cyprinoidei | Cyprinidae | Enteromius | mattozi | 90.44 | 3292 | 0.47 | 17 | 6.15 | 224 | 0.05 | 2 | 2.88 | 105 | 3309 | 226 | 6.2 | C:3309[S:3292,D:17],F:226,M:105,n:3640 | NCBI | multiple assembly methods | + | + | + | + | + | + |
| GCA_032357445.1 | Ingroup | Cypriniformes | Cyprinoidei | Cyprinidae | Enteromius | radiatus | 95.47 | 3475 | 0.36 | 13 | 2.66 | 97 | 0.05 | 2 | 1.46 | 53 | 3488 | 99 | 2.71 | C:3488[S:3475,D:13],F:99,M:53,n:3640 | NCBI | multiple assembly methods | + | + | + | + | + | + |
| LR15606 | Ingroup | Cypriniformes | Cyprinoidei | Cyprinidae | Enteromius | sp1 | 57.88 | 2107 | 0.3 | 11 | 22.94 | 835 | 0.27 | 10 | 18.6 | 677 | 2118 | 845 | 23.21 | C:2118[S:2107,D:11],F:845,M:677,n:3640 | This study | spades | + | + | + | + | + | + |
| GCA_032359435.1 | Ingroup | Cypriniformes | Cyprinoidei | Cyprinidae | Enteromius | unitaeniatus | 91.62 | 3335 | 0.36 | 13 | 5.33 | 194 | 0.03 | 1 | 2.66 | 97 | 3348 | 195 | 5.36 | C:3348[S:3335,D:13],F:195,M:97,n:3640 | NCBI | multiple assembly methods | + | + | + | + | + | + |
| LR01655 | Ingroup | Cypriniformes | Cyprinoidei | Cyprinidae | Epalzeorhynchos | kalopterum | 30.11 | 1096 | 0.3 | 11 | 26.84 | 977 | 0.55 | 20 | 42.2 | 1536 | 1107 | 997 | 27.39 | C:1107[S:1096,D:11],F:997,M:1536,n:3640 | This study | spades | + | + | + | + | + | + |
| LR15563 | Ingroup | Cypriniformes | Cyprinoidei | Cyprinidae | Folifer | brevifilis | 63.79 | 2322 | 0.44 | 16 | 20.49 | 746 | 0.3 | 11 | 14.97 | 545 | 2338 | 757 | 20.79 | C:2338[S:2322,D:16],F:757,M:545,n:3640 | This study | spades | + | + | + | + | + | + |
| KUFOS.RR.2022.P.4 | Ingroup | Cypriniformes | Cyprinoidei | Cyprinidae | Garra | periyarensis | 47.45 | 1727 | 0.3 | 11 | 25.58 | 931 | 0.22 | 8 | 26.46 | 963 | 1738 | 939 | 25.8 | C:1738[S:1727,D:11],F:939,M:963,n:3640 | This study | spades | - | + | + | + | + | + |
| GCA_027564155.1 | Ingroup | Cypriniformes | Cyprinoidei | Cyprinidae | Gymnocypris | eckloni-scoliostomus | 90.41 | 3291 | 6.18 | 225 | 1.18 | 43 | 0 | 0 | 2.23 | 81 | 3516 | 43 | 1.18 | C:3516[S:3291,D:225],F:43,M:81,n:3640 | NCBI | multiple assembly methods | + | + | + | + | + | + |
| LR15635 | Ingroup | Cypriniformes | Cyprinoidei | Cyprinidae | Haludaria | fasciata | 64.09 | 2333 | 2.2 | 80 | 17.72 | 645 | 0.25 | 9 | 15.74 | 573 | 2413 | 654 | 17.97 | C:2413[S:2333,D:80],F:654,M:573,n:3640 | This study | abyss | + | + | + | + | + | + |
| LR15078 | Ingroup | Cypriniformes | Cyprinoidei | Cyprinidae | Hampala | ampalong | 79.78 | 2904 | 0.44 | 16 | 12.75 | 464 | 0.11 | 4 | 6.92 | 252 | 2920 | 468 | 12.86 | C:2920[S:2904,D:16],F:468,M:252,n:3640 | This study | spades | + | + | + | + | + | + |
| KUFOS.RR.2022.P.3 | Ingroup | Cypriniformes | Cyprinoidei | Cyprinidae | Hypselobarbus | periyarensis | 33.87 | 1233 | 11.37 | 414 | 25.63 | 933 | 0.47 | 17 | 28.65 | 1043 | 1647 | 950 | 26.1 | C:1647[S:1233,D:414],F:950,M:1043,n:3640 | This study | spades | - | + | + | + | + | + |
| GCA_012976165.1 | Ingroup | Cypriniformes | Cyprinoidei | Cyprinidae | Labeo | catla | 96.35 | 3507 | 1.15 | 42 | 1.29 | 47 | 0 | 0 | 1.21 | 44 | 3549 | 47 | 1.29 | C:3549[S:3507,D:42],F:47,M:44,n:3640 | NCBI | multiple assembly methods | + | + | + | + | + | + |
| GCF_022985175.1 | Ingroup | Cypriniformes | Cyprinoidei | Cyprinidae | Labeo | rohita | 99.18 | 3610 | 0.27 | 10 | 0.38 | 14 | 0 | 0 | 0.16 | 6 | 3620 | 14 | 0.38 | C:3620[S:3610,D:10],F:14,M:6,n:3640 | NCBI | multiple assembly methods | + | + | + | + | + | + |
| LR15278 | Ingroup | Cypriniformes | Cyprinoidei | Cyprinidae | Labeo | cylindricus | 59.67 | 2172 | 0.27 | 10 | 22.17 | 807 | 0.05 | 2 | 17.83 | 649 | 2182 | 809 | 22.22 | C:2182[S:2172,D:10],F:809,M:649,n:3640 | This study | spades | + | + | + | + | + | + |
| GCA_034780575.1 | Ingroup | Cypriniformes | Cyprinoidei | Cyprinidae | Labeobarbus | marequensis | 80.71 | 2938 | 4.92 | 179 | 8.57 | 312 | 0.03 | 1 | 5.77 | 210 | 3117 | 313 | 8.6 | C:3117[S:2938,D:179],F:313,M:210,n:3640 | NCBI | multiple assembly methods | + | + | + | + | + | + |
| LR12884 | Ingroup | Cypriniformes | Cyprinoidei | Cyprinidae | Labiobarbus | ocellatus | 27.36 | 996 | 0.22 | 8 | 28.41 | 1034 | 0.47 | 17 | 43.54 | 1585 | 1004 | 1051 | 28.88 | C:1004[S:996,D:8],F:1051,M:1585,n:3640 | This study | spades | + | + | + | + | + | + |
| KUFOS.RR.2022.P.2 | Ingroup | Cypriniformes | Cyprinoidei | Cyprinidae | Lepidopygopsis | typus | 40.74 | 1483 | 15.36 | 559 | 23.1 | 841 | 0.41 | 15 | 20.38 | 742 | 2042 | 856 | 23.51 | C:2042[S:1483,D:559],F:856,M:742,n:3640 | This study | spades | - | + | + | + | + | + |
| LR15104 | Ingroup | Cypriniformes | Cyprinoidei | Cyprinidae | Lobocheilos | aureolineatus | 38.6 | 1405 | 0.14 | 5 | 25.93 | 944 | 0.33 | 12 | 35 | 1274 | 1410 | 956 | 26.26 | C:1410[S:1405,D:5],F:956,M:1274,n:3640 | This study | spades | + | + | + | + | + | + |
| LR15565 | Ingroup | Cypriniformes | Cyprinoidei | Cyprinidae | Mystacoleucus | argenteus | 77.34 | 2815 | 0.36 | 13 | 13.32 | 485 | 0.14 | 5 | 8.85 | 322 | 2828 | 490 | 13.46 | C:2828[S:2815,D:13],F:490,M:322,n:3640 | This study | spades | + | + | + | + | + | + |
| LR15612 | Ingroup | Cypriniformes | Cyprinoidei | Cyprinidae | Neolissochilus | sp | 38.52 | 1402 | 18.63 | 678 | 22.14 | 806 | 0.41 | 15 | 20.3 | 739 | 2080 | 821 | 22.55 | C:2080[S:1402,D:678],F:821,M:739,n:3640 | This study | spades | + | + | + | + | + | + |
| KUFOS.AN.2021.7.1 | Ingroup | Cypriniformes | Cyprinoidei | Cyprinidae | Neolissochilus | wynaadensis | 24.64 | 897 | 5.33 | 194 | 29.59 | 1077 | 0.47 | 17 | 39.97 | 1455 | 1091 | 1094 | 30.06 | C:1091[S:897,D:194],F:1094,M:1455,n:3640 | This study | spades | - | + | + | + | + | + |

| | | | | | | | | | | | | | | | | | | | | | | | | | | | | |
|---|---|---|---|---|---|---|---|---|---|---|---|---|---|---|---|---|---|---|---|---|---|---|---|---|---|---|---|---|
| 143LR15682 | Ingroup | Cypriniformes | Cyprinoidei | Cyprinidae | Oliotius | oligolepis | 81.1 | 2952 | 1.18 | 43 | 10.93 | 398 | 0.08 | 3 | 6.7 | 244 | 2995 | 401 | 11.01 | C:2995[S:2952,D:43],F:401,M:244,n:3640 | This study | abyss | + | + | + | + | + | + |
| GCF_012432095.1 | Ingroup | Cypriniformes | Cyprinoidei | Cyprinidae | Onychostoma | macrolepis | 98.6 | 3589 | 0.38 | 14 | 0.33 | 12 | 0 | 0 | 0.69 | 25 | 3603 | 12 | 0.33 | C:3603[S:3589,D:14],F:12,M:25,n:3640 | NCBI | multiple assembly methods | + | + | + | + | + | + |
| EA1291 | Ingroup | Cypriniformes | Cyprinoidei | Cyprinidae | Oreichthys | sp | 56.48 | 2056 | 0.66 | 24 | 21.35 | 777 | 0.25 | 9 | 21.26 | 774 | 2080 | 786 | 21.6 | C:2080[S:2056,D:24],F:786,M:774,n:3640 | This study | spades | + | + | + | + | + | + |
| LR09367 | Ingroup | Cypriniformes | Cyprinoidei | Cyprinidae | Osteobrama | cunma | 71.43 | 2600 | 0.47 | 17 | 16.07 | 585 | 0.33 | 12 | 11.7 | 426 | 2617 | 597 | 16.4 | C:2617[S:2600,D:17],F:597,M:426,n:3640 | This study | spades | + | + | + | + | + | + |
| LR15096 | Ingroup | Cypriniformes | Cyprinoidei | Cyprinidae | Osteochilus | vittatoides | 21.13 | 769 | 0.03 | 1 | 26.84 | 977 | 0.55 | 20 | 51.46 | 1873 | 770 | 997 | 27.39 | C:770[S:769,D:1],F:997,M:1873,n:3640 | This study | spades | + | + | + | + | + | + |
| GCA_003573665.1 | Ingroup | Cypriniformes | Cyprinoidei | Cyprinidae | Oxygymnocypris | stewartii | 49.29 | 1794 | 48.54 | 1767 | 1.02 | 37 | 0 | 0 | 1.15 | 42 | 3561 | 37 | 1.02 | C:3561[S:1794,D:1767],F:37,M:42,n:3640 | NCBI | multiple assembly methods | + | + | + | + | + | + |
| LR01656 | Ingroup | Cypriniformes | Cyprinoidei | Cyprinidae | Paracrossochilus | sp | 42.09 | 1532 | 0.22 | 8 | 26.98 | 982 | 0.41 | 15 | 30.3 | 1103 | 1540 | 997 | 27.39 | C:1540[S:1532,D:8],F:997,M:1103,n:3640 | This study | spades | + | + | + | + | + | + |
| LR15618 | Ingroup | Cypriniformes | Cyprinoidei | Cyprinidae | Paraspinibarbus | sp | 43.13 | 1570 | 19.12 | 696 | 20.85 | 759 | 0.27 | 10 | 16.62 | 605 | 2266 | 769 | 21.12 | C:2266[S:1570,D:696],F:769,M:605,n:3640 | This study | spades | + | + | + | + | + | + |
| KUFOS.AN.2021.5.1 | Ingroup | Cypriniformes | Cyprinoidei | Cyprinidae | Pethia | conchonius | 44.64 | 1625 | 1.65 | 60 | 24.56 | 894 | 0.27 | 10 | 28.87 | 1051 | 1685 | 904 | 24.83 | C:1685[S:1625,D:60],F:904,M:1051,n:3640 | This study | abyss | - | + | + | + | + | + |
| LR06584 | Ingroup | Cypriniformes | Cyprinoidei | Cyprinidae | Pethia | sp1 | 75.8 | 2759 | 0.47 | 17 | 14.01 | 510 | 0.22 | 8 | 9.51 | 346 | 2776 | 518 | 14.23 | C:2776[S:2759,D:17],F:518,M:346,n:3640 | This study | spades | + | + | + | + | + | + |
| LR15646 | Ingroup | Cypriniformes | Cyprinoidei | Cyprinidae | Pethia | sp2 | 62.94 | 2291 | 0.91 | 33 | 19.78 | 720 | 0.27 | 10 | 16.1 | 586 | 2324 | 730 | 20.05 | C:2324[S:2291,D:33],F:730,M:586,n:3640 | This study | abyss | + | + | + | + | + | + |
| GCA_004124795.1 | Ingroup | Cypriniformes | Cyprinoidei | Cyprinidae | Poropuntius | huangchuchieni | 37.12 | 1351 | 0.33 | 12 | 25.16 | 916 | 0.36 | 13 | 37.03 | 1348 | 1363 | 929 | 25.52 | C:1363[S:1351,D:12],F:929,M:1348,n:3640 | NCBI | multiple assembly methods | + | + | + | + | + | + |
| LR15083 | Ingroup | Cypriniformes | Cyprinoidei | Cyprinidae | Poropuntius | sp | 66.62 | 2425 | 0.49 | 18 | 18.49 | 673 | 0.25 | 9 | 14.15 | 515 | 2443 | 682 | 18.74 | C:2443[S:2425,D:18],F:682,M:515,n:3640 | This study | spades | + | + | + | + | + | + |
| LR15065 | Ingroup | Cypriniformes | Cyprinoidei | Cyprinidae | Probarbus | jullieni | 49.53 | 1803 | 25.8 | 939 | 15.66 | 570 | 0.19 | 7 | 8.82 | 321 | 2742 | 577 | 15.85 | C:2742[S:1803,D:939],F:577,M:321,n:3640 | This study | abyss | + | + | + | + | + | + |
| 125LR15728 | Ingroup | Cypriniformes | Cyprinoidei | Cyprinidae | Puntigrus | partipentazona | 62.39 | 2271 | 1.98 | 72 | 18.46 | 672 | 0.11 | 4 | 17.06 | 621 | 2343 | 676 | 18.57 | C:2343[S:2271,D:72],F:676,M:621,n:3640 | This study | abyss | + | + | + | + | + | + |
| GCF_018831695.1 | Ingroup | Cypriniformes | Cyprinoidei | Cyprinidae | Puntigrus | tetrazona | 95.03 | 3459 | 4.48 | 163 | 0.38 | 14 | 0 | 0 | 0.11 | 4 | 3622 | 14 | 0.38 | C:3622[S:3459,D:163],F:14,M:4,n:3640 | NCBI | multiple assembly methods | + | + | + | + | + | + |
| LR15108 | Ingroup | Cypriniformes | Cyprinoidei | Cyprinidae | Puntioplites | bulu | 66.59 | 2424 | 0.6 | 22 | 18.19 | 662 | 0.22 | 8 | 14.4 | 524 | 2446 | 670 | 18.41 | C:2446[S:2424,D:22],F:670,M:524,n:3640 | This study | spades | + | + | + | + | + | + |
| EA1300 | Ingroup | Cypriniformes | Cyprinoidei | Cyprinidae | Puntius | sp | 53.71 | 1955 | 0.44 | 16 | 21.04 | 766 | 0.33 | 12 | 24.48 | 891 | 1971 | 778 | 21.37 | C:1971[S:1955,D:16],F:778,M:891,n:3640 | This study | spades | + | + | + | + | + | + |
| LR15257 | Ingroup | Cypriniformes | Cyprinoidei | Cyprinidae | Rohanella | titteya | 55.05 | 2004 | 1.84 | 67 | 20.71 | 754 | 0.36 | 13 | 22.03 | 802 | 2071 | 767 | 21.07 | C:2071[S:2004,D:67],F:767,M:802,n:3640 | This study | abyss | + | + | + | + | + | + |
| 124LR15727 | Ingroup | Cypriniformes | Cyprinoidei | Cyprinidae | Rohtee | ogilbii | 71.81 | 2614 | 0.41 | 15 | 15.85 | 577 | 0.19 | 7 | 11.73 | 427 | 2629 | 584 | 16.04 | C:2629[S:2614,D:15],F:584,M:427,n:3640 | This study | spades | + | + | + | + | + | + |
| 145LR15691 | Ingroup | Cypriniformes | Cyprinoidei | Cyprinidae | Sahyadria | denisonii | 63.93 | 2327 | 0.38 | 14 | 17.58 | 640 | 0.25 | 9 | 17.86 | 650 | 2341 | 649 | 17.83 | C:2341[S:2327,D:14],F:649,M:650,n:3640 | This study | spades | + | + | + | + | + | + |
| LR15061 | Ingroup | Cypriniformes | Cyprinoidei | Cyprinidae | Sawbwa | resplendens | 58.41 | 2126 | 2.03 | 74 | 21.68 | 789 | 0.19 | 7 | 17.69 | 644 | 2200 | 796 | 21.87 | C:2200[S:2126,D:74],F:796,M:644,n:3640 | This study | abyss | + | + | + | + | + | + |
| LR15564 | Ingroup | Cypriniformes | Cyprinoidei | Cyprinidae | Scaphiodonichthys | acanthopterus | 72.01 | 2621 | 1.21 | 44 | 15.71 | 572 | 0.14 | 5 | 10.93 | 398 | 2665 | 577 | 15.85 | C:2665[S:2621,D:44],F:577,M:398,n:3640 | This study | abyss | + | + | + | + | + | + |
| LR15101 | Ingroup | Cypriniformes | Cyprinoidei | Cyprinidae | Schismatorhynchos | heterorhynchos | 58.38 | 2125 | 0.63 | 23 | 21.79 | 793 | 0.16 | 6 | 19.04 | 693 | 2148 | 799 | 21.95 | C:2148[S:2125,D:23],F:799,M:693,n:3640 | This study | spades | + | + | + | + | + | + |
| LR09392 | Ingroup | Cypriniformes | Cyprinoidei | Cyprinidae | Semiplotus | sp | 64.09 | 2333 | 0.38 | 14 | 19.51 | 710 | 0.36 | 13 | 15.66 | 570 | 2347 | 723 | 19.87 | C:2347[S:2333,D:14],F:723,M:570,n:3640 | This study | spades | + | + | + | + | + | + |
| GCF_001515605.1 | Ingroup | Cypriniformes | Cyprinoidei | Cyprinidae | Sinocyclocheilus | anshuiensis | 44.89 | 1634 | 54.01 | 1966 | 0.71 | 26 | 0 | 0 | 0.38 | 14 | 3600 | 26 | 0.71 | C:3600[S:1634,D:1966],F:26,M:14,n:3640 | NCBI | multiple assembly methods | + | + | + | + | + | + |
| GCF_001515645.1 | Ingroup | Cypriniformes | Cyprinoidei | Cyprinidae | Sinocyclocheilus | grahami | 59.07 | 2150 | 36.48 | 1328 | 2.75 | 100 | 0.03 | 1 | 1.68 | 61 | 3478 | 101 | 2.78 | C:3478[S:2150,D:1328],F:101,M:61,n:3640 | NCBI | multiple assembly methods | + | + | + | + | + | + |
| GCA_018148995.1 | Ingroup | Cypriniformes | Cyprinoidei | Cyprinidae | Sinocyclocheilus | maitianheensis | 42.69 | 1554 | 56.37 | 2052 | 0.66 | 24 | 0 | 0 | 0.27 | 10 | 3606 | 24 | 0.66 | C:3606[S:1554,D:2052],F:24,M:10,n:3640 | NCBI | multiple assembly methods | + | + | + | + | + | + |
| GCF_001515625.1 | Ingroup | Cypriniformes | Cyprinoidei | Cyprinidae | Sinocyclocheilus | rhinocerous | 47.83 | 1741 | 50.41 | 1835 | 1.21 | 44 | 0 | 0 | 0.55 | 20 | 3576 | 44 | 1.21 | C:3576[S:1741,D:1835],F:44,M:20,n:3640 | NCBI | multiple assembly methods | + | + | + | + | + | + |
| 111LR15711 | Ingroup | Cypriniformes | Cyprinoidei | Cyprinidae | Spinibarbichthys | denticulatus | 22.75 | 828 | 5.3 | 193 | 30.49 | 1110 | 0.49 | 18 | 40.96 | 1491 | 1021 | 1128 | 30.98 | C:1021[S:828,D:193],F:1128,M:1491,n:3640 | This study | spades | + | + | + | + | + | + |
| LR15617 | Ingroup | Cypriniformes | Cyprinoidei | Cyprinidae | Spinibarbichthys | sp | 31.68 | 1153 | 10.14 | 369 | 28.46 | 1036 | 0.58 | 21 | 29.15 | 1061 | 1522 | 1057 | 29.04 | C:1522[S:1153,D:369],F:1057,M:1061,n:3640 | This study | spades | + | + | + | + | + | + |
| 126LR15731 | Ingroup | Cypriniformes | Cyprinoidei | Cyprinidae | Striuntius | lineatus | 72.77 | 2649 | 0.44 | 16 | 15.16 | 552 | 0.19 | 7 | 11.43 | 416 | 2665 | 559 | 15.35 | C:2665[S:2649,D:16],F:559,M:416,n:3640 | This study | spades | + | + | + | + | + | + |
| LR06590 | Ingroup | Cypriniformes | Cyprinoidei | Cyprinidae | Systomus | binduchitra | 72.5 | 2639 | 0.52 | 19 | 15.38 | 560 | 0.11 | 4 | 11.48 | 418 | 2658 | 564 | 15.49 | C:2658[S:2639,D:19],F:564,M:418,n:3640 | This study | spades | + | + | + | + | + | + |
| KUFOS.RR.2022.P.1 | Ingroup | Cypriniformes | Cyprinoidei | Cyprinidae | Tariqilabeo | periyarensis | 39.42 | 1435 | 0.27 | 10 | 29.04 | 1057 | 0.58 | 21 | 30.69 | 1117 | 1445 | 1078 | 29.62 | C:1445[S:1435,D:10],F:1078,M:1117,n:3640 | This study | spades | - | + | + | + | + | + |
| LR12880 | Ingroup | Cypriniformes | Cyprinoidei | Cyprinidae | Thynnichthys | sp | 40.6 | 1478 | 0.25 | 9 | 26.95 | 981 | 0.3 | 11 | 31.9 | 1161 | 1487 | 992 | 27.25 | C:1487[S:1478,D:9],F:992,M:1161,n:3640 | This study | spades | + | + | + | + | + | + |
| LR09401 | Ingroup | Cypriniformes | Cyprinoidei | Cyprinidae | Tor | sp1 | 42.23 | 1537 | 22.66 | 825 | 19.59 | 713 | 0.47 | 17 | 15.05 | 548 | 2362 | 730 | 20.06 | C:2362[S:1537,D:825],F:730,M:548,n:3640 | This study | spades | + | + | + | + | + | + |
| LR15100 | Ingroup | Cypriniformes | Cyprinoidei | Cyprinidae | Tor | sp2 | 38.87 | 1415 | 17.88 | 651 | 22.99 | 837 | 0.47 | 17 | 19.78 | 720 | 2066 | 854 | 23.46 | C:2066[S:1415,D:651],F:854,M:720,n:3640 | This study | spades | + | + | + | + | + | + |
| LR01638 | Ingroup | Cypriniformes | Cyprinoidei | Danionidae | Amblypharyngodon | chulabhornae | 82.17 | 2991 | 0.44 | 16 | 10.41 | 379 | 0.16 | 6 | 6.81 | 248 | 3007 | 385 | 10.57 | C:3007[S:2991,D:16],F:385,M:248,n:3640 | This study | spades | + | + | + | + | + | + |
| fAmbSpp1 | Ingroup | Cypriniformes | Cyprinoidei | Danionidae | Amblypharyngodon | sp | 81.1 | 2952 | 0.69 | 25 | 11.26 | 410 | 0.08 | 3 | 6.87 | 250 | 2977 | 413 | 11.34 | C:2977[S:2952,D:25],F:413,M:250,n:3640 | NCBI-SRA | abyss | + | + | + | + | + | + |
| LR15576 | Ingroup | Cypriniformes | Cyprinoidei | Danionidae | Barilius | mesopotamicus | 80.11 | 2916 | 0.47 | 17 | 12.14 | 442 | 0.16 | 6 | 7.12 | 259 | 2933 | 448 | 12.3 | C:2933[S:2916,D:17],F:448,M:259,n:3640 | This study | spades | + | + | + | + | + | + |
| LR15871 | Ingroup | Cypriniformes | Cyprinoidei | Danionidae | Boraras | brigittae | 97.58 | 3552 | 1.02 | 37 | 0.47 | 17 | 0 | 0 | 0.93 | 34 | 3589 | 17 | 0.47 | C:3589[S:3552,D:37],F:17,M:34,n:3640 | This study | hifiasm.primary.HiC.scaffolded | + | + | + | + | + | + |
| fBorMac1 | Ingroup | Cypriniformes | Cyprinoidei | Danionidae | Boraras | maculatus | 47.09 | 1714 | 1.13 | 41 | 19.95 | 726 | 0.41 | 15 | 31.43 | 1144 | 1755 | 741 | 20.36 | C:1755[S:1714,D:41],F:741,M:1144,n:3640 | NCBI-SRA | abyss | + | + | + | + | + | + |
| LR12206 | Ingroup | Cypriniformes | Cyprinoidei | Danionidae | Boraras | micros | 39.73 | 1446 | 0.3 | 11 | 22.83 | 831 | 0.49 | 18 | 36.65 | 1334 | 1457 | 849 | 23.32 | C:1457[S:1446,D:11],F:849,M:1334,n:3640 | This study | spades | + | + | + | + | + | + |
| LR12190 | Ingroup | Cypriniformes | Cyprinoidei | Danionidae | Boraras | urophthalmoides | 38.13 | 1388 | 0.19 | 7 | 23.54 | 857 | 0.41 | 15 | 37.72 | 1373 | 1395 | 872 | 23.95 | C:1395[S:1388,D:7],F:872,M:1373,n:3640 | This study | spades | + | + | + | + | + | + |
| LR09154 | Ingroup | Cypriniformes | Cyprinoidei | Danionidae | Brevibora | cheeya1 | 35.71 | 1300 | 0.27 | 10 | 24.95 | 908 | 0.33 | 12 | 38.74 | 1410 | 1310 | 920 | 25.28 | C:1310[S:1300,D:10],F:920,M:1410,n:3640 | This study | spades | + | + | + | + | + | + |
| LR10130 | Ingroup | Cypriniformes | Cyprinoidei | Danionidae | Brevibora | cheeya2 | 27.34 | 995 | 0.25 | 9 | 24.34 | 886 | 0.38 | 14 | 47.69 | 1736 | 1004 | 900 | 24.72 | C:1004[S:995,D:9],F:900,M:1736,n:3640 | This study | spades | + | + | + | + | + | + |
| LR11616 | Ingroup | Cypriniformes | Cyprinoidei | Danionidae | Brevibora | dorsiocellata | 38.1 | 1387 | 0.36 | 13 | 24.23 | 882 | 0.44 | 16 | 36.87 | 1342 | 1400 | 898 | 24.67 | C:1400[S:1387,D:13],F:898,M:1342,n:3640 | This study | spades | + | + | + | + | + | + |
| LR15567 | Ingroup | Cypriniformes | Cyprinoidei | Danionidae | Cabdio | morar | 62.64 | 2280 | 4.37 | 159 | 18.71 | 681 | 0.38 | 14 | 13.9 | 506 | 2439 | 695 | 19.09 | C:2439[S:2280,D:159],F:695,M:506,n:3640 | This study | abyss | + | + | + | + | + | + |
| fCheCac1 | Ingroup | Cypriniformes | Cyprinoidei | Danionidae | Chela | cachius | 23.82 | 867 | 0.44 | 16 | 26.07 | 949 | 0.38 | 14 | 49.29 | 1794 | 883 | 963 | 26.45 | C:883[S:867,D:16],F:963,M:1794,n:3640 | NCBI-SRA | abyss | + | + | + | + | + | + |
| 131LR08589 | Ingroup | Cypriniformes | Cyprinoidei | Danionidae | Chelaethiops | elongatus | 77.55 | 2823 | 2.36 | 86 | 11.35 | 413 | 0.22 | 8 | 8.52 | 310 | 2909 | 421 | 11.57 | C:2909[S:2823,D:86],F:421,M:310,n:3640 | This study | abyss | + | + | + | + | + | + |
| LR15596 | Ingroup | Cypriniformes | Cyprinoidei | Danionidae | Chelaethiops | sp1 | 85.91 | 3127 | 0.52 | 19 | 8.96 | 326 | 0.08 | 3 | 4.53 | 165 | 3146 | 329 | 9.04 | C:3146[S:3127,D:19],F:329,M:165,n:3640 | This study | spades | + | + | + | + | + | + |
| LR15597 | Ingroup | Cypriniformes | Cyprinoidei | Danionidae | Chelaethiops | sp2 | 88.19 | 3210 | 0.3 | 11 | 7.88 | 287 | 0.03 | 1 | 3.6 | 131 | 3221 | 288 | 7.91 | C:3221[S:3210,D:11],F:288,M:131,n:3640 | This study | spades | + | + | + | + | + | + |
| GCF_903798145.1 | Ingroup | Cypriniformes | Cyprinoidei | Danionidae | Danio | aesculapii | 96.18 | 3501 | 0.93 | 34 | 0.96 | 35 | 0 | 0 | 1.92 | 70 | 3535 | 35 | 0.96 | C:3535[S:3501,D:34],F:35,M:70,n:3640 | NCBI | multiple assembly methods | + | + | + | + | + | + |
| GCA_903798035.1 | Ingroup | Cypriniformes | Cyprinoidei | Danionidae | Danio | albolineatus | 79.89 | 2908 | 1.04 | 38 | 9.45 | 344 | 0.05 | 2 | 9.56 | 348 | 2946 | 346 | 9.5 | C:2946[S:2908,D:38],F:346,M:348,n:3640 | NCBI | multiple assembly methods | + | + | + | + | + | + |
| GCA_903798125.1 | Ingroup | Cypriniformes | Cyprinoidei | Danionidae | Danio | choprae | 74.26 | 2703 | 0.55 | 20 | 11.51 | 419 | 0.05 | 2 | 13.63 | 496 | 2723 | 421 | 11.56 | C:2723[S:2703,D:20],F:421,M:496,n:3640 | NCBI | multiple assembly methods | + | + | + | + | + | + |
| LR01642 | Ingroup | Cypriniformes | Cyprinoidei | Danionidae | Danio | erythromicron | 50.47 | 1837 | 0.44 | 16 | 22.23 | 809 | 0.3 | 11 | 26.57 | 967 | 1853 | 820 | 22.53 | C:1853[S:1837,D:16],F:820,M:967,n:3640 | This study | spades | + | + | + | + | + | + |
| GCA_903798115.1 | Ingroup | Cypriniformes | Cyprinoidei | Danionidae | Danio | jaintianensis | 88.32 | 3215 | 0.47 | 17 | 5.05 | 184 | 0 | 0 | 6.15 | 224 | 3232 | 184 | 5.05 | C:3232[S:3215,D:17],F:184,M:224,n:3640 | NCBI | multiple assembly methods | + | + | + | + | + | + |
| GCA_903798195.1 | Ingroup | Cypriniformes | Cyprinoidei | Danionidae | Danio | kyathit | 96.9 | 3527 | 0.93 | 34 | 0.74 | 27 | 0 | 0 | 1.43 | 52 | 3561 | 27 | 0.74 | C:3561[S:3527,D:34],F:27,M:52,n:3640 | NCBI | multiple assembly methods | + | + | + | + | + | + |
| LR02056 | Ingroup | Cypriniformes | Cyprinoidei | Danionidae | Danio | margaritatus | 50.38 | 1834 | 0.36 | 13 | 23.3 | 848 | 0.33 | 12 | 25.63 | 933 | 1847 | 860 | 23.63 | C:1847[S:1834,D:13],F:860,M:933,n:3640 | This study | spades | + | + | + | + | + | + |
| GCA_944039275.1 | Ingroup | Cypriniformes | Cyprinoidei | Danionidae | Danio | rerio | 99.04 | 3605 | 0.33 | 12 | 0.41 | 15 | 0 | 0 | 0.22 | 8 | 3617 | 15 | 0.41 | C:3617[S:3605,D:12],F:15,M:8,n:3640 | NCBI | multiple assembly methods | + | + | + | + | + | + |
| GCA_903798205.1 | Ingroup | Cypriniformes | Cyprinoidei | Danionidae | Danio | tinwini | 83.3 | 3032 | 0.41 | 15 | 8.57 | 312 | 0.03 | 1 | 7.69 | 280 | 3047 | 313 | 8.6 | C:3047[S:3032,D:15],F:313,M:280,n:3640 | NCBI | multiple assembly methods | + | + | + | + | + | + |
| GCA_903798025.1 | Ingroup | Cypriniformes | Cyprinoidei | Danionidae | Danionella | cerebrum | 93.35 | 3398 | 0.33 | 12 | 2.97 | 108 | 0.03 | 1 | 3.32 | 121 | 3410 | 109 | 3 | C:3410[S:3398,D:12],F:109,M:121,n:3640 | NCBI | multiple assembly methods | + | + | + | + | + | + |
| GCA_900490495.1 | Ingroup | Cypriniformes | Cyprinoidei | Danionidae | Danionella | dracula | 90.44 | 3292 | 2.03 | 74 | 1.48 | 54 | 0 | 0 | 6.04 | 220 | 3366 | 54 | 1.48 | C:3366[S:3292,D:74],F:54,M:220,n:3640 | NCBI | multiple assembly methods | + | + | + | + | + | + |
| fDevDev2 | Ingroup | Cypriniformes | Cyprinoidei | Danionidae | Devario | devario | 25.82 | 940 | 0.25 | 9 | 27.06 | 985 | 0.41 | 15 | 46.46 | 1691 | 949 | 1000 | 27.47 | C:949[S:940,D:9],F:1000,M:1691,n:3640 | NCBI-SRA | spades | + | + | + | + | + | + |
| 144LR15698 | Ingroup | Cypriniformes | Cyprinoidei | Danionidae | Devario | sondhii | 36.9 | 1343 | 0.3 | 11 | 26.81 | 976 | 0.41 | 15 | 35.58 | 1295 | 1354 | 991 | 27.22 | C:1354[S:1343,D:11],F:991,M:1295,n:3640 | This study | spades | + | + | + | + | + | + |
| fDanAeq2 | Ingroup | Cypriniformes | Cyprinoidei | Danionidae | Devario | sp | 17.53 | 638 | 0.19 | 7 | 23.57 | 858 | 0.44 | 16 | 58.27 | 2121 | 645 | 874 | 24.01 | C:645[S:638,D:7],F:874,M:2121,n:3640 | NCBI-SRA | abyss | + | + | + | + | + | + |
| LR15598 | Ingroup | Cypriniformes | Cyprinoidei | Danionidae | Engraulicypris | sp1 | 87.09 | 3170 | 0.47 | 17 | 8.19 | 298 | 0.05 | 2 | 4.2 | 153 | 3187 | 300 | 8.24 | C:3187[S:3170,D:17],F:300,M:153,n:3640 | This study | spades | + | + | + | + | + | + |
| LR15656 | Ingroup | Cypriniformes | Cyprinoidei | Danionidae | Engraulicypris | sp2 | 75.47 | 2747 | 1.95 | 71 | 13.08 | 476 | 0.19 | 7 | 9.31 | 339 | 2818 | 483 | 13.27 | C:2818[S:2747,D:71],F:483,M:339,n:3640 | This study | abyss | + | + | + | + | + | + |
| 116LR15716 | Ingroup | Cypriniformes | Cyprinoidei | Danionidae | Esomus | danricus | 78.63 | 2862 | 0.44 | 16 | 11.04 | 402 | 0.08 | 3 | 9.81 | 357 | 2878 | 405 | 11.12 | C:2878[S:2862,D:16],F:405,M:357,n:3640 | This study | spades | + | + | + | + | + | + |
| LR01653 | Ingroup | Cypriniformes | Cyprinoidei | Danionidae | Esomus | metallicus | 85.99 | 3130 | 2.09 | 76 | 7.06 | 257 | 0.11 | 4 | 4.75 | 173 | 3206 | 261 | 7.17 | C:3206[S:3130,D:76],F:261,M:173,n:3640 | This study | abyss | + | + | + | + | + | + |
| LR12503 | Ingroup | Cypriniformes | Cyprinoidei | Danionidae | Esomus | sp | 86.15 | 3136 | 0.41 | 15 | 8.02 | 292 | 0.14 | 5 | 5.27 | 192 | 3151 | 297 | 8.16 | C:3151[S:3136,D:15],F:297,M:192,n:3640 | This study | spades | + | + | + | + | + | + |
| fHorAtu1 | Ingroup | Cypriniformes | Cyprinoidei | Danionidae | Horadandia | atukorali | 63.21 | 2301 | 4.15 | 151 | 17.61 | 641 | 0.25 | 9 | 14.78 | 538 | 2452 | 650 | 17.86 | C:2452[S:2301,D:151],F:650,M:538,n:3640 | NCBI-SRA | abyss | + | + | + | + | + | + |
| LR01646 | Ingroup | Cypriniformes | Cyprinoidei | Danionidae | Inlecypris | auropurpurea | 26.07 | 949 | 0.16 | 6 | 27.14 | 988 | 0.44 | 16 | 46.18 | 1681 | 955 | 1004 | 27.58 | C:955[S:949,D:6],F:1004,M:1681,n:3640 | This study | spades | + | + | + | + | + | + |
| LR01674 | Ingroup | Cypriniformes | Cyprinoidei | Danionidae | Kottelatia | brittani | 71.32 | 2596 | 0.74 | 27 | 14.18 | 516 | 0.05 | 2 | 13.71 | 499 | 2623 | 518 | 14.23 | C:2623[S:2596,D:27],F:518,M:499,n:3640 | This study | abyss | + | + | + | + | + | + |
| fLauMye1 | Ingroup | Cypriniformes | Cyprinoidei | Danionidae | Laubuka | siamensis | 48.71 | 1773 | 0.47 | 17 | 25.49 | 928 | 0.3 | 11 | 25.03 | 911 | 1790 | 939 | 25.79 | C:1790[S:1773,D:17],F:939,M:911,n:3640 | NCBI-SRA | abyss | + | + | + | + | + | + |
| 132LR09413 | Ingroup | Cypriniformes | Cyprinoidei | Danionidae | Laubuka | sp1 | 44.78 | 1630 | 0.36 | 13 | 27.09 | 986 | 0.19 | 7 | 27.58 | 1004 | 1643 | 993 | 27.28 | C:1643[S:1630,D:13],F:993,M:1004,n:3640 | This study | spades | + | + | + | + | + | + |
| KUFOS.AN.2021.3.1 | Ingroup | Cypriniformes | Cyprinoidei | Danionidae | Laubuka | sp2 | 39.7 | 1445 | 0.52 | 19 | 28.08 | 1022 | 0.3 | 11 | 31.4 | 1143 | 1464 | 1033 | 28.38 | C:1464[S:1445,D:19],F:1033,M:1143,n:3640 | This study | spades | - | + | + | + | + | + |
| LR15601 | Ingroup | Cypriniformes | Cyprinoidei | Danionidae | Leptocypris | sp | 75.52 | 2749 | 0.36 | 13 | 14.23 | 518 | 0.22 | 8 | 9.67 | 352 | 2762 | 526 | 14.45 | C:2762[S:2749,D:13],F:526,M:352,n:3640 | This study | spades | + | + | + | + | + | + |
| fLucSpp1 | Ingroup | Cypriniformes | Cyprinoidei | Danionidae | Luciosoma | sp1 | 79.31 | 2887 | 1.54 | 56 | 11.43 | 416 | 0.27 | 10 | 7.45 | 271 | 2943 | 426 | 11.7 | C:2943[S:2887,D:56],F:426,M:271,n:3640 | NCBI-SRA | abyss | + | + | + | + | + | + |
| LR01659 | Ingroup | Cypriniformes | Cyprinoidei | Danionidae | Luciosoma | sp2 | 76.54 | 2786 | 0.38 | 14 | 13.27 | 483 | 0.19 | 7 | 9.62 | 350 | 2800 | 490 | 13.46 | C:2800[S:2786,D:14],F:490,M:350,n:3640 | This study | spades | + | + | + | + | + | + |
| LR15098 | Ingroup | Cypriniformes | Cyprinoidei | Danionidae | Malayochela | maassi | 56.59 | 2060 | 0.41 | 15 | 22.12 | 805 | 0.33 | 12 | 20.55 | 748 | 2075 | 817 | 22.45 | C:2075[S:2060,D:15],F:817,M:748,n:3640 | This study | spades | + | + | + | + | + | + |
| LR15568 | Ingroup | Cypriniformes | Cyprinoidei | Danionidae | Microdevario | kubotai | 56.73 | 2065 | 0.69 | 25 | 21.51 | 783 | 0.27 | 10 | 20.8 | 757 | 2090 | 793 | 21.78 | C:2090[S:2065,D:25],F:793,M:757,n:3640 | This study | spades | + | + | + | + | + | + |
| LR15650 | Ingroup | Cypriniformes | Cyprinoidei | Danionidae | Microrasbora | rubescens | 32.77 | 1193 | 0.3 | 11 | 27.03 | 984 | 0.41 | 15 | 39.48 | 1437 | 1204 | 999 | 27.44 | C:1204[S:1193,D:11],F:999,M:1437,n:3640 | This study | spades | + | + | + | + | + | + |
| 121LR15723 | Ingroup | Cypriniformes | Cyprinoidei | Danionidae | Nematabramis | everetti | 72.61 | 2643 | 0.41 | 15 | 16.37 | 596 | 0.19 | 7 | 10.41 | 379 | 2658 | 603 | 16.56 | C:2658[S:2643,D:15],F:603,M:379,n:3640 | This study | spades | + | + | + | + | + | + |
| LR15099 | Ingroup | Cypriniformes | Cyprinoidei | Danionidae | Nematabramis | steindachneri | 60.03 | 2185 | 0.41 | 15 | 20.33 | 740 | 0.41 | 15 | 18.82 | 685 | 2200 | 755 | 20.74 | C:2200[S:2185,D:15],F:755,M:685,n:3640 | This study | spades | + | + | + | + | + | + |
| LR14062 | Ingroup | Cypriniformes | Cyprinoidei | Danionidae | Neochela | dadiburjori | 33.49 | 1219 | 0.25 | 9 | 25.22 | 918 | 0.41 | 15 | 40.63 | 1479 | 1228 | 933 | 25.63 | C:1228[S:1219,D:9],F:933,M:1479,n:3640 | This study | spades | + | + | + | + | + | + |
| LR15289 | Ingroup | Cypriniformes | Cyprinoidei | Danionidae | Opsaridium | sp1 | 84.84 | 3088 | 0.52 | 19 | 9.97 | 363 | 0.05 | 2 | 4.62 | 168 | 3107 | 365 | 10.02 | C:3107[S:3088,D:19],F:365,M:168,n:3640 | This study | spades | + | + | + | + | + | + |
| LR15290 | Ingroup | Cypriniformes | Cyprinoidei | Danionidae | Opsaridium | sp2 | 86.37 | 3144 | 0.49 | 18 | 9.31 | 339 | 0.11 | 4 | 3.71 | 135 | 3162 | 343 | 9.42 | C:3162[S:3144,D:18],F:343,M:135,n:3640 | This study | spades | + | + | + | + | + | + |
| LR15602 | Ingroup | Cypriniformes | Cyprinoidei | Danionidae | Opsaridium | sp3 | 86.65 | 3154 | 0.44 | 16 | 8.9 | 324 | 0.03 | 1 | 3.98 | 145 | 3170 | 325 | 8.93 | C:3170[S:3154,D:16],F:325,M:145,n:3640 | This study | spades | + | + | + | + | + | + |
| 142LR15666 | Ingroup | Cypriniformes | Cyprinoidei | Danionidae | Opsarius | sp1 | 90.71 | 3302 | 0.58 | 21 | 6.1 | 222 | 0.05 | 2 | 2.55 | 93 | 3323 | 224 | 6.15 | C:3323[S:3302,D:21],F:224,M:93,n:3640 | This study | spades | + | + | + | + | + | + |
| LR02136 | Ingroup | Cypriniformes | Cyprinoidei | Danionidae | Opsarius | sp2 | 89.4 | 3254 | 0.49 | 18 | 6.43 | 234 | 0.03 | 1 | 3.65 | 133 | 3272 | 235 | 6.46 | C:3272[S:3254,D:18],F:235,M:133,n:3640 | This study | spades | + | + | + | + | + | + |
| LR12240 | Ingroup | Cypriniformes | Cyprinoidei | Danionidae | Opsarius | sp3 | 93.24 | 3394 | 0.3 | 11 | 4.67 | 170 | 0.05 | 2 | 1.73 | 63 | 3405 | 172 | 4.72 | C:3405[S:3394,D:11],F:172,M:63,n:3640 | This study | spades | + | + | + | + | + | + |
| LR15081 | Ingroup | Cypriniformes | Cyprinoidei | Danionidae | Pectenocypris | sp | 66.84 | 2433 | 0.52 | 19 | 17.36 | 632 | 0.27 | 10 | 15 | 546 | 2452 | 642 | 17.63 | C:2452[S:2433,D:19],F:642,M:546,n:3640 | This study | abyss | + | + | + | + | + | + |
| fRaiGut1 | Ingroup | Cypriniformes | Cyprinoidei | Danionidae | Raiamas | guttatus | 85.96 | 3129 | 0.41 | 15 | 8.98 | 327 | 0.08 | 3 | 4.56 | 166 | 3144 | 330 | 9.06 | C:3144[S:3129,D:15],F:330,M:166,n:3640 | NCBI-SRA | spades | + | + | + | + | + | + |
| LR15292 | Ingroup | Cypriniformes | Cyprinoidei | Danionidae | Raiamas | moorii | 86.95 | 3165 | 0.66 | 24 | 8.54 | 311 | 0.08 | 3 | 3.76 | 137 | 3189 | 314 | 8.62 | C:3189[S:3165,D:24],F:314,M:137,n:3640 | This study | abyss | + | + | + | + | + | + |
| 105LR15605 | Ingroup | Cypriniformes | Cyprinoidei | Danionidae | Raiamas | sp | 87.17 | 3173 | 0.49 | 18 | 8.35 | 304 | 0.11 | 4 | 3.87 | 141 | 3191 | 308 | 8.46 | C:3191[S:3173,D:18],F:308,M:141,n:3640 | This study | spades | + | + | + | + | + | + |
| LR10786 | Ingroup | Cypriniformes | Cyprinoidei | Danionidae | Rasbora | cephalotaenia | 57.86 | 2106 | 5.27 | 192 | 17.12 | 623 | 0.16 | 6 | 19.59 | 713 | 2298 | 629 | 17.28 | C:2298[S:2106,D:192],F:629,M:713,n:3640 | This study | abyss | + | + | + | + | + | + |
| KUFOS.AN.2021.2.1 | Ingroup | Cypriniformes | Cyprinoidei | Danionidae | Rasbora | dandia | 49.51 | 1802 | 0.71 | 26 | 21.18 | 771 | 0.25 | 9 | 28.35 | 1032 | 1828 | 780 | 21.43 | C:1828[S:1802,D:26],F:780,M:1032,n:3640 | This study | spades | - | + | + | + | + | + |
| fRasDan1 | Ingroup | Cypriniformes | Cyprinoidei | Danionidae | Rasbora | daniconius | 64.73 | 2356 | 1.26 | 46 | 14.86 | 541 | 0.27 | 10 | 18.87 | 687 | 2402 | 551 | 15.13 | C:2402[S:2356,D:46],F:551,M:687,n:3640 | NCBI-SRA | abyss | + | + | + | + | + | + |
| LR15347 | Ingroup | Cypriniformes | Cyprinoidei | Danionidae | Rasbora | kalbarensis | 49.7 | 1809 | 0.38 | 14 | 20.22 | 736 | 0.3 | 11 | 29.4 | 1070 | 1823 | 747 | 20.52 | C:1823[S:1809,D:14],F:747,M:1070,n:3640 | This study | spades | + | + | + | + | + | + |
| LR15877 | Ingroup | Cypriniformes | Cyprinoidei | Danionidae | Rasbora | kalochroma | 96.73 | 3521 | 2.23 | 81 | 0.44 | 16 | 0 | 0 | 0.6 | 22 | 3602 | 16 | 0.44 | C:3602[S:3521,D:81],F:16,M:22,n:3640 | This study | hifiasm.primary.HiC.scaffolded | + | + | + | + | + | + |
| LR10804 | Ingroup | Cypriniformes | Cyprinoidei | Danionidae | Rasbora | kottelati | 20.38 | 742 | 0.3 | 11 | 24.73 | 900 | 0.58 | 21 | 54.01 | 1966 | 753 | 921 | 25.31 | C:753[S:742,D:11],F:921,M:1966,n:3640 | This study | spades | + | + | + | + | + | + |
| LR09739 | Ingroup | Cypriniformes | Cyprinoidei | Danionidae | Rasbora | myersi | 25.36 | 923 | 1.43 | 52 | 25.3 | 921 | 0.55 | 20 | 47.36 | 1724 | 975 | 941 | 25.85 | C:975[S:923,D:52],F:941,M:1724,n:3640 | This study | abyss | + | + | + | + | + | + |
| fRasRas1 | Ingroup | Cypriniformes | Cyprinoidei | Danionidae | Rasbora | rasbora | 36.26 | 1320 | 1.21 | 44 | 23.85 | 868 | 0.25 | 9 | 38.43 | 1399 | 1364 | 877 | 24.1 | C:1364[S:1320,D:44],F:877,M:1399,n:3640 | NCBI-SRA | abyss | + | + | + | + | + | + |
| LR09847 | Ingroup | Cypriniformes | Cyprinoidei | Danionidae | Rasbora | trilineata | 27.17 | 989 | 0.58 | 21 | 23.6 | 859 | 0.55 | 20 | 48.1 | 1751 | 1010 | 879 | 24.15 | C:1010[S:989,D:21],F:879,M:1751,n:3640 | This study | abyss | + | + | + | + | + | + |
| LR05463 | Ingroup | Cypriniformes | Cyprinoidei | Danionidae | Rasboroides | pallidus | 76.07 | 2769 | 0.58 | 21 | 12.58 | 458 | 0.14 | 5 | 10.63 | 387 | 2790 | 463 | 12.72 | C:2790[S:2769,D:21],F:463,M:387,n:3640 | This study | spades | + | + | + | + | + | + |
| LR12212 | Ingroup | Cypriniformes | Cyprinoidei | Danionidae | Rasbosoma | spilocerca | 44.89 | 1634 | 0.52 | 19 | 24.12 | 878 | 0.36 | 13 | 30.11 | 1096 | 1653 | 891 | 24.48 | C:1653[S:1634,D:19],F:891,M:1096,n:3640 | This study | spades | + | + | + | + | + | + |
| KUFOS.AN.2021.6.1 | Ingroup | Cypriniformes | Cyprinoidei | Danionidae | Salmostoma | balookee | 84.48 | 3075 | 0.49 | 18 | 9.75 | 355 | 0.05 | 2 | 5.22 | 190 | 3093 | 357 | 9.8 | C:3093[S:3075,D:18],F:357,M:190,n:3640 | This study | spades | - | + | + | + | + | + |
| EA0634 | Ingroup | Cypriniformes | Cyprinoidei | Danionidae | Thryssocypris | tonlesapensis | 84.31 | 3069 | 0.33 | 12 | 9.07 | 330 | 0.08 | 3 | 6.21 | 226 | 3081 | 333 | 9.15 | C:3081[S:3069,D:12],F:333,M:226,n:3640 | This study | spades | + | + | + | + | + | + |
| LR08172 | Ingroup | Cypriniformes | Cyprinoidei | Danionidae | Trigonopoma | gracile | 58.65 | 2135 | 0.58 | 21 | 18.49 | 673 | 0.22 | 8 | 22.06 | 803 | 2156 | 681 | 18.71 | C:2156[S:2135,D:21],F:681,M:803,n:3640 | This study | abyss | + | + | + | + | + | + |
| LR10181 | Ingroup | Cypriniformes | Cyprinoidei | Danionidae | Trigonopoma | pauciperforatum | 21.98 | 800 | 0.93 | 34 | 24.26 | 883 | 0.58 | 21 | 52.25 | 1902 | 834 | 904 | 24.84 | C:834[S:800,D:34],F:904,M:1902,n:3640 | This study | abyss | + | + | + | + | + | + |
| LR11656 | Ingroup | Cypriniformes | Cyprinoidei | Danionidae | Trigonopoma | sp | 18.87 | 687 | 0.3 | 11 | 25.19 | 917 | 0.41 | 15 | 55.22 | 2010 | 698 | 932 | 25.6 | C:698[S:687,D:11],F:932,M:2010,n:3640 | This study | spades | + | + | + | + | + | + |
| LR08812 | Ingroup | Cypriniformes | Cyprinoidei | Danionidae | Trigonostigma | espei | 35.27 | 1284 | 0.11 | 4 | 24.29 | 884 | 0.47 | 17 | 39.86 | 1451 | 1288 | 901 | 24.76 | C:1288[S:1284,D:4],F:901,M:1451,n:3640 | This study | spades | + | + | + | + | + | + |
| LR11086 | Ingroup | Cypriniformes | Cyprinoidei | Danionidae | Trigonostigma | heteromorpha | 33.96 | 1236 | 0.11 | 4 | 24.56 | 894 | 0.44 | 16 | 40.93 | 1490 | 1240 | 910 | 25 | C:1240[S:1236,D:4],F:910,M:1490,n:3640 | This study | spades | + | + | + | + | + | + |
| GCA_949357685.1 | Ingroup | Cypriniformes | Cyprinoidei | Gobionidae | Gobio | gobio | 97.94 | 3565 | 0.55 | 20 | 0.69 | 25 | 0 | 0 | 0.82 | 30 | 3585 | 25 | 0.69 | C:3585[S:3565,D:20],F:25,M:30,n:3640 | NCBI | multiple assembly methods | + | + | + | + | + | + |
| GCA_023029165.1 | Ingroup | Cypriniformes | Cyprinoidei | Gobionidae | Gobiocypris | rarus | 96.1 | 3498 | 2.12 | 77 | 0.69 | 25 | 0 | 0 | 1.1 | 40 | 3575 | 25 | 0.69 | C:3575[S:3498,D:77],F:25,M:40,n:3640 | NCBI | multiple assembly methods | + | + | + | + | + | + |
| LR15569 | Ingroup | Cypriniformes | Cyprinoidei | Gobionidae | Hemibarbus | maculatus | 59.23 | 2156 | 0.33 | 12 | 22.14 | 806 | 0.14 | 5 | 18.16 | 661 | 2168 | 811 | 22.28 | C:2168[S:2156,D:12],F:811,M:661,n:3640 | This study | abyss | + | + | + | + | + | + |
| LR15570 | Ingroup | Cypriniformes | Cyprinoidei | Gobionidae | Ladislavia | taczanowskii | 38.02 | 1384 | 0.27 | 10 | 26.57 | 967 | 0.36 | 13 | 34.78 | 1266 | 1394 | 980 | 26.93 | C:1394[S:1384,D:10],F:980,M:1266,n:3640 | This study | spades | + | + | + | + | + | + |
| LR15062 | Ingroup | Cypriniformes | Cyprinoidei | Gobionidae | Microphysogobio | tafangensis | 45.58 | 1659 | 0.22 | 8 | 24.64 | 897 | 0.25 | 9 | 29.31 | 1067 | 1667 | 906 | 24.89 | C:1667[S:1659,D:8],F:906,M:1067,n:3640 | This study | spades | + | + | + | + | + | + |
| GCA_018749465.1 | Ingroup | Cypriniformes | Cyprinoidei | Gobionidae | Paracanthobrama | guichenoti | 93.19 | 3392 | 5.14 | 187 | 0.77 | 28 | 0 | 0 | 0.91 | 33 | 3579 | 28 | 0.77 | C:3579[S:3392,D:187],F:28,M:33,n:3640 | NCBI | multiple assembly methods | + | + | + | + | + | + |
| GCA_024679245.1 | Ingroup | Cypriniformes | Cyprinoidei | Gobionidae | Pseudorasbora | parva | 98.68 | 3592 | 0.49 | 18 | 0.41 | 15 | 0 | 0 | 0.41 | 15 | 3610 | 15 | 0.41 | C:3610[S:3592,D:18],F:15,M:15,n:3640 | NCBI | multiple assembly methods | + | + | + | + | + | + |
| GCA_023566225.1 | Ingroup | Cypriniformes | Cyprinoidei | Gobionidae | Romanogobio | albipinnatus | 96.48 | 3512 | 0.85 | 31 | 0.49 | 18 | 0 | 0 | 2.17 | 79 | 3543 | 18 | 0.49 | C:3543[S:3512,D:31],F:18,M:79,n:3640 | NCBI | multiple assembly methods | + | - | + | + | + | + |
| LR15571 | Ingroup | Cypriniformes | Cyprinoidei | Gobionidae | Sarcocheilichthys | hainanensis | 41.02 | 1493 | 0.33 | 12 | 27.28 | 993 | 0.49 | 18 | 30.88 | 1124 | 1505 | 1011 | 27.77 | C:1505[S:1493,D:12],F:1011,M:1124,n:3640 | This study | spades | + | + | + | + | + | + |
| LR15579 | Ingroup | Cypriniformes | Cyprinoidei | Gobionidae | Squalidus | sp | 42.47 | 1546 | 0.36 | 13 | 26.21 | 954 | 0.27 | 10 | 30.69 | 1117 | 1559 | 964 | 26.48 | C:1559[S:1546,D:13],F:964,M:1117,n:3640 | This study | spades | + | + | + | + | + | + |
| LR15106 | Ingroup | Cypriniformes | Cyprinoidei | Leptobarbidae | Leptobarbus | sp | 62.86 | 2288 | 0.74 | 27 | 19.97 | 727 | 0.11 | 4 | 16.32 | 594 | 2315 | 731 | 20.08 | C:2315[S:2288,D:27],F:731,M:594,n:3640 | This study | abyss | + | + | + | + | + | + |
| GCA_963993115.1 | Ingroup | Cypriniformes | Cyprinoidei | Leuciscidae | Abramis | brama | 98.13 | 3572 | 0.88 | 32 | 0.44 | 16 | 0 | 0 | 0.55 | 20 | 3604 | 16 | 0.44 | C:3604[S:3572,D:32],F:16,M:20,n:3640 | NCBI | multiple assembly methods | + | + | + | + | + | + |
| 2180_2012 | Ingroup | Cypriniformes | Cyprinoidei | Leuciscidae | Alburnoides | bipunctatus | 41.87 | 1524 | 0.22 | 8 | 24.84 | 904 | 0.36 | 13 | 32.72 | 1191 | 1532 | 917 | 25.2 | C:1532[S:1524,D:8],F:917,M:1191,n:3640 | This study | spades | + | + | + | + | + | + |
| 108LR15742 | Ingroup | Cypriniformes | Cyprinoidei | Leuciscidae | Chrosomus | erythrogaster | 40.19 | 1463 | 0.3 | 11 | 25.69 | 935 | 0.36 | 13 | 33.46 | 1218 | 1474 | 948 | 26.05 | C:1474[S:1463,D:11],F:948,M:1218,n:3640 | This study | spades | + | + | + | + | + | + |
| GCA_027497735.1 | Ingroup | Cypriniformes | Cyprinoidei | Leuciscidae | Cyprinella | lepida | 95.19 | 3465 | 0.25 | 9 | 2.31 | 84 | 0 | 0 | 2.25 | 82 | 3474 | 84 | 2.31 | C:3474[S:3465,D:9],F:84,M:82,n:3640 | NCBI | multiple assembly methods | + | + | + | + | + | + |
| GCA_027497675.1 | Ingroup | Cypriniformes | Cyprinoidei | Leuciscidae | Cyprinella | lutrensis | 86.87 | 3162 | 0.58 | 21 | 6.79 | 247 | 0.05 | 2 | 5.71 | 208 | 3183 | 249 | 6.84 | C:3183[S:3162,D:21],F:249,M:208,n:3640 | NCBI | multiple assembly methods | + | + | + | + | + | + |
| GCA_027361055.1 | Ingroup | Cypriniformes | Cyprinoidei | Leuciscidae | Cyprinella | venusta | 95.05 | 3460 | 0.22 | 8 | 2.58 | 94 | 0.03 | 1 | 2.12 | 77 | 3468 | 95 | 2.61 | C:3468[S:3460,D:8],F:95,M:77,n:3640 | NCBI | multiple assembly methods | + | + | + | + | + | + |
| LR15252 | Ingroup | Cypriniformes | Cyprinoidei | Leuciscidae | Dionda | flavipinnis | 65.96 | 2401 | 0.33 | 12 | 17.77 | 647 | 0.25 | 9 | 15.69 | 571 | 2413 | 656 | 18.02 | C:2413[S:2401,D:12],F:656,M:571,n:3640 | This study | abyss | + | + | + | + | + | + |
| 110LR15746 | Ingroup | Cypriniformes | Cyprinoidei | Leuciscidae | Gila | pandora | 49.62 | 1806 | 0.25 | 9 | 23.24 | 846 | 0.3 | 11 | 26.59 | 968 | 1815 | 857 | 23.54 | C:1815[S:1806,D:9],F:857,M:968,n:3640 | This study | spades | + | + | + | + | + | + |
| GCA_030378365.1 | Ingroup | Cypriniformes | Cyprinoidei | Leuciscidae | Hybognathus | amarus | 92.25 | 3358 | 1.62 | 59 | 2.47 | 90 | 0.03 | 1 | 3.63 | 132 | 3417 | 91 | 2.5 | C:3417[S:3358,D:59],F:91,M:132,n:3640 | NCBI | multiple assembly methods | + | + | + | + | + | + |
| GCA_023718395.1 | Ingroup | Cypriniformes | Cyprinoidei | Leuciscidae | Leucaspius | delineatus | 97.75 | 3558 | 0.74 | 27 | 0.52 | 19 | 0 | 0 | 0.99 | 36 | 3585 | 19 | 0.52 | C:3585[S:3558,D:27],F:19,M:36,n:3640 | NCBI | multiple assembly methods | + | + | + | + | + | + |

| | | | | | | | | | | | | | | | | | | | | | | | | | | | | |
|---|---|---|---|---|---|---|---|---|---|---|---|---|---|---|---|---|---|---|---|---|---|---|---|---|---|---|---|---|
| GCA_021554675.1 | Ingroup | Cypriniformes | Cyprinoidei | Leuciscidae | Leuciscus | idus | 98.1 | 3571 | 0.77 | 28 | 0.44 | 16 | 0 | 0 | 0.69 | 25 | 3599 | 16 | 0.44 | C:3599[S:3571,D:28],F:16,M:25,n:3640 | NCBI | multiple assembly methods | + | + | + | + | + | + |
| 2223_2013 | Ingroup | Cypriniformes | Cyprinoidei | Leuciscidae | Leuciscus | leuciscus | 41.13 | 1497 | 0.22 | 8 | 26.43 | 962 | 0.52 | 19 | 31.7 | 1154 | 1505 | 981 | 26.95 | C:1505[S:1497,D:8],F:981,M:1154,n:3640 | This study | spades | + | + | + | + | + | + |
| GCA_030578275.1 | Ingroup | Cypriniformes | Cyprinoidei | Leuciscidae | Meda | fulgida | 98.49 | 3585 | 0.41 | 15 | 0.6 | 22 | 0 | 0 | 0.49 | 18 | 3600 | 22 | 0.6 | C:3600[S:3585,D:15],F:22,M:18,n:3640 | NCBI | multiple assembly methods | + | + | + | + | + | + |
| LR15254 | Ingroup | Cypriniformes | Cyprinoidei | Leuciscidae | Notropis | chalybaeus | 28.93 | 1053 | 0.16 | 6 | 26.29 | 957 | 0.38 | 14 | 44.23 | 1610 | 1059 | 971 | 26.67 | C:1059[S:1053,D:6],F:971,M:1610,n:3640 | This study | spades | + | + | + | + | + | + |
| GCA_949152265.1 | Ingroup | Cypriniformes | Cyprinoidei | Leuciscidae | Phoxinus | phoxinus | 98.46 | 3584 | 0.41 | 15 | 0.47 | 17 | 0 | 0 | 0.66 | 24 | 3599 | 17 | 0.47 | C:3599[S:3584,D:15],F:17,M:24,n:3640 | NCBI | multiple assembly methods | + | + | + | + | + | + |
| GCF_016745375.1 | Ingroup | Cypriniformes | Cyprinoidei | Leuciscidae | Pimephales | promelas | 94.75 | 3449 | 3.43 | 125 | 1.04 | 38 | 0 | 0 | 0.77 | 28 | 3574 | 38 | 1.04 | C:3574[S:3449,D:125],F:38,M:28,n:3640 | NCBI | multiple assembly methods | + | + | + | + | + | + |
| 109LR15744 | Ingroup | Cypriniformes | Cyprinoidei | Leuciscidae | Ptychocheilus | lucius | 53.35 | 1942 | 0.27 | 10 | 22.64 | 824 | 0.47 | 17 | 23.27 | 847 | 1952 | 841 | 23.11 | C:1952[S:1942,D:10],F:841,M:847,n:3640 | This study | spades | + | + | + | + | + | + |
| LR15251 | Ingroup | Cypriniformes | Cyprinoidei | Leuciscidae | Rhinichthys | cataractae | 45.19 | 1645 | 0.25 | 9 | 25.49 | 928 | 0.47 | 17 | 28.6 | 1041 | 1654 | 945 | 25.96 | C:1654[S:1645,D:9],F:945,M:1041,n:3640 | This study | spades | + | + | + | + | + | + |
| LR15572 | Ingroup | Cypriniformes | Cyprinoidei | Leuciscidae | Rhynchocypris | lagowskii | 36.9 | 1343 | 0.27 | 10 | 25.52 | 929 | 0.41 | 15 | 36.9 | 1343 | 1353 | 944 | 25.93 | C:1353[S:1343,D:10],F:944,M:1343,n:3640 | This study | spades | + | + | + | + | + | + |
| GCA_951802725.1 | Ingroup | Cypriniformes | Cyprinoidei | Leuciscidae | Rutilus | rutilus | 98.43 | 3583 | 0.49 | 18 | 0.55 | 20 | 0 | 0 | 0.52 | 19 | 3601 | 20 | 0.55 | C:3601[S:3583,D:18],F:20,M:19,n:3640 | NCBI | multiple assembly methods | + | + | + | + | + | + |
| GCA_024453875.1 | Ingroup | Cypriniformes | Cyprinoidei | Leuciscidae | Scardinius | erythrophthalmus | 97.45 | 3547 | 0.91 | 33 | 0.55 | 20 | 0 | 0 | 1.1 | 40 | 3580 | 20 | 0.55 | C:3580[S:3547,D:33],F:20,M:40,n:3640 | NCBI | multiple assembly methods | + | + | + | + | + | + |
| GCA_031834385.1 | Ingroup | Cypriniformes | Cyprinoidei | Leuciscidae | Semotilus | atromaculatus | 98.24 | 3576 | 1.15 | 42 | 0.41 | 15 | 0 | 0 | 0.19 | 7 | 3618 | 15 | 0.41 | C:3618[S:3576,D:42],F:15,M:7,n:3640 | NCBI | multiple assembly methods | + | + | + | + | + | + |
| GCA_949319135.1 | Ingroup | Cypriniformes | Cyprinoidei | Leuciscidae | Squalius | cephalus | 98.71 | 3593 | 0.55 | 20 | 0.52 | 19 | 0 | 0 | 0.22 | 8 | 3613 | 19 | 0.52 | C:3613[S:3593,D:20],F:19,M:8,n:3640 | NCBI | multiple assembly methods | + | + | + | + | + | + |
| 1450_2013 | Ingroup | Cypriniformes | Cyprinoidei | Leuciscidae | Telestes | muticellus | 38.76 | 1411 | 0.25 | 9 | 26.29 | 957 | 0.3 | 11 | 34.4 | 1252 | 1420 | 968 | 26.59 | C:1420[S:1411,D:9],F:968,M:1252,n:3640 | This study | spades | + | + | + | + | + | + |
| GCA_030578255.1 | Ingroup | Cypriniformes | Cyprinoidei | Leuciscidae | Tiaroga | cobitis | 97.01 | 3531 | 2.53 | 92 | 0.41 | 15 | 0 | 0 | 0.05 | 2 | 3623 | 15 | 0.41 | C:3623[S:3531,D:92],F:15,M:2,n:3640 | NCBI | multiple assembly methods | + | + | + | + | + | + |
| GCA_022828995.1 | Ingroup | Cypriniformes | Cyprinoidei | Leuciscidae | Vimba | vimba | 98.35 | 3580 | 0.71 | 26 | 0.38 | 14 | 0 | 0 | 0.55 | 20 | 3606 | 14 | 0.38 | C:3606[S:3580,D:26],F:14,M:20,n:3640 | NCBI | multiple assembly methods | + | - | + | + | + | + |
| LR12004 | Ingroup | Cypriniformes | Cyprinoidei | Paedocyprididae | Paedocypris | carbunculus | 57.64 | 2098 | 0.88 | 32 | 16.98 | 618 | 0.22 | 8 | 24.29 | 884 | 2130 | 626 | 17.2 | C:2130[S:2098,D:32],F:626,M:884,n:3640 | NCBI-SRA | abyss | + | + | + | + | + | + |
| GCA_028031935.1 | Ingroup | Cypriniformes | Cyprinoidei | Paedocyprididae | Paedocypris | micromegethes1 | 93.63 | 3408 | 0.74 | 27 | 1.29 | 47 | 0 | 0 | 4.34 | 158 | 3435 | 47 | 1.29 | C:3435[S:3408,D:27],F:47,M:158,n:3640 | NCBI | multiple assembly methods | + | + | + | + | + | + |
| LR7898 | Ingroup | Cypriniformes | Cyprinoidei | Paedocyprididae | Paedocypris | micromegethes2 | 71.54 | 2604 | 0.47 | 17 | 13.08 | 476 | 0.14 | 5 | 14.78 | 538 | 2621 | 481 | 13.22 | C:2621[S:2604,D:17],F:481,M:538,n:3640 | NCBI-SRA | spades | + | + | + | + | + | + |
| LR15325 | Ingroup | Cypriniformes | Cyprinoidei | Paedocyprididae | Paedocypris | sp2 | 72.5 | 2639 | 1.35 | 49 | 10.99 | 400 | 0.08 | 3 | 15.08 | 549 | 2688 | 403 | 11.07 | C:2688[S:2639,D:49],F:403,M:549,n:3640 | This study | abyss | + | + | + | + | + | + |
| LR15863A | Ingroup | Cypriniformes | Cyprinoidei | Paedocyprididae | Paedocypris | sp1 | 93.52 | 3404 | 1.07 | 39 | 1.1 | 40 | 0.03 | 1 | 4.29 | 156 | 3443 | 41 | 1.13 | C:3443[S:3404,D:39],F:41,M:156,n:3640 | This study | hifiasm.primary.HiC.scaffolded | + | + | + | + | + | + |
| LR09467 | Ingroup | Cypriniformes | Cyprinoidei | Psilorhynchidae | Psilorhynchus | sp1 | 87.23 | 3175 | 0.52 | 19 | 7.2 | 262 | 0.08 | 3 | 4.97 | 181 | 3194 | 265 | 7.28 | C:3194[S:3175,D:19],F:265,M:181,n:3640 | This study | spades | + | + | + | + | + | + |
| LR15089 | Ingroup | Cypriniformes | Cyprinoidei | Psilorhynchidae | Psilorhynchus | sp2 | 81.46 | 2965 | 0.33 | 12 | 11.15 | 406 | 0.11 | 4 | 6.95 | 253 | 2977 | 410 | 11.26 | C:2977[S:2965,D:12],F:410,M:253,n:3640 | This study | spades | + | + | + | + | + | + |
| 100LR08982 | Ingroup | Cypriniformes | Cyprinoidei | Sundadanionidae | Fangfangia | spinicleithralis | 55.58 | 2023 | 0.41 | 15 | 22.31 | 812 | 0.16 | 6 | 21.54 | 784 | 2038 | 818 | 22.47 | C:2038[S:2023,D:15],F:818,M:784,n:3640 | This study | spades | + | + | + | + | + | + |
| LR15870 | Ingroup | Cypriniformes | Cyprinoidei | Sundadanionidae | Sundadanio | atomus | 97.12 | 3535 | 0.85 | 31 | 0.66 | 24 | 0 | 0 | 1.37 | 50 | 3566 | 24 | 0.66 | C:3566[S:3535,D:31],F:24,M:50,n:3640 | This study | hifiasm.primary.HiC.scaffolded | + | + | + | + | + | + |
| LR10372 | Ingroup | Cypriniformes | Cyprinoidei | Sundadanionidae | Sundadanio | goblinus | 40.33 | 1468 | 0.27 | 10 | 25.99 | 946 | 0.25 | 9 | 33.16 | 1207 | 1478 | 955 | 26.24 | C:1478[S:1468,D:10],F:955,M:1207,n:3640 | This study | spades | + | + | + | + | + | + |
| LR15082 | Ingroup | Cypriniformes | Cyprinoidei | Sundadanionidae | Sundadanio | margarition | 42.88 | 1561 | 0.38 | 14 | 26.04 | 948 | 0.27 | 10 | 30.41 | 1107 | 1575 | 958 | 26.31 | C:1575[S:1561,D:14],F:958,M:1107,n:3640 | This study | spades | + | + | + | + | + | + |
| 101LR12751 | Ingroup | Cypriniformes | Cyprinoidei | Sundadanionidae | Sundadanio | retiarius | 44.23 | 1610 | 0.41 | 15 | 25.22 | 918 | 0.38 | 14 | 29.75 | 1083 | 1625 | 932 | 25.6 | C:1625[S:1610,D:15],F:932,M:1083,n:3640 | This study | spades | + | + | + | + | + | + |
| LR15110 | Ingroup | Cypriniformes | Cyprinoidei | Tanichthyidae | Tanichthys | albonudes | 24.15 | 879 | 0.14 | 5 | 26.57 | 967 | 0.49 | 18 | 48.65 | 1771 | 884 | 985 | 27.06 | C:884[S:879,D:5],F:985,M:1771,n:3640 | This study | spades | + | + | + | + | + | + |
| LR02462 | Ingroup | Cypriniformes | Cyprinoidei | Tanichthyidae | Tanichthys | micagemmae | 27.86 | 1014 | 0.08 | 3 | 26.79 | 975 | 0.33 | 12 | 44.95 | 1636 | 1017 | 987 | 27.12 | C:1017[S:1014,D:3],F:987,M:1636,n:3640 | This study | spades | + | + | + | + | + | + |
| 1545_2010 | Ingroup | Cypriniformes | Cyprinoidei | Tincidae | Tinca | tinca | 46.26 | 1684 | 0.25 | 9 | 25.55 | 930 | 0.14 | 5 | 27.8 | 1012 | 1693 | 935 | 25.69 | C:1693[S:1684,D:9],F:935,M:1012,n:3640 | This study | spades | + | + | + | + | + | + |
| GCA_003731715.1 | Ingroup | Cypriniformes | Cyprinoidei | Xenocyprididae | Anabarilius | grahami | 98.13 | 3572 | 0.44 | 16 | 1.04 | 38 | 0 | 0 | 0.38 | 14 | 3588 | 38 | 1.04 | C:3588[S:3572,D:16],F:38,M:14,n:3640 | NCBI | multiple assembly methods | + | + | + | + | + | + |
| 118LR15719 | Ingroup | Cypriniformes | Cyprinoidei | Xenocyprididae | Candidia | pingtungensis | 64.84 | 2360 | 0.49 | 18 | 20.58 | 749 | 0.25 | 9 | 13.85 | 504 | 2378 | 758 | 20.83 | C:2378[S:2360,D:18],F:758,M:504,n:3640 | This study | spades | + | + | + | + | + | + |
| GCA_024489055.1 | Ingroup | Cypriniformes | Cyprinoidei | Xenocyprididae | Chanodichthys | erythropterus | 98.82 | 3597 | 0.6 | 22 | 0.33 | 12 | 0 | 0 | 0.25 | 9 | 3619 | 12 | 0.33 | C:3619[S:3597,D:22],F:12,M:9,n:3640 | NCBI | multiple assembly methods | + | + | + | + | + | + |
| GCF_019924925.1 | Ingroup | Cypriniformes | Cyprinoidei | Xenocyprididae | Ctenopharyngodon | idella | 96.4 | 3509 | 2.64 | 96 | 0.3 | 11 | 0 | 0 | 0.66 | 24 | 3605 | 11 | 0.3 | C:3605[S:3509,D:96],F:11,M:24,n:3640 | NCBI | multiple assembly methods | + | + | + | + | + | + |
| GCA_028476615.1 | Ingroup | Cypriniformes | Cyprinoidei | Xenocyprididae | Culter | alburnus | 97.23 | 3539 | 1.21 | 44 | 1.07 | 39 | 0 | 0 | 0.49 | 18 | 3583 | 39 | 1.07 | C:3583[S:3539,D:44],F:39,M:18,n:3640 | NCBI | multiple assembly methods | + | + | + | + | + | + |
| GCA_037101425.1 | Ingroup | Cypriniformes | Cyprinoidei | Xenocyprididae | Elopichthys | bambusa | 99.04 | 3605 | 0.49 | 18 | 0.38 | 14 | 0 | 0 | 0.08 | 3 | 3623 | 14 | 0.38 | C:3623[S:3605,D:18],F:14,M:3,n:3640 | NCBI | multiple assembly methods | + | - | + | + | + | + |
| 103LR15622 | Ingroup | Cypriniformes | Cyprinoidei | Xenocyprididae | Hemiculter | sp | 35.25 | 1283 | 0.11 | 4 | 27.45 | 999 | 0.41 | 15 | 36.79 | 1339 | 1287 | 1014 | 27.86 | C:1287[S:1283,D:4],F:1014,M:1339,n:3640 | This study | spades | + | + | + | + | + | + |
| GCA_004764525.1 | Ingroup | Cypriniformes | Cyprinoidei | Xenocyprididae | Hypophthalmichthys | molitrix | 90.55 | 3296 | 0.36 | 13 | 4.78 | 174 | 0.05 | 2 | 4.26 | 155 | 3309 | 176 | 4.83 | C:3309[S:3296,D:13],F:176,M:155,n:3640 | NCBI | multiple assembly methods | + | + | + | + | + | + |
| GCA_019925145.1 | Ingroup | Cypriniformes | Cyprinoidei | Xenocyprididae | Hypophthalmichthys | nobilis | 98.27 | 3577 | 0.41 | 15 | 0.47 | 17 | 0 | 0 | 0.85 | 31 | 3592 | 17 | 0.47 | C:3592[S:3577,D:15],F:17,M:31,n:3640 | NCBI | multiple assembly methods | + | + | + | + | + | + |
| GCF_018812025.1 | Ingroup | Cypriniformes | Cyprinoidei | Xenocyprididae | Megalobrama | amblycephala | 98.19 | 3574 | 1.02 | 37 | 0.63 | 23 | 0 | 0 | 0.16 | 6 | 3611 | 23 | 0.63 | C:3611[S:3574,D:37],F:23,M:6,n:3640 | NCBI | multiple assembly methods | + | + | + | + | + | + |
| 102LR15621 | Ingroup | Cypriniformes | Cyprinoidei | Xenocyprididae | Opsariichthys | bidens | 63.08 | 2296 | 0.44 | 16 | 21.32 | 776 | 0.27 | 10 | 14.89 | 542 | 2312 | 786 | 21.59 | C:2312[S:2296,D:16],F:786,M:542,n:3640 | This study | spades | + | + | + | + | + | + |
| 128LR15724 | Ingroup | Cypriniformes | Cyprinoidei | Xenocyprididae | Opsariichthys | pachycephalus | 48.43 | 1763 | 0.33 | 12 | 24.89 | 906 | 0.52 | 19 | 25.82 | 940 | 1775 | 925 | 25.41 | C:1775[S:1763,D:12],F:925,M:940,n:3640 | This study | spades | + | + | + | + | + | + |
| LR15574 | Ingroup | Cypriniformes | Cyprinoidei | Xenocyprididae | Oxygaster | pointoni | 63.02 | 2294 | 0.49 | 18 | 18.85 | 686 | 0.19 | 7 | 17.45 | 635 | 2312 | 693 | 19.04 | C:2312[S:2294,D:18],F:693,M:635,n:3640 | This study | spades | + | + | + | + | + | + |
| GCA_034642465.1 | Ingroup | Cypriniformes | Cyprinoidei | Xenocyprididae | Zacco | platypus | 94.07 | 3424 | 3.63 | 132 | 0.44 | 16 | 0 | 0 | 1.87 | 68 | 3556 | 16 | 0.44 | C:3556[S:3424,D:132],F:16,M:68,n:3640 | NCBI | multiple assembly methods | + | + | + | + | + | + |
| LR15097 | Ingroup | Cypriniformes | Gyrinocheiloidei | Gyrinocheilidae | Gyrinocheilus | pustulosus | 91.4 | 3327 | 0.55 | 20 | 5.47 | 199 | 0.05 | 2 | 2.53 | 92 | 3347 | 201 | 5.52 | C:3347[S:3327,D:20],F:201,M:92,n:3640 | This study | spades | + | + | + | + | + | + |
| 104EA0138 | Ingroup | Cypriniformes | Gyrinocheiloidei | Gyrinocheilidae | Gyrinocheilus | sp | 92.83 | 3379 | 0.71 | 26 | 4.07 | 148 | 0.05 | 2 | 2.34 | 85 | 3405 | 150 | 4.12 | C:3405[S:3379,D:26],F:150,M:85,n:3640 | This study | abyss | + | + | + | + | + | + |
| GCA_034509495.1 | Outgroup | Characiformes | | Acestrorhynchidae | Acestrorhynchus | altus | 94.86 | 3453 | 3.16 | 115 | 0.99 | 36 | 0 | 0 | 0.99 | 36 | 3568 | 36 | 0.99 | C:3568[S:3453,D:115],F:36,M:36,n:3640 | NCBI | multiple assembly methods | + | + | + | + | + | + |
| GCF_023375975.1 | Outgroup | Characiformes | | Characidae | Astyanax | mexicanus | 99.07 | 3606 | 0.49 | 18 | 0.27 | 10 | 0 | 0 | 0.16 | 6 | 3624 | 10 | 0.27 | C:3624[S:3606,D:18],F:10,M:6,n:3640 | NCBI | multiple assembly methods | + | + | + | + | + | + |
| GCA_032718565.1 | Outgroup | Characiformes | | Distichodontidae | Distichodus | sexfasciatus | 91.92 | 3346 | 0.93 | 34 | 3.08 | 112 | 0.05 | 2 | 4.01 | 146 | 3380 | 114 | 3.13 | C:3380[S:3346,D:34],F:114,M:146,n:3640 | NCBI | multiple assembly methods | + | + | + | + | + | + |
| GCA_017165825.1 | Outgroup | Characiformes | | Hepsetidae | Hepsetus | odoe | 96.35 | 3507 | 1.37 | 50 | 1.59 | 58 | 0.03 | 1 | 0.66 | 24 | 3557 | 59 | 1.62 | C:3557[S:3507,D:50],F:59,M:24,n:3640 | NCBI | multiple assembly methods | + | + | + | + | + | + |
| GCA_034552555.1 | Outgroup | Characiformes | | Prochilodontidae | Semaprochilodus | insignis | 81.02 | 2949 | 1.18 | 43 | 8.98 | 327 | 0.05 | 2 | 8.76 | 319 | 2992 | 329 | 9.03 | C:2992[S:2949,D:43],F:329,M:319,n:3640 | NCBI | multiple assembly methods | + | + | + | + | + | + |
| GCF_900700375.1 | Outgroup | Clupeiformes | | Denticipitidae | Denticeps | clupeoides | 89.95 | 3274 | 7.64 | 278 | 0.71 | 26 | 0 | 0 | 1.7 | 62 | 3552 | 26 | 0.71 | C:3552[S:3274,D:278],F:26,M:62,n:3640 | NCBI | multiple assembly methods | + | + | + | + | + | + |
| GCA_027475355.1 | Outgroup | Clupeiformes | | Engraulidae | Coilia | nasus | 95.49 | 3476 | 0.8 | 29 | 0.85 | 31 | 0 | 0 | 2.86 | 104 | 3505 | 31 | 0.85 | C:3505[S:3476,D:29],F:31,M:104,n:3640 | NCBI | multiple assembly methods | + | + | + | + | + | + |
| GCF_902362185.1 | Outgroup | Gonorynchiformes | | Chanidae | Chanos | chanos | 98.79 | 3596 | 0.44 | 16 | 0.3 | 11 | 0 | 0 | 0.47 | 17 | 3612 | 11 | 0.3 | C:3612[S:3596,D:16],F:11,M:17,n:3640 | NCBI | multiple assembly methods | + | + | + | + | + | + |
| GCA_028565775.1 | Outgroup | Gymnotiformes | | Apteronotidae | Sternarchorhynchus | goeldii | 94.78 | 3450 | 0.41 | 15 | 2.72 | 99 | 0.08 | 3 | 2.01 | 73 | 3465 | 102 | 2.8 | C:3465[S:3450,D:15],F:102,M:73,n:3640 | NCBI | multiple assembly methods | + | + | + | + | + | + |
| GCA_025802075.1 | Outgroup | Gymnotiformes | | Apteronotidae | Tenebrosternarchus | preto | 94.51 | 3440 | 0.36 | 13 | 3.35 | 122 | 0 | 0 | 1.79 | 65 | 3453 | 122 | 3.35 | C:3453[S:3440,D:13],F:122,M:65,n:3640 | NCBI | multiple assembly methods | + | + | + | + | + | + |
| GCA_020368025.1 | Outgroup | Gymnotiformes | | Hypopomidae | Brachyhypopomus | occidentalis | 97.86 | 3562 | 0.22 | 8 | 0.44 | 16 | 0 | 0 | 1.48 | 54 | 3570 | 16 | 0.44 | C:3570[S:3562,D:8],F:16,M:54,n:3640 | NCBI | multiple assembly methods | + | + | + | + | + | + |
| GCF_030014155.1 | Outgroup | Siluriformes | | Bagridae | Tachysurus | vachellii | 98.54 | 3587 | 0.52 | 19 | 0.33 | 12 | 0 | 0 | 0.6 | 22 | 3606 | 12 | 0.33 | C:3606[S:3587,D:19],F:12,M:22,n:3640 | NCBI | multiple assembly methods | + | + | + | + | + | + |
| GCA_030522415.1 | Outgroup | Siluriformes | | Clariidae | Channallabes | apus | 97.86 | 3562 | 0.74 | 27 | 0.38 | 14 | 0 | 0 | 1.02 | 37 | 3589 | 14 | 0.38 | C:3589[S:3562,D:27],F:14,M:37,n:3640 | NCBI | multiple assembly methods | + | + | + | + | + | + |
| GCA_030347435.1 | Outgroup | Siluriformes | | Clariidae | Clarias | fuscus | 97.83 | 3561 | 0.93 | 34 | 0.47 | 17 | 0 | 0 | 0.77 | 28 | 3595 | 17 | 0.47 | C:3595[S:3561,D:34],F:17,M:28,n:3640 | NCBI | multiple assembly methods | + | + | + | + | + | + |
| GCA_023630585.1 | Outgroup | Siluriformes | | Cranoglanididae | Cranoglanis | bouderius | 98.41 | 3582 | 0.74 | 27 | 0.33 | 12 | 0 | 0 | 0.52 | 19 | 3609 | 12 | 0.33 | C:3609[S:3582,D:27],F:12,M:19,n:3640 | NCBI | multiple assembly methods | + | + | + | + | + | + |
| GCF_023375685.1 | Outgroup | Siluriformes | | Ictaluridae | Ictalurus | furcatus | 97.14 | 3536 | 0.77 | 28 | 0.47 | 17 | 0 | 0 | 1.62 | 59 | 3564 | 17 | 0.47 | C:3564[S:3536,D:28],F:17,M:59,n:3640 | NCBI | multiple assembly methods | + | - | + | + | + | + |
| GCA_025084515.1 | Outgroup | Siluriformes | | Loricariidae | Ancistrus | triradiatus | 88.54 | 3223 | 0.74 | 27 | 4.37 | 159 | 0.03 | 1 | 6.32 | 230 | 3250 | 160 | 4.4 | C:3250[S:3223,D:27],F:160,M:230,n:3640 | NCBI | multiple assembly methods | + | + | + | + | + | + |
| GCA_024760905.1 | Outgroup | Siluriformes | | Plotosidae | Plotosus | lineatus | 97.66 | 3555 | 0.58 | 21 | 0.6 | 22 | 0 | 0 | 1.15 | 42 | 3576 | 22 | 0.6 | C:3576[S:3555,D:21],F:22,M:42,n:3640 | NCBI | multiple assembly methods | + | + | + | + | + | + |
| GCF_014805685.1 | Outgroup | Siluriformes | | Siluridae | Silurus | meridionalis | 96.73 | 3521 | 0.66 | 24 | 0.47 | 17 | 0 | 0 | 2.14 | 78 | 3545 | 17 | 0.47 | C:3545[S:3521,D:24],F:17,M:78,n:3640 | NCBI | multiple assembly methods | + | + | + | + | + | + |

**Supplementary Table 4.** Summary of datasets used in the phylogenomic analyses, showing the number of taxa, retained loci, and alignment lengths.

| Dataset | Number.of.taxa | Number.of.loci | Sequence.Type | Alignment.length | Median.alignement.l | Median.Missing.data | Median.Occupancy | Genomic.region | Concatenated.phylogeny.IQTREE2 | loci.trees.IQTREE2 | Coalescent.Species.Tree.ASTRAL4 | Coalescent.Species.Tree.wASTRAL | Coalescent.Species.Tree.CASTER.SITE | Coalescent.Species.Tree.CASTER.PAIR | loci.trees.RAXML-NG | Coalescent.Species.Tree.ASTRAL-Pro3 | Gene.wise.Phylogenetic.signal | Gene.genealogy.interrogation |
|---|---|---|---|---|---|---|---|---|---|---|---|---|---|---|---|---|---|---|
| BUSCO.NT.C123 | 320 | 539 | nucleotide | 428220 | 678 | 0.013 | 0.857 | exons | + | + | + | + | + | + | - | - | - | - |
| BUSCO.NT.C12 | 320 | 539 | nucleotide | 258496 | 442 | 0.013 | 0.863 | exons | + | + | + | + | + | + | - | - | + | + |
| BUSCO.AA | 320 | 539 | amino acid | 137323 | 226 | 0.014 | 0.857 | exons | + | + | + | + | + | + | - | - | - | - |
| HUGHES.NT.C123 | 330 | 607 | nucleotide | 184734 | 246 | 0.004 | 0.997 | exons | + | + | + | + | + | + | - | - | - | - |
| HUGHES.NT.C12 | 330 | 607 | nucleotide | 127622 | 166 | 0.004 | 0.997 | exons | + | + | + | + | + | + | - | - | + | + |
| HUGHES.AA | 330 | 607 | amino acid | 68469 | 87 | 0.005 | 0.997 | exons | + | + | + | + | + | + | - | - | - | - |
| MALMSTRØM.NT.C123 | 335 | 976 | nucleotide | 194961 | 192 | 0.006 | 0.777 | exons | + | + | + | + | + | + | - | - | - | - |
| MALMSTRØM.NT.C12 | 335 | 976 | nucleotide | 133914 | 128 | 0.007 | 0.778 | exons | + | + | + | + | + | + | - | - | + | + |
| MALMSTRØM.AA | 335 | 976 | amino acid | 66776 | 64 | 0.007 | 0.778 | exons | + | + | + | + | + | + | - | - | - | - |
| UCE | 335 | 628 | nucleotide | 283487 | 442 | 0.022 | 0.851 | UCE | + | + | + | + | + | + | - | - | + | + |
| BUSCO.NT.C12+UCE | 335 | 1167 | nucleotide | 541983 | | | | exons+UCE | + | - | + | - | - | - | - | - | - | - |
| ROADIES-Rep1 | 335 | 1337 | nucleotide | 898040 | | | | Genome-wide | - | - | - | - | - | - | + | + | - | - |
| ROADIES-Rep2 | 335 | 2662 | nucleotide | 1769786 | | | | Genome-wide | - | - | - | - | - | - | + | + | - | - |
| ROADIES-Rep3 | 335 | 1352 | nucleotide | 903840 | | | | Genome-wide | - | - | - | - | - | - | + | + | - | - |

**Supplementary Table 5.** Different GHOST models tested for the combined first and second codon positions of BUSCO.NT.C12 and UCE nucleotide dataset with 140 species.

| GHOST_Model | Log-likelihood_tree | No.free.parameters | AIC_score | AICc_score | BIC_score |
|---|---|---|---|---|---|
| GTR+H2 | -8213283.764 | 563 | 16427693.5 | 16427694.7 | 16434000.8 |
| GTR*H2 | -8301487.355 | 568 | 16604110.7 | 16604111.9 | 16610474 |
| GTR+H4 | -8131034.58 | 1119 | 16264307.2 | 16264311.79 | 16276843.3 |
| GTR*H4 | -8243833.087 | 1134 | 16489934.2 | 16489938.93 | 16502638.4 |
| GTR+H6 | -8121220.464 | 1675 | 16245790.9 | 16245801.32 | 16264555.9 |
| GTR*H6 | -8200123.786 | 1700 | 16403647.6 | 16403658.28 | 16422692.7 |
| GTR+FO+H4 | -8130804.651 | 1119 | 16263847.3 | 16263851.94 | 16276383.4 |
| GTR+FO*H4 | -8533385.138 | 1143 | 17069056.3 | 17069061.11 | 17081861.3 |
| GTR+FO*H2 | -8349368.899 | 571 | 16699879.8 | 16699881 | 16706276.7 |
| GTR+FO*H6 | -8243762.204 | 1715 | 16490954.4 | 16490965.3 | 16510167.5 |

**Supplementary Table 6.** Relationship between log-transformed branch lengths and gene concordance factors (gCF) and site concordance factors (sCF).

| Dataset | Response | Predictor | Pearson correlation | t-statistics | Estimate (Slope) | Std. Error | Intercept | R square | Adjusted R square | p-value |
|---|---|---|---|---|---|---|---|---|---|---|
| BUSCO.NT.C123 | gCF | log transofrmed branch length | 0.8185489 | 25.291 | 18.645 | 0.7372 | 159.5109 | 0.67 | 0.669 | 2.20E-16 |
| BUSCO.NT.C123 | sCF | log transofrmed branch length | 0.452357 | 9.0023 | 7.5782 | 0.8418 | 89.1503 | 0.2046 | 0.2021 | 2.20E-16 |
| BUSCO.NT.C12 | gCF | log transofrmed branch length | 0.8510681 | 28.768 | 18.5819 | 0.6459 | 155.0679 | 0.7243 | 0.7234 | 2.20E-16 |
| BUSCO.NT.C12 | sCF | log transofrmed branch length | 0.4558947 | 9.091 | 8.2425 | 0.9067 | 99.4576 | 0.2078 | 0.2053 | 2.20E-16 |
| BUSCO.AA | gCF | log transofrmed branch length | 0.8432779 | 27.846 | 18.5851 | 0.6674 | 141.3713 | 0.7111 | 0.7102 | 2.20E-16 |
| BUSCO.AA | sCF | log transofrmed branch length | 0.4922061 | 10.036 | 9.1143 | 0.9082 | 99.8097 | 0.2423 | 0.2399 | 2.20E-16 |
| HUGHES.NT.C123 | gCF | log transofrmed branch length | 0.7980242 | 23.873 | 18.4187 | 0.7715 | 142.1073 | 0.6368 | 0.6357 | 2.20E-16 |
| HUGHES.NT.C123 | sCF | log transofrmed branch length | 0.4170132 | 8.2713 | 6.7812 | 0.8198 | 85.241 | 0.1739 | 0.1714 | 3.45E-15 |
| HUGHES.NT.C12 | gCF | log transofrmed branch length | 0.7580946 | 20.957 | 11.3306 | 0.5407 | 95.617 | 0.5747 | 0.5734 | 2.20E-16 |
| HUGHES.NT.C12 | sCF | log transofrmed branch length | 0.4686588 | 9.5643 | 7.7537 | 0.8107 | 99.2514 | 0.2196 | 0.2172 | 2.20E-16 |
| HUGHES.AA | gCF | log transofrmed branch length | 0.7610123 | 21.148 | 11.4855 | 0.5431 | 88.3046 | 0.5791 | 0.5778 | 2.20E-16 |
| HUGHES.AA | sCF | log transofrmed branch length | 0.5079148 | 10.63 | 8.6558 | 0.8143 | 101.4236 | 0.258 | 0.2557 | 2.20E-16 |
| MALMSTRØM.NT.C123 | gCF | log transofrmed branch length | 0.7760932 | 22.357 | 17.0229 | 0.7614 | 129.5982 | 0.6023 | 0.6011 | 2.20E-16 |
| MALMSTRØM.NT.C123 | sCF | log transofrmed branch length | 0.3738255 | 7.3217 | 6.0016 | 0.8197 | 80.533 | 0.1397 | 0.1371 | 1.89E-12 |
| MALMSTRØM.NT.C12 | gCF | log transofrmed branch length | 0.684516 | 17.057 | 7.8149 | 0.4582 | 67.9817 | 0.4686 | 0.467 | 2.20E-16 |
| MALMSTRØM.NT.C12 | sCF | log transofrmed branch length | 0.4306826 | 8.6689 | 6.736 | 0.777 | 93.103 | 0.1855 | 0.183 | 2.20E-16 |
| MALMSTRØM.AA | gCF | log transofrmed branch length | 0.6908903 | 17.36 | 7.6085 | 0.4383 | 60.1054 | 0.4773 | 0.4757 | 2.20E-16 |
| MALMSTRØM.AA | sCF | log transofrmed branch length | 0.5204207 | 11.071 | 8.6437 | 0.7807 | 100.4202 | 0.2708 | 0.2686 | 2.20E-16 |
| UCE | gCF | log transofrmed branch length | 0.7752588 | 22.229 | 13.4982 | 0.6072 | 129.0194 | 0.601 | 0.5998 | 2.20E-16 |
| UCE | sCF | log transofrmed branch length | 0.4735845 | 9.7679 | 6.8075 | 0.6969 | 91.3928 | 0.2243 | 0.2219 | 2.20E-16 |
